# C-6 Modified 2-F-Fucose Derivatives as Inhibitors of Fucosyl Transferases

**DOI:** 10.1101/2025.08.18.670827

**Authors:** Yanyan Liu, Geert-Jan Boons

## Abstract

Fluorinated analogs of guanosine-diphosphate-β-L-fucose (GDP-Fuc) have received considerable attention for the development of inhibitors of fucosyltransferases (FUTs). These compounds can be recognized by FUTs but do not or slowly transfer the fluorinated fucosyl residue because the electron-withdrawing fluorine(s) destabilize the oxocarbenium-like transition state. Fluorinated GDP-Fuc analogs can also act as feedback inhibitor of the *de novo* biosynthesis pathway of GDP-Fuc. To investigate the biological significance of distinct glycoconjugate classes, it is important to develop inhibitors that can selectively target specific FUT enzymes. Here, we report the design, synthesis, and biological evaluation of a series of GDP-2-F-Fuc analogs modified at C-6 of Fuc by various amides and ethers. We also prepared and examined corresponding prodrugs as potential FUT inhibitors of cellular glycosylation. Our findings reveal that two of the inhibitors potently inhibited FUT1, 3, 6, and 9, while displaying minimal activity against FUT8. However, the corresponding prodrugs did not inhibit cellular fucosylation, which is probably due to a lack of GDP-fucose pyrophosphorylase activity. The results demonstrate that modifications at the C-6 position of Fuc can confer selectivity, although further investigations of alternative functional groups are required to enhance cellular tolerance and efficacy.

## Introduction

Fucosylation of glycans and glycoconjugates is crucial for numerous biological processes, including cell adhesion, tissue development, angiogenesis, immunity, malignancy, and tumor metastasis.^[1–3]^ Fucosyltransferases (Futs) are enzymes that catalyze the transfer of L-fucose (Fuc) from guanosine-diphosphate-β-L-fucose (GDP-Fuc) to diverse sugar acceptors, including oligosaccharides, glycoproteins, and glycolipids.^[4]^ In humans, thirteen distinct FUT genes have been identified, categorized into subtypes based on the glycosidic linkages they form. These include α1,2-fucosyltransferases (FUT1 and FUT2), α1,3/4-fucosyltransferases (FUT3-7 and FUT9), α1,6-fucosyltransferases (FUT8), and *O*-fucosyltransferases (O-FUTs), each exhibiting unique substrate specificities and sites of action. For instance, α1,2- and α1,3/4-fucosylations typically occur as terminal or sub-terminal modifications, while α1,6-fucosylation occurs at the core of *N*-glycans. Fucosylation by POFUT1 and POFUT2 involves the attachment of a fucoside to specific serine and threonine residues within protein domains such as epidermal growth factor (EGF)-like and thrombospondin repeating units.

The development of inhibitors of fucosylation as therapeutic agents for various diseases has received attention.^[5–10]^ Several fluorinated GDP-fucose analogs have been reported, including GDP-2-F-Fuc, GDP-2,2-di-F-Fuc, GDP-3-F-Fuc, GDP-6-F-Fuc, GDP-6,6-di-F-Fuc, and GDP-6,6,6-tri-F-Fuc.^[8,11–16]^ These compounds can be recognized by FUTs but do not or slowly transfer the fucosyl residue because the electron-withdrawing fluorine(s) destabilize the oxocarbenium-like transition state. Fluorinated fucose analogs or their acetylated derivatives can be taken up by cells, and several derivatives can be converted by the salvage pathway into the corresponding GDP derivatives, thereby inhibiting cellular fucosylation. In addition, fluorinated fucose-1-phosphate prodrugs have been described to by-pass generating intracellularly the corresponding GDP derivatives.^[7,13,17–18]^ Fluorinated GDP-Fuc analogs can also act as feedback inhibitors of the *de novo* biosynthesis pathway of GDP-Fuc^[7,17]^, thereby further diminishing fucosylation of glycoconjugates of cells.

Despite these advances, fluorinated GDP-Fuc analogs still have modest potencies. Furthermore, there is a need to develop inhibitors that target specific fucosyl transferases. To address these challenges, we explored whether further modification of GDP-Fuc-2-F at the C-6 position of fucose can provide more potent and selective inhibitors (compounds **2a**-**h**; Figure 1). It was expected that the C-6 modification of these compounds would confer selectivity, whereas C-2 fluorination would ensure inhibitory activity. There are indications that FUTs tolerate modification at the C-6 position. A GDP-Fuc analog having a C-6 hydroxyl at fucose (GDP-L-galactose) can readily be transferred by α1,3-fucosyltransferases, leading to the formation of a dimeric SLe^x^ undecasaccharide.^[19]^ Also, 6-deoxy-6-*N*-(2-naphthylacetyl)-GDP-fucose is an excellent donor for FUT6, with a K_m_ of 0.94 μM, and can also be accepted by

**Figure 1.**
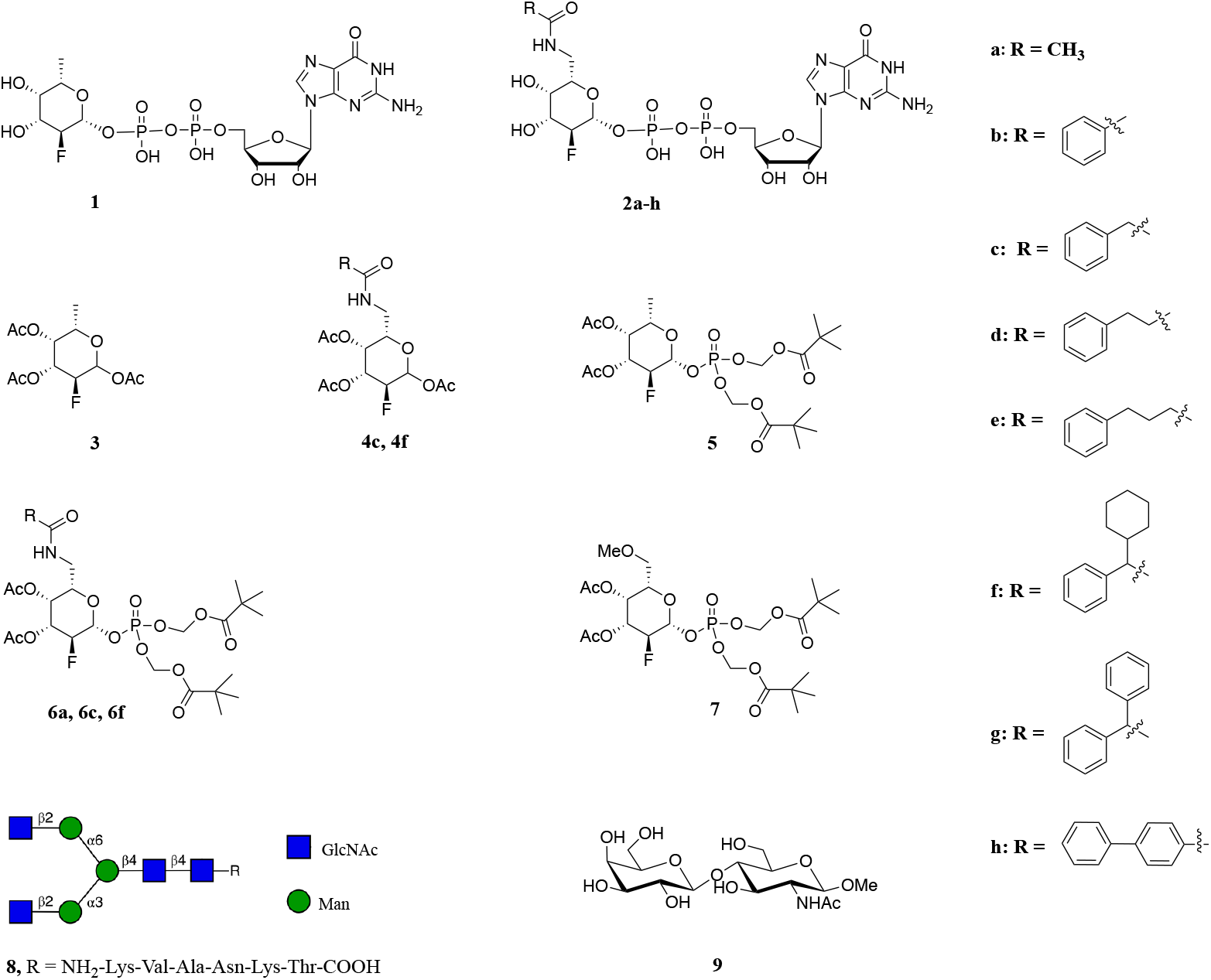
Structures of target compounds. Compounds **1** and **2a**-**h** are GDP-fucose analogs as potential FUT inhibitors. Compounds **3**-**7** are prodrug forms intended for cellular assays. Compounds **8** and **9** serve as acceptor substrates in enzymatic assays for determining inhibitory constants.

FUT8 with a K_m_ of 175 μM, facilitating the direct monitoring of α1,6-fucosylation at the reducing-terminal GlcNAc residue.^[20]^ Similarly, 7-alkynyl-fucose is efficiently converted by the biosynthetic machinery of cells into its GDP-activated form and can be incorporated into *N*-glycans, allowing for the sensitive detection of fucosylated proteins in living cells.^[21]^ Peracetylated 6-alkynyl and 6-alkenyl fucose analogs can intracellularly be converted to their GDP-Fuc forms and incorporated into Notch EGF repeating units by PoFUT1.^[14]^ A fluorescently labeled derivative, 6-*N*-(ATTO 550-acetamide)-GDP-Fuc, which incorporates the cationic dye ATTO 550, is recognized by FUT9, allowing for real-time monitoring of enzyme activity *via* fluorescence cross-correlation spectroscopy.^[22]^ Other 6-N-fluorescent tagged GDP-Fuc derivates can also been accepted by FUTs.^[23]^ High-throughput screening has identifies several 6-substituted GDP-Fuc analogs that can inhibit FUTs inhibitors.^[6,24]^

Based on these observations, we designed and synthesized eight 2-F-GDP-Fuc analogs (compounds **2a**-**h**; Figure 1) featuring *N*-substituents at C-6 linked *via* an amide, ranging from methyl to phenyl with varying alkyl chains and branched modifications. It was expected that the C-2 modification would confer inhibitory activity, whereas the C-6 substituent would provide selectivity. We also prepared the corresponding prodrugs, including the per-acetyl-fucose versions (compounds **3, 4c**, and **4f**; Figure 1) and fucose-1-phosphate derivatives (compounds **5, 6a, 6c**, and **6f**; Figure 1), having the phosphate masked as bis(pivaloyloxymethyl) (POM) carbonate. The latter functionality is used as a phosphate-masking moiety in the development of membrane-permeable prodrugs. The POM carbonate is cleaved intracellularly by esterases to yield a carboxylate intermediate that spontaneously degrades to release a phosphate monoester. It is employed in several FDA-approved antiviral agents such as adefovir dipivoxil.^[25]^ We have employed the POM masking group for effective cellular utility of 2,2-di-F-Fuc-1-phosphate.^[16]^ Similarly, a POM-based prodrug approach also improved the performance of 6-CF_3_-Fuc-1-phosphate in cell-based assays.^[13]^ It bypasses the necessity for anomeric phosphorylation by L-fucose kinase, which is sensitive to chemical modifications. Compound **1** served as a control. The inhibitory effects of the GDP-Fuc analogs have been evaluated against six different human FUTs. We also examined the ability of prodrugs **3**-**6** to inhibit cell surface fucosylation.

## Results and Discussion

### Synthesis and Evaluation of 6-Modified-2-F-GDP-Fuc as Inhibitors of FUTs

The preparation of GDP-2-F-Fuc (**1)** was performed by optimization of a reported method as outlined in Scheme S1.^[26]^ The synthesis of compounds **2a**-**h** focuses on two key aspects: the introduction of *N*-substituents at the C-6 position and the incorporation of fluorine at the C-2 position. A common method for amino substitution of hydroxyl groups involves initially converting the hydroxyl group into a leaving group,^[27–28]^ which can then be replaced by an azido group,^[22,29]^ which can subsequently be reduced to an amino group.^[30]^ The introduction of the fluorine atom requires the formation of a double bond between C1 and C2,^[31–32]^ which serves as the reactive site for electrophilic fluorination by Selectfluor. This step was performed before azido installation due to the instability of azido-containing intermediates (Figure 2). We chose tosyl (Ts) as the leaving group, which is sufficiently stable that it can be retained until the step just before coupling with GMP, at which point it is replaced by azide. This approach allows for a common synthetic route for compounds **2a**-**h**, reducing the overall workload.

**Figure 2.**
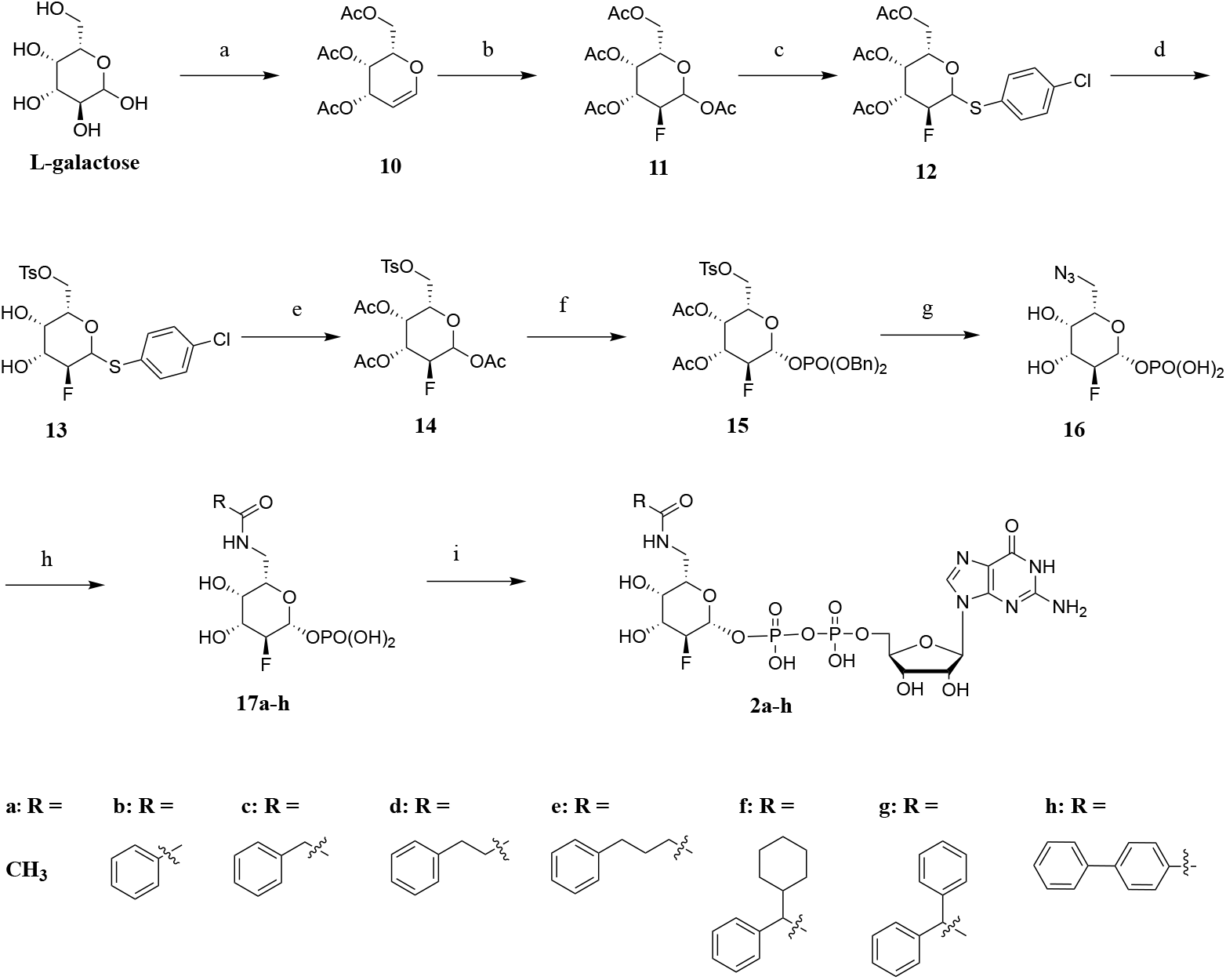
Synthesis of GDP-fucose analogs with modifications at C6 and a fluorine at C2. a) i. Acetic anhydride, pyridine, perchloric acid, HBr/AcOH; ii. EA/saturated NaH_2_PO_4_ (1:2), zinc dust, 87% over two steps. b) i. Selectfluor, nitromethane, H_2_O, microwave; ii. acetic anhydride, pyridine, 65% over two steps. c) i. HBr/AcOH; ii. tetrabutylammonium, hydrogen sulfate, *p*-chlorothiophenol, chloroform, 1 M NaOH, 56% over two steps. d) i. NaOMe, MeOH; ii. 4-toluenesulfonyl chloride, pyridine, 68% over two steps. e) i. acetic anhydride, pyridine; ii. *N*-bromosuccinimide, acetone, H_2_O; iii. acetic anhydride, pyridine, 83% over 3 steps. f) i. HBr/AcOH; ii. silver carbonate, dibenzyl phosphate, 3 Å MS, CH_3_CN, 84% over two steps. g) i. H_2_, Pd/C, MeOH; ii. MeOH: H_2_O: Et_3_N = 3:7:1; iii. NaN_3_, DMF, 70 °C, 81% over 3 steps. h) i. *t*-BuOH, 10 mM NH_4_HCO_3_, Pd/C, H_2_; ii. Corresponding NHS ester, DIPEA, THF, H_2_O, 50-70% over two steps. i) GMP-morpholidate, 1*H*-tetrazole, pyridine, 25-45%

This synthesis began with commercially available L-galactose, which was acetylated using acetic anhydride in the presence of perchloric acid as the catalyst. The anomeric acetyl ester was replaced by bromide by treatment with HBr in acetic acid, resulting in the formation of acetobromo-L-galactose. The galactosyl bromide was then subjected to reductive elimination by being treated with zinc dust, generating the desired product **10** in a yield of 87%.^[33]^ The fluorination of compound **10** was achieved by a two-step process, beginning with electrophilic fluorination utilizing Selectfluor in a nitromethane-water mixture. This reaction was followed by the acetylation of the resulting anomeric hydroxyl group, leading to the formation of compound **11** in an overall yield of 65%. The next step involved the selective introduction of a leaving group at the C-6 position. To achieve this goal, it was essential to first block the reactive anomeric hydroxyl. This was achieved by initially converting **11** into the corresponding glycosyl bromide by treatment with HBr in acetic acid, followed by substitution of the bromide by *p*-chlorothiophenol in the presence of tetrabutylammonium hydrogen sulfate, resulting in the formation of *p*-chlorophenol thioglycoside **12**, which was isolated in a yield of 56%.^[26]^ After deacetylation of **12** using standard conditions, the C-6 hydroxyl was selectively tosylated by reaction with the *p*-toluenesulfonyl chloride in pyridine, resulting in the formation of **13** (68% yield).^[34]^ Subsequently, the thioglycoside of **13** was cleaved by treatment with *N*-bromosuccinimide (NBS) in a mixture of acetone and H_2_O, and the resulting anomeric hydroxyl was acetylated using acetic anhydride in pyridine, generating full-protected compound **14** (83% yield). An anomeric phosphate was installed by a two-step procedure involving treatment of **14** with HBr in acetic acid to generate the corresponding anomeric bromide, which was activated with silver carbonate and subjected to nucleophilic substitution with dibenzyl phosphate. This S_N_2 displacement resulted in the exclusive formation of **15** as the β-anomer in a yield of 84%.^[35]^ The β-configuration was confirmed by the coupling constant between H1 and F (*J*_H1-F_ = 7.7 Hz). The next step involved deprotection^[36–37]^ of compound **15**, followed by azide substitution to afford **16**.^[38]^ Catalytic hydrogenation over Pd/C was employed to remove the benzyl protecting groups on the phosphate, followed by the deacetylation of the C3 and C4 esters utilizing triethylamine. Subsequently, treatment with sodium azide in DMF facilitated the displacement of the tosylate by azide, resulting in the formation of 6-azido-2-F-fucose-1-phosphate (**16**) in 81% yield. Compound **16** serves as the intermediate for the synthesis of the targeted inhibitors. Thus, the azide was reduced to a free amine by hydrogenation over Pd/C and then reacted with a range of NHS-activated esters^[39–41]^ to introduce the desired modification at C-6.^[42]^ The final step involved the coupling of the anomeric phosphate with GMP-morpholidate, resulting in the generation of all desired GDP-fucose analogues **2a**-**h** in yields ranging from 25 to 45%. The rather low yields for these compounds can be attributed to two factors: first, the reaction is inherently slow, and even with prolonged reaction times, residual starting material remains. Second, the negatively charged nature of sugar nucleotides complicates purification, often resulting in loss of material. The preparation of acceptors **8** and **9** is described in the supporting information. (Figure S1 and Scheme S3).

We also attempted to synthesize inhibitors **2a**-**h** using enzymatic methods. FKP (Bacterial L-fucokinase/GDP-fucose pyrophosphorylase) is a bifunctional enzyme that facilitates the conversion of fucose to Fuc-1-P, which is then transformed into GDP-Fuc. FKP demonstrates broad substrate specificity and efficiently acts on various L-fucose derivatives, such as 2-F-Fuc, 6-N_3_-Fuc, and some other 6-modified-Fuc derivatives.^[17,43]^. However, when we combined C-2 and C-6 modifications, we discovered that FKP was unable to catalyze the conversion of either 6-N_3_-2-F-Fuc or 6-N_3_-2-F-Fuc-1-P (compound **16**) into the corresponding sugar nucleotide product. We further tested compounds **17a**-**h** as substrates for FKP, but these compounds were also not converted by this enzyme.

Following the synthesis of all target inhibitors, we conducted experiments to evaluate their inhibitory activity for human FUTs. Using a commercially available GDP-Glo Glycosyltransferase assay kit, we assessed the kinetic parameters of FUT1, 3, 6, 8, and 9 in the presence of inhibitors **2a**-**h**. The GDP-Glo assay provides a convenient approach for detecting GDP production during glycosyltransferase reactions.^[44–46]^ In these experiments, GDP-Fuc served as the donor substrate for all FUTs. Compound **9** was used as the acceptor for FUT1, 3, 5, 6, and 9, while **8** was used for FUT8. The summarized results are outlined in Table 1.

**Table 1.** Summary of K_i_ values for compounds **2a**-**h** against FUT1, 3, 6, 8, and 9.

| Ki/μM | FUT 8 | FUT6 | FUT1 | FUT3 | FUT9 |
| --- | --- | --- | --- | --- | --- |
| **1** | 111.8±8.5 | 26.3±2.5 | 35.9±10.4 | 17.9±1.9 | 27.8±1.7 |
| **2a** | 68.6±7.2 | 13.5±2.2 | 5.4±0.9 | 9.9±1.8 | 21.5±1.6 |
| **2b** | 106.4±13.1 | 12.9±1.5 | 7.1 ±4.3 | 16.3±1.3 | 28.1±3.7 |
| **2c** | 208.5±20.9 | 4.3±0.2 | 4.1±0.9 | 6.2±0.9 | 10.6±1.3 |
| **2d** | 80.7±9.0 | 3.7±0.2 | 2.2±0.4 | 8.8±0.9 | 7.8±4.5 |
| **2e** | 57.1±9.5 | 9.7±2.9* | 1.5±0.2 | 2.1±0.4 | 14.2±1.3 |
| **2f** | 518.7±72.1 | 3.4±0.2 | 11.4±3.3 | 8.8±1.1 | 5.4±0.8 |
| **2g** | 172.6±50.6 | 8.0±1.2* | 1.3±0.4 | 1.3±0.2 | 3.8±0.2 |
| **2h** | 273.0±47.1 | 10.4±1.3 | 6.1±1.6 | 28.8±2.3 | 629.8±233.7 |

\* These values were analyzed using a non-competitive inhibition model. All other values were analyzed with the competitive inhibition model.

All the new inhibitors exhibit inhibitory activity for the employed FUTs. Specifically, inhibitors **2a**-**h** substantially increased inhibitory activity for human FUT1, 3, 6, and 9 compared to **1**. This finding indicates the potential of these compounds as FUT inhibitors. Notably, inhibitors **2a** and **2e** showed greater inhibition of FUT8 compared to **1**, however, they also exhibited strong inhibition of other FUTs. Conversely, inhibitors **2c** and **2f** displayed some selectivity, with K_i_ values for FUT8 of 208 and 518 μM, respectively, and K_i_ values for FUT1, 3, 6, and 9 ranging from 3 to 11 μM, substantially lower than those for FUT8. Both inhibitors **2c** and **2f** share a phenethyl structure, suggesting that the spacing of a single carbon atom between the phenyl group and the amide bond critically influences recognition by FUT8. When a flexible alkyl ring is attached to this carbon (as in inhibitor **2f**), the inhibitory effect is markedly reduced, whereas a rigid phenyl ring (as in inhibitor **2g**) does not produce this reduction. When the carbon chain between the phenyl group and the amide bond is extended to two (inhibitor **2d**) or three carbons (inhibitor **2e**), inhibitory effects are increased, indicating that longer phenyl alkyl chains can effectively bind to FUT8.

### Synthesis and Evaluation of Prodrugs for Cell Surface Fucosylation

Based on enzymatic kinetics testing, two representative compounds were selected for prodrug development. Prodrugs **4c, 4f, 6c**, and **6f** were synthesized based on inhibitors **2c** and **2f**, as these show the highest selectivity between FUT8 and other FUTs. Additionally, prodrug **6a** was synthesized to evaluate whether intracellular metabolic pathways are compatible with the amide bond at the C6 position. To investigate the influence of bond type at position C-6, we also designed and synthesized prodrug **7**. Prodrug **6a** consists of a 2-F-fucose containing compound linked to a methyl group *via* an amide, whereas prodrug **7** features a 2-F-fucose moiety connected to a methyl group *via* an ether. The amide is more rigid and can act as both a hydrogen bond donor and acceptor,^[47–48]^ while the ether can only function as a hydrogen bond acceptor.^[49–50]^

First, prodrug **5** was synthesized based on compound **3**. Various methods^[25,51–54]^ for introducing Di-POM moieties were explored, and it was determined that bromination of per-acetylated sugar, followed by phosphorylation, was most efficient with the fewest reaction steps. DiPOM-phosphate was synthesized *via* a three-step published procedure (Scheme S2).^[55]^ Prodrug **5** was synthesized from compound **3** by an anomeric phosphorylation using anomeric bromide as the electrophile and DiPOM-phosphate as the nucleophile, affording **5** in 65% yield.

Compounds **4a, 4c, 4f, 6a, 6c**, and **6f** were prepared starting from **13** (Figure 3A). The tosyl of **13** was substituted with azide by treatment with sodium azide, resulting in the formation of **18** in a 70% yield. The azide of **18** was then reduced to a free amine group using triphenylphosphine, and the resulting amine was reacted with appropriate NHS esters to introduce the planned modifications at the C-6 position. Subsequent acetylation of the C-3 and C4-hydroxyls with acetic anhydride, followed by hydrolysis of the thioglycoside by treatment with *N*-bromosuccinimide and acetylation of the resulting alcohol, resulted in desired compounds **4a, 4c**, and **4f**. Prodrugs **6a, 6c**, and **6f** were synthesized by a phosphorylation reaction between **4** and DiPOM-phosphate.

**Figure 3.**
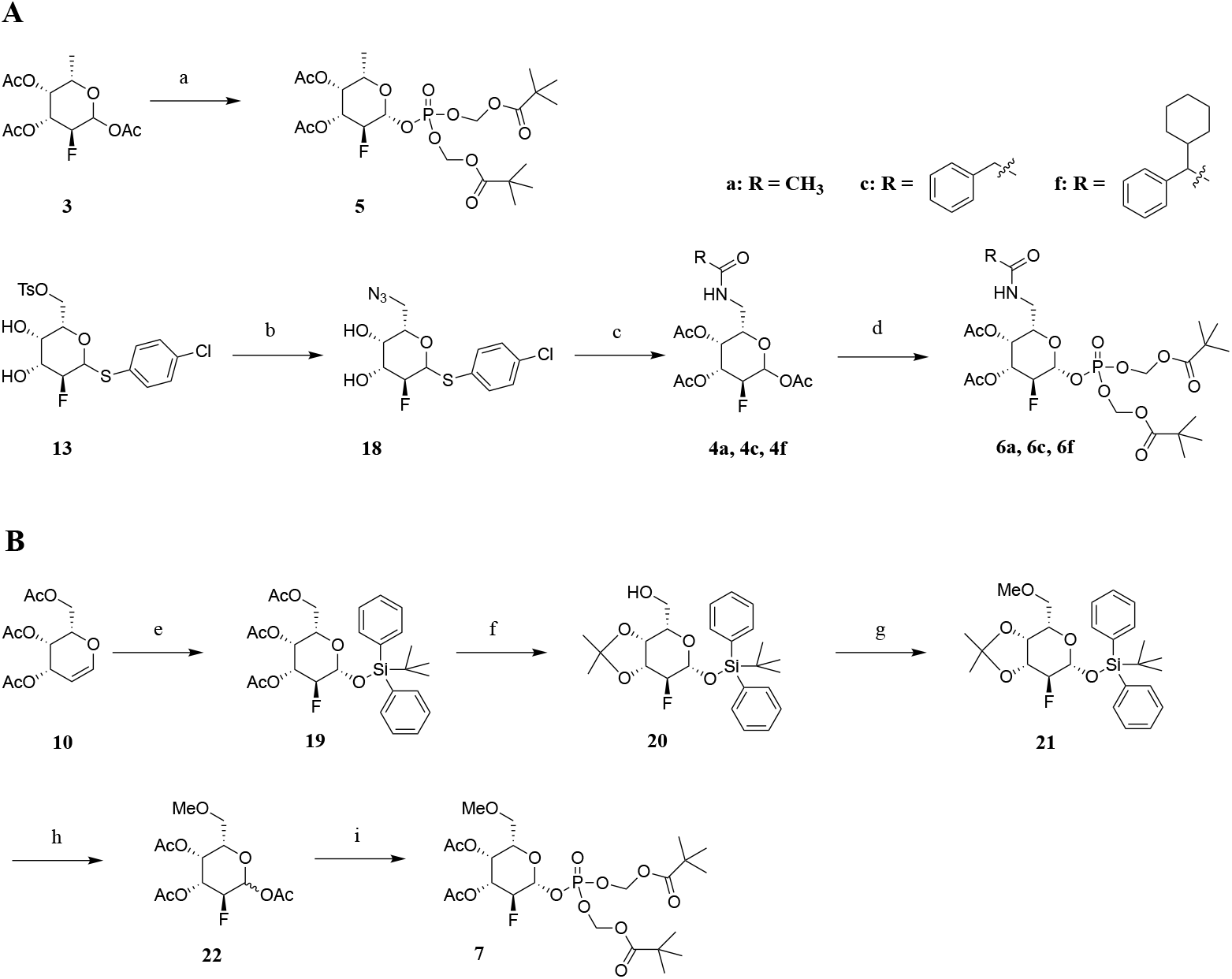
A) Synthesis of 6-N-2-F-prodrugs. a) i. HBr/AcOH; ii. silver carbonate, freshly prepared diPOM-phosphate, 3Å MS, CH_3_CN, 65% over two steps. b) NaN_3_, DMF, 70 °C, 70%. c) i. riphenylphosphine, THF; ii. H_2_O, DIPEA, corresponding active NHS ester; iii. acetic anhydride, pyridine; iv. *N*-bromosuccinimide, acetone, H_2_O; v. acetic anhydride, pyridine, 50-60% over 5 steps. d) i. HBr/AcOH, ii. silver carbonate, freshly prepared diPOM-phosphate, 3Å MS, CH_3_CN, 40-50% over two steps. B) Synthesis of prodrug **7**. e) i. Selectfluor, nitromethane, H_2_O, microwave, 100°C; ii. Tert-butyl diphenyl-silyl chloride, imidazole, DMF, 45% over 2 steps. f) i. sodium methoxide, methanol, ii. *p*-toluenesulfonic acid, acetone, 41% over 2 steps. g) Sodium hydride, iodomethane, DMF, 83%. h) i. 90% TFA; ii. tetrabutylammonium fluoride, THF, acetate acid, iii. pyridine, acetic anhydride, 95% over 3 steps. i) i. HBr/AcOH, ii. silver carbonate, DiPOM-phosphate, 3 Å MS, CH_3_CN, 65% over two steps.

The synthesis of **7** followed a similar strategy as for the corresponding amide by initially introducing fluorine at the C-2 position, followed by the installation of the desired ether bond at C-6. The synthetic route began with the glycal **10** (Figure 3B). Electrophilic fluorination was achieved by treatment with Selectfluor, resulting in the formation of a lactol that was protected as a tert-*t*-butyl diphenyl silyl group (TBDPS)^[56]^ by reaction with *t*-butyl diphenyl silyl chloride, forming **19** in a yield of 45%. To install the methyl ester at the position C-6, compound **19** was deacetylated using standard conditions, and the resulting free hydroxyls at C3 and C4 were protected as an acetonide, resulting in the formation of compound **20** in 41% yield.^[57–59]^ The methyl ether at the C6 position was introduced by treatment of **20** with iodomethane in the presence of NaH in DMF, resulting in the formation of **21** in a yield of 83%. Subsequently, the *t*-butyldiphenylsilyl group and acetonide were removed by sequential treatment with 90% trifluoroacetic acid (TFA), followed by tetrabutylammonium fluoride in THF and acetic acid. The resulting triol was then acetylated using acetic anhydride, yielding compound **22** in a yield of 95%.^[60–61]^ Finally, **22** was converted into an anomeric bromide, which was used in a phosphorylation reaction with DiPOM-phosphate in the presence of silver carbonate to afford the desired 6-OMe-2-F prodrug **7** (65% yield).

The prodrugs were evaluated for their inhibitor activity for cell surface fucosylation of HL-60 cells. Flow cytometry was employed using staining with the lectin AAL that specifically recognizes α-L-fucosides. Besides prodrugs **3** and **5**, which were used as controls, none of the newly synthesized prodrugs exhibited inhibitory effects in the cell-based experiment (Figure 4A). Several explanations may account for these results. One possibility is that these prodrugs are not converted into the corresponding GDP-fucose analogues, leading to a lack of cellular activity. Alternatively, even if the analogs were generated, the amide bond at the C6 position might interfere with the recognition by the enzyme GMDs required for feedback inhibition, preventing these prodrugs from effectively modulating intracellular GDP-fucose concentration.

**Figure 4.**
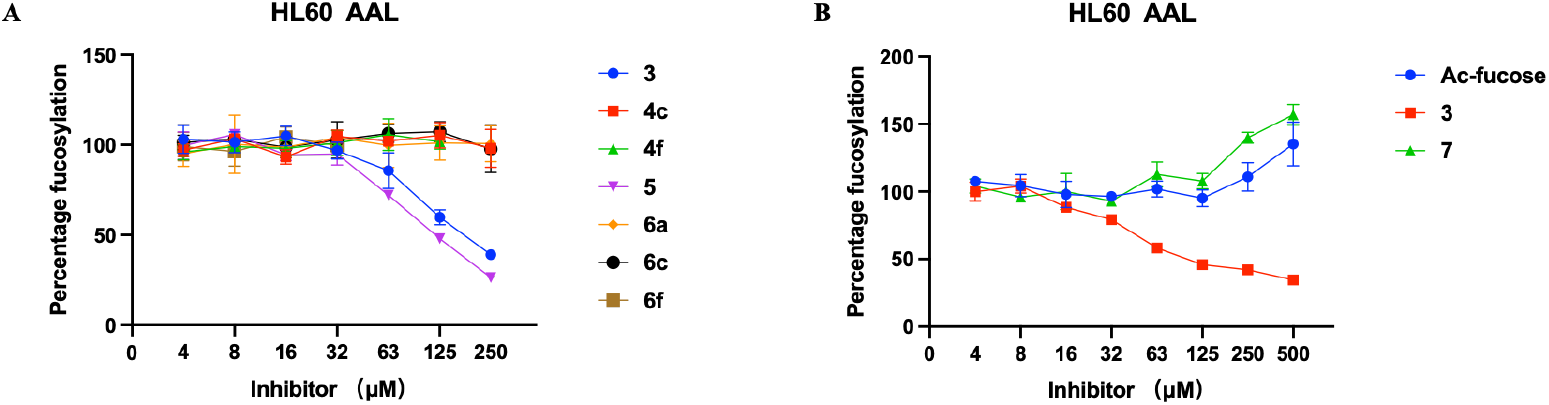
Flow cytometry results of inhibition of fucosylation on HL60 cells. Cells were treated with prodrugs ranging in concentrations from 4 to 500 μM or DMSO as control for 3 days. The data were normalized to cells treated with DMSO only as 100% fucosylation and unstained cells as 0%. The data presented are indicative of three independent experiments conducted in triplicate, illustrating the mean values ± s.d. (n=3). Error bars are displayed for each dataset; these may not be apparent in some cases due to their size being smaller than the data points marker.

Their direct inhibitory effects on the FUTs may have been insufficient to alter cell surface glycan expression. Another potential explanation is that, due to substantial structural modifications, these compounds might not exhibit selectivity for fucose-related pathways but could instead affect other metabolic functions within the cells.

Prodrug **7** was also applied to the HL60 cell line. The only difference between prodrug **6a** and **7** is the modification of C-6 by an amide and ether, respectively. Unexpectedly, while other prodrugs showed no effect on cellular fucosylation, prodrug **7** promoted cell surface fucosylation (Figure 4B). The origin of this observation needs further investigation. We also conducted experiments to assess the impact of the prodrugs on cell proliferation.^[62–63]^ We found that prodrugs **4f** and **6f** significantly affected proliferation at certain concentrations (Figure S4. The other prodrugs did not significantly impact proliferation.

## Conclusions

A series of novel FUT inhibitors (**2a**-**h**) was designed and synthesized. These compounds feature an amine at the C6 position of 2-F-fucose, with various phenyl and alkyl groups attached at C-6 through an amine bond. Enzyme kinetic analysis revealed that most inhibitors exhibited more potent activity for various human FUTs compared to the control inhibitor GDP-2-F-Fuc (**1**). Notably, **2c** and **2f** showed an increase in selectivity between FUT8 and other FUTs, with Ki values of 208 and 518 μM for FUT8, respectively. Their K_i_ values for FUT1, 3, 6, and 9 ranged from 3 to 11 μM, significantly lower than for FUT8. Both inhibitors **2c** and **2f** contain a phenethyl group, indicating that a one-carbon spacing between the phenyl ring and the amide is critical for FUT8 recognition. Introducing a flexible alkyl ring at this position (as in inhibitor **2f**) substantially reduces inhibitory potency for FUT8, while attaching a rigid phenyl ring (inhibitor **2g**) did not induce such a reduction. Extending the carbon chain between the phenyl and the amide bond to two (inhibitor **2d**) or three carbons (inhibitor **2e**) increased inhibitory activity, indicating that longer phenyl-alkyl chains enhance the binding affinity to FUT8. These results indicated that a strategy of maintaining fluorine at C2 for inhibition, combined with a modification at C6, can modulate FUTs recognition, is effective. Moreover, inhibitors **2c** and **2f** are both selective, showing a preference for inhibiting FUT6 over FUT8.

Based on these results, we developed prodrugs **4c, 4f, 6a, 6c**, and **6f** to investigate their potential to modulate cell surface fucosylation. Additionally, prodrugs **7** was synthesized to compare the effects of an amide and an ether bond at the C6 position. In cellular experiments, **4c, 4f, 6a, 6c**, and **6f** did not inhibit cell surface fucosylation. Compounds, **4f** and **6f** did, however, reeduce cell proliferation at high concentration. Prodrug **7** appeared to slightly promote cell surface fucosylation. These results indicate that while an amide at the C6 position enhances FUTs inhibition and selectivity, it is not an appropriate substrate for the cellular fucose biopathway. Future research will focus on exploring additional modifications based on this strategy to arrive at cell-based inhibitors. A possible strategy is to employ carba-fucose as a starting point to install various modifications at C-6. Carba-fucose is a possible starting point because it is a substrate of bifunctional L-fucokinase/GDP-fucose pyrophosphorylase and can therefore be converted into the corresponding GDP derivatives, which in turn is an inhibitor of various fucosyl transferases.^[9,64]^

### Experimental Section

Experimental details can be found in the Supporting Information and includes general methods, synthetic protocols and compound characterization, enzyme kinetics, cell culture, and flow cytometry. The Supporting Information (PDF) also contains supplementary figures and schemes, copies of NMR spectra, and graphics depicting K_i_ measurements.

## Acknowledgements

Dr. Kelley Moremen (University of Georgia, USA) provided the expression vectors for the various fucosyl transferases, which were expressed by Dr. Gerlof P Bosman (Utrecht University) and Mrs Linda H.C. Quarles van Ufford (Utrecht University) by reported procedures. Y.L. was supported by the Chinese Scholarship Council.

## Conflict of Interest

The authors declare no conflict of interest.

## Data Availability Statement

The data that support the findings of this study are available in the supplementary material of this article.

## Notes

### Competing Interest Statement

The authors have declared no competing interest.

