## Supplementary material for "C-6 Modified 2-F-Fucose Derivatives as Inhibitors of Fucosyl Transferases": SI

#### Table of contents

page

**1) Supplementary Schemes and Figures** S2

**Scheme S1.** Synthesis of GDP-2-F-Fuc S2

**Scheme S2.** Synthesis of compound 28 (diPOM-phosphate) S2

**Scheme S3.** Synthesis of substrate **9** S3

**Figure S1.** Preparation of substrate **8** S3

**Figure S2.** $^{1}H$ NMR spectrum of compound **8** S4

**Figure S3.** Anomeric region $^{13}C$ $^{1}H$ HSQC spectrum (assignment **8**) S4

**Figure S4**. Proliferation and viability of HL60 S5

**2) Experimental Procedures** S5

General Methods S5

Synthetic Protocols and Compound Characterization S6

Kinetic Studies S42

Cell Culture S42

Flow Cytometry S43

Cell Proliferation and Viability S43

**3) Reference** S43

**4) NMR Spectra** S43

**5) Graphs Depicting the Determination of Ki** S83

### 1) Supplementary Schemes and Figures

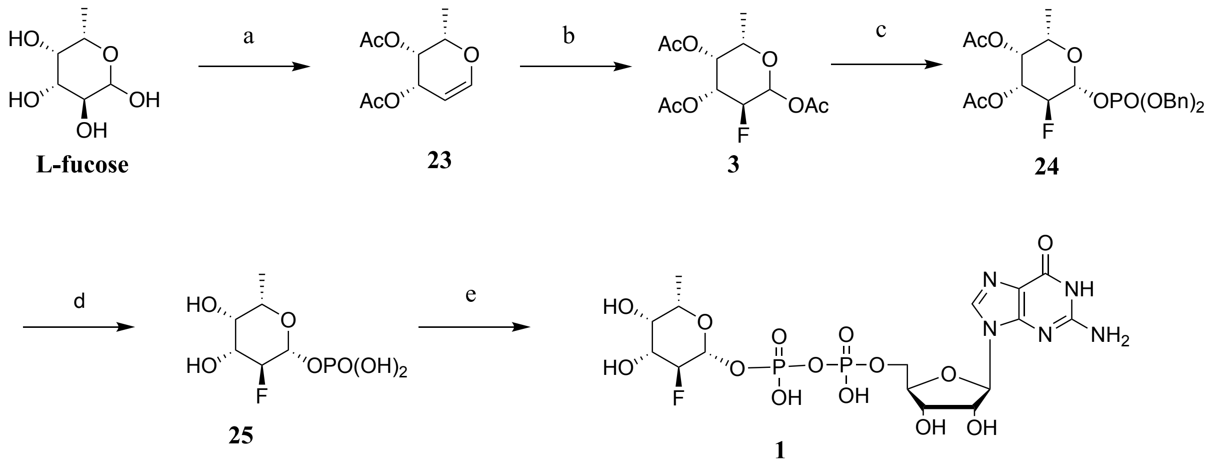


**Scheme S1.** Synthesis of GDP-2-F-Fuc. a) i. Acetic anhydride, Pyridine, Perchloric acid, HBr/AcOH, ii. EA/saturated $NaH_2PO_4$ (1:2), zinc dust, 73% over two steps. b) i. Selectfluor, DMF, $H_2O$, ii. Acetic anhydride, Pyridine, 73% over two steps. c) i. HBr/AcOH, ii. Silver carbonate, dibenzyl phosphate, 3 Å MS, $CH_3CN$, 70% over two steps. d) i. Pd/C, $H_2$, MeOH, ii. MeOH: H2O: Et3N = 3:7:1, 88% over two steps. e) GMP-morpholidate, 1H-Tetrazole, pyridine, 18%.

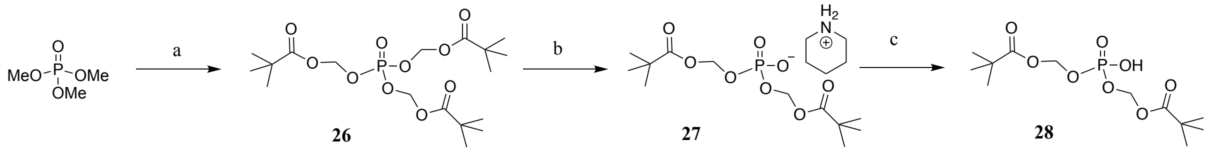


**Scheme S2.** Synthesis of bis(pivaloyloxymethyl) (POM) phosphate. a) Sodium iodide, chloromethyl pivalate, $CH_3CN$, reflux, 55%. b) Piperidine, 95%. c) $H^+$ resin, 95%.

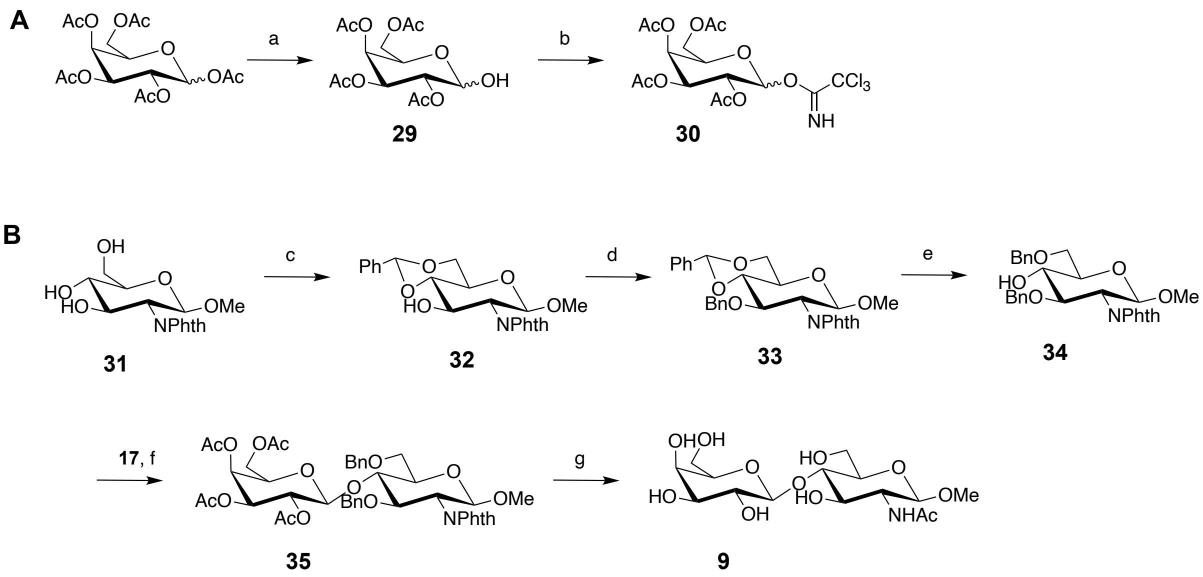


**Scheme S3.** Synthesis of substrate **9**. **A** Synthesis of building block **30**. a) Benzylamine, THF, 84%. b) Trichloroacetonitrile, Potassium carbonate, DCM, 77%. **B** Synthesis of LacNAc substrate **9.** c) Benzaldehyde dimethyl acetal, p-TsOH, $CH_3CN$, 95%. d) BnBr, NaH, DMF, 50%. e) Triethyl silane, trifluoroacetic acid, DCM, 44%. f) TMSOTf, DCM, 4 Å ms, 55%. g) i. MeONa/MeOH, ii. ethylenediamine, EtOH, iii. MeOH, $Ac_2O$, iv. $H_2$, Pd/C, MeOH, $H_2O$, 95%.

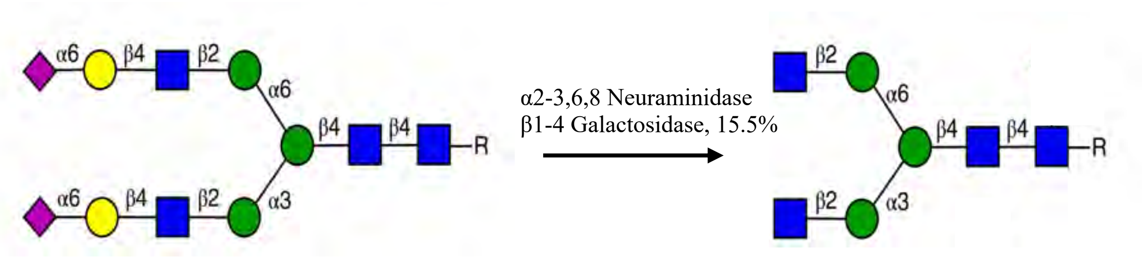


**SGP, extrated from egg yolk** **8**

**Figure S1.** Preparation of substrate **8**.

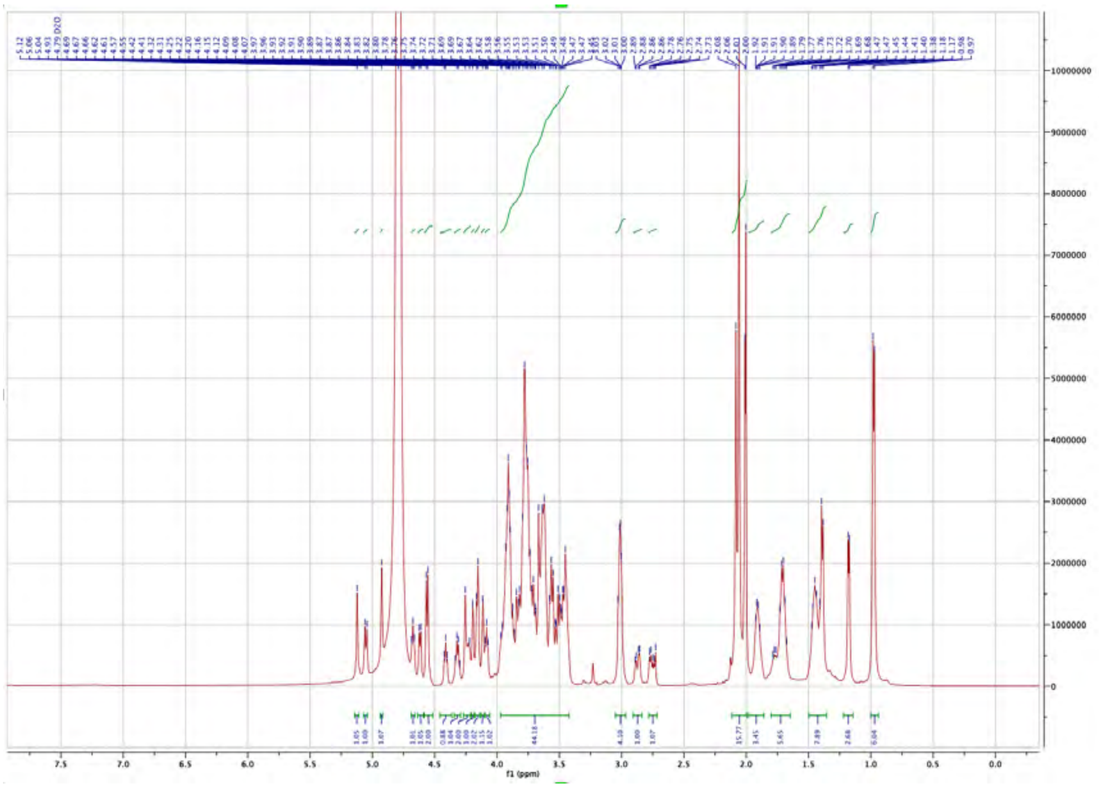

**Figure S2.** $^{1}$H NMR spectrum of compound **8** (600 MHz, Deuterium Oxide).

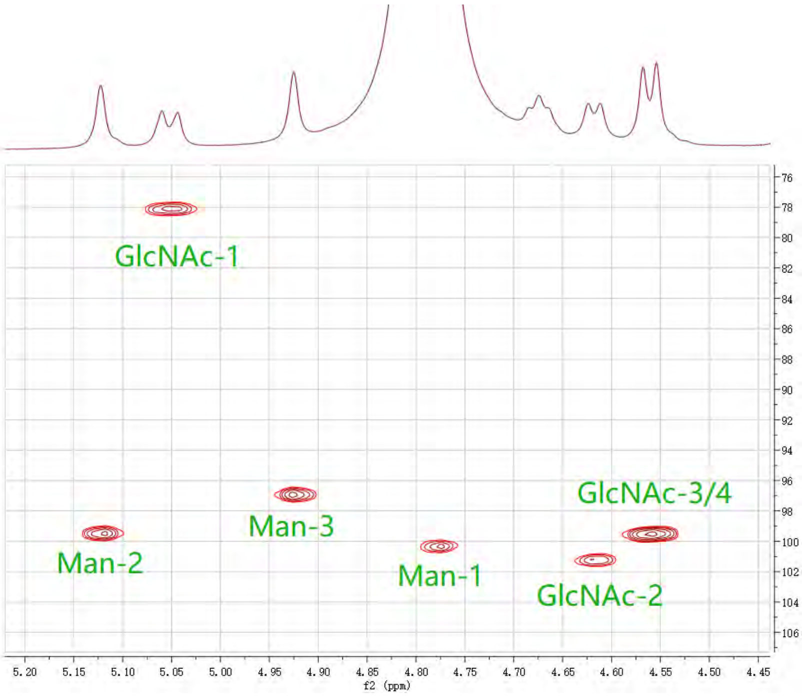


**Figure S3.** Anomeric region of the $^{13}$C -$^{1}$H HSQC spectrum (complete anomeric assignment of **8**).

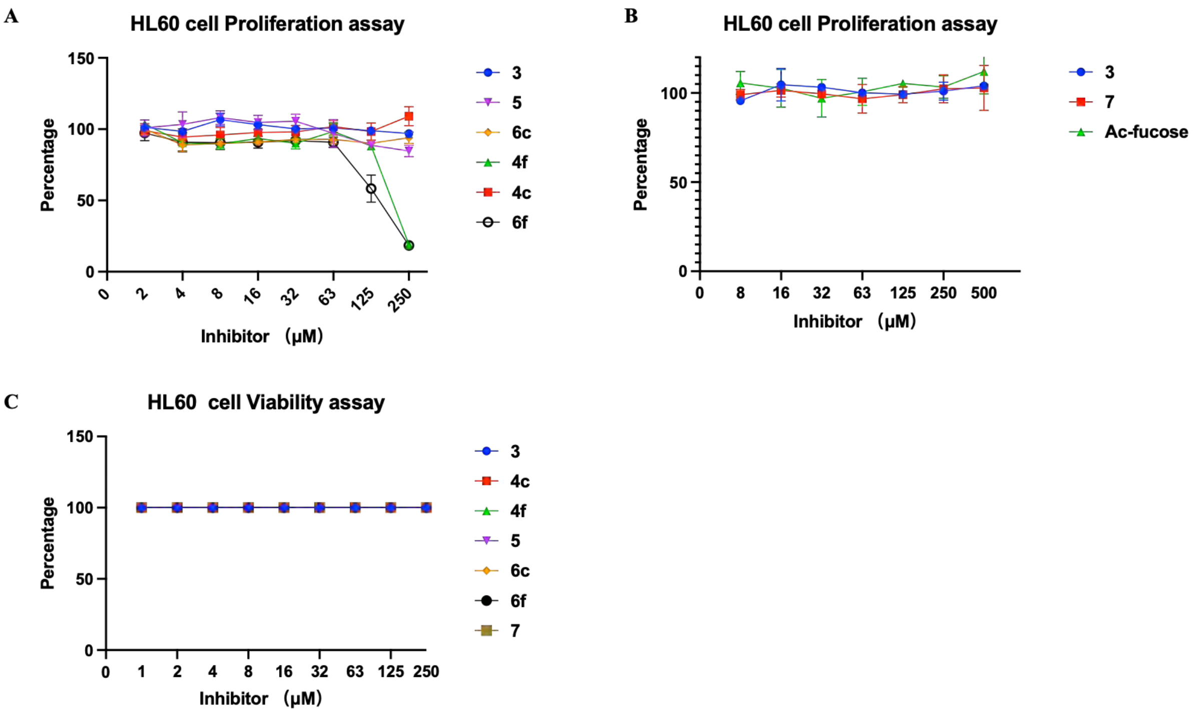


**Figure S4.** Proliferation (**A**, **B**) and viability (**C**) results with HL-60 cells.

#### 2) Experimental Procedures

##### General Methods

All chemicals were purchased from Sigma-Aldrich, Fisher Scientific, or Biosynth Carbosynth. NMR spectra were recorded with an Agilent 400 or a Bruker 600. 1H NMR data are presented in the order: Chemical shift, multiplicity (s = singlet, d = doublet, t = triplet, dd = doublet of doublets, m = multiplet), and coupling constants (J) are reported in Hertz (Hz). High-resolution mass spectrometry (HRMS) was recorded on an Agilent Technologies 6560 Ion mobility Q-TOF. Column chromatography was performed on silica gel G60 (Silicycle, 60-200 μm, 60 Å), and size exclusion chromatography was performed on Bio-Gel P-2 (45-90 μm) by using 10-100 mM $NH_4HCO_3$ in Mili-Q $H_2O$ as eluent. Thin layer chromatography (TLC) analysis was conducted on Silica gel 60 F254 (EMD Chemicals Inc.) coated aluminum sheets and detected by using UV light (254 nm), staining with 5% sulfuric acid in ethanol or p-anisaldehyde solution. Acid-washed molecular sieves were flame-activated under vacuum before reactions. Ion-exchange resin was purchased from Sigma-Aldrich. Before use, the $H^+$ resin (Amberlite IRC 120H, hydrogen form) was washed and activated, sequentially washed with Mili-Q $H_2O$, 1N NaOH, Mili-Q $H_2O$, 1N HCL, and Mili-Q $H_2O$.

#### Synthetic Protocols and Compound Characterization

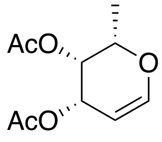


**23**

**3,4-Di-*O*-acetyl-fucal**

L-fucose (15 g, 90 mmol) was dissolved in acetic anhydride (75 mL), followed by the dropwise addition of $HClO_4$ (0.75 mL) at 0 °C. The reaction mixture was stirred at room temperature (RT) for 2 h, then cooled back to 0 °C, and HBr (130 mL, 33% in AcOH) was added. After stirring at RT for 1 h, the reaction mixture was concentrated under vacuum. The resulting residue was dissolved in ethyl acetate/saturated $NaH_2PO_4$ (1:2, 390 mL), and zinc dust (66 g) was added. The mixture was stirred for 12 h at RT, then filtered. The resulting solution was extracted with ethyl acetate, the organic phase was separated, dried with $Na_2SO_4$, and filtered again. The filtrate was concentrated under vacuum. Purification through silica gel column chromatography afforded **23** as a colorless oil (14.3 g, 73%).

$^1$H NMR (600 MHz, Chloroform-*d*) δ 6.45 (dd, *J* = 6.3, 1.9 Hz, 1H, H-1), 5.56 (m, 1H, H-3), 5.27 (dt, *J* = 4.7, 1.3 Hz, 1H,H-4), 4.63 (dt, *J* = 6.3, 2.0 Hz, 1H, H-2), 4.20 (q, *J* = 6.2 Hz, 1H, H-5), 2.15 (s, 3H, $CH_3$-Ac), 2.00 (s, 3H, $CH_3$-Ac), 1.26 (d, *J* = 6.6 Hz, 3H, $CH_3$-Fuc). $^{13}$C NMR (151 MHz, Chloroform-*d*) δ 170.82, 170.52, 146.24, 98.39, 71.65, 66.38, 65.16, 20.98, 20.83, 16.65.

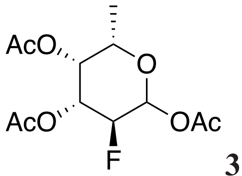


**3**

**2-Deoxy-2-fluoro-per-*O*-acetyl-fucose**

Compound **23** (6 g, 28 mmol) was dissolved in a mixture of DMF and $H_2O$ (1:1, 100 mL), followed by the addition of Selectfluor (16 g, 46 mmol). The mixture was stirred at 50 °C for 5 h. Then the mixture was diluted with ethyl acetate and washed with sat. aqueous $NaHCO_3$. The organic layer was then concentrated *in vacuo*. The resulting residue was dissolved in pyridine (50 mL) and acetic anhydride (50 mL). After stirring for 2 h at RT, the reaction mixture was concentrated *in vacuo*. Purification through silica gel column chromatography afforded **3** as a colorless oil (5.9 g, 73%).

$^{1}H$ NMR (600 MHz, Chloroform-*d*) δ 6.42 (t, *J* = 4.1 Hz, 1.3H, αH-1), 5.77 (m, 1H, βH-1), 5.40 (m, 1.4H, αH-3), 5.36 (m, 1.3H, αH-4), 5.30 (m, 1H, βH-4), 5.16 (m, 1H, βH-3), 4.92 (dt, *J* = 10.2, 3.9 Hz, 0.7H, αH-2), 4.83 (dt, *J* = 10.2, 3.9 Hz, 0.7H, αH-2), 4.67 (ddd, *J* = 9.7, 8.0, 4.7 Hz, 0.5H, βH-2), 4.58 (ddd, *J* = 9.7, 8.0, 4.7 Hz, 0.5H, βH-2), 4.24 (m, 1.4H, αH-5), 3.98 (q, *J* = 6.5, 6.0 Hz, 1H, βH-5), 2.24 – 2.14 (m, 14H, α/β $CH_3$-Ac), 2.06 (m, 7H, α/β $CH_3$-Ac), 1.21 (dd, *J* = 6.5 Hz, 3H, β $CH_3$-Fuc), 1.14 (dd, *J* = 6.6, 4H, α $CH_3$-Fuc). $^{13}C$ NMR (151 MHz, Chloroform-*d*) δ 170.45, 170.43, 170.24, 169.99, 169.23, 169.14, 91.75 (d, *J* = 24.3 Hz, βC-1), 89.35 (d, *J* = 22.3 Hz, αC-1), 86.93 (d, *J* = 188.0 Hz, βC-2), 84.37 (d, *J* = 190.7 Hz, αC-2), 71.49 (d, *J* = 18.4 Hz, βC-3), 71.21 (d, *J* = 7.7 Hz, αC-4), 70.72 (d, *J* = 8.2 Hz, βC-4), 70.44, 68.74 (d, *J* = 18.6 Hz, αC-3), 67.27, 21.06, 21.00, 20.81, 20.73, 20.67, 15.91, 15.88.
ESI MS(m/z): $[M + Na]^+$ calcd for $C_{12}H_{17}FO_7Na$, 315.0856; found 315.0231

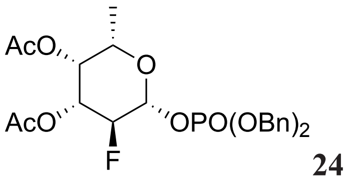


**24**

**2-Deoxy-2-fluoro-3,4-di-*O*-acetyl-β-1-(dibenzylphosphoryl)-L-fucopyranose**

HBr (33% in AcOH, 1 mL) was added to compound **3** (200 mg, 0.68 mmol), and after stirring for 0.5 h at RT, the mixture was concentrated *in vacuo*. The residue was dissolved in 5 mL $CH_3CN$, and 500 mg of 3 Å molecular sieves were added. The mixture was stirred at RT for 10 min. Then, silver carbonate (375 mg, 1.36 mmol) and dibenzyl phosphate (375 mg, 1.36 mmol) were added, the reaction mixture was stirred at RT, under nitrogen gas protection, and in darkness. After 12 h, the mixture was diluted with ethyl acetate and then filtered through Celite. The filtrate was concentrated *in vacuo*. The residue was purified by silica gel chromatography, yielding **24** (244 mg, 70%) as a colorless oil.

$^{1}H$ NMR (600 MHz, Chloroform-*d*) δ 7.44 – 7.31 (m, 10H, Ar), 5.37 (td, *J* = 7.5, 3.9 Hz, 1H, H-1), 5.28 (t, *J* = 3.2 Hz, 1H, H-4), 5.18 – 5.03 (m, 5H, $CH_2$, H-3), 4.65 (dd, *J* = 9.9, 7.7 Hz, 0.5H, H-2), 4.56 (dd, *J* = 9.9, 7.7 Hz, 0.5H, H-2), 3.92 (q, *J* = 6.4 Hz, 1H, H-5), 2.17 (s, 3H, $CH_3$-Ac), 2.06 (s, 3H, $CH_3$-Ac), 1.20 (d, *J* = 6.4 Hz, 3H, $CH_3$-Fuc).
ESI MS(m/z): [M + Na]+ calcd for $C_{24}H_{28}FO_9PNa$, 533.1353; found 533.0648

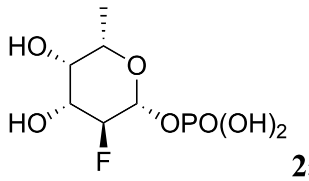

**25**

**2-Deoxy-2-fluoro-β-fucopyranosyl phosphate**

The mixture of Compound **24** (55 mg, 0.1 mmol) in dry MeOH (2 mL) was hydrogenated in an $H_2$ atmosphere with 10% activated Pd/C (40 mg) as the catalyst. After 16 h, the mixture was filtered, and the filtrate was concentrated *in vacuo*. The residue was dissolved in a mixture of MeOH: $H_2O$: $Et_3N$ (7:3:1, 3.3 mL) and stirred for 6 h at RT. The mixture was then concentrated *in vacuo* and purified by silica gel chromatography, affording **25** (24 mg, 88%) as a white solid.

$^1H$ NMR (400 MHz, Deuterium Oxide) δ 5.10 (td, *J* = 8.0, 3.7 Hz, 1H, H-1), 4.43 (dd, *J* = 9.6, 7.7 Hz, 0.5H, H-2), 4.30 (dd, *J* = 9.5, 7.7 Hz, 0.5H, H-2), 3.98 (m, 1H, H-3), 3.88 (q, *J* = 6.5 Hz, 1H, H-5), 3.83 (t, *J* = 3.3 Hz, 1H, H-4), 3.21 (q, *J* = 7.3 Hz, 9H, $CH_2$-$Et_3N$), 1.34 – 1.24 (m, 15H, CH3, $CH_3$-$Et_3N$). $^{13}C$ NMR (101 MHz, Deuterium Oxide) δ 95.12 (d, *J* = 23.9 Hz), 90.80, 71.71 (d, *J* = 9.0 Hz), 71.50, 71.21 (d, *J* = 17.0 Hz), 46.60, 15.14, 8.15.

ESI MS(m/z): $[2M - H]^-$ calcd for $C_{12}H_{23}F_2O_{14}P_2$, 491.0537; found 491.0350

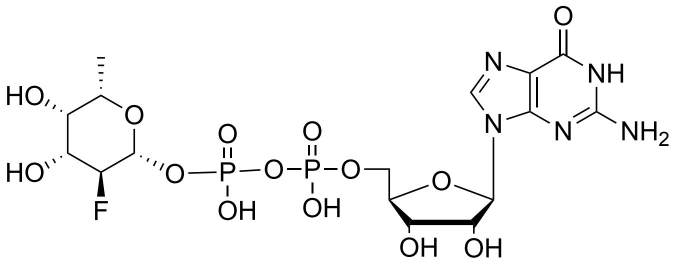

**1**

**GDP-β-2-fluoro-fucose**

Compound **27** (8.5 mg, 0.03 mmol) and GMP-morpholidate (50 mg, 0.07 mmol) were co-evaporated with dry pyridine and dried under vacuum. Then, 1H-Tetrazole (8 mg, 0.1 mmol) and dry pyridine (5 mL) were added. The solution was stirred at RT under $N_2$. After 2 days, the mixture was diluted with water and applied to a Bio-Gel P2 column, eluted with 50 mM $NH_4HCO_3$ to afford the compound **1** (3.7 mg, 18%) as a white solid.

$^1H$ NMR (400 MHz, Deuterium Oxide) δ 8.12 (s, 1H, $H_{Guo}$-8'), 5.94 (d, *J* = 6.3 Hz, 1H, $H_{Guo}$-1), 5.18 (td, *J* = 7.9, 3.5 Hz, 1H, $H_{Fuc}$-1), 4.87 (m, 1H, $H_{Guo}$-2), 4.54 (dd, *J* = 5.2, 3.1 Hz, 1H, $H_{Guo}$-3), 4.44 (dd, *J* = 9.6, 7.5 Hz, 0.5H, $H_{Fuc}$-2), 4.36(m, 1H, $H_{Guo}$-4), 4.31 (dd, *J* = 9.5, 7.5 Hz, 0.5H, $H_{Fuc}$-2), 4.22 (dd, *J* = 5.4, 3.5 Hz, 2H, $H_{Guo}$-5), 3.92 (ddd, *J* = 14.4, 9.6, 3.6 Hz, 1H,

$H_{Fuc}$-3), 3.82 (m, 1H, $H_{Fuc}$-5), 3.78 (t, *J* = 3.5 Hz, 1H, $H_{Fuc}$-4), 1.23 (d, *J* = 6.4 Hz, 3H, $CH_3$-Fuc).
ESI MS(m/z): $[M - H]^-$ calcd for $C_{16}H_{23}FN_5O_{14}P_2$, 590.0706; found 590.0812

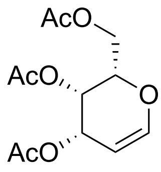

**10**

**3,4,6-Tri-*O*-acetyl-L-galactal**

L-galactose (400 mg, 2.2 mmol) was dissolved in acetic anhydride (4 mL), and then $HClO_4$ (40 μL) was added at 0 °C. The reaction was stirred at RT for 2 h, then cooled to 0 °C, and HBr (3 mL, 33% in AcOH) was added. After stirring for 1 h at RT, the mixture was concentrated *in vacuo*. Then the residue was dissolved in ethyl acetate/saturated $NaH_2PO_4$ (1:2, 18 mL), and Zinc dust (3 g) was added. The mixture was stirred for 12 h at RT, then filtered. The filtrate was extracted with ethyl acetate, dried with $Na_2SO_4$, and filtered again. The filtrate was concentrated *in vacuo*. Silica gel column chromatography afforded **10** as a colorless oil (524 mg, 87%).
$^1$H NMR (400 MHz, Chloroform-*d*) δ 6.45 (dd, *J* = 6.3, 1.8 Hz, 1H, H-1), 5.55 (m, 1H, H-3), 5.42 (dt, *J* = 4.5, 1.7 Hz, 1H, H-4), 4.72 (ddd, *J* = 6.3, 2.7, 1.5 Hz, 1H, H-2), 4.30 (m, 1H, H-5), 4.23 (m, 2H, H-6), 2.12 (s, 3H, $CH_3$-Ac), 2.08 (s, 3H, $CH_3$-Ac), 2.02 (s, 3H, $CH_3$-Ac).
$^{13}$C NMR (101 MHz, Chloroform-*d*) δ 170.54, 170.27, 170.12, 145.39, 98.81, 72.77, 63.86, 63.73, 61.89, 20.77, 20.72, 20.62.
ESI MS(m/z): $[M + Na]^+$ calcd for $C_{12}H_{16}O_7Na$, 295.0794; found 295.0376

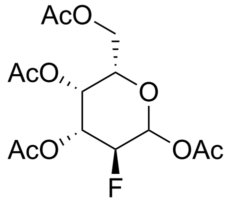

**11**

**2-Deoxy-2-fluoro-per-*O*-acetyl-L-galactose**

Compound **10** (56.6 mg, 0.2 mmol) was dissolved in $MeNO_2/H_2O$ (4:1, 0.5 mL), and then Selectfluor (110.5 mg, 0.3 mmol) was added. The reaction mixture was placed in a microwave reactor and stirred for 5 min at 100 °C. After cooling down to RT, the mixture was diluted with ethyl acetate and washed with water. The organic layer was dried over $Na_2SO_4$ and concentrated *in vacuo*. The resulting residue was dissolved in pyridine (2 mL) and acetic anhydride (2 mL). After stirring for 0.5 h at RT, the reaction mixture was concentrated *in vacuo*. Silica gel chromatography afforded **11** (47 mg, 65%) as a colorless oil.

$^{1}$H NMR (400 MHz, Chloroform-*d*) δ 6.46 (d, *J* = 3.9 Hz, 1H, αH-1), 5.78 (dd, *J* = 8.0, 4.1 Hz, 2.3H, βH-1), 5.51 (td, *J* = 3.5, 1.4 Hz, 1H, αH-4), 5.45 (m, 2.3H, βH-4), 5.40 (td, *J* = 10.6, 3.5 Hz, 1H, αH-3), 5.17 (ddd, *J* = 13.2, 9.8, 3.6 Hz, 2.3H, βH-3), 4.95 (dd, *J* = 10.2, 4.0 Hz, 0.5H, αH-2), 4.83 (dd, *J* = 10.2, 4.0 Hz, 0.5H, αH-2), 4.70 (dd, *J* = 9.8, 8.0 Hz, 1.2H, βH-2), 4.57 (dd, *J* = 9.9, 8.1 Hz, 1.2H, βH-2), 4.31 (td, *J* = 6.8, 1.4 Hz, 1H, αH-5), 4.17 – 4.04 (m, 9H, βH-5, βH-6, αH-6), 2.19 (m, 10H, $CH_3$-Ac), 2.14 (m, 10H, $CH_3$-Ac), 2.05 (m, 10H, $CH_3$-Ac).

$^{13}$C NMR (101 MHz, Chloroform-*d*) δ 170.45, 170.18, 170.07, 170.03, 169.93, 168.97, 168.94, 91.70 (d, *J* = 24.6 Hz, βC-1), 89.08 (d, *J* = 22.4 Hz, αC-1), 86.82 (d, *J* = 188.4 Hz, βC-2), 84.26 (d, *J* = 191.3 Hz, αC-2), 71.85 (βC-5), 71.07 (d, *J* = 18.7 Hz, βC-3), 68.69 (αC-5), 68.29 (d, *J* = 18.9 Hz, αC-3), 68.02 (d, *J* = 7.8 Hz, αC-4), 67.57 (d, *J* = 8.3 Hz, βC-4), 61.09 (αC-6), 60.95 (βC-6), 21.00, 20.94, 20.76, 20.68, 20.64.

$^{19}$F NMR (376 MHz, Chloroform-*d*) δ -208.15 (ddt, *J* = 51.4, 12.9, 3.0 Hz), -209.15 (ddd, *J* = 49.2, 11.0, 3.4 Hz).

ESI MS(m/z): $[M + Na]^+$ calcd for $C_{14}H_{19}FO_9Na$, 373.0911; found 373.0214

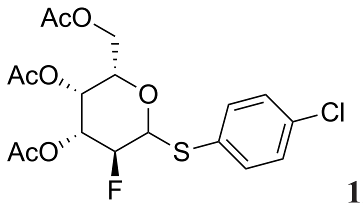


**12**

**Para-chlorothiophenyl 2-deoxy-2-fluoro-per-*O*-acetyl-β-L-galactopyranoside**

Compound **11** (103 mg, 0.29 mmol) was dissolved in HBr (33% in AcOH, 3 mL) and stirred for 1 h at RT. The mixture was then concentrated *in vacuo* and dissolved in $CHCl_3$ (2 mL). The solution was cooled to 0 ºC, and a cold mixture of tetrabutylammonium hydrogen sulfate (108 mg, 0.38 mmol) and p-chlorothiophenol (108 mg, 0.7 mmol) in 1M NaOH (0.5 mL) was added. The resulting brown mixture was allowed to warm to RT and stirred for 18 h. The reaction mixture was diluted with DCM and washed with 1N NaOH, then dried over $Na_2SO_4$ and concentrated *in vacuo*. The resulting residue was purified by silica gel chromatography, yielding **12** (78 mg, 56%) as a white solid.

$^{1}$H NMR (600 MHz, Chloroform-*d*) δ 7.53 (d, *J* = 8.5 Hz, 2H, Ar), 7.32 (d, *J* = 8.5 Hz, 2H, Ar), 5.45 – 5.38 (m, 1H, H-4), 5.12 (ddd, *J* = 13.1, 9.4, 3.5 Hz, 1H, H-3), 4.70 (dd, *J* = 9.7, 2.7 Hz, 1H, H-1), 4.47 (t, *J* = 9.5 Hz, 0.5H, H-2), 4.38 (t, *J* = 9.5 Hz, 0.5H, H-2), 4.17 (dd, *J* =

11.4, 6.9 Hz, 1H, H-6), 4.09 (dd, *J* = 11.3, 6.2 Hz, 1H, H-6), 3.94 (m, 1H, H-5), 2.08 (s, 3H, $CH_3$-Ac), 2.05 (s, 3H, $CH_3$-Ac), 2.03 (s, 3H, $CH_3$-Ac).

$^{13}$C NMR (151 MHz, Chloroform-*d*) δ 170.48, 170.01, 135.34, 135.28, 129.41, 129.27, (86.21, 84.97, d, *J* = 188.4 Hz, C-2), (85.07, 84.91, d, *J* = 24.1 Hz, C-1), 74.74 (C-5), 72.13 (d, *J* = 19.6 Hz, C-3), 68.01 (d, *J* = 8.2 Hz, C-4), 61.52 (C-6), 20.81, 20.73, 20.63.

ESI MS(m/z): $[M + Na]^+$ calcd for $C_{18}H_{20}ClFSNaO_7$, 457.0500; found 456.9933

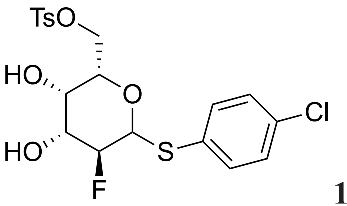


**13**

**Para-chlorothiophenyl 2-deoxy-2-fluoro-6-*O*-tosyl-β-L-galactopyranoside**

Compound **12** (320 mg, 0.74 mmol) was dissolved in MeOH (10 mL), and a catalytic quantity of 1M NaOMe in methanol was added. The reaction mixture was stirred for 1 h at RT, then neutralized with $H^+$ form resin. The resin was removed by filtration, and the filtrate was concentrated *in vacuo*. The resulting residue was dissolved in pyridine (6 mL), and 4-toluenesulfonyl chloride (168.5 mg, 0.88 mmol) was added. The reaction mixture was stirred for 24 h under $N_2$ at RT. Afterwards, the mixture was diluted with ethyl acetate and washed with brine and 1N HCl. The organic layer was dried over $Na_2SO_4$ and concentrated *in vacuo*. The residue was purified by silica gel chromatography, affording **13** (231 mg, 68%) as a white solid.

$^1$H NMR (400 MHz, Chloroform-*d*) δ 7.85 – 7.74 (m, 2H, Ar), 7.49 – 7.41 (m, 2H, Ar), 7.36 (m, 2H, Ar), 7.24 (m, 2H, Ar), 4.55 (dd, *J* = 9.7, 1.9 Hz, 1H, H-1), 4.36 (m, 0.5H, H-2), 4.32 – 4.16 (m, 2.5H, H-2, H-6), 4.06 (m, 1H, H-4), 3.91 – 3.75 (m, 2H, H-3, H-5), 2.45 (s, 3H, $CH_3$-OTs). $^{13}$C NMR (101 MHz, Chloroform-*d*) δ 145.51, 134.94, 134.80, 132.56, 130.18, 129.75, 129.31, 128.16, 89.02 (d, *J* = 183.9 Hz, C-2), 84.59 (d, *J* = 24.2 Hz, C-1), 75.89, 72.92 (d, *J* = 19.4 Hz, C-3), 68.87 (d, *J* = 8.5 Hz, C-4), 67.60, 21.83.

ESI MS(m/z): $[M + Na]^+$ calcd for $C_{19}H_{20}ClFS_2NaO_6$, 485.0272; found 484.9734

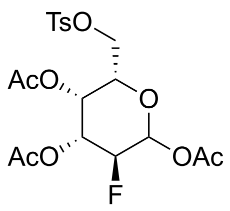

**14**

**2-Deoxy-2-fluoro-6-*O*-tosyl-per-*O*-acetyl-β-L-galactopyranoside**

Compound **13** (345 mg, 0.75 mmol) was dissolved in pyridine (5 mL) and acetic anhydride (3 mL). After stirring for 0.5 h at RT, the reaction mixture was concentrated *in vacuo*. The residue was dissolved in acetone (9 mL) and $H_2O$ (1.5 mL), and then N-Bromosuccinimide (1.3 g, 7.3 mmol) was added. The reaction mixture was stirred for 12 h at RT under $N_2$. Afterward, sat. aq. $NaHCO_3$ (0.5 mL) was added, the mixture was diluted with ethyl acetate and washed with brine. The organic phase was dried over $Na_2SO_4$ and concentrated *in vacuo*. The residue was dissolved in pyridine (2 mL) and acetic anhydride (2 mL). After stirring for 1 h at RT, the mixture was concentrated *in vacuo*. Purification through silica gel column chromatography afforded **14** as a colorless oil (287 mg, 83%).

$^{1}$H NMR (400 MHz, Chloroform-*d*) δ 7.75 (d, *J* = 6.8 Hz, 4.4H, Ar), 7.35 (d, *J* = 8.0 Hz, 4.4H, Ar), 6.40 (d, *J* = 3.9 Hz, 1.2H, αH-1), 5.74 (dd, *J* = 8.0, 4.1 Hz, 1H, βH-1), 5.50 (t, *J* = 3.3 Hz, 1.2H, αH-4), 5.43 (t, *J* = 3.1 Hz, 1H, βH-4), 5.36 (td, *J* = 10.6, 3.5 Hz, 1.2H, αH-3), 5.13 (ddd, *J* = 13.1, 9.9, 3.5 Hz, 1H, βH-3), 4.89 (dd, *J* = 10.3, 4.0 Hz, 0.6H, αH-2), 4.77 (dd, *J* = 10.2, 4.0 Hz, 0.6H, αH-2), 4.65 (dd, *J* = 9.9, 8.0 Hz, 0.5H, βH-2), 4.53 (dd, *J* = 9.9, 8.0 Hz, 0.5H, βH-2), 4.31 (t, *J* = 6.5 Hz, 1.3H, αH-5), 4.15 – 3.91 (m, 5.6H, βH-5, βH-6, αH-6), 2.45 (s, 7H, $CH_3$-Ac), 2.16 (d, *J* = 2.2 Hz, 7H, $CH_3$-Ac), 2.05 (dd, *J* = 7.7, 2.7 Hz, 14H, $CH_3$-Ac).

$^{13}$C NMR (101 MHz, Chloroform-*d*) δ 170.04, 169.80, 169.77, 169.72, 168.85, 168.76, 145.48, 145.45, 132.36, 132.33, 130.13, 130.09, 128.21, 128.19, 91.57 (d, *J* = 24.8 Hz, βC-1), 88.83 (d, *J* = 22.8 Hz, αC-1), 86.62 (d, *J* = 188.5 Hz, βC-2), 84.06 (d, *J* = 191.4 Hz, αC-2), 71.38, 70.81 (d, *J* = 18.8 Hz, βC-3), 68.60, 68.04 (d, *J* = 19.0 Hz, αC-3), 67.80 (d, *J* = 7.9 Hz, αC-4), 67.19 (d, *J* = 8.3 Hz, βC-4), 65.80, 65.21, 21.81, 20.97, 20.87, 20.72, 20.64, 20.52.

ESI MS(m/z): $[M + Na]^+$ calcd for $C_{19}H_{23}FO_{10}SNa$, 485.0894; found 485.0437

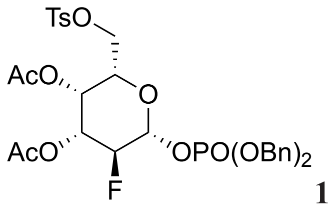

**15**

**2-Deoxy-2-fluoro-3,4-di-*O*-acetyl-6-*O*-tosyl-β-1-(dibenzylphosphoryl)-L-galactopyranose**

HBr (33% in AcOH, 2 mL) was added to compound **14** (269 mg, 0.58 mmol), and after stirring for 0.5 h at RT, the mixture was concentrated *in vacuo*. The residue was dissolved in 5 mL dry $CH_3CN$, and 500 mg of 3 Å molecular sieves were added. The mixture was stirred at RT for 10 min. Then, in one portion, silver carbonate (320 mg, 1.16 mmol) and dibenzyl phosphate (320 mg, 1.16 mmol) were added. The reaction mixture was stirred at RT, under nitrogen gas protection, and in the dark. After 12 h, the mixture was diluted with ethyl acetate and then filtered through Celite. The filtrate was concentrated *in vacuo*. The residue was purified by silica gel chromatography, yielding **15** (337 mg, 84%) in white form.

$^1$H NMR (600 MHz, Chloroform-*d*) δ 7.72 (d, $J$ = 8.3 Hz, 2H, Ar), 7.40 – 7.29 (m, 12H, Ar), 5.40 (t, $J$ = 3.1 Hz, 1H, H-4), 5.34 (td, $J$ = 7.7, 3.9 Hz, 1H, H-1), 5.14 – 5.03 (m, 5H, H-3, $CH_2$-Bn), 4.62 (dd, $J$ = 9.9, 7.7 Hz, 0.5H, H-2), 4.53 (dd, $J$ = 9.9, 7.7 Hz, 0.5H, H-2), 4.07 – 3.95 (m, 3H, H-5, H-6), 2.43 (s, 3H, $CH_3$-Ac), 2.06 (m, 6H, $CH_3$-Ac).

$^{13}$C NMR (151 MHz, Chloroform-*d*) δ 169.78, 169.74, 145.56, 135.52, 135.47, 135.38, 135.33, 132.21, 130.18, 128.85, 128.78, 128.75, 128.71, 128.16, 128.13, 96.30 (dd, $J$ = 24.6, 4.8 Hz, C-1), 87.64 (dd, $J$ = 189.4, 9.0 Hz, C-2), 71.46, 70.47 (d, $J$ = 18.5 Hz, C-3), 69.91 (dd, $J$ = 9.4, 5.4 Hz, $CH_2$-Bn), 67.09 (d, $J$ = 8.1 Hz, C-4), 65.23, 21.81, 20.64, 20.54.

ESI MS(m/z): $[M + Na]^+$ calcd for $C_{31}H_{34}FO_{12}PSNa$, 703.1390; found 703.0974

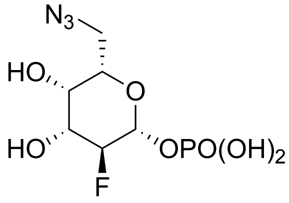

**16**

**2-Deoxy-2-fluoro-6-deoxy-6-$N_3$-β-L-galactopyranosyl 1-phosphate**

The mixture of compound **15** (41 mg, 0.06 mmol) in dry MeOH (3 mL) was hydrogenated in an $H_2$ atmosphere with 10% activated Pd/C (40 mg) as the catalyst. After 2 h, the mixture was filtered, and the filtrate was concentrated *in vacuo*. The residue was dissolved in a mixture of MeOH: H2O: Et3N (7:3:1, 5.5 mL) and stirred for 6 h at RT. The mixture was then concentrated *in vacuo*. The resulting residue was dissolved in DMF (2 mL), and sodium azide (39 mg, 0.6 mmol) was added. The reaction mixture was stirred at 70 °C for 12 h. Afterward, the mixture was concentrated *in vacuo*. The residue was purified by silica gel chromatography, yielding **16** (14 mg, 81%) as a colorless oil.

$^{1}$H NMR (400 MHz, Deuterium Oxide) δ 5.38 (dd, $J$ = 15.3, 8.6 Hz, 1H, H-1), 4.91 (d, $J$ = 5.0 Hz, 0.5H, H-2), 4.55 (d, $J$ = 10.0 Hz, 1H, H-6), 4.52 (d, $J$ = 5.1 Hz, 1H, H-3), 4.48 (t, $J$ = 2.5 Hz, 1H, H-4), 4.42 (dd, $J$ = 4.1, 1.9 Hz, 1H, H-5), 4.00 (dd, $J$ = 9.5, 2.7 Hz, 1H, H-6), 3.36 (s, 0.3H, OH). $^{13}$C NMR (101 MHz, Deuterium Oxide) δ 93.51 (dd, $J$ = 36.3, 4.5 Hz, C-1), 90.47 (dd, $J$ = 178.3, 8.0 Hz, C-2), 77.59 (d, $J$ = 25.3 Hz, C-3), 77.57 (C-4), 70.12 (C-6), 69.65 (C-5).

ESI MS(m/z): [M - H]$^-$ calcd for $C_6H_{10}FN_3O_7P$, 286.0246; found 286.0202

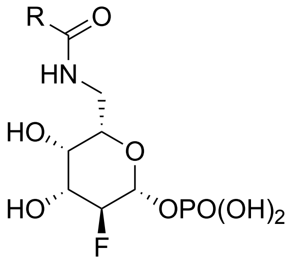


**17a-h**

**General procedure for the synthesis of 17a-h**

Compound **16** (0.03 mmol) was dissolved in t-BuOH (0.5 mL) and 10 mM $NH_4HCO_3$ (0.5 mL), Pd/C (10 mg) was added, and the reaction was stirred in an $H_2$ atmosphere. After 3 h, the reaction mixture was filtered and concentrated *in vacuo*. The resulting residue was dissolved in THF (0.5 mL) and $H_2O$ (0.5 mL). The corresponding NHS ester (0.07 mmol) and DIPEA (10 μL, 0.06 mmol) were added. The mixture was stirred for 2 h at RT and then concentrated *in vacuo*. The residue was purified by silica gel chromatography, affording **17a-h** (50%-70%) as white foam.

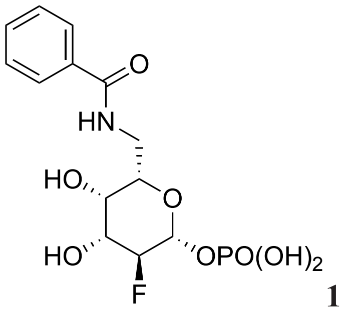


**17b**

**2-Deoxy-2-fluoro-6-deoxy-6-*N*-benzamide-β-L-galactopyranosyl 1-phosphate**

$^{1}$H NMR (600 MHz, Deuterium Oxide) δ 7.75 (d, *J* = 7.5 Hz, 2H, Ar), 7.60 (t, *J* = 7.1 Hz, 1H, Ar), 7.55 – 7.47 (m, 2H, Ar), 5.11 (dq, *J* = 7.9, 3.6, 3.1 Hz, 1H, H-1), 4.44 (t, *J* = 8.6 Hz, 0.5H, H-2), 4.36 (t, *J* = 8.7 Hz, 0.5H, H-2), 4.05 – 3.96 (m, 2H, H-4, H-3), 3.93 (m, 1H, H-5), 3.76 (dd, *J* = 14.1, 4.5 Hz, 1H, H-6a), 3.53 (m, 1H, H-6b).

$^{13}$C NMR (151 MHz, Deuterium Oxide) δ 171.43, 133.44, 132.12, 128.70, 127.15, 95.25 (d, *J* = 24.4 Hz, C-1), 91.58 (d, *J* = 181.9 Hz, C-2), 73.67, (C-5), 70.92 (d, *J* = 17.4 Hz, C-3), 69.54 (d, *J* = 9.1 Hz, C-4), 40.21 (C-6).

ESI MS(m/z): $[M - H]^-$ calcd for $C_{13}H_{16}FNO_8P$, 364.0603; found 364.0464

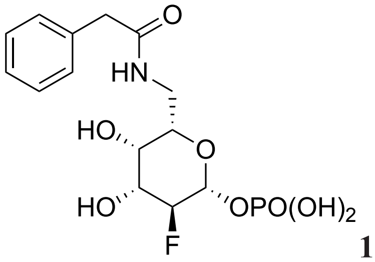


**17c**

**2-Deoxy-2-fluoro-6-deoxy-6-*N*-phenylacetamide-β-L-galactopyranosyl 1-phosphate**

$^{1}$H NMR (600 MHz, Deuterium Oxide) δ 7.40 (t, *J* = 7.4 Hz, 2H, Ar), 7.35 – 7.30 (m, 3H, Ar), 5.06 (td, *J* = 8.1, 3.6 Hz, 1H, H-1), 4.37 (dd, *J* = 9.6, 7.6 Hz, 0.5H, H-2), 4.28 (dd, *J* = 9.4, 7.6 Hz, 0.5H, H-2), 3.97 – 3.89 (m, 2H, H-3, H-4), 3.76 (dd, *J* = 8.4, 4.4 Hz, 1H, H-5), 3.63 (s, 2H, $CH_2$), 3.59 (dd, *J* = 14.1, 4.4 Hz, 1H, H-6a), 3.30 (dd, *J* = 14.1, 8.4 Hz, 1H, H-6b).

$^{13}$C NMR (101 MHz, Deuterium Oxide) δ 175.03, 135.18, 129.08, 128.88, 127.20, 94.96 (d, *J* = 21.4 Hz, C-1), 92.06 (d, *J* = 182.7 Hz, C-2), 73.35, 71.07 (d, *J* = 17.4 Hz, C-4), 69.79 (d, *J* = 8.9 Hz, C-3), 42.05, 40.07.

ESI MS(m/z): $[M - H]^-$ calcd for $C_{14}H_{18}FNO_8P$, 378.0760; found 378.0700

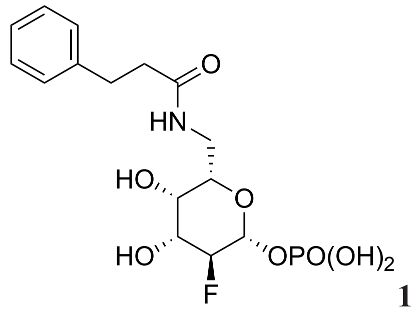

**17d**

**2-Deoxy-2-fluoro-6-deoxy-6-*N*-phenethylamide-β-L-galactopyranosyl 1-phosphate**

$^{1}$H NMR (600 MHz, Deuterium Oxide) δ 7.37 (d, *J* = 15.5 Hz, 2H, Ar), 7.28 (t, *J* = 8.6 Hz, 3H, Ar), 5.01 – 4.96 (m, 1H, H-1), 4.32 (m, 0.5H, H-2), 4.23 (m, 0.5H, H-2), 3.82 (ddd, *J* = 13.0, 9.6, 3.6 Hz, 1H, H-3), 3.54 (m, 1H, H-4), 3.48 (t, *J* = 6.0 Hz, 1H, H-5), 3.41 (dd, *J* = 13.9, 5.6 Hz, 1H, H-6a), 3.22 (dd, *J* = 13.7, 7.8 Hz, 1H, H-6b), 2.93 (t, *J* = 7.3 Hz, 2H, $CH_2$), 2.61 – 2.55 (m, 2H, $CH_2$).

$^{13}$C NMR (151 MHz, Deuterium Oxide) δ 176.06, 140.42, 128.64, 128.50, 126.47, 94.90 (d, *J* = 23.8 Hz, C-1), 92.00 (d, *J* = 181.1 Hz, C-2), 73.06, 71.00 (d, *J* = 17.6 Hz, C-3), 69.27 (d, *J* = 9.0 Hz, C-4), 39.30, 37.05, 31.27.

ESI MS(m/z): $[M - H]^-$ calcd for $C_{15}H_{20}FNO_8P$, 392.0961; found 392.0871

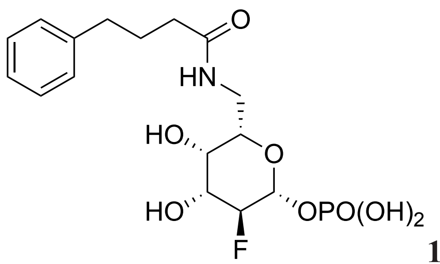

**17e**

**2-Deoxy-2-fluoro-6-deoxy-6-*N*-phenylpropylamide-β-L-galactopyranosyl 1-phosphate**

$^{1}$H NMR (600 MHz, Deuterium Oxide) δ 7.36 (t, *J* = 7.6 Hz, 2H, Ar), 7.27 (dd, *J* = 12.0, 6.9 Hz, 3H, Ar), 5.08 (td, *J* = 7.9, 3.7 Hz, 1H, H-1), 4.41 (dd, *J* = 9.5, 7.6 Hz, 0.5H, H-2), 4.32 (dd, *J* = 9.5, 7.6 Hz, 0.5H, H-2), 3.98 – 3.90 (m, 2H, H-3, H-4), 3.71 (dd, *J* = 8.2, 4.6 Hz, 1H, H-5), 3.48 (dd, *J* = 14.1, 4.4 Hz, 1H, H-6a), 3.25 (dd, *J* = 14.1, 8.4 Hz, 1H, H-6b), 2.64 (t, *J* = 7.5 Hz, 2H, $CH_2$), 2.27 (t, *J* = 7.3 Hz, 2H, $CH_2$), 1.92 (p, *J* = 7.4 Hz, 2H, $CH_2$).

$^{13}$C NMR (151 MHz, Deuterium Oxide) δ 177.08, 141.97, 128.66, 128.58, 126.09, 95.20 (d, *J* = 19.9 Hz, C-1), 91.63 (d, *J* = 178.7 Hz, C-2), 73.66, 70.90 (d, *J* = 17.2 Hz, C-3), 69.53 (d, *J* = 9.3 Hz, C-4), 39.65, 35.00, 34.36, 26.74.

ESI MS(m/z): $[M - H]^-$ calcd for $C_{16}H_{22}FNO_8P$, 406.1073; found 406.0759

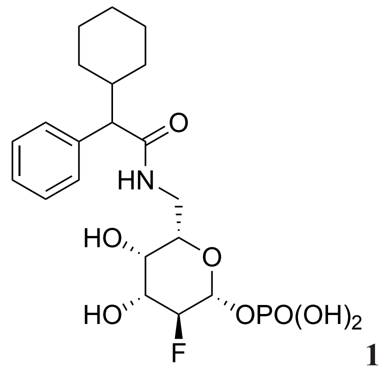

**17f**

**2-Deoxy-2-fluoro-6-deoxy-6-*N*-(2-cyclohexyl-phenylacetamide)-β-L-galactopyranosyl 1-phosphate**

(R, S mixture) $^{1}$H NMR (600 MHz, Deuterium Oxide) δ 7.44 – 7.37 (m, 8H, Ar), 7.34 (m, 2H, Ar), 5.06 (td, *J* = 8.0, 3.6 Hz, 1H, H-1), 5.02 (td, *J* = 8.1, 3.6 Hz, 1H, H-1), 4.36 (ddd, *J* = 10.5, 7.7, 3.3 Hz, 1H, H-2), 4.27 (ddd, *J* = 10.9, 7.6, 3.4 Hz, 1H, H-2), 3.89 (m, 2H, H-3), 3.84 (m, 1H, H-4), 3.77 (t, *J* = 3.3 Hz, 1H, H-4), 3.71 (dd, *J* = 7.1, 5.4 Hz, 1H, H-5), 3.61 (m, 2H, H-6, H-5), 3.55 (dd, *J* = 14.0, 5.0 Hz, 1H, H-6), 3.31 (d, *J* = 11.8 Hz, 1H, CH), 3.27 (dd, *J* = 14.0, 7.8 Hz, 1H, H-6), 3.17 (dd, *J* = 14.1, 7.8 Hz, 1H, H-6), 2.05 (m, 2H, CH), 1.77 – 1.68 (m, 4H, $CH_2$), 1.65 – 1.57 (m, 4H, $CH_2$), 1.32 (m, 4H, $CH_2$), 1.21 – 1.06 (m, 6H, $CH_2$), 0.80 (d, *J* = 12.2 Hz, 2H, $CH_2$).

$^{13}$C NMR (151 MHz, Deuterium Oxide) δ 177.12, 177.02, 138.77, 128.71, 128.69, 128.35, 127.34, 94.93 (d, *J* = 28.3 Hz, C-1), 91.92 (d, *J* = 180.9 Hz, C-2), 73.55, 73.36, 71.03 (d, *J* = 17.3 Hz, C-3), 69.69 (d, *J* = 8.9 Hz, C-4), 58.99, 58.93, 39.66, 39.61, 39.31, 39.17, 31.19, 30.05, 25.81, 25.53, 25.45.

ESI MS(m/z): $[M - H]^-$ calcd for $C_{20}H_{28}FNO_8P$, 460.1542; found 460.1459

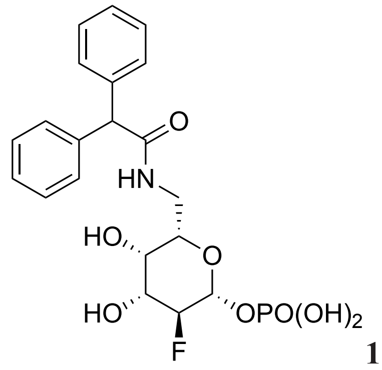

**17g**

**2-Deoxy-2-fluoro-6-deoxy-6-*N*-diphenylacetamide-β-L-galactopyranosyl 1-phosphate**

$^{1}$H NMR (600 MHz, Deuterium Oxide) δ 7.42 – 7.37 (m, 4H, Ar), 7.36 – 7.33 (m, 2H, Ar), 7.30 (d, *J* = 9.8 Hz, 4H, Ar), 5.19 (s, 1H, CH), 5.02 (td, *J* = 8.0, 3.7 Hz, 1H, H-1), 4.34 (dd, *J*

= 9.6, 7.6 Hz, 0.5H, H-2), 4.26 (dd, *J* = 9.6, 7.6 Hz, 0.5H, H-2), 3.91 (ddd, *J* = 13.7, 9.6, 3.6 Hz, 1H, H-3), 3.86 (t, *J* = 3.4 Hz, 1H, H-4), 3.72 (dd, *J* = 7.6, 5.3 Hz, 1H, H-5), 3.65 (dd, *J* = 14.0, 5.2 Hz, 1H, H-6a), 3.32 (dd, *J* = 13.9, 7.5 Hz, 1H, H-6b).

$^{13}$C NMR (151 MHz, Deuterium Oxide) δ 182.39, 175.46, 160.31, 138.95, 128.86, 128.72, 127.49, 94.95 (d, *J* = 27.2 Hz, C-1), 92.03 (d, *J* = 181.4 Hz, C-2), 73.24, 71.01 (d, *J* = 17.4 Hz, C-3), 69.63 (d, *J* = 9.2 Hz, C-4), 57.08, 39.93.

ESI MS(m/z): $[M - H]^-$ calcd for $C_{20}H_{22}FNO_8P$, 454.1073; found 454.0954

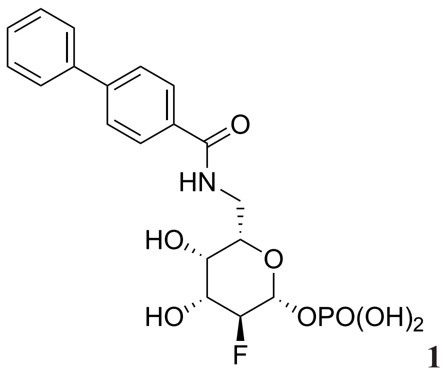


**17h**

**2-Deoxy-2-fluoro-6-deoxy-6-*N*-(biphenyl-4-amide)-β-L-galactopyranosyl 1-phosphate**

$^{1}$H NMR (600 MHz, Deuterium Oxide) δ 7.89 (d, *J* = 8.3 Hz, 2H, Ar), 7.82 (d, *J* = 8.3 Hz, 2H, Ar), 7.78 (d, *J* = 8.0 Hz, 2H, Ar), 7.57 (t, *J* = 7.7 Hz, 2H, Ar), 7.49 (t, *J* = 7.6 Hz, 1H, Ar), 5.10 (td, *J* = 7.6, 3.1 Hz, 1H, H-1), 4.43 (t, *J* = 8.6 Hz, 0.5H, H-2), 4.34 (t, *J* = 8.7 Hz, 0.5H, H-2), 4.06 (t, *J* = 3.4 Hz, 1H, H-4), 4.01 (ddd, *J* = 13.5, 9.6, 3.6 Hz, 1H, H-3), 3.93 (dd, *J* = 8.2, 4.5 Hz, 1H, H-5), 3.86 (dd, *J* = 14.1, 4.4 Hz, 1H, H-6a), 3.57 – 3.51 (m, 1H, H-6b).

ESI MS(m/z): $[M - H]^-$ calcd for $C_{19}H_{20}FNO_8P$, 440.0916; found 440.0765

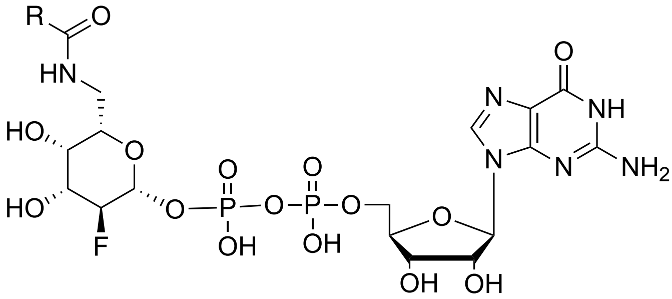


**2a-h**

**General procedure for the synthesis of 2a-h**

A mixture of 0.03 mmol of fucose-1-phosphate analogs (**17a-h**) and GMP-morpholidate (57 mg, 0.08 mmol) was co-evaporated with dry pyridine and then dried under vacuum. 1H-Tetrazole (11 mg, 0.15 mmol) and dry pyridine (3 mL) were added to the mixture, and the solution was stirred at RT under $N_2$. After 2 days, the mixture was diluted with water and applied to a Bio-Gel P2 column, eluted with 50 mM $NH_4HCO_3$. This procedure yielded compounds **2a-h** (25-45%) as white solid products.

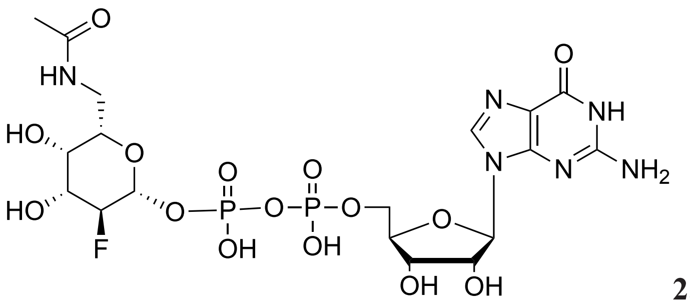

**2a**

**GDP-β-2-deoxy-2-fluoro-6-deoxy-6-*N*-acetamide-L-galactose**

$^1$H NMR (600 MHz, Deuterium Oxide) δ 8.09 (s, 1H, CH, $H_{Guo}$-8'), 5.90 (d, *J* = 6.3 Hz, 1H, $H_{Guo}$-1), 5.15 (td, *J* = 7.9, 3.7 Hz, 1H, $H_{Fuc}$-1), 4.49 (dd, *J* = 5.2, 3.2 Hz, 1H, $H_{Guo}$-3), 4.42 (m, 0.5H, $H_{Fuc}$-2), 4.33 (m, 1.5H, $H_{Fuc}$-2, $H_{Guo}$-4), 4.19 (dd, *J* = 5.5, 3.7 Hz, 2H, $H_{Guo}$-5), 3.90 – 3.84 (m, 2H, $H_{Fuc}$-4, $H_{Fuc}$-3), 3.69 (dd, *J* = 9.0, 3.8 Hz, 1H, $H_{Fuc}$-5), 3.51 (dd, *J* = 14.2, 3.8 Hz, 1H, $H_{Fuc}$-6a), 3.21 (dd, *J* = 14.2, 8.9 Hz, 1H, $H_{Fuc}$-6b), 1.96 (s, 3H, $CH_3$). (The peak of $H_{Guo}$-2 was overlapping with the water peak).

ESI MS(m/z): [M - H]$^-$ calcd for $C_{18}H_{26}FN_6O_{15}P_2$, 674,0921; found 647.0872

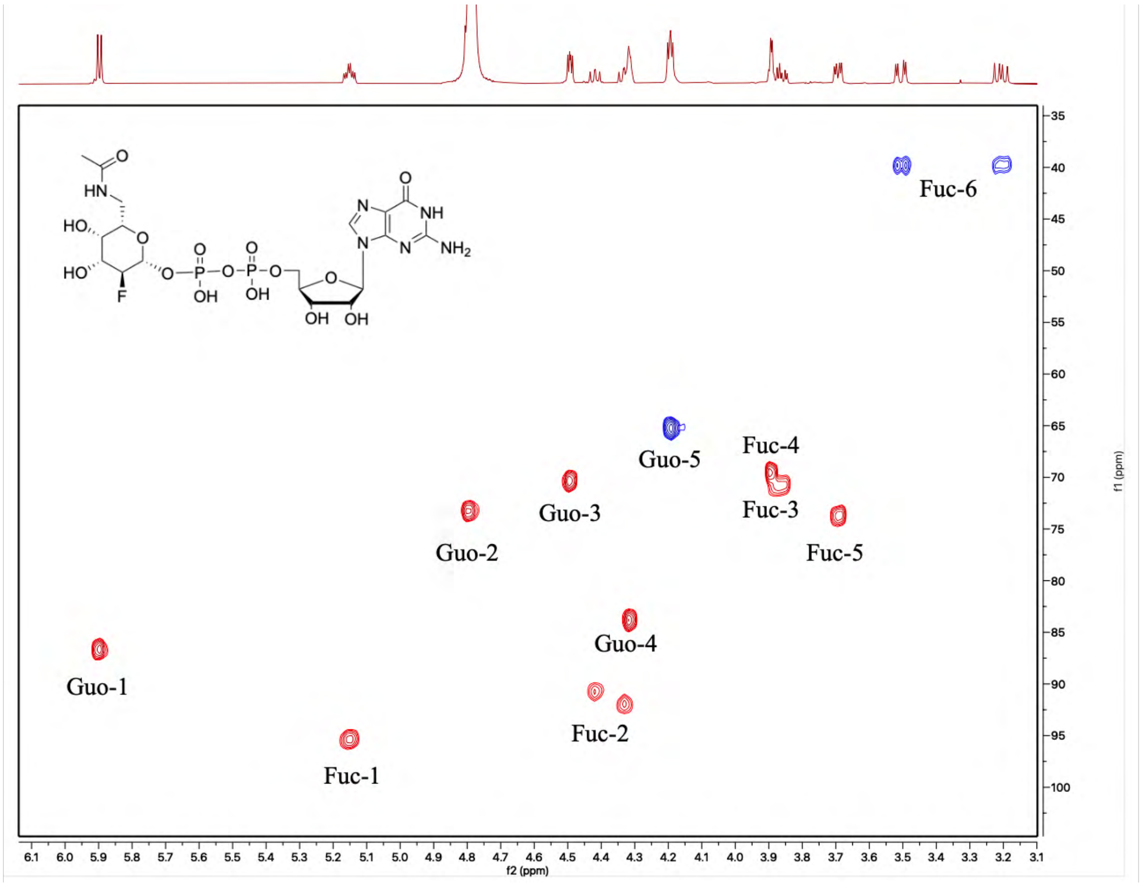


$^{13}$C -$^1$H HSQC spectrum of compound **2a** (sugar region).

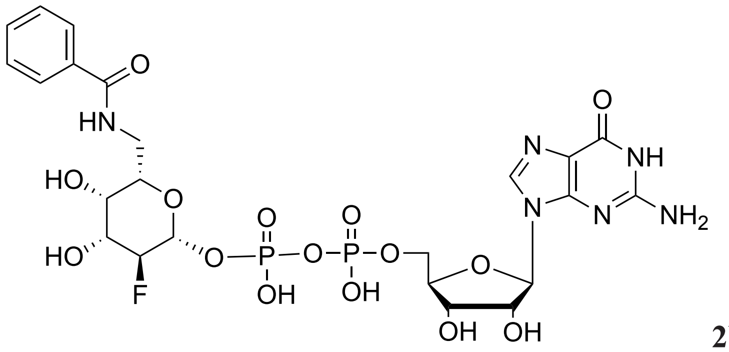


**2b**

**GDP-β-2-deoxy-2-fluoro-6-deoxy-6-*N*-benzamide-L-galactose**

$^{1}$H NMR (600 MHz, Deuterium Oxide) δ 8.12 (s, 1H, $H_{Guo}$-8'), 7.70 (d, $J$ = 7.0 Hz, 2H, Ar), 7.47 (t, $J$ = 7.4 Hz, 1H, Ar), 7.40 (t, $J$ = 7.8 Hz, 2H, Ar), 5.81 (d, $J$ = 5.9 Hz, 1H, $H_{Guo}$-1), 5.21 (td, $J$ = 7.7, 3.7 Hz, 1H, $H_{Fuc}$-1), 4.75 (t, $J$ = 5.5 Hz, 1H, $H_{Guo}$-2), 4.50 (dd, $J$ = 9.6, 7.6 Hz, 1H, $H_{Fuc}$-2), 4.45 (dd, $J$ = 5.2, 3.5 Hz, 1H, $H_{Guo}$-3), 4.42 (dd, $J$ = 9.6, 7.6 Hz, 1H, $H_{Fuc}$-2), 4.29 – 4.23 (m, 2H, $H_{Guo}$-4, $H_{Guo}$-5a), 4.15 (ddd, $J$ = 11.3, 5.7, 3.6 Hz, 1H, $H_{Guo}$-5b), 4.02 (d, $J$ = 3.5 Hz, 1H, $H_{Fuc}$-4), 3.95 (ddd, $J$ = 13.3, 9.6, 3.6 Hz, 1H, $H_{Fuc}$-3), 3.87 (dd, $J$ = 8.4, 3.6 Hz, 1H, $H_{Fuc}$-5), 3.79 (dd, $J$ = 14.2, 3.8 Hz, 1H, $H_{Fuc}$-6a), 3.45 (dd, $J$ = 14.2, 8.6 Hz, 1H, $H_{Fuc}$-6b).

ESI MS(m/z): [M - H]$^{-}$ calcd for $C_{23}H_{29}FN_6O_{15}P_2$, 709.1077; found 709.0745

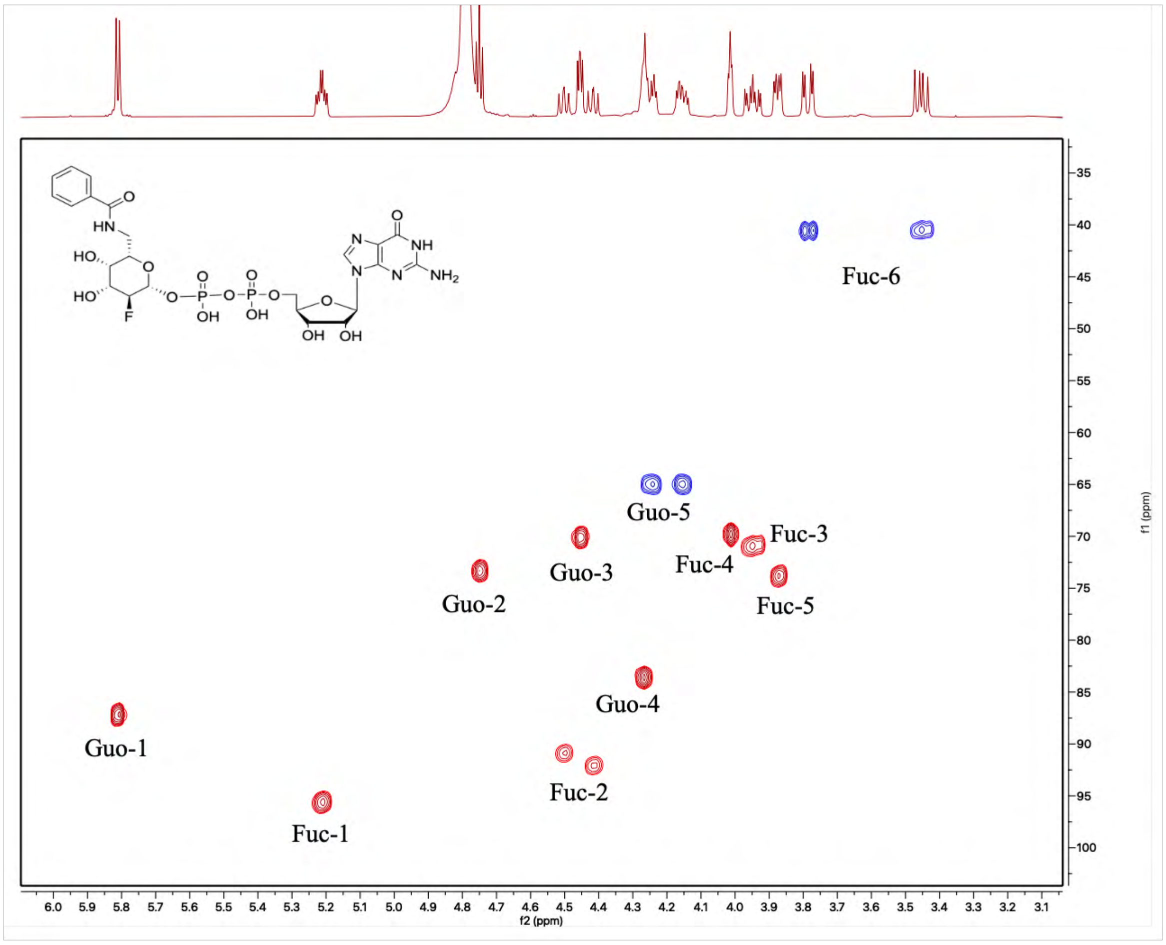


$^{13}$C -$^{1}$H HSQC spectrum of compound **2b** (sugar region).

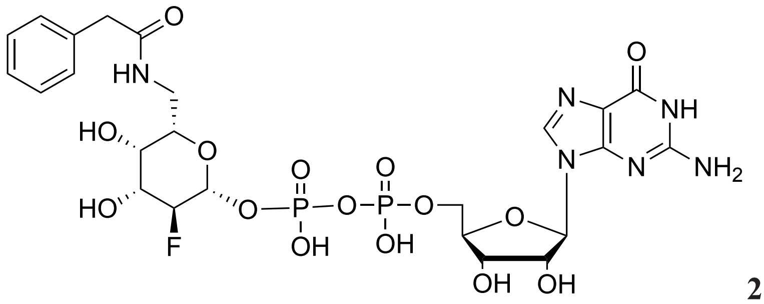

**2c**

**GDP-β-2-deoxy-2-fluoro-6-deoxy-6-*N*-phenylacetamide-L-galactose**

$^{1}$H NMR (600 MHz, Deuterium Oxide) δ 8.05 (s, 1H, $H_{Guo}$-8'), 7.29 (m, 2H, Ar), 7.25 (t, $J$ = 7.2 Hz, 1H, Ar), 7.17 (m, 2H, Ar), 5.85 (d, $J$ = 6.0 Hz, 1H, $H_{Guo}$-1), 5.19 (td, $J$ = 8.0, 3.7 Hz, 1H, $H_{Fuc}$-1), 4.71 (t, $J$ = 5.6 Hz, 1H, $H_{Guo}$-2), 4.48 (m, 1H, $H_{Guo}$-3), 4.46 (m, 0.5H, $H_{Fuc}$-2), 4.36 (m, 0.5H, $H_{Fuc}$-2), 4.32 (m, 1H, $H_{Guo}$-4), 4.23 (m, 2H, $H_{Guo}$-5), 3.90 (m, 2H, $H_{Fuc}$-3, $H_{Fuc}$-4), 3.74 (dd, $J$ = 9.2, 3.7 Hz, 1H, $H_{Fuc}$-5), 3.54 (dd, $J$ = 14.2, 3.7 Hz, 1H, $H_{Fuc}$-6a), 3.52 – 3.43 (m, 2H, $CH_2$), 3.27 (dd, $J$ = 14.2, 9.2 Hz, 1H, $H_{Fuc}$-6b).

$^{13}$C NMR (151 MHz, Deuterium Oxide) δ 174.94, 160.29, 158.98, 153.91, 151.54, 137.43, 134.91, 128.88, 128.62, 126.93, 116.25, 95.59 (d, $J$ = 24.6 Hz), 91.46 (d, $J$ = 182.3 Hz), 86.87, 83.72, 73.81, 73.58, 70.90 (d, $J$ = 17.4 Hz), 70.33, 69.62 (d, $J$ = 9.0 Hz), 65.24, 41.94, 40.05.

ESI MS(m/z): $[M - H]^-$ calcd for $C_{24}H_{30}FN_6O_{15}P_2$, 723.1234; found 723.1148

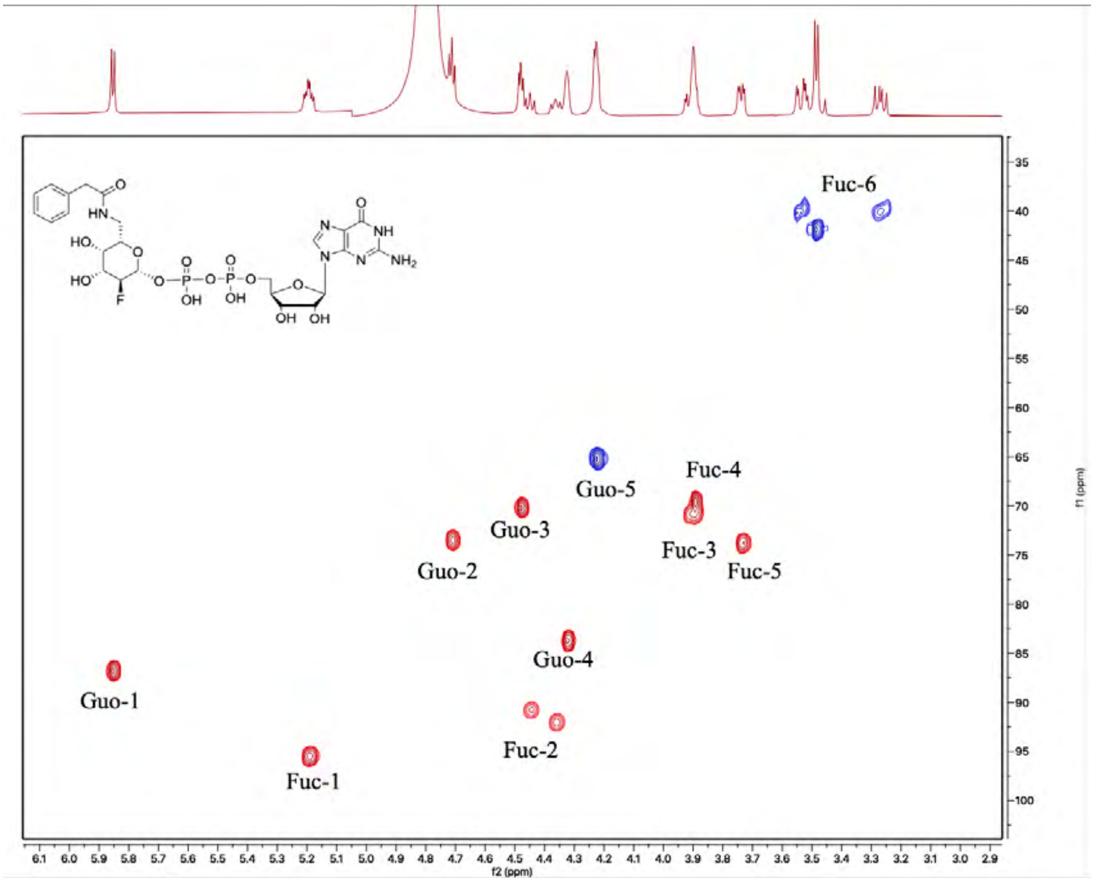


$^{13}$C -$^{1}$H HSQC spectrum of compound **2c** (sugar region).

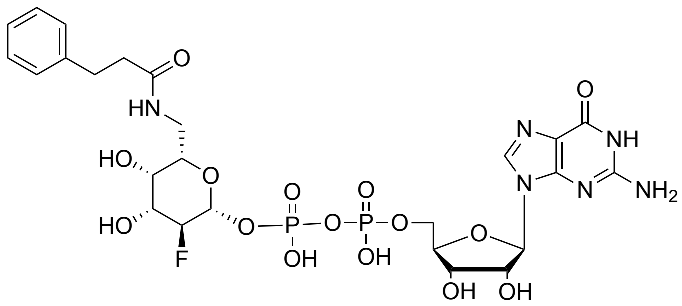

**2d**

**GDP-β-2-deoxy-2-fluoro-6-deoxy-6-*N*-phenethylamide-L-galactose**

$^1$H NMR (600 MHz, Deuterium Oxide) δ 8.12 (s, 1H, $H_{Guo}$-8'), 7.27 (t, *J* = 7.6 Hz, 2H, Ar), 7.19 (m, 3H, Ar), 5.89 (d, *J* = 6.1 Hz, 1H, $H_{Guo}$-1), 5.14 (td, *J* = 7.9, 3.6 Hz, 1H, $H_{Fuc}$-1), 4.51 (dd, *J* = 5.2, 3.4 Hz, 1H, $H_{Guo}$-3), 4.42 (dd, *J* = 9.6, 7.6 Hz, 0.5H, $H_{Fuc}$-2), 4.34 (m, 1.5H, $H_{Guo}$-4, $H_{Fuc}$-2), 4.27 – 4.17 (m, 2H, $H_{Guo}$-5), 3.83 (ddd, *J* = 13.8, 9.6, 3.6 Hz, 1H, $H_{Fuc}$-3), 3.64 (t, *J* = 3.2 Hz, 1H, $H_{Fuc}$-4), 3.50 (dd, *J* = 8.4, 4.5 Hz, 1H, $H_{Fuc}$-5), 3.45 (dd, *J* = 14.0, 4.5 Hz, 1H, $H_{Fuc}$-6a), 3.19 (dd, *J* = 13.9, 8.3 Hz, 1H, $H_{Fuc}$-6b), 2.84 (t, *J* = 7.6 Hz, 2H, $CH_2$), 2.53 (h, *J* = 6.5 Hz, 2H, $CH_2$). (The peak of $H_{Guo}$-2 was overlapping with the water peak)

ESI MS(m/z): [M - H]$^-$ calcd for $C_{25}H_{32}FN_6O_{15}P_2$, 737.1390; found 727.1134

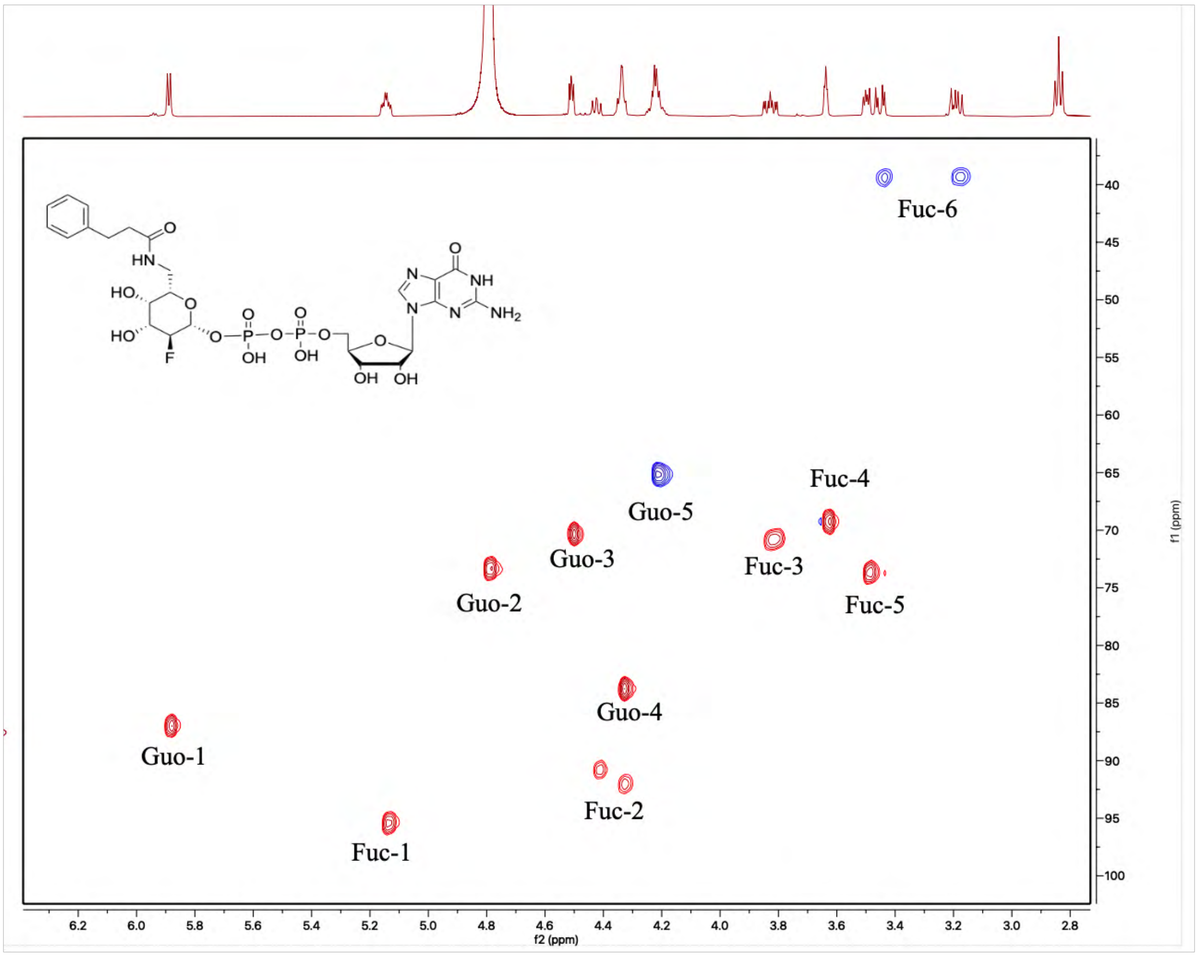


$^{13}$C -$^1$H HSQC spectrum of compound **2d** (sugar region).

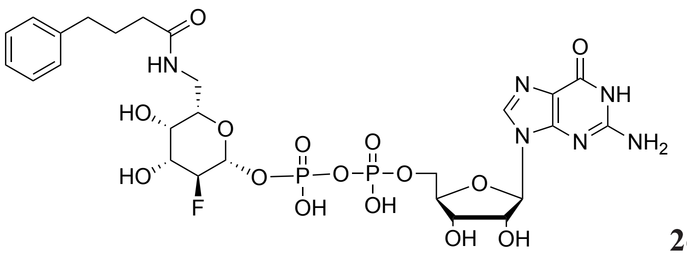

**2e**

**GDP-β-2-deoxy-2-fluoro-6-deoxy-6-*N*-phenylpropylamide-L-galactose**

$^{1}$H NMR (600 MHz, Deuterium Oxide) δ 8.08 (s, 1H, $H_{Guo}$-8'), 7.29 (t, *J* = 7.5 Hz, 2H, Ar), 7.23 – 7.19 (m, 1H, Ar), 7.17 (m, 2H, Ar), 5.88 (d, *J* = 6.1 Hz, 1H, $H_{Guo}$-1), 5.19 (td, *J* = 7.8, 3.5 Hz, 1H, $H_{Fuc}$-1), 4.76 (m, 1H, $H_{Guo}$-2), 4.49 (dd, *J* = 5.2, 3.4 Hz, 1H, $H_{Guo}$-3), 4.46 (dd, *J* = 9.2, 7.6 Hz, 0.5H, $H_{Fuc}$-2), 4.37 (m, 0.5H, $H_{Fuc}$-2), 4.33 (m 1H, $H_{Guo}$-4), 4.21 (m, 2H, $H_{Guo}$-5), 3.95 – 3.89 (m, 2H, $H_{Fuc}$-3, $H_{Fuc}$-4), 3.70 (dd, *J* = 9.1, 3.8 Hz, 1H, $H_{Fuc}$-5), 3.52 (dd, *J* = 14.2, 3.7 Hz, 1H, $H_{Fuc}$-6a), 3.22 (dd, *J* = 14.1, 9.0 Hz, 1H, $H_{Fuc}$-6b), 2.50 (t, *J* = 7.7 Hz, 2H, $CH_2$), 2.25 (t, *J* = 7.4 Hz, 2H, $CH_2$), 1.82 (p, *J* = 7.9 Hz, 2H, $CH_2$).

$^{13}$C NMR (151 MHz, Deuterium Oxide) δ 176.98, 160.27, 158.78, 153.93, 153.80, 141.88, 137.56, 128.41, 128.38, 125.87, 95.54 (d, *J* = 25.1 Hz), 91.49 (d, *J* = 190.1 Hz), 86.89, 83.77, 73.86, 73.55, 70.92 (d, *J* = 17.1 Hz), 70.42, 69.70 (d, *J* = 8.9 Hz), 65.35, 39.88, 35.05, 34.39, 26.72.

ESI MS(m/z): $[M - H]^-$ calcd for $C_{26}H_{34}FN_6O_{15}P_2$, 751.1547; found 751.1041

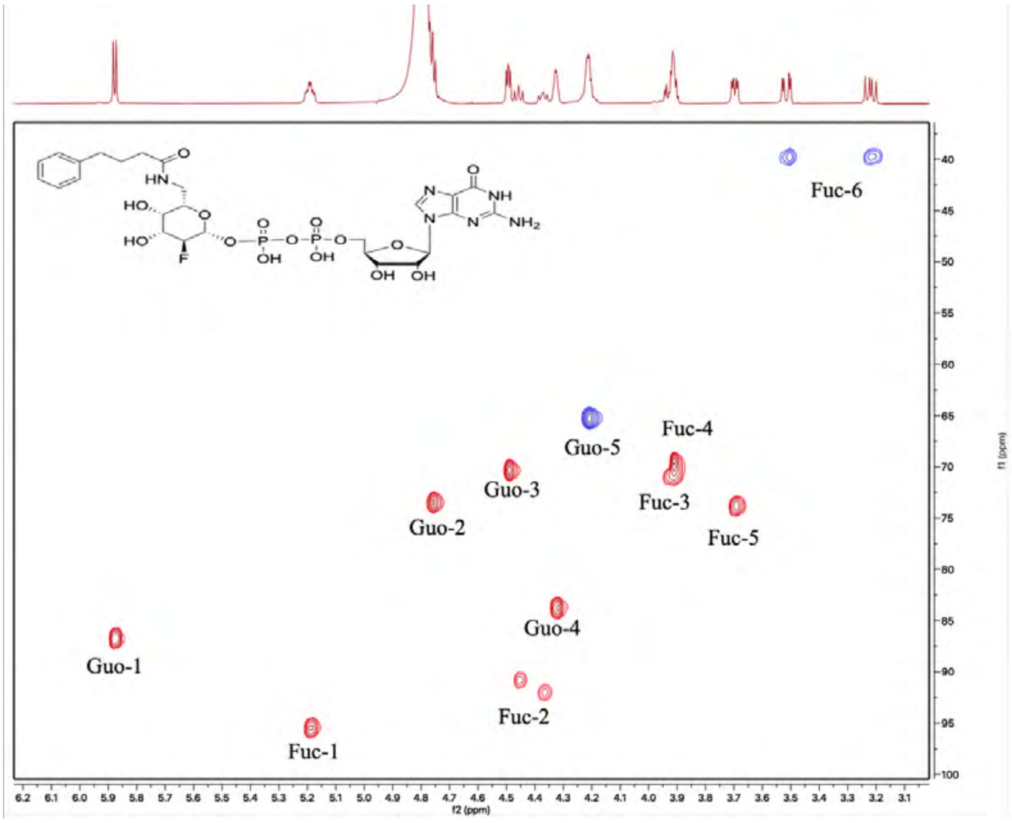


$^{13}$C -$^{1}$H HSQC spectrum of compound **2e** (sugar region).

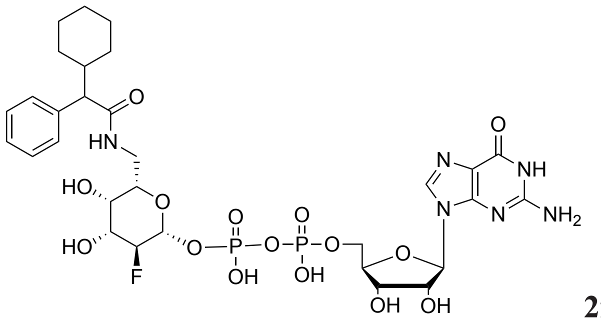

**2f**

**GDP-β-2-deoxy-2-fluoro-6-deoxy-6-*N*-(2-cyclohexyl-phenylacetamide)-L-galactose**

(R, S mixture) $^{1}H$ NMR (600 MHz, Deuterium Oxide) δ 8.08 (s, 1H, $H_{Guo}$-8'), 7.38 (m, 5H, Ar), 5.88 (d, $J$ = 6.3 Hz, 1H, $H_{Guo}$-1), 5.13 (td, $J$ = 8.0, 3.7 Hz, 1H, $H_{Fuc}$-1), 4.50 (dd, $J$ = 5.2, 3.2 Hz, 1H, $H_{Guo}$-3), 4.42 (m, 0.5H, $H_{Fuc}$-2), 4.36 – 4.29 (m, 1.5H, $H_{Fuc}$-2, $H_{Guo}$-4), 4.26 – 4.19 (m, 2H, $H_{Guo}$-5), 3.83 – 3.75 (m, 2H, $H_{Fuc}$-3, $H_{Fuc}$-4), 3.60 – 3.50 (m, 2H, $H_{Fuc}$-5, $H_{Fuc}$-6a), 3.30 – 3.21 (m, 2H, $H_{Fuc}$-6b, CH), 1.98 (m, 1H, CH), 1.71 – 1.57 (m, 4H, $CH_2$), 1.34 – 1.19 (m, 2H, $CH_2$), 1.08 (m, 3H, $CH_2$), 0.65 (m, 1H, $CH_2$). (The peak of $H_{Guo}$-2 was overlapping with the water peak).

ESI MS(m/z): $[M - H]^{-}$ calcd for $C_{30}H_{40}FN_6O_{15}P_2$, 805.2016; found 805.1762

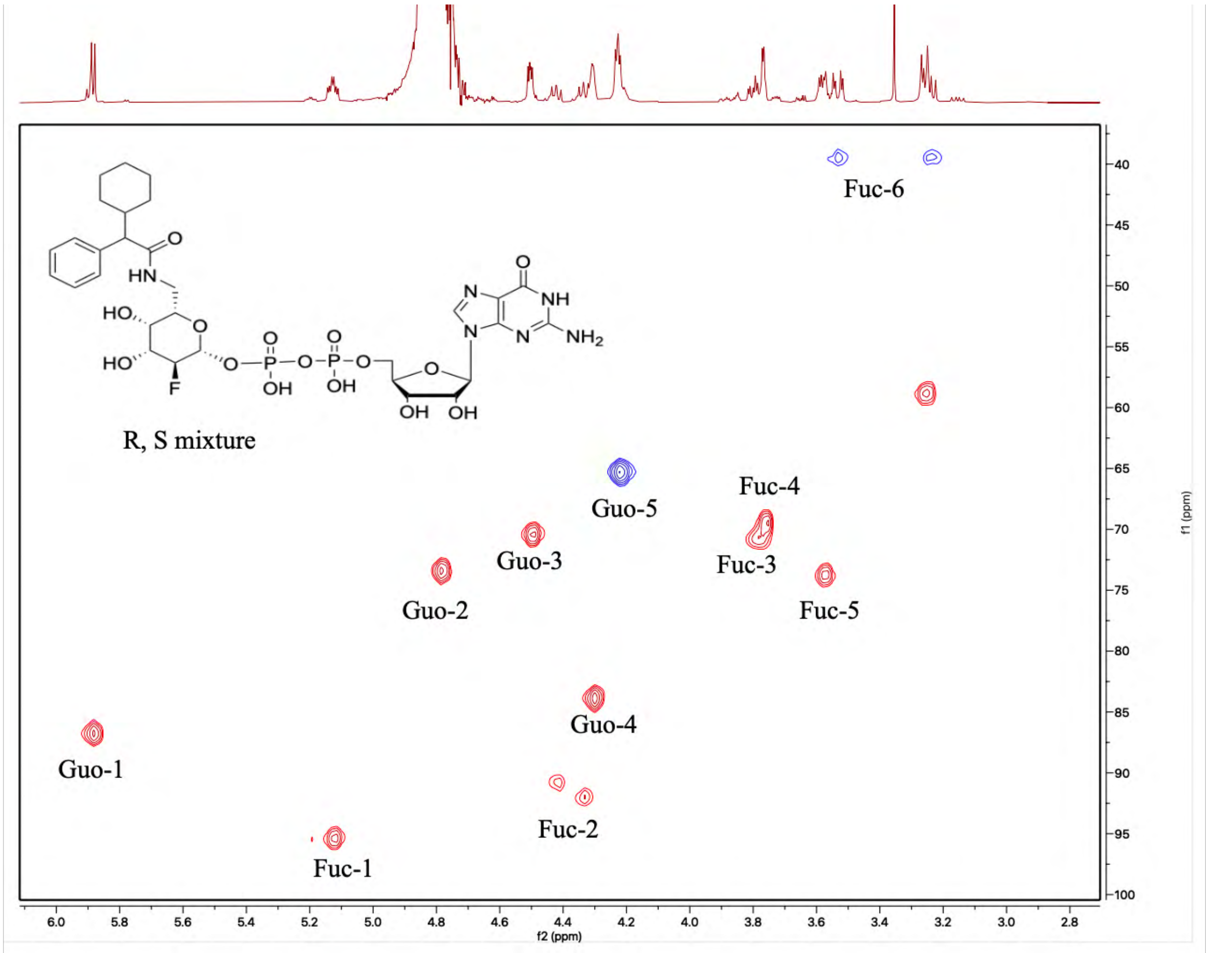


$^{13}C$ -$^{1}H$ HSQC spectrum of compound **2f** (sugar region).

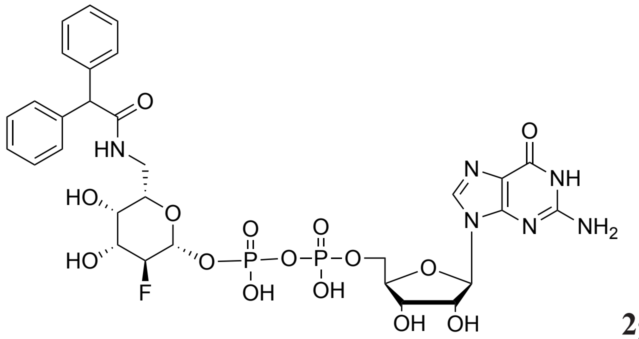

**2g**

**GDP-β-2-deoxy-2-fluoro-6-deoxy-6-*N*-diphenylacetamide-L-galactose**

[1]H NMR (600 MHz, Deuterium Oxide) δ 8.02 (s, 1H, $H_{Guo}$-8’), 7.41 – 7.21 (m, 10 H, Ar), 5.81 (d, *J* = 6.2 Hz, 1H, $H_{Guo}$-1), 5.19 (td, *J* = 8.0, 3.7 Hz, 1H, $H_{Fuc}$-1), 5.16 (s, 1H, CH), 4.71 (m, 1H, $H_{Guo}$-2), 4.44 (m, 1.5H, $H_{Guo}$-3, $H_{Fuc}$-2), 4.39 – 4.32 (m, 0.5H, $H_{Fuc}$-2), 4.25 (m, 1H, $H_{Guo}$-4), 4.17 (m, 2H, $H_{Guo}$-5), 3.91 – 3.83 (m, 2H, $H_{Fuc}$-3, $H_{Fuc}$-4), 3.74 (dd, *J* = 8.6, 4.1 Hz, 1H, $H_{Fuc}$-5), 3.61 (dd, *J* = 14.2, 4.2 Hz, 1H, $H_{Fuc}$-6a), 3.33 (dd, *J* = 14.2, 8.6 Hz, 1H, $H_{Fuc}$-6b).

ESI MS(m/z): $[M - H]^-$ calcd for $C_{30}H_{34}FN_6O_{15}P_2$, 799.1547; found 799.1193

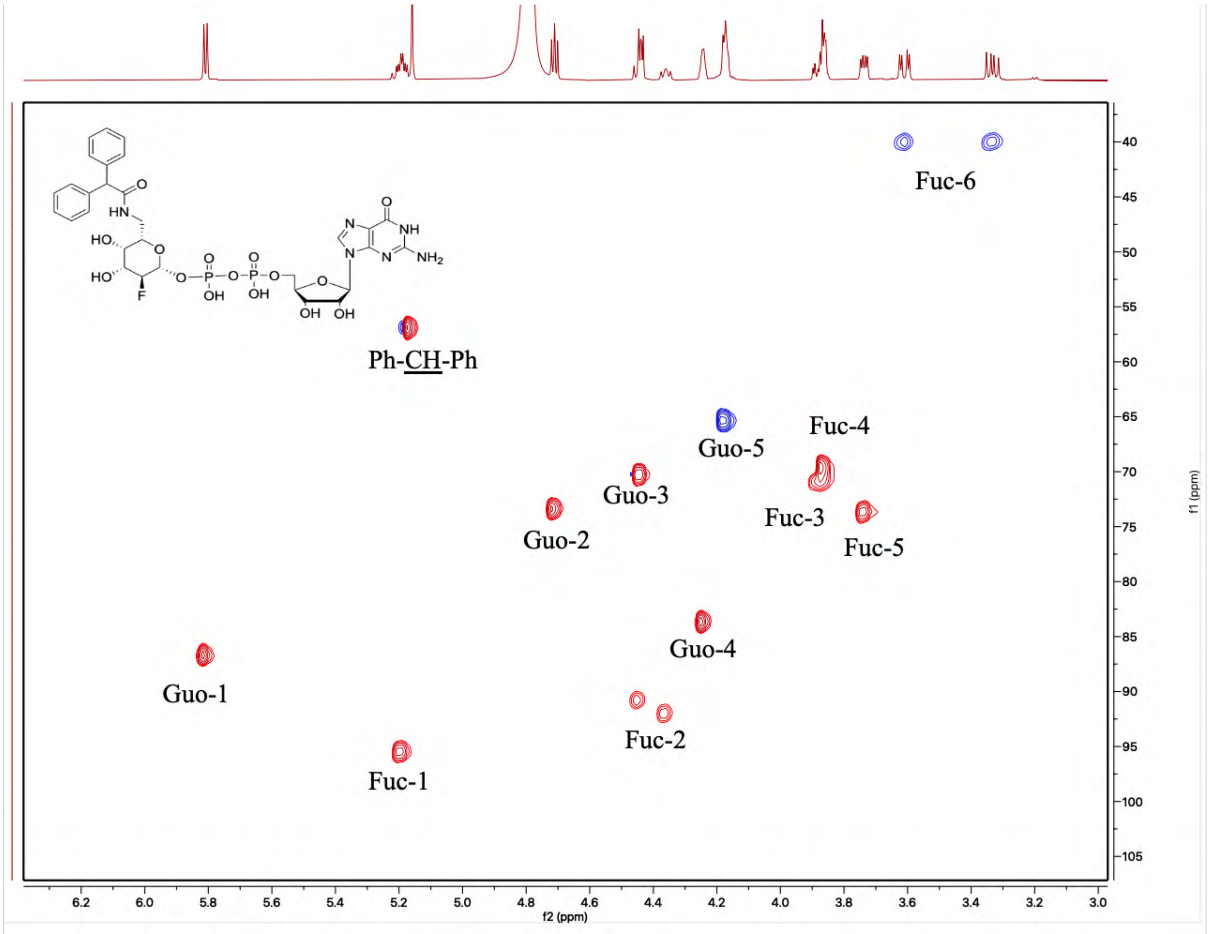


[13]C -[1]H HSQC spectrum of compound **2g** (sugar region).

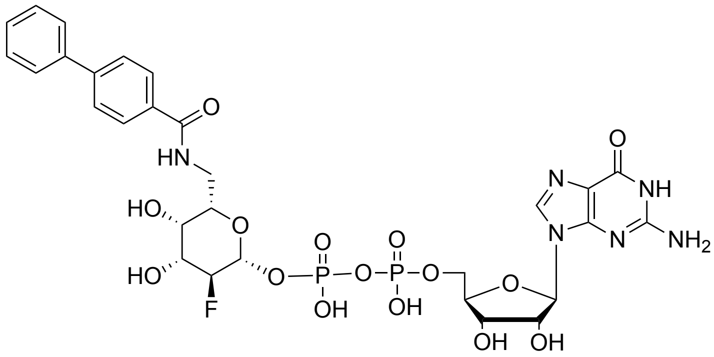

**2h**

**GDP-β-2-deoxy-2-fluoro-6-deoxy-6-*N*-(biphenyl-4-amide)-L-galactose**

$^1$H NMR (600 MHz, Deuterium Oxide) δ 7.91 – 7.79 (m, 3H, $H_{Guo}$-8', Ar), 7.58 (dd, *J* = 22.9, 7.6 Hz, 4H, Ar), 7.46 – 7.36 (m, 3H, Ar), 5.64 (d, *J* = 5.7 Hz, 1H, $H_{Guo}$-1), 5.25 (td, *J* = 7.6, 3.7 Hz, 1H, $H_{Fuc}$-1), 4.60 – 4.52 (m, 1.5H, $H_{Guo}$-2, $H_{Fuc}$-2), 4.47 (m, 0.5H, $H_{Fuc}$-2), 4.43 (m, 1H, $H_{Guo}$-3), 4.27 – 4.15 (m, 3H, $H_{Guo}$-4, $H_{Guo}$-5), 4.07 (t, *J* = 3.3 Hz, 1H, $H_{Fuc}$-4), 4.02 (ddd, *J* = 13.7, 9.5, 3.6 Hz, 1H, $H_{Fuc}$-3), 3.95 (dd, *J* = 9.8, 2.6 Hz, 1H, $H_{Fuc}$-5), 3.88 (dd, *J* = 14.2, 2.6 Hz, 1H, $H_{Fuc}$-6a), 3.50 (dd, *J* = 14.2, 9.7 Hz, 1H, $H_{Fuc}$-6b).

ESI MS(m/z): $[M - H]^-$ calcd for $C_{29}H_{32}FN_6O_{15}P_2$, 785.1390; found 785.1036

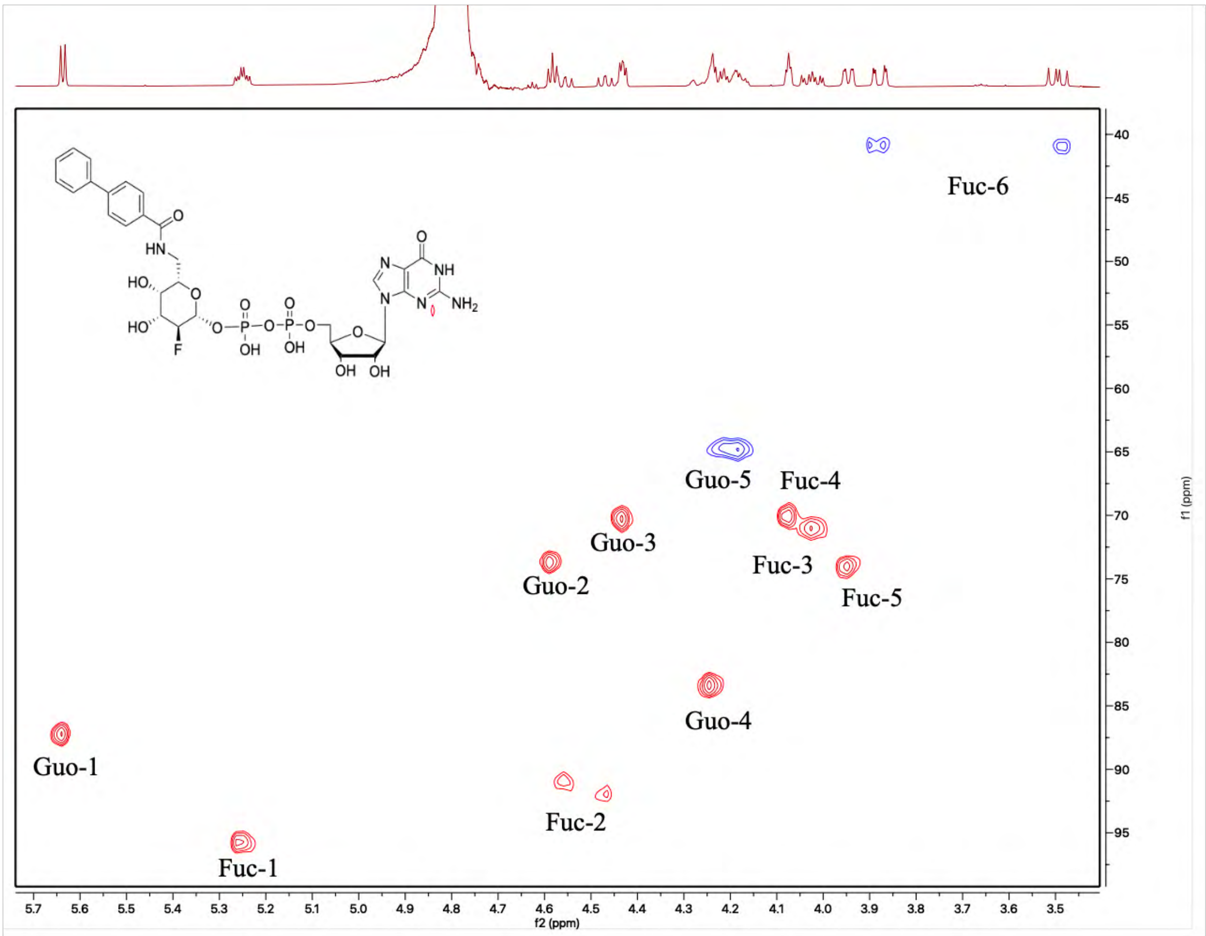


$^{13}$C -$^1$H HSQC spectrum of compound **2h** (sugar region).

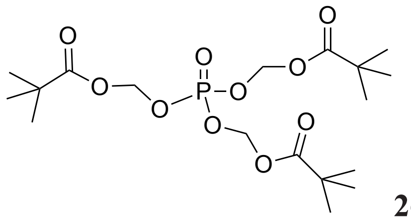

**26**

**Tris(pivaloyloxymethyl) (POM) phosphate**

Trimethyl phosphate (1.2 mL, 11 mmol) was dissolved in $CH_3CN$ (9 mL), and then NaI (4.8 g, 33 mmol) and chloromethylpivalate (6 mL, 41 mmol) were added. The reaction mixture was warmed to reflux and stirred for 30 h. Afterward, the reaction mixture was diluted with 150 mL of $Et_2O$ and washed with water. The organic layer was dried over $Na_2SO_4$ and concentrated under vacuum. Purification through silica gel column chromatography afforded **26** as a colorless oil (2.6 g, 55%).

$^1$H NMR (600 MHz, Chloroform-*d*) δ 5.66 (d, $J$ = 13.8 Hz, 6H, $CH_2$), 1.24 (s, 27H, $CH_3$). $^{13}$C NMR (151 MHz, Chloroform-*d*) δ 176.76, 82.92 (d, $J_{P\text{-}C}$ = 4.9 Hz), 38.89, 26.95.

ESI MS(m/z): $[M + Na]^+$ calcd for $C_{18}H_{33}NaO_{10}P$, 463.1709; found 463.1903

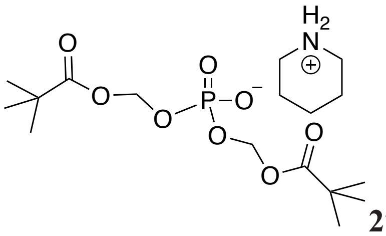

**27**

**Bis(pivaloyloxymethyl) (POM) phosphate piperidine salt**

Compound **26** (2.6 g, 6 mmol) was dissolved in 20 mL of piperidine and stirred at RT for 24 h. Afterward, the reaction mixture was concentrated under vacuum, and the crude product **27** was directly used for the next step. (95% yield)

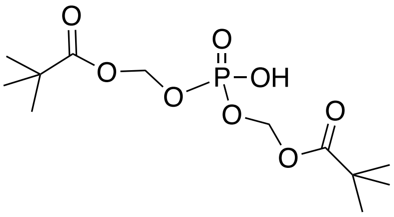

**28**

**Bis(pivaloyloxymethyl) (POM) phosphate**

Compound **27** was dissolved in MILIQ and loaded onto an $H^+$ resin column. It was then eluted with MilliQ-$H_2O$. The eluted solution was collected and concentrated to give compound **28** as a colorless oil (95% yield). Because compound **37** is acidic, it is not stable during storage and needs to be freshly prepared when used.

$^1$H NMR (600 MHz, Chloroform-*d*) δ 5.64 (d, *J* = 14.1 Hz, 4H, $CH_2$), 1.23 (d, *J* = 1.4 Hz, 18H, $CH_3$). $^{13}$C NMR (151 MHz, Chloroform-*d*) δ 176.98, 82.83 (d, $J_{P-C}$= 5.3 Hz), 38.88, 26.95.

ESI MS(m/z): $[M + Na]^+$ calcd for $C_{12}H_{23}NaO_8P$, 349.1028; found 349.1804

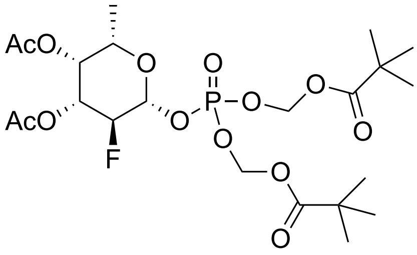


**5**

**2-Deoxy-2-fluoro-3,4-di-*O*-acetyl-β-1-(dipivaloyloxymethylphosphoryl)-L-fucopyranose**

Compound **3** (600 mg, 2 mmol) was dissolved in HBr (33% in AcOH, 2 mL) and stirred for 1 h at RT. Afterward, the mixture was concentrated *in vacuo*. The resulting residue was dissolved in 5 mL dry $CH_3CN$, and 500 mg of 3 Å molecular sieves were added. The mixture was stirred at RT for 10 min. Then, in one portion, silver carbonate (1.3 g, 4 mmol) and diPOM-phosphate, compound **28** (1.5 g, 4 mmol) were added. The reaction mixture was stirred at RT, under nitrogen gas protection, and in the dark. After 16 h, the mixture was diluted with ethyl acetate and then filtered through Celite. The filtrate was concentrated *in vacuo*. The residue was purified by silica gel chromatography, yielding **5** (725 mg, 65%) in white form.

$^1$H NMR (600 MHz, Chloroform-*d*) δ 5.73 – 5.62 (m, 4H, $CH_2$), 5.36 (m, 1H, H-1), 5.27 (m, 1H, H-4), 5.11 (ddd, *J* = 13.1, 9.9, 3.6 Hz, 1H, H-3), 4.61 (dd, *J* = 9.9, 7.6 Hz, 0.5H, H-2), 4.52 (dd, *J* = 9.9, 7.6 Hz, 0.5H, H-2), 3.95 (m, 1H, H-5), 2.16 (s, 3H, $CH_3$-Ac), 2.05 (s, 3H, $CH_3$-Ac), 1.23 (m, 21H, $CH_3$-Fuc, $CH_3$-POM,). $^{13}$C NMR (151 MHz, Chloroform-*d*) δ 176.73, 170.39, 169.95, 96.77 (dd, *J* = 24.2, 4.6 Hz, C-1), 87.72 (dd, *J* = 189.5, 9.2 Hz, C-2), 82.92, 71.02 (d, *J* = 18.6 Hz, C-3), 70.62 (C-5), 70.42 (d, *J* = 8.3 Hz, C-3), 38.86, 26.93, 26.90, 20.70, 20.66, 15.88. $^{19}$F NMR (565 MHz, Chloroform-*d*) δ -207.86 (dd, *J* = 51.8, 12.8 Hz).

ESI MS(m/z): $[M + Na]^+$ calcd for $C_{22}H_{36}FO_{13}PNa$, 581.1775; found 581.0457

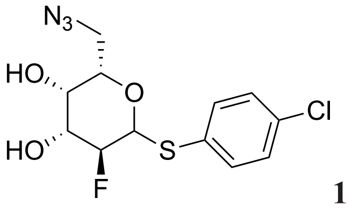


**18**

**Para-chlorothiophenyl 2-deoxy-2-fluoro-6-N$_3$-β-L-galactopyranoside**

Compound **13** (350 mg, 0.75 mmol) was dissolved in DMF (3 mL), and sodium azide (520 mg, 8 mmol) was added. The reaction was stirred for 16 h at 70 °C. The solvent was then removed *in vacuo*, and the residue was dissolved in ethyl acetate and filtered. The filtrate was concentrated *in vacuo*, and the resulting residue was purified by silica gel chromatography, affording **18** (174 mg, 70%) as a white solid.

$^1$H NMR (600 MHz, Chloroform-*d*) δ 7.53 (d, *J* = 8.5 Hz, 2H, Ar), 7.31 (d, *J* = 8.5 Hz, 2H, Ar), 4.60 (dd, *J* = 9.7, 1.7 Hz, 1H, H-1), 4.39 (t, *J* = 9.2 Hz, 0.5H, H-2), 4.30 (t, *J* = 9.2 Hz, 0.5H, H-2), 4.00 (m, 1H, H-4), 3.86 (ddt, *J* = 12.8, 8.1, 3.7 Hz, 1H, H-3), 3.67 (ddd, *J* = 19.5, 12.1, 7.4 Hz, 2H, H-6), 3.43 (dd, *J* = 12.3, 4.4 Hz, 1H, H-5), 2.66 (d, *J* = 4.4 Hz, 1H, OH-3), 2.44 (d, *J* = 2.9 Hz, 1H, OH-4).

$^{13}$C NMR (101 MHz, Chloroform-*d*) δ 135.33, 135.04, 129.55, 129.36, 89.17 (d, *J* = 183.4 Hz, C-2), 84.82 (d, *J* = 24.0 Hz, C-1), 77.36, 73.29 (d, *J* = 19.3 Hz, C-3), 69.81 (d, *J* = 8.6 Hz, C-4), 51.12.

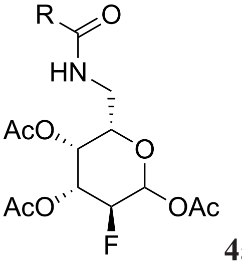


**4a**, **c**, **f**

**General procedure for the synthesis of 4a**, **c**, **f**

Compound **18** (1 equivalent) was dissolved in THF, followed by the addition of 1.1 equivalents of triphenylphosphine and a catalytic amount of water. The reaction mixture was stirred at RT for 12 h. Subsequently, an equal volume of water, relative to THF, was added to the reaction mixture, along with 1 equivalent of DIPEA and 1.5 equivalents of the corresponding active NHS ester. After stirring at RT for 6 h, the reaction was dried, diluted with ethyl acetate, and washed with brine. The organic layer was then concentrated under vacuum. The resulting residue was dissolved in pyridine and acetic anhydride, and the mixture was stirred at RT for 1 h. Afterward, the reaction mixture was dried again. The resulting residue was dissolved in an acetone and $H_2O$ (6:1) mixture, and then 10 equivalents of N-Bromosuccinimide were added. The reaction mixture was stirred for 12 h at RT under $N_2$. Subsequently, sat. aq. NaHCO3 was added, and the mixture was diluted with ethyl acetate and washed with brine. The organic phase was dried over $Na_2SO_4$ and concentrated *in vacuo*. The resulting residue was dissolved in pyridine and acetic anhydride, and the mixture was stirred at RT for 1 h. Afterward, the reaction mixture was dried again. Purification through silica gel column chromatography afforded **4a**, **4c**, and **4f** as colorless oils (50-60%).

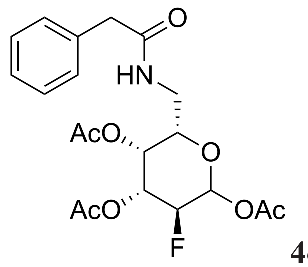


**4c**

**2-Deoxy-2-fluoro-6-*N*-benzylamide-per-*O*-acetyl-L-galactopyranoside**

$^{1}$H NMR (400 MHz, Chloroform-*d*) δ 7.49 – 7.16 (m, 16H, Ar), 6.37 (d, *J* = 4.0 Hz, 2.2H, αH-1), 5.70 (dd, *J* = 8.1, 4.1 Hz, 1H, βH-1), 5.64 (m, 3H, NH), 5.41 – 5.29 (m, 5.4H, αH-4, αH-3, βH-4), 5.12 (ddd, *J* = 12.8, 10.0, 3.3 Hz, 1H, βH-3), 4.91 (dd, *J* = 9.1, 4.0 Hz, 1.1H, αH-2), 4.79 (dd, *J* = 10.0, 4.1 Hz, 1.1H, αH-2), 4.66 (m, 0.5H, βH-2), 4.53 (m, 0.5H, βH-2), 4.11 (t, *J* = 6.9 Hz, 2.2H, αH-5), 3.87 (dd, *J* = 8.1, 5.5 Hz, 1H, βH-5), 3.55 (d, *J* = 10.3 Hz, 6.4H, $CH_2$-Ar), 3.47 (m, 1H, βH-6), 3.36 (dt, *J* = 13.2, 6.5 Hz, 2.2H, αH-6), 3.26 – 3.10 (m, 3.2H, αH-6, βH-6), 2.37 – 1.96 (m, 29H, $CH_3$-Ac).

$^{13}$C NMR (151 MHz, Chloroform-*d*) δ 171.48, 171.44, 170.39, 170.27, 170.03, 169.80, 169.08, 168.78, 134.59, 129.69, 129.51, 129.24, 127.60, 127.56, 91.77 (d, *J* = 24.7 Hz, βC-1), 89.02 (d, *J* = 22.3 Hz, αC-1), 86.87 (d, *J* = 188.4 Hz, βC-2), 84.32 (d, *J* = 191.3 Hz, αC-2), 72.52, 71.08 (d, *J* = 18.8 Hz, βC-3), 69.26, 68.90 (d, *J* = 7.8 Hz, αC-4), 68.35 (d, *J* = 8.3 Hz, βC-4), 68.34 (d, *J* = 19.0 Hz, αC-3), 43.84, 43.80, 38.95, 38.93, 21.02, 20.92, 20.78, 20.70, 20.68.

ESI MS(m/z): $[M + Na]^+$ calcd for $C_{20}H_{24}FNO_8Na$, 448.1384; found 448.0696

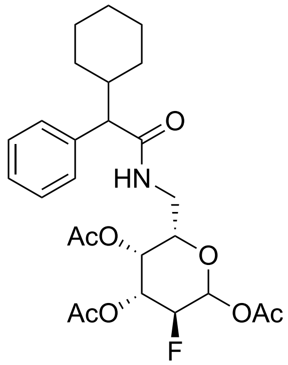


**4f**

**2-Deoxy-2-fluoro-6-*N*-(2-cyclohexyl-phenylacetamide)-per-*O*-acetyl-L-galactopyranoside**

$^{1}$H NMR (600 MHz, Chloroform-*d*) (R, S mixture, β, α mixture) δ 7.31 – 7.28 (m, 12H, Ar), 7.23 (m, 4H, Ar), 7.16 (m, 1.5H, Ar), 6.40 (d, *J* = 3.9 Hz, 1H, αH-1), 6.34 (d, *J* = 4.0 Hz, 1H, αH-1), 5.81 (m, 3.2H, NH), 5.70 (ddd, *J* = 10.8, 8.0, 4.2 Hz, 1.5H, βH-1), 5.41 (t, *J* = 2.8 Hz, 1H, αH-4), 5.38 – 5.28 (m, 3.5H, βH-4, αH-3), 5.22 (t, *J* = 2.6 Hz, 1H αH-4), 5.10 (dddd, *J* = 13.1, 9.8, 5.1, 3.5 Hz, 1.5H, βH-3), 4.89 (ddd, *J* = 10.3, 6.2, 4.1 Hz, 1H, αH-2), 4.84 – 4.77 (m, 1H, αH-2), 4.65 (ddd, *J* = 12.9, 9.9, 8.0 Hz, 1H, βH-2), 4.56 (ddd, *J* = 12.9, 9.8, 8.0 Hz, 1H, βH-2), 4.11 (dd, *J* = 8.4, 4.8 Hz, 1H, αH-5), 4.00 (m, 1H, αH-5), 3.82 (m, 1.5H, βH-5), 3.4 (m, 3.5H, αH-6, βH-6), 3.19 (m, 2.5H, αH-6, βH-6), 3.05 (ddd, *J* = 13.8, 8.4, 5.1 Hz, 1H, αH-6), 2.89 (dd, *J* = 10.2, 2.5 Hz, 1.5H, CH), 2.86 (dd, *J* = 10.4, 1.9 Hz, 2H, CH), 2.23 – 1.90 (m, 31.5H, $CH_3$-Ac), 1.87 – 1.53 (m, 16H, $CH_2$), 1.40 – 1.26 (m, 8H, $CH_2$), 1.21 – 1.07 (m, 7H, $CH_2$), 1.03 – 0.89 (m, 4H, $CH_2$)

$^{13}$C NMR (151 MHz, Chloroform-*d*) δ 173.76, 173.74, 173.69, 173.60, 170.54, 170.39, 170.36, 170.27, 169.90, 169.87, 169.71, 169.63, 169.20, 169.14, 168.82, 168.76, 138.59, 129.18, 128.67, 128.65, 128.52, 128.48, 128.43, 128.38, 127.28, 127.21, 91.81 (d, *J* = 24.6 Hz, αC-1), 89.12 (d, *J* = 22.3 Hz, βC-1), 88.92 (d, *J* = 22.3 Hz, βC-1), 86.98 (d, *J* = 182.9 Hz, βC-2), 84.38 (d, *J* = 191.5 Hz, αC-2), 72.79 (βC-5), 72.61 (βC-5), 71.03 (d, *J* = 19.5 Hz, βC-3), 69.85 (αC-5), 69.64 (αC-5), 69.26 (d, *J* = 7.7 Hz, αC-4), 68.66 (d, *J* = 8.9 Hz, βC-4), 68.31 (d, *J* = 16.5 Hz, αC-3), 68.19 (d, *J* = 16.7 Hz, αC-3), 60.97, 60.90, 40.65, 40.54, 40.52, 39.20, 39.02, 38.99, 38.83, 32.19, 32.08, 31.88, 30.77, 30.74, 29.42, 26.47, 26.22, 26.15, 26.11, 20.98, 20.96, 20.95, 20.83, 20.76, 20.72, 20.66, 20.62.

ESI MS(m/z): $[M + Na]^+$ calcd for $C_{26}H_{34}FNO_8Na$, 530.2166; found 530.1304

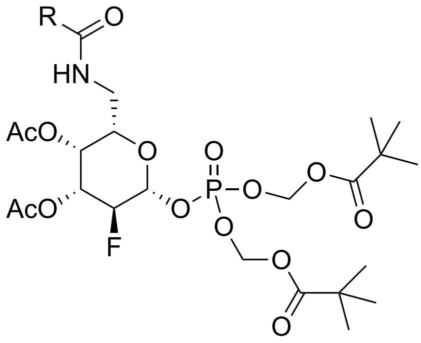

**6a**, **6c**, **6f**

**General procedure for the synthesis of prodrugs 6a**, **c**, **f**

Per acetylated fucose analogues **4a**, **4c**, and **4f** (1 equivalent, 0.1 mmol each) was dissolved in HBr (2 mL, 33% in AcOH) and stirred for 1 h at RT. Afterward, the mixture was concentrated *in vacuo*. The resulting residue was dissolved in dry $CH_3CN$, and 3 Å molecular sieves were added. The mixture was stirred at RT for 10 min. Then, in one portion, 2 equivalents of silver carbonate (55 mg, 0.2 mmol) and 2 equivalents of diPOM-phosphate (**28)** (65 mg, 0.2 mmol) were added. The reaction mixture was stirred at RT, under nitrogen gas protection, and in the dark. After 12 h, the mixture was diluted with ethyl acetate and then filtered through Celite. The filtrate was concentrated *in vacuo*. The residue was purified by silica gel chromatography, yielding the corresponding products **6a**, **6c**, and **6f** (60-80%) as white solids.

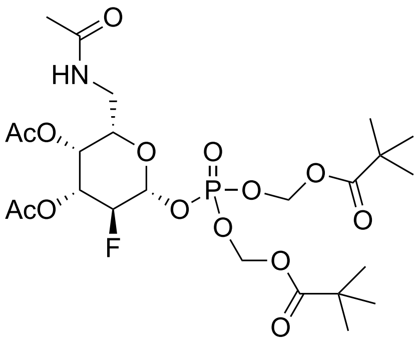

**6a**

**2-Deoxy-2-fluoro-3,4-di-*O*-acetyl-6-deoxy-6-*N*-acetamide-β-1 (dipivaloyloxymethylphosphoryl)-L-galactopyranose**

$^1$H NMR (600 MHz, Chloroform-*d*) δ 6.41 (dd, *J* = 7.8, 4.5 Hz, 1H, NH), 5.74 – 5.65 (m, 4H, $CH_2$), 5.40 (t, *J* = 2.6 Hz, 1H, H-4), 5.33 (td, *J* = 7.4, 4.0 Hz, 1H, H-1), 5.11 (m, 1H, H-1), 4.61 (dd, *J* = 9.9, 7.6 Hz, 0.5H, H-2), 4.52 (dd, *J* = 9.9, 7.6 Hz, 1H, H-2), 3.89 (dd, *J* = 8.5, 4.3 Hz, 1H, H-5), 3.62 (ddd, *J* = 14.2, 7.8, 4.3 Hz, 1H, H-6a), 3.14 (m, 1H, H-6b), 2.15 (s, 3H, $CH_3$-Ac), 2.05 (s, 3H, $CH_3$-Ac), 2.00 (s, 3H, $CH_3$-Ac), 1.25 (d, *J* = 1.8 Hz, 18H, $CH_3$-POM).

$^{13}C$ NMR (151 MHz, Chloroform-*d*) δ 177.10, 177.06, 170.76, 170.10, 169.71, 96.82 (d, *J* = 24.7 Hz, C-1), 87.74 (d, *J* = 190.1, C-2), 82.97, 82.95, 73.29, 70.63 (d, *J* = 18.5 Hz, C-3), 68.48 (d, *J* = 7.8 Hz, C-4), 39.19, 38.96, 38.93, 26.96, 26.93, 23.15, 20.66.

ESI MS(m/z): $[M + Na]^+$ calcd for $C_{24}H_{39}FNO_{14}PNa$, 638.1990; found 638.0991

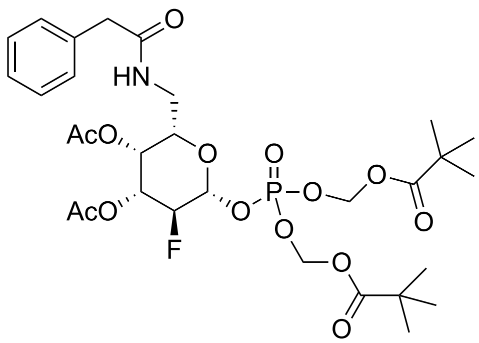


**6c**

**2-Deoxy-2-fluoro-3,4-di-*O*-acetyl-6-deoxy-6-*N*-phenylacetamide-β-1 (dipivaloyloxymethylphosphoryl)-L-galactopyranose**

$^{1}H$ NMR (600 MHz, Chloroform-*d*) δ 7.36 – 7.32 (m, 2H, Ar), 7.30 – 7.27 (m, 3H, Ar), 6.26 (dd, *J* = 7.6, 4.8 Hz, 1H, NH), 5.67 (d, *J* = 13.7 Hz, 2H, CH2), 5.63 (dq, *J* = 13.1, 5.0 Hz, 2H, CH2), 5.34 (m, 1H, H-4), 5.31 (td, *J* = 7.4, 3.9 Hz, 1H, H-1), 5.08 (ddd, *J* = 13.0, 9.9, 3.5 Hz, 1H, H-3), 4.59 (dd, *J* = 9.9, 7.6 Hz, 0.5H, H-2), 4.50 (dd, *J* = 9.9, 7.6 Hz, 0.5H, H-2), 3.87 (dd, *J* = 7.6, 5.3 Hz, 1H, H-5), 3.60 – 3.54 (m, 3H, $CH_2$, H-6a), 3.18 – 3.12 (m, 1H, H-6b), 2.11 (s, 3H, $CH_3$-Ac), 2.05 (s, 3H, $CH_3$-Ac), 1.24 (d, *J* = 2.9 Hz, 18H, $CH_3$-POM).

$^{13}C$ NMR (151 MHz, Chloroform-*d*) δ 176.97, 171.66, 170.09, 169.69, 134.88, 129.50, 129.09, 127.38, 96.76 (d, *J* = 24.4 Hz, C-1), 87.74 (d, *J* = 192.5 Hz, C-2), 82.98, 82.95, 73.08 (C-5), 70.59 (d, *J* = 19.4 Hz, C-3), 68.33 (d, *J* = 8.0 Hz, C-4), 43.65, 39.24, 38.93, 38.92, 29.86, 26.96, 26.93, 20.66, 20.65.

ESI MS(m/z): $[M + Na]^+$ calcd for $C_{30}H_{43}FNO_{14}PNa$, 714.2303; found 714.1312

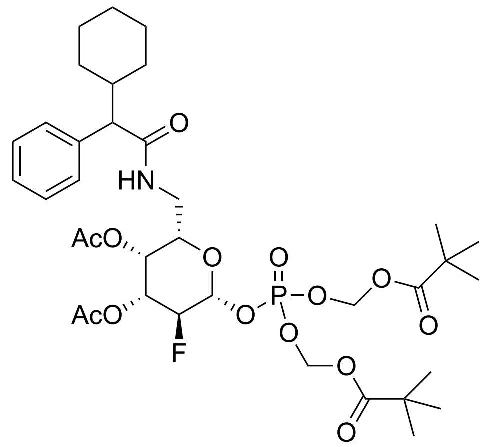

**6f**

**2-Deoxy-2-fluoro-3,4-di-*O*-acetyl-6-deoxy-6-*N*-(2-cyclohexyl-phenylacetamide)-β-1 (dipivaloyloxymethylphosphoryl)-L-galactopyranose**

$^1$H NMR (400 MHz, Chloroform-*d*) (R, S mixture) δ 7.33 – 7.16 (m, 10H, Ar), 6.49 (q, *J* = 5.7 Hz, 2H, NH), 5.72 – 5.56 (m, 8H, $CH_2$), 5.28 (m, 3H, H-4, H-1), 5.18 (td, *J* = 7.5, 4.0 Hz, 1H, H-1), 5.03 (m, 2H, H-3), 4.57 (dd, *J* = 9.8, 7.6 Hz, 1H, H-2), 4.45 (dd, *J* = 9.8, 7.6 Hz, 1H, H-2), 3.83 (dd, *J* = 8.7, 4.0 Hz, 1H, H-5), 3.72 (dd, *J* = 8.5, 4.4 Hz, 1H, H-5), 3.51 (m, 2H, H-6), 3.16 (ddd, *J* = 13.5, 8.3, 4.5 Hz, 1H, H-6), 3.03 (ddd, *J* = 13.6, 8.5, 4.5 Hz, 1H, H-6), 2.96 (d, *J* = 10.4 Hz, 2H, CH), 2.06 (m, 12H, $CH_3$-Ac), 1.83 – 1.51 (m, 11H, $CH_3$-POM), 1.39 – 1.22 (m, 20H, CH, $CH_3$-POM), 1.15 – 0.90 (m, 7H, $CH_3$-POM).

$^{13}$C NMR (151 MHz, Chloroform-*d*) δ 177.08, 177.04, 177.02, 174.07, 170.14, 170.01, 169.58, 169.56, 139.07, 138.86, 128.59, 128.53, 128.48, 127.14, 127.12, 96.79 (d, *J* = 24.7 Hz, C-1), 87.82 (d, *J* = 189.9 Hz, C-2), 82.95, 82.92, 73.43, 73.23, 70.50 (d, *J* = 18.7 Hz, C-3), 68.54 (d, *J* = 8.1 Hz, C-4), 60.49, 60.43, 40.56, 40.48, 39.37, 39.10, 38.95, 38.92, 32.18, 30.81, 26.98, 26.93, 26.55, 26.51, 26.21, 26.17, 26.13, 20.62, 20.56.

ESI MS(m/z): $[M + Na]^+$ calcd for $C_{36}H_{53}FNO_{14}PNa$, 796.3085; found 796.1873

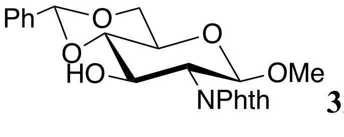

**32**

**Methyl 4,6-benzylidene-2-deoxy-2-*N*-phthalimido-β-D-glucopyranoside**

Compound **31** (1.3 g, 4 mmol) was dissolved in $CH_3CN$ (10 mL), and benzaldehyde dimethyl acetal (0.4 mL, 6 mmol) and p-TsOH·$H_2O$ (38 mg, 0.2 mmol) were then added. The reaction mixture was stirred at RT for 4 h. Then, triethylamine was added to quench the reaction. The solution was dried and chromatographed to give compound **32** (1.6 g, 100%) as a white solid.

$^1$H NMR (600 MHz, Chloroform-*d*) δ 7.86 (m, 2H, Ar), 7.73 (m, 2H, Ar), 7.50 (m, 2H, Ar),

7.38 (m, 3H, Ar), 5.57 (d, *J* = 5.5 Hz, 1H, PhCH), 5.20 (dd, *J* = 8.5, 5.7 Hz, 1H, H-1), 4.66 – 4.59 (m, 1H, H-3), 4.41 (dt, *J* = 10.7, 5.2 Hz, 1H, H-6a), 4.27 – 4.19 (m, 1H, H-2), 3.84 (td, *J* = 10.1, 5.3 Hz, 1H, H-6b), 3.69 – 3.57 (m, 2H, H-4, H-5), 3.44 (d, *J* = 5.9 Hz, 3H, OMe). $^{13}$C NMR (151 MHz, Chloroform-*d*) δ 137.08, 134.28, 131.86, 129.54, 128.55, 126.44, 123.71, 102.12, 99.97, 82.43, 68.85, 68.77, 66.25, 57.26, 56.56.

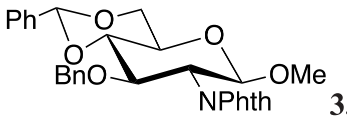

**33**

**Methyl 4,6-benzylidene-3-*O*-benzyl-2-deoxy-2-*N*-phthalimido-β-D-glucopyranoside**

Compound **32** (1.6 g, 4 mmol) was dissolved in DMF (20 mL) and cooled to 0 ºC. Then, sodium hydride (60% dispersion in mineral oil, 0.4 g, 10 mmol) was added in portions. After stirring for 20 min, Benzyl bromide (2 mL, 16 mmol) was added dropwise. Then the reaction mixture was allowed to warm back to RT and stirred for 1 h. Afterward, MeOH was added to quench the reaction. The reaction mixture was diluted with ethyl acetate and washed with brine. The organic layer was collected and dried. Purification by silica gel chromatography afforded **33** (1.1 g, 50%) as a white solid.

$^{1}$H NMR (600 MHz, Chloroform-*d*) δ 7.71 (d, *J* = 5.3 Hz, 3H, Ar), 7.53 (dd, *J* = 8.1, 1.6 Hz, 2H, Ar), 7.43 – 7.33 (m, 4H, Ar), 7.03 – 6.97 (m, 2H, Ar), 6.93 (t, *J* = 7.3 Hz, 1H, Ar), 6.88 (m, 2H, Ar), 5.63 (s, 1H, PhCH), 5.13 (d, *J* = 8.5 Hz, 1H, H-1), 4.80 (d, *J* = 12.3 Hz, 1H, $CH_2$), 4.50 (d, *J* = 12.4 Hz, 1H, $CH_2$), 4.42 (ddd, *J* = 10.4, 6.8, 1.9 Hz, 2H, H-6a, H-3), 4.21 (dd, *J* = 10.4, 8.5 Hz, 1H, H-2), 3.87 (t, *J* = 10.3 Hz, 1H, H-6b), 3.82 (t, *J* = 9.2 Hz, 1H, H-4), 3.65 (td, *J* = 9.8, 5.0 Hz, 1H, H-5), 3.40 (s, 3H, OMe). $^{13}$C NMR (151 MHz, Chloroform-*d*) δ 138.02, 137.45, 133.98, 131.80, 129.18, 128.45, 128.17, 127.53, 126.19, 123.50, 101.46, 99.97, 83.27, 74.70, 74.23, 68.94, 66.18, 57.16, 55.83.

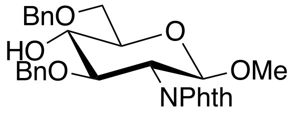

**34**

**Methyl 3,6-*O*-dibenzyl-2-deoxy-2-*N*-phthalimido-β-D-glucopyranoside**

Compound **33** (1 g, 2 mmol) was dissolved in anhydrous DCM (10 mL) and cooled to 0 ºC. Triethyl silane (3.3 mL, 20 mmol) and trifluoroacetic acid (1.5 mL, 20 mmol) were added sequentially. The reaction mixture was stirred at 0 ºC for 1 h. Afterward, the solution was washed with water and sat. aq. $NaHCO_3$ and brine. The organic layer was collected, dried over

$Na_2SO_4$, concentrated, and purified by column chromatography, giving compound **34** (0.46 g, 44%) as a colorless oil.

$^1$H NMR (600 MHz, Chloroform-*d*) δ 7.69 (s, 4H, Ar), 7.42 – 7.29 (m, 5H, Ar), 7.09 – 7.02 (m, 2H, Ar), 7.00 – 6.90 (m, 3H, Ar), 5.07 (d, *J* = 7.8 Hz, 1H, H-1), 4.74 (d, *J* = 12.2 Hz, 1H, $CH_2$-Bn), 4.66 (d, *J* = 11.9 Hz, 1H, $CH_2$-Bn), 4.60 (d, *J* = 11.9 Hz, 1H, $CH_2$-Bn), 4.53 (d, *J* = 12.2 Hz, 1H, $CH_2$-Bn), 4.23 (dd, *J* = 10.7, 8.5 Hz, 1H, H-3), 4.14 (dd, *J* = 10.8, 8.4 Hz, 1H, H-2), 3.87 – 3.78 (m, 3H, H-6, H-4), 3.65 (dt, *J* = 9.8, 5.0 Hz, 1H, H-5), 3.38 (s, 3H, OMe), 2.89 (d, *J* = 2.5 Hz, 1H, OH).

$^{13}$C NMR (151 MHz, Chloroform-*d*) δ 138.30, 137.72, 128.69, 128.31, 128.09, 128.01, 127.98, 127.58, 99.36, 78.81, 74.72, 74.48, 73.95, 73.57, 70.86, 56.85, 55.38.

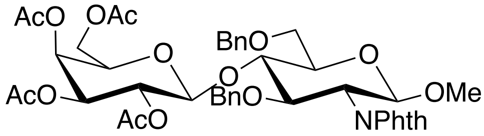

**35**

**Methyl [peracetyl-β-D-galactopyranosyl]-(1→4)-3,6-*O*-dibenzyl-2-deoxy-2-*N*-phthalimido-β-D-glucopyranoside**

Compound **34** (0.46 g, 1 mmol) and compound **30** (0.67 g, 1.4 mmol) were dissolved in anhydrous DCM (5 mL), and 4 Å molecular sieve (500 mg) was added. The reaction mixture was stirred at RT for 10 min and then cooled to -20 $^o$C. Trimethylsilyl trifluoromethanesulfonate (33 μL, 0.2 mmol) was added, and stirring continued for 15 min. Afterward, the reaction was quenched with triethylamine and filtered through Celite. The solution was concentrated under vacuum. Purification by silica gel chromatography afforded **35** (0.5 g, 55%) as a colorless oil.

$^1$H NMR (600 MHz, Chloroform-*d*) δ 7.78 – 7.59 (m, 4H, Ar), 7.41 (d, *J* = 6.1 Hz, 4H, Ar), 7.34 (m, 1H, Ar), 7.01 (d, *J* = 6.5 Hz, 2H, Ar), 6.87 (m, 3H, Ar), 5.26 (d, *J* = 3.5 Hz, 1H, $H_{Gal}$-4), 5.14 (dd, *J* = 10.4, 8.0 Hz, 1H, $H_{Gal}$-2), 5.01 (d, *J* = 8.5 Hz, 1H, $H_{Glcl}$-1), 4.86 – 4.81 (m, 2H, $H_{Gal}$-3, $PhCH_2$), 4.79 (d, *J* = 12.3 Hz, 1H, $PhCH_2$), 4.58 (d, *J* = 8.0 Hz, 1H, $H_{Gal}$-1), 4.50 (d, *J* = 12.1 Hz, 1H, $PhCH_2$), 4.42 (d, *J* = 12.3 Hz, 1H, $PhCH_2$), 4.24 (dd, *J* = 10.7, 8.5 Hz, 1H, $H_{Glcl}$-3), 4.13 (dd, *J* = 10.8, 8.5 Hz, 1H, $H_{Glcl}$-2), 4.10 – 4.05 (m, 1H, $H_{Glcl}$-4), 4.00 – 3.91 (m, 2H, $H_{Gal}$-6), 3.82 – 3.76 (m, 2H, $H_{Glcl}$-6), 3.63 (t, *J* = 6.9 Hz, 1H, $H_{Gal}$-5), 3.53 (dt, *J* = 9.9, 2.5 Hz, 1H, $H_{Glcl}$-5), 3.38 (s, 3H, OMe), 2.06 (s, 3H, $CH_3$-Ac), 2.02 (s, 6H, $CH_3$-Ac), 1.97 (s, 3H, $CH_3$-Ac). $^{13}$C NMR (151 MHz, Chloroform-*d*) δ 170.48, 170.39, 170.21, 169.35, 138.67, 138.00, 128.77, 128.25, 128.22, 128.04, 127.96, 127.22, 100.46, 99.40, 78.15, 76.82, 74.91,

74.57, 73.79, 71.13, 70.57, 69.65, 67.59, 67.03, 60.85, 56.82, 55.64, 20.96, 20.83, 20.77, 20.73.
ESI MS(m/z): [M + Na]+ calcd for $C_{43}H_{47}NO_{16}Na$, 856.2793; found 856.4816

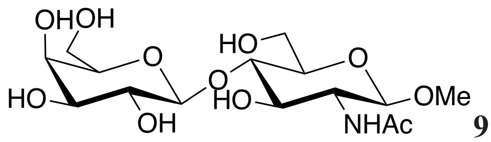

**9**

**Methyl [β-D-galactopyranosyl]-(1→4)- 2-deoxy-2-acetamido-β-D-glucopyranoside**

Compound **35** (0.5 g, 0.6 mmol) was dissolved in MeOH (10 mL), and freshly prepared sodium methoxide was added dropwise until the pH reached approximately 10. The reaction mixture was stirred at RT for 30 min, and then it was neutralized with $H^+$ resin. The solution was filtered through Celite and concentrated under vacuum. The resulting residue was dissolved in ethanol (40 mL), and ethylenediamine (1.3 mL, 20 mmol) was added. The reaction mixture was refluxed for 5 h. Afterward, the solution was concentrated *in vacuo*. The residue was dissolved in methanol (10 mL), and acetic anhydride (0.5 mL, 5 mmol) was added. The reaction mixture was stirred at RT for 6 h. Afterward, the reaction was concentrated under vacuum. The resulting residue was dissolved in a mixture of methanol (5 mL) and water (5 mL) and hydrogenated in an $H_2$ atmosphere with 10% activated $Pd(OH)_2/C$ (30 mg) as a catalyst for 3 h. Then, the solution was filtered through Celite and concentrated under vacuum. Purification by silica gel chromatography afforded **9** (252 mg, 95%) as a white powder.

$^1$H NMR (600 MHz, Methanol-$d_4$) δ 4.38 (d, $J$ = 7.6 Hz, 1H, $H_{Gal}$-1), 4.31 (d, $J$ = 8.4 Hz, 1H, $H_{Glc}$-4), 3.92 (dd, $J$ = 12.2, 2.5 Hz, 1H, $H_{Glc}$-6a), 3.86 (dd, $J$ = 12.1, 4.3 Hz, 1H, $H_{Glc}$-6b), 3.81 (d, $J$ = 3.3 Hz, 1H, $H_{Gal}$-4), 3.76 (dd, $J$ = 11.5, 7.6 Hz, 1H, $H_{Gal}$-6a), 3.72 (dd, $J$ = 10.1, 8.4 Hz, 1H, $H_{Glc}$-2), 3.68 (dd, $J$ = 11.6, 4.5 Hz, 1H, $H_{Gal}$-6b), 3.62 – 3.60 (m, 2H, $H_{Glc}$-3, $H_{Glc}$-4), 3.58 (dd, $J$ = 7.8, 4.4 Hz, 1H, $H_{Gal}$-5), 3.53 (dd, $J$ = 9.7, 7.6 Hz, 1H, $H_{Gal}$-2), 3.48 (dd, $J$ = 9.7, 3.3 Hz, 1H, $H_{Gal}$-3), 3.46 (s, 3H, OMe), 3.44 – 3.38 (m, 1H, $H_{Glc}$-5), 1.97 (s, 3H, NHAc). $^{13}$C NMR (151 MHz, Methanol-$d_4$) δ 173.61, 105.09, 103.63, 80.83, 77.17, 76.60, 74.83, 74.34, 72.61, 70.34, 62.53, 61.92, 57.02, 56.51, 22.91.

ESI MS(m/z): [M + Na]+ calcd for $C_{15}H_{27}NO_{11}Na$, 420.1482; found 419.6008

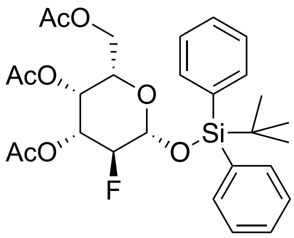

**19**

**2-Deoxy-2-fluoro-per-acetyl-1-*O*-(tert-butyldiphenylsilyl)-L-galactopyranose**

Compound **10** (200 mg, 0.8 mmol) was dissolved in a mixture of nitromethane and water (4:1), then Selectfluor (348 mg, 1 mmol) was added. The reaction mixture was placed in a microwave reactor and heated to 100 °C for 10 min, then the reaction mixture was diluted with ethyl acetate and washed with brine. The organic layer was dried over $Na_2SO_4$ and evaporated. The resulting residue was dissolved in DMF (2 mL) and cooled to 0 °C, tert-butyl diphenyl-silyl chloride (318 µL, 1.2 mmol) and imidazole (167 mg, 2.4 mmol) were added. The reaction mixture was stirred at 0 °C for 1 h and then allowed to warm to RT for 12 h. Afterwards, the reaction mixture was diluted with ethyl ether and washed with water. The organic layer was dried. Purification through silica gel column chromatography afforded **19** as a white solid (205 mg, 45%).

$^1$H NMR (600 MHz, Chloroform-*d*) δ 7.75 – 7.72 (m, 2H, Ar), 7.71 – 7.67 (m, 2H, Ar), 7.46 – 7.42 (m, 2H, Ar), 7.41 – 7.37 (m, 4H, Ar), 5.33 (ddd, *J* = 3.7, 2.5, 1.2 Hz, 1H, H-4), 4.96 (m, 1H, H-3), 4.69 (dd, *J* = 7.4, 3.3 Hz, 1H, H-1), 4.64 (dd, *J* = 9.7, 7.4 Hz, 0.5H, H-2), 4.55 (dd, *J* = 9.9, 7.4 Hz, 0.5H, H-2), 4.04 (d, *J* = 6.9 Hz, 2H, H-6), 3.65 (td, *J* = 6.7, 1.2 Hz, 1H, H-5), 2.16 (s, 3H, $CH_3$-Ac), 2.03 (s, 3H, $CH_3$-Ac), 1.97 (s, 3H, $CH_3$-Ac), 1.11 (s, 9H, $CH_3$-POM).

$^{13}$C NMR (151 MHz, Chloroform-*d*) δ 170.47, 170.30, 170.12, 135.97, 135.94, 132.72, 132.41, 130.22, 130.14, 127.86, 127.73, 95.77 (d, *J* = 22.6 Hz, C-1), 89.96 (d, *J* = 187.0 Hz, C-2), 71.14 (d, *J* = 18.8 Hz, C-3), 70.78, 68.05 (d, *J* = 8.1 Hz, C-4), 61.38, 26.80, 20.78, 20.75, 20.72, 19.41.

ESI MS(m/z): $[M + Na]^+$ calcd for $C_{28}H_{35}FNaO_8Si$, 569.1983; found 569.1436

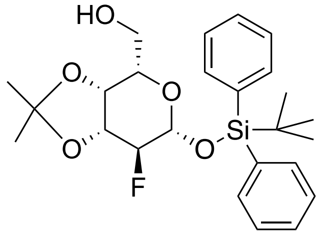

**20**

**2-Deoxy-2-fluoro-3,4-*O*-isopropylidene-1-*O*-(tert-butyldiphenylsilyl)-L-galactopyranose**

Compound **19** (150 mg, 0.27 mmol) was dissolved in methanol, and freshly prepared sodium methoxide was added dropwise until the pH of the reaction mixture reached 10. After stirring at RT for 0.5 h, the reaction was neutralized with H+ resin and filtered. The filtrate was concentrated under vacuum and then dissolved in acetone (5 mL). To this solution, p-Toluenesulfonic acid (8 mg, 0.02 mmol) was added. The reaction was stirred at RT for 12h under a nitrogen gas atmosphere. The reaction was quenched with several drops of saturated $NaHCO_3$ aqueous solution and concentrated under vacuum. The residue was dissolved in ethyl acetate and washed with brine. The organic layer was collected and concentrated. Purification through silica gel column chromatography afforded **20** as a white solid (63 mg, 41%).

$^1$H NMR (600 MHz, Chloroform-*d*) δ 7.77 – 7.74 (m, 2H, Ar), 7.72 – 7.68 (m, 2H, Ar), 7.46 – 7.42 (m, 2H, Ar), 7.41 – 7.36 (m, 4H, Ar), 4.59 (dd, *J* = 7.8, 2.2 Hz, 1H, H-1), 4.43 (dd, *J* = 7.8, 6.7 Hz, 0.5H, H-2), 4.35 (dd, *J* = 7.8, 6.6 Hz, 0.5H, H-3), 4.17 (m, 1H, H-3), 4.06 (dd, *J* = 5.8, 1.3 Hz, 1H, H-4), 3.77 (m, 1H, H-6a), 3.59 (ddd, *J* = 12.0, 10.4, 3.8 Hz, 1H, H-6b), 3.47 (ddd, *J* = 8.0, 3.8, 2.3 Hz, 1H, H-5), 1.55 (s, 3H, $CH_3$), 1.32 (s, 3H, $CH_3$), 1.10 (s, 9H, $CH_3$-POM).

$^{13}$C NMR (151 MHz, Chloroform-*d*) δ 135.95, 133.43, 132.55, 130.19, 127.92, 127.77, 110.98, 95.27 (d, *J* = 19.0 Hz, C-1), 94.58 (d, *J* = 142.8 Hz, C-2), 77.82 (d, *J* = 23.2 Hz, C-3), 74.55 (d, *J* = 8.2 Hz, C-4), 73.89, 62.22, 28.07, 26.82, 26.29, 19.26.

ESI MS(m/z): $[M + Na]^+$ calcd for $C_{26}H_{33}FNaO_5Si$, 483.1979; found 483.2039

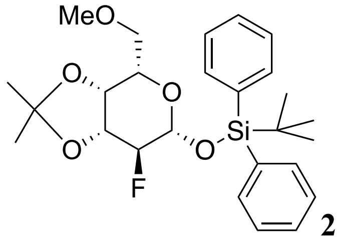


**21**

**2-Deoxy-2-fluoro-3,4-*O*-isopropylidene-6-*O*-methyl-1-*O*-(tert-butyldiphenylsilyl)-L-galactopyranose**

Compound **20** (30 mg, 0.06 mmol) was dissolved in DMF under a nitrogen atmosphere while being placed in an ice bath. Sodium hydride (60% dispersion in mineral oil, 13 mg, 0.3 mmol) was added portion-wise to this solution. After stirring for 5 min, iodomethane (40 μL, 0.6 mmol) was added. The reaction mixture was allowed to warm to RT and was stirred for 12 h. Afterwards, the reaction was cooled to 0 °C and quenched with water. Then the solution was diluted with ethyl acetate and washed with brine. The organic layer was dried over $Na_2SO_4$ and concentrated under vacuum. Purification through silica gel column chromatography afforded **21** as a white solid (25 mg, 83%).

$^1H$ NMR (400 MHz, Chloroform-*d*) δ 7.82 – 7.74 (m, 2H, Ar), 7.75 – 7.69 (m, 2H, Ar), 7.48 – 7.31 (m, 6H, Ar), 4.51 (dd, *J* = 7.8, 2.5 Hz, 1H, H-1), 4.46 (dd, *J* = 7.8, 6.3 Hz, 0.5H, H-2), 4.33 (m, 0.5H, H-2), 4.15 (t, *J* = 6.0 Hz, 0.5H, H-3), 4.10 (m, 1.5H, H-3, H-4), 3.65 (dd, *J* = 9.5, 5.4 Hz, 1H, H-6a), 3.58 (m, 1H, H-5), 3.50 (ddd, *J* = 9.5, 6.3, 0.9 Hz, 1H, H-6b), 3.33 (s, 3H, OMe), 1.56 (s, 3H, $CH_3$), 1.34 (s, 3H, $CH_3$), 1.10 (s, 9H, $CH_3$-POM).

$^{13}C$ NMR (101 MHz, Chloroform-*d*) δ 136.11, 136.01, 132.98, 132.77, 130.01, 129.94, 127.77, 127.64, 110.64, 95.10 (d, *J* = 23.7 Hz, C-1), 94.82 (d, *J* = 185.0 Hz, C-2), 77.73, 74.57 (d, *J* = 8.3 Hz, C-4), 72.22, 71.32, 59.48, 28.13, 26.83, 26.32, 19.38.

ESI MS(m/z): $[M + Na]^+$ calcd for $C_{26}H_{35}FNO_5Si$, 497.2135; found 497.1650

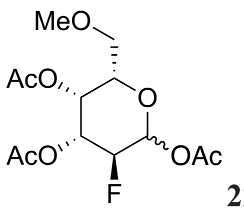


**22**

**2-Deoxy-2-fluoro-per-acetyl-6-*O*-methyl-L-galactopyranose**

Compound **21** (20 mg, 0.04 mmol) was dissolved in a mixture of TFA and water (9:1) and stirred at RT for 1 h. Then, the solution was co-evaporated with toluene to remove the acid and dried under vacuum. The resulting residue was dissolved in THF (2 mL), and a solution of Tetrabutylammonium fluoride (1.0 M in THF, 0.2 mL) and acetate acid (24 μL, 0.4 mmol) was added. The reaction mixture was stirred at RT under a nitrogen atmosphere for 6 h, and then the solution was concentrated under vacuum. The residue was dissolved in a mixture of pyridine and acetic anhydride and stirred for 4 h at RT. Afterward, the reaction mixture was diluted with ethyl acetate and washed with water. The organic layer was dried over $Na_2SO_4$ and concentrated under vacuum. Purification through silica gel column chromatography afforded **22** as a white solid (13 mg, 95%).

$^1H$ NMR (600 MHz, Chloroform-*d*) δ 6.48 (d, *J* = 3.9 Hz, 3.3H, αH-1), 5.79 (dd, *J* = 8.0, 4.1 Hz, 1H, βH-1), 5.55 (td, *J* = 3.5, 1.3 Hz, 3.3H, αH-4), 5.49 (m, 1H, βH-4), 5.41 (m, 3.3H, αH-3), 5.17 (ddd, *J* = 13.2, 9.9, 3.6 Hz, 1H, βH-3), 4.94 (dd, *J* = 10.2, 4.0 Hz, 1.7H, αH-2), 4.85 (dd, *J* = 10.2, 4.0 Hz, 1.7H, αH-2), 4.69 (dd, *J* = 9.9, 8.0 Hz, 0.5H, βH-2), 4.60 (dd, *J* = 9.9, 8.0 Hz, 0.5H, βH-2), 4.23 (td, *J* = 6.1, 0.8 Hz, 3.3H, αH-5), 3.97 (t, *J* = 6.2 Hz, 1H, βH-5), 3.50 (dd, *J* = 10.1, 5.9 Hz, 1H, βH-6a), 3.44 (dd, *J* = 10.0, 6.2 Hz, 3.3H, αH-6a), 3.41 (dd, *J* = 10.1, 6.2 Hz, 1H, βH-6b), 3.36 (dd, *J* = 10.0, 6.0 Hz, 3.3H, αH-6b), 3.31 (s, 13H, $CH_3$-OMe), 2.17 (s, 13H, $CH_3$-Ac), 2.15 (s, 13H, $CH_3$-Ac), 2.06 (s, 13H, $CH_3$-Ac).

$^{13}$C NMR (151 MHz, Chloroform-*d*) δ 170.16, 170.08, 169.06, 89.26 (d, *J* = 22.6 Hz, αC-1), 84.52 (d, *J* = 190.9 Hz, αC-2), 70.42, 70.04, 68.84 (d, *J* = 7.7 Hz, αC-4), 68.50 (d, *J* = 18.7 Hz, αC-3), 59.55, 21.07, 20.82, 20.70.

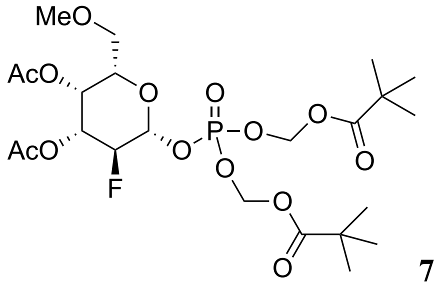

**7**

**2-Deoxy-2-fluoro-3,4-di-*O*-acetyl-6-*O*-methyl-β-1 (dipivaloyloxymethylphosphoryl)-L-galactopyranose**

Compound **22** (13 mg, 0.04 mmol) was dissolved in HBr (33% in AcOH, 1 mL) and stirred for 1 h at RT. Afterward, the mixture was concentrated *in vacuo*. The resulting residue was dissolved in 2 mL dry $CH_3CN$, and 200 mg of 3 Å molecular sieves were added. The mixture was stirred at RT for 10 min. Then, in one portion, silver carbonate (55 mg, 0.2 mmol) and diPOM-phosphate, compound **28** (55 mg, 0.2 mmol) were added. The reaction mixture was stirred at RT, under nitrogen gas protection, and in the dark. After 12 h, the mixture was diluted with ethyl acetate and then filtered through Celite. The filtrate was concentrated *in vacuo*. The residue was purified by silica gel chromatography, yielding **7** (15 mg, 65%) as a white solid.

$^{1}$H NMR (600 MHz, Chloroform-*d*) δ 5.75 – 5.61 (m, 4H, $CH_2$-POM), 5.47 (m, 1H, H-4), 5.38 (td, *J* = 7.3, 3.9 Hz, 1H, H-1), 5.13 (ddd, *J* = 13.1, 9.9, 3.5 Hz, 1H, H-3), 4.63 (dd, *J* = 9.9, 7.6 Hz, 0.5H, H-2), 4.55 (dd, *J* = 9.9, 7.6 Hz, 0.5H, H-2), 3.96 (t, *J* = 6.3 Hz, 1H, H-5), 3.51 (dd, *J* = 10.0, 6.1 Hz, 1H, H-6a), 3.42 (dd, *J* = 10.0, 6.4 Hz, 1H, H-6b), 3.31 (s, 3H, OMe), 2.14 (s, 3H, $CH_3$-Ac), 2.06 (s, 3H, $CH_3$-Ac), 1.24 (d, *J* = 4.6 Hz, 18H, $CH_3$-POM).

$^{13}$C NMR (151 MHz, Chloroform-*d*) δ 176.73, 169.95, 169.84, 96.83 (d, *J* = 24.5 Hz, C-1), 87.88 (d, *J* = 189.8 Hz, C-2), 83.03, 83.00, 73.06, 70.84 (d, *J* = 17.8 Hz, C-3), 69.80, 67.89 (d, *J* = 7.9 Hz, C-4), 59.47, 38.89, 26.96, 26.93, 20.70, 20.67.

ESI MS(m/z): $[M + Na]^+$ calcd for $C_{23}H_{38}FNaO_{14}P$, 611.1881; found 611.0770

**Preparation of substrate 8**

SGP (50 mg, 0.02 mmol) was dissolved in a sodium acetate buffer (PH 5.5, 50 mM), containing calcium chloride (10 mM). Neuraminidase was added to the mixture, and the reaction was incubated at 37 °C with shaking until ESI indicated complete removal of sialic acid. The pH of the reaction mixture was then adjusted to 4.5 using acetic acid. Subsequently, BSA and β-galactosidase were introduced into the mixture, and the reaction was incubated at 37 °C with shaking until ESI indicated that all galactose residues had been removed. The protein in the reaction mixture was separated by centrifugation using a 10K MWCO filter. The filtrate was lyophilized and subjected to multiple rounds of purification via size exclusion chromatography (Bio-Rad P2 gel and P4 gel), eluting with a 0.1 M ammonium bicarbonate solution. Fractions containing the pure products were collected and lyophilized, resulting in the isolation of compound **8** (5.3 mg, 15.5%) as a white powder.

ESI MS(m/z): $[M - 2H]^{2-}$ calcd for $C_{78}H_{133}N13O_{44}^{2-}$, 977.9290; found 977.9101

**Kinetic Studies**

The recombinant human fucosyltransferases were expressed following the reported procedures.[1] All Ki values determinations were performed with the GDP-Glo™ Glycosyltransferase Assay Kit from Promega. Kinetic parameters were determined by maintaining a saturating concentration of acceptors while varying concentrations of GDP-fucose (3-100 μM) and fluorinated GDP-fucose (3-300 μM); specific values for each Ki can be found in the graphs corresponding to each inhibitor and FUT (see Section 5) Graphs Depicting the Determination of Ki). The amount of enzyme used is determined based on the test reaction to ensure that the reaction process is within the linear rate range. Transfer reactions involving inhibitors were performed in a 25 μL reaction volume in PBS buffer, containing appropriate amounts of FUT, acceptor, inhibitor, and GDP-fucose. Enzyme assay mixtures were incubated for 1 h at 37 °C, and the reaction was stopped by adding 25 μL of GDP Detection Reagent. The assay plate was shaked for 30 sec using an automated plate shaker and then incubated at RT for 60 min. Luminescence was measured using a plate reader (BMG Labtech, POLARstar Omega) with the gain set at 3600, the lens as emission filter, a measurement interval of 1 or 2 sec, and the top optic. A control group, which included all reaction components except FUT, was used as a reaction blank. All experiments were performed in triplicate.

**Cell Culture**

HL-60 cells (ATCC CCL-240) were cultured in RPMI 1640 medium supplemented with 10% (v/v) fetal bovine serum (FBS) and 1% penicillin-streptomycin at 37 °C and 5% $CO_2$ in a humidified atmosphere.

**Flow Cytometry**

For the cell surface staining experiments, HL-60 cells were seeded into a 96-well plate (10,000 cells/100 μL medium per well). After allowing the cells to settle for several hours, 100 μL of medium containing the inhibitor at the desired concentration was added. An equivalent concentration of DMSO was added to the control group, as all inhibitor stock solutions were prepared with DMSO. Cells were allowed to grow for 72 h. After this, cells were harvested and washed with FACS buffer (BSA/PBS, 1%, w/v). The staining steps were performed on ice for 30 min with biotinylated AAL (1 μg/mL; Vector Labs, B1395) in FACS buffer, followed by streptavidin Alexa Fluor-488 (1 μg/mL; Invitrogen, S11223) in FACS buffer. After staining, cells were washed 3 times with FACS buffer and then resuspended for flow cytometry. Experiments were performed in triplicate and repeated at least 3 times. The data were normalized to control cells treated with DMSO only (100%) and unstained cells (0%).

**Cell Proliferation and Viability**

Cell viability was assessed using trypan blue staining, while cell proliferation was determined with the CellTiter 96 Aqueous One Solution Cell Proliferation Assay kit (MTS assay). Briefly, cells were incubated with desired concentrations of inhibitors for 3 days, after which the assay solutions were added. The assays were performed following the manufacturer's instructions. Data were recorded using a Promega plate reader.

#### 4) NMR Spectra

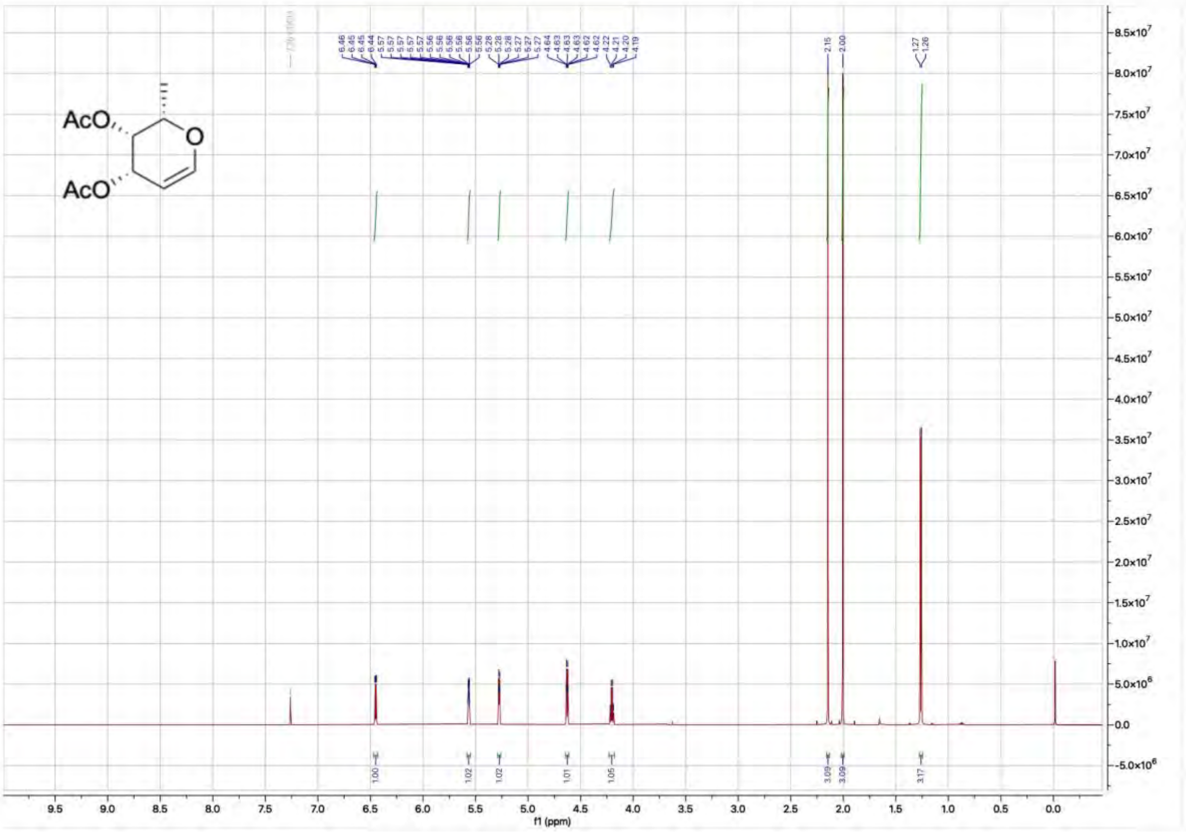

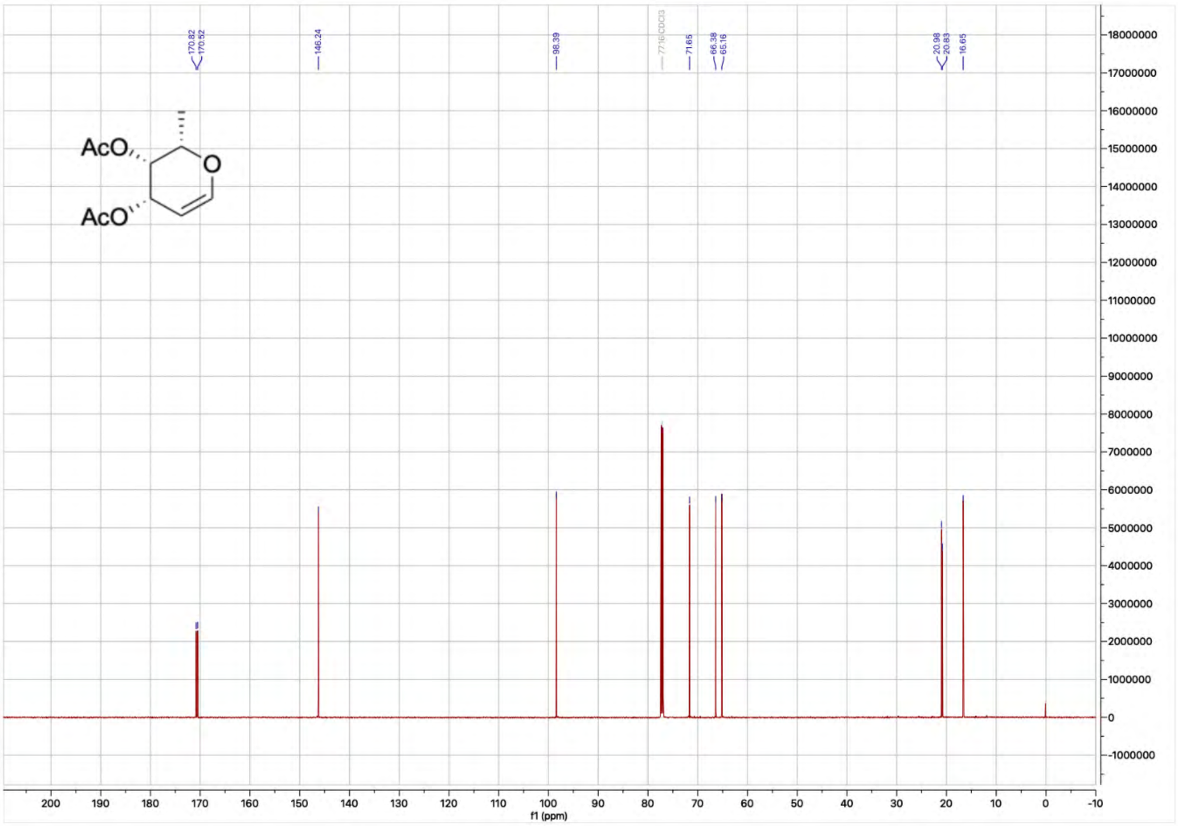

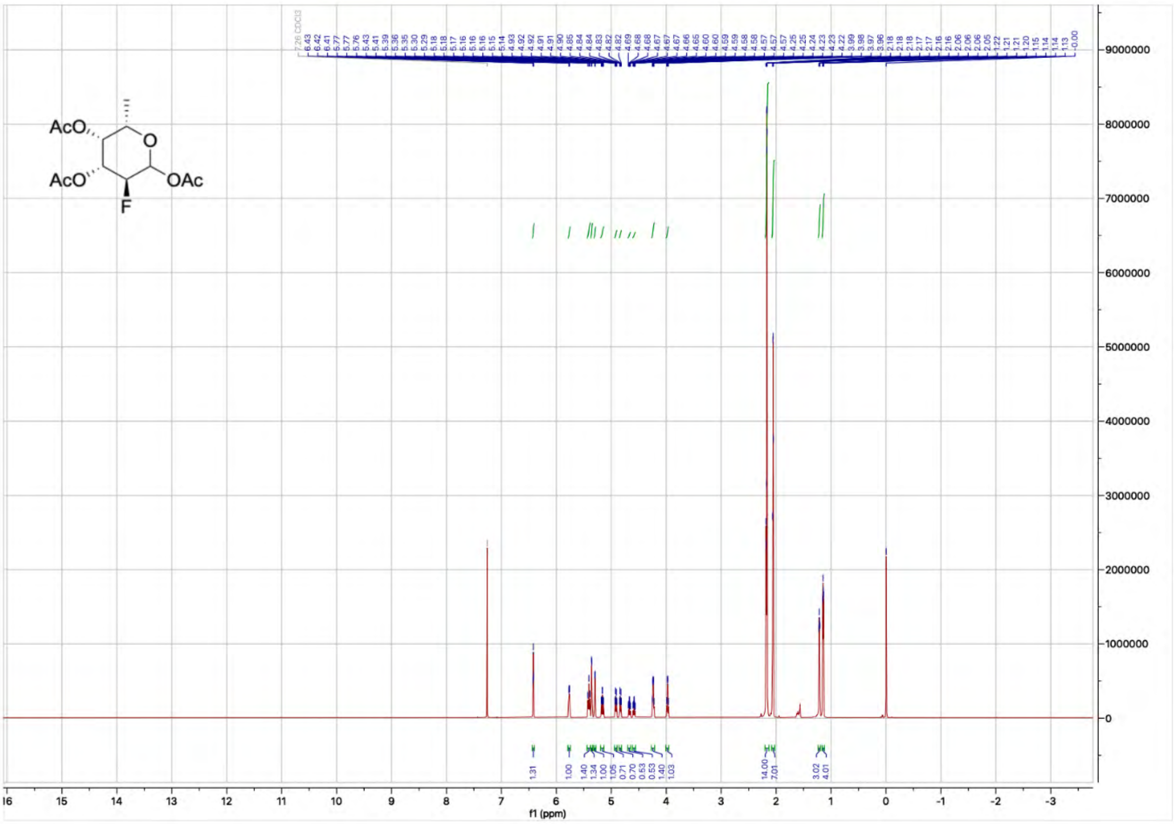

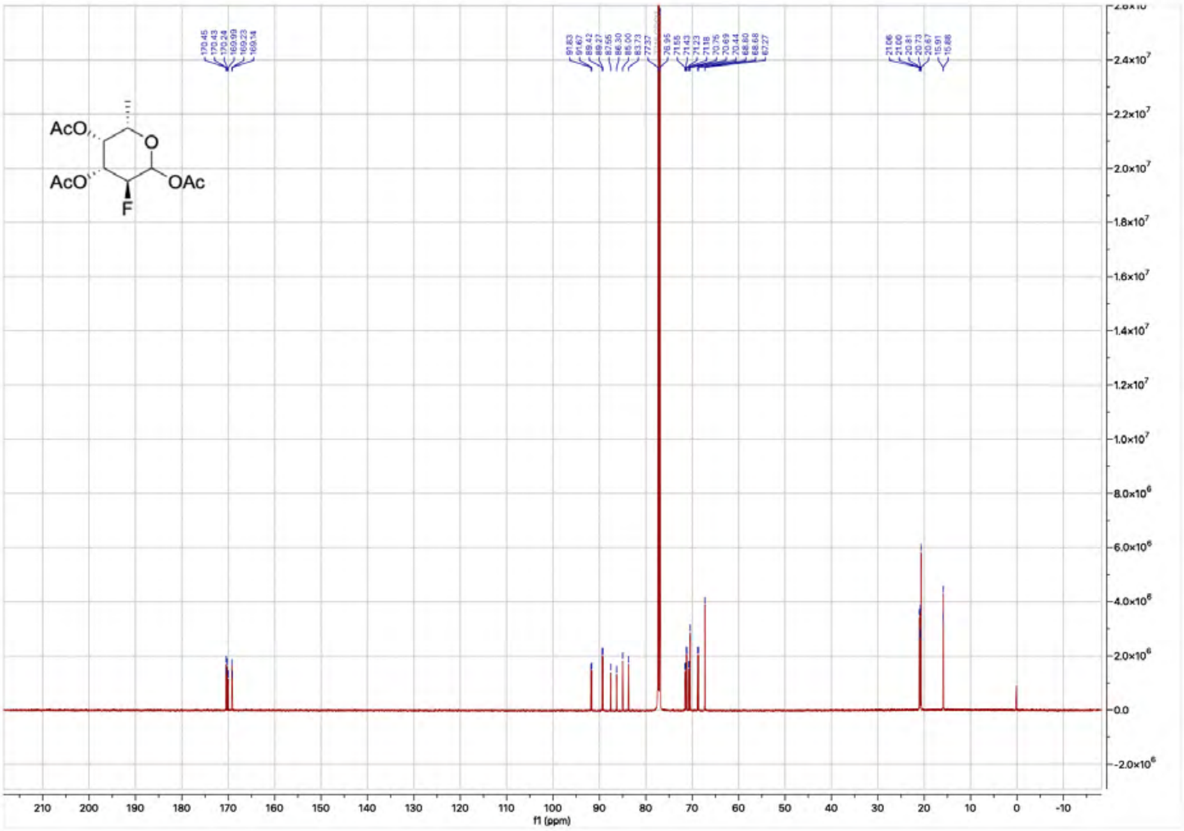

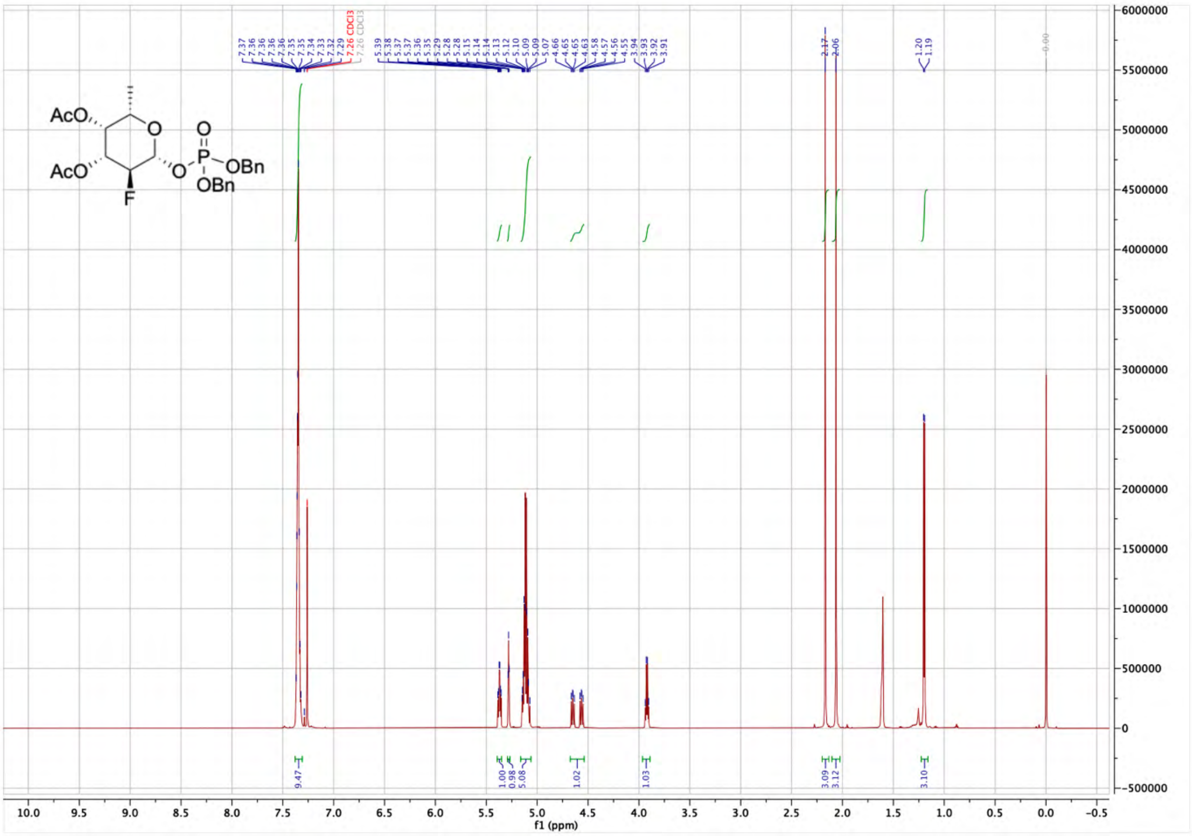

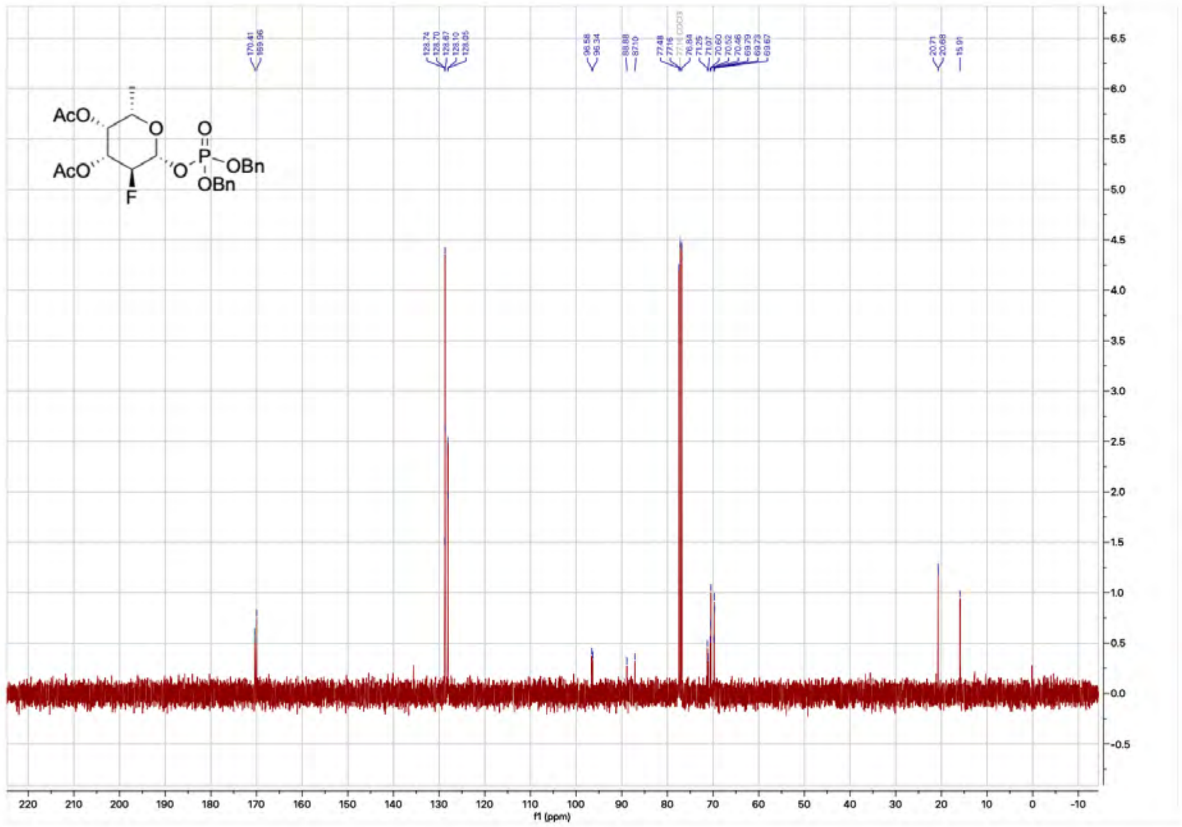

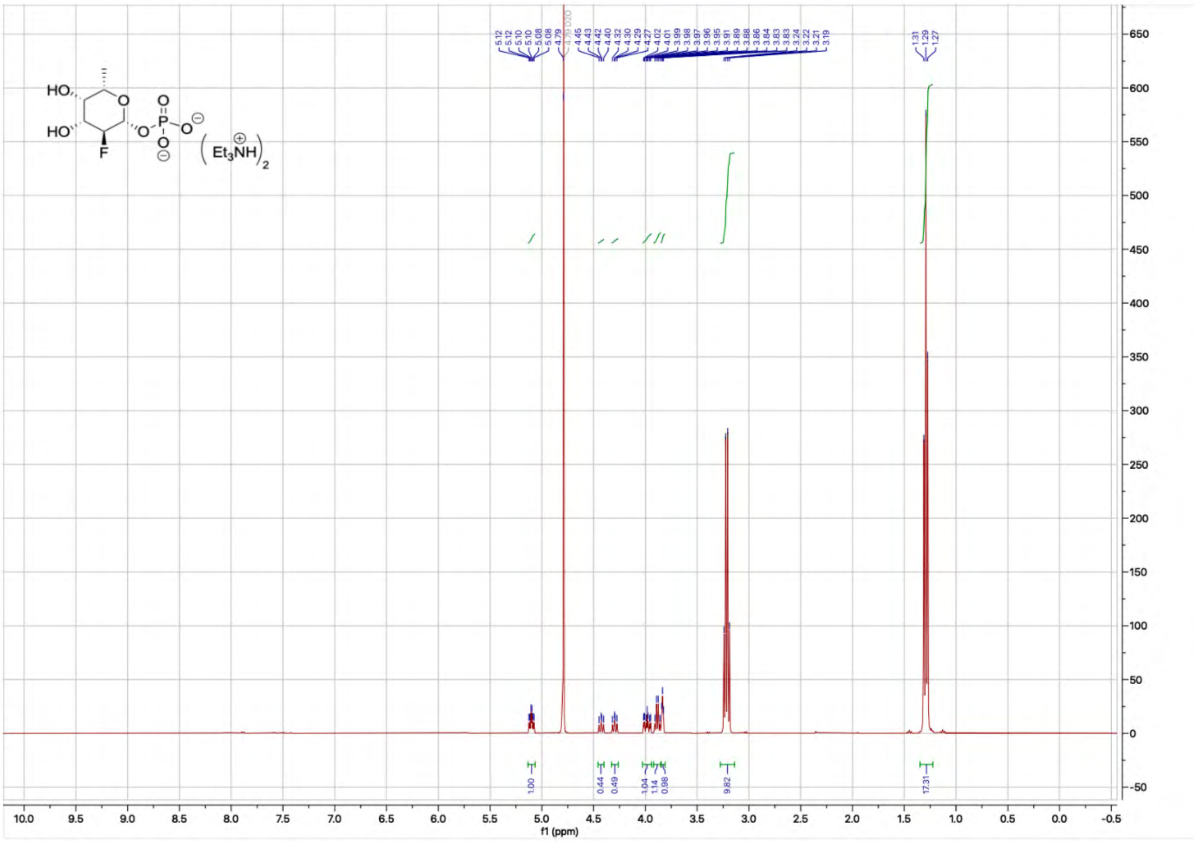

HO
HO
F
O
P
Et3NH
f1 (ppm)


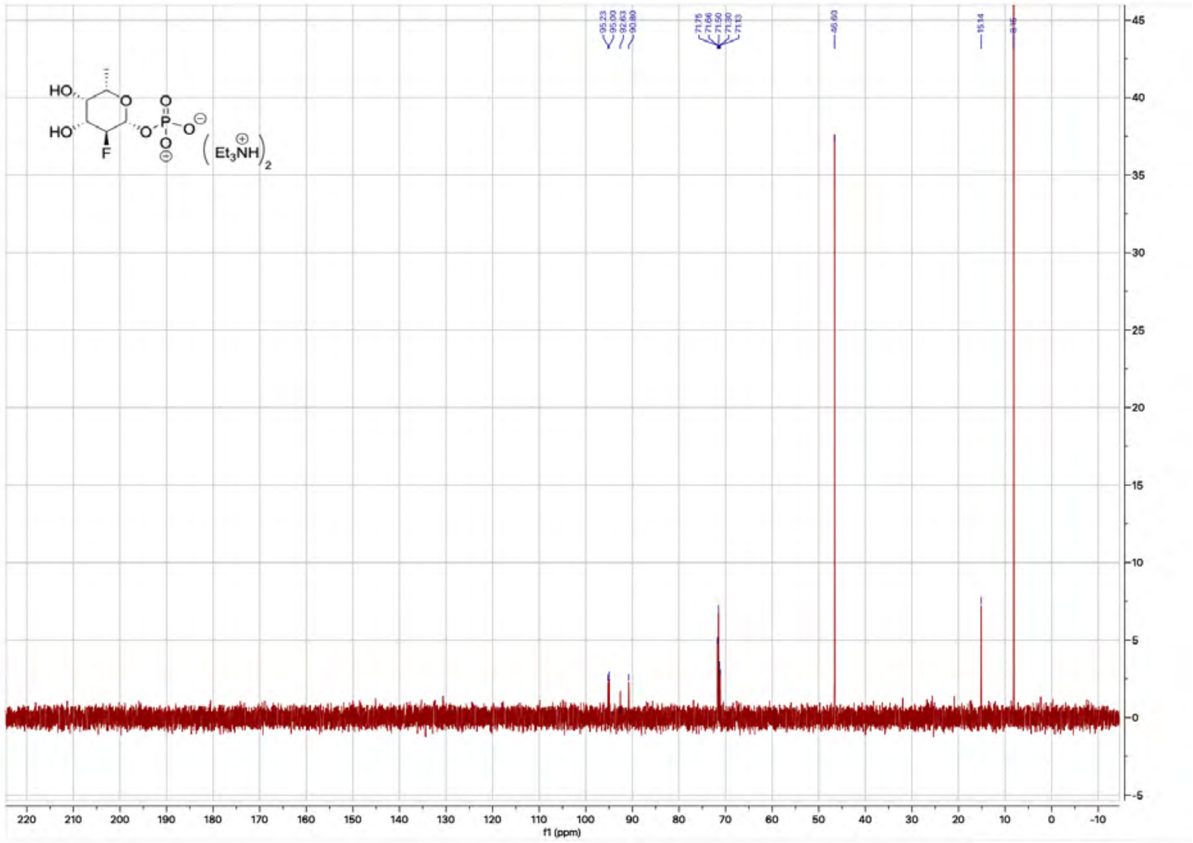

HO
HO
F
O
P
Et3NH
f1 (ppm)

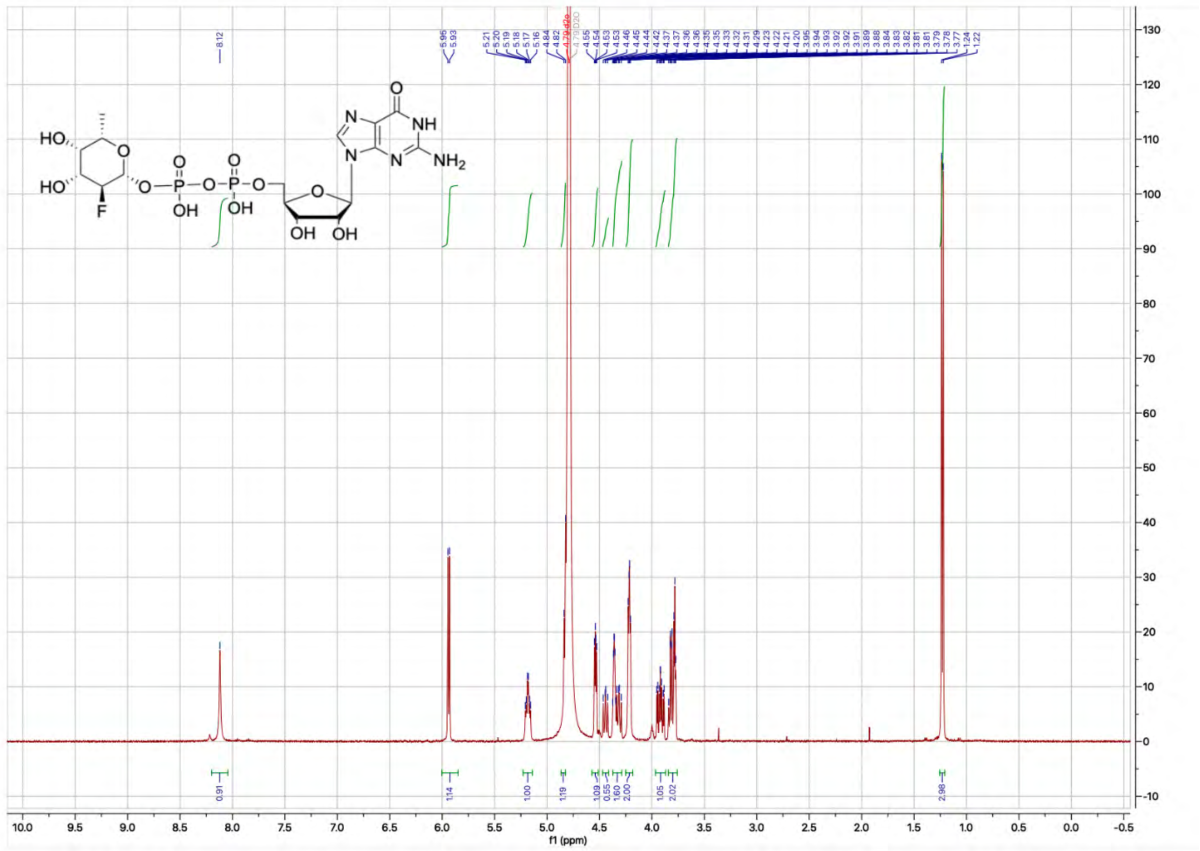
f1 (ppm)

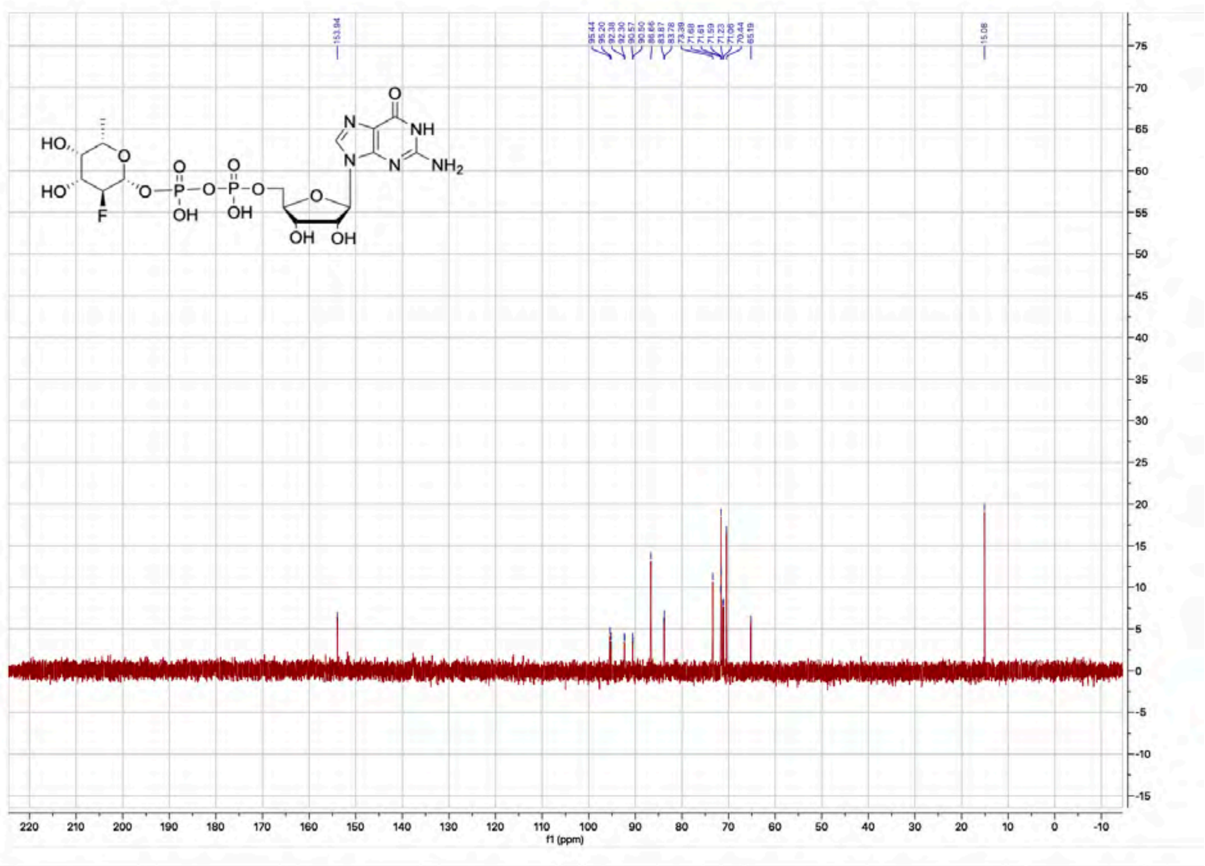
153.94
15.08
f1 (ppm)

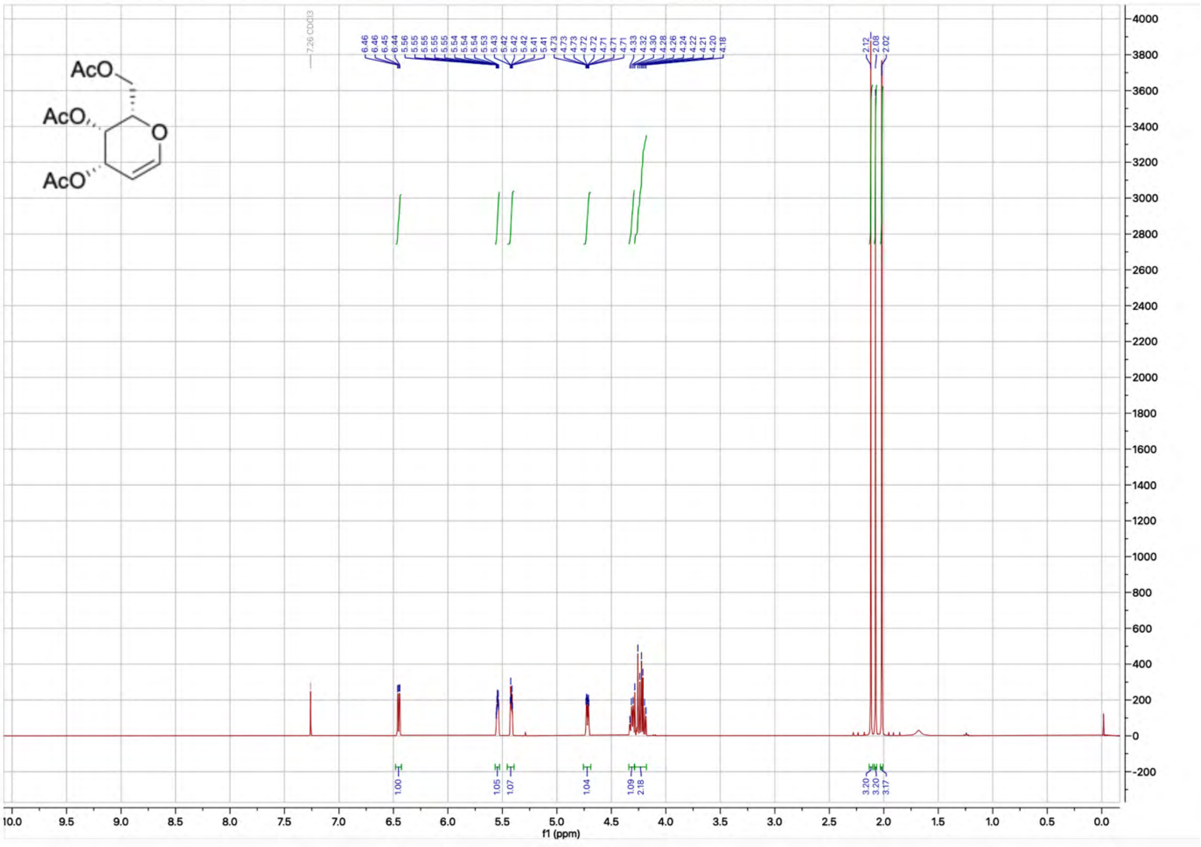

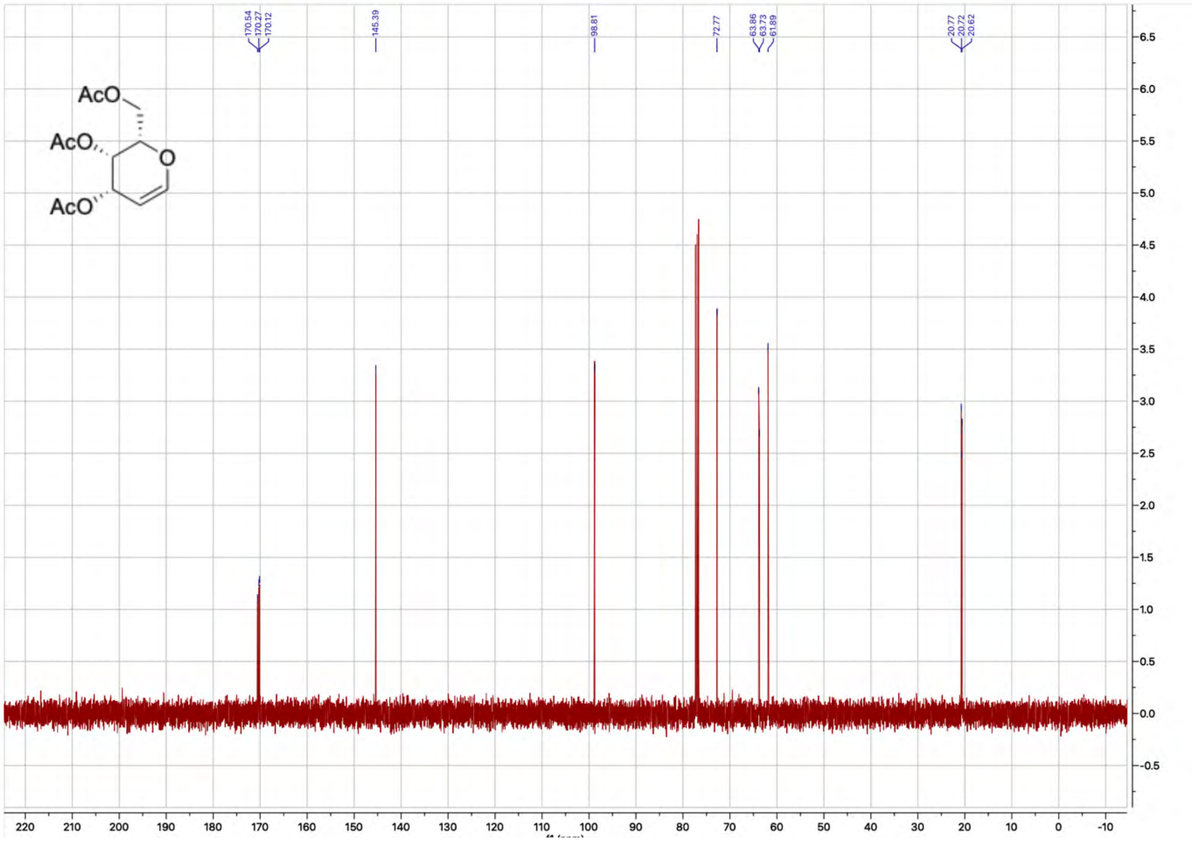

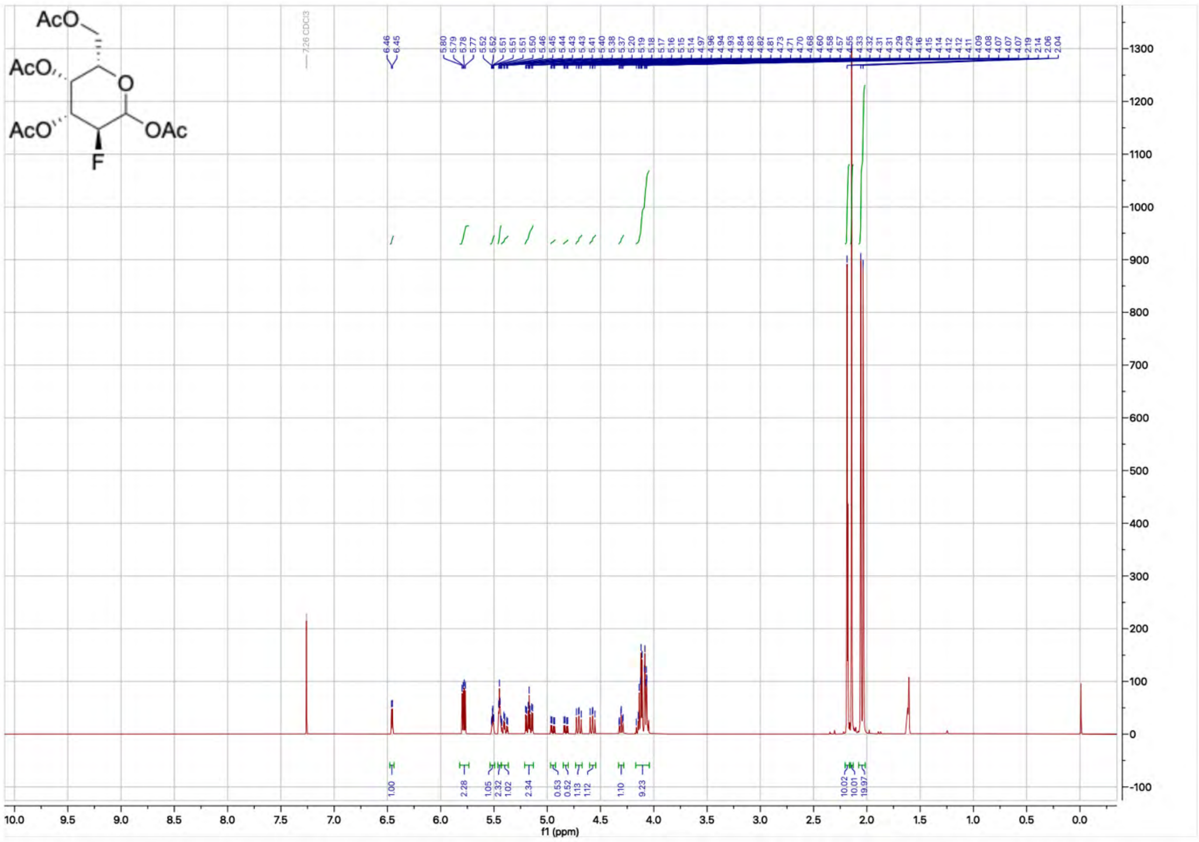

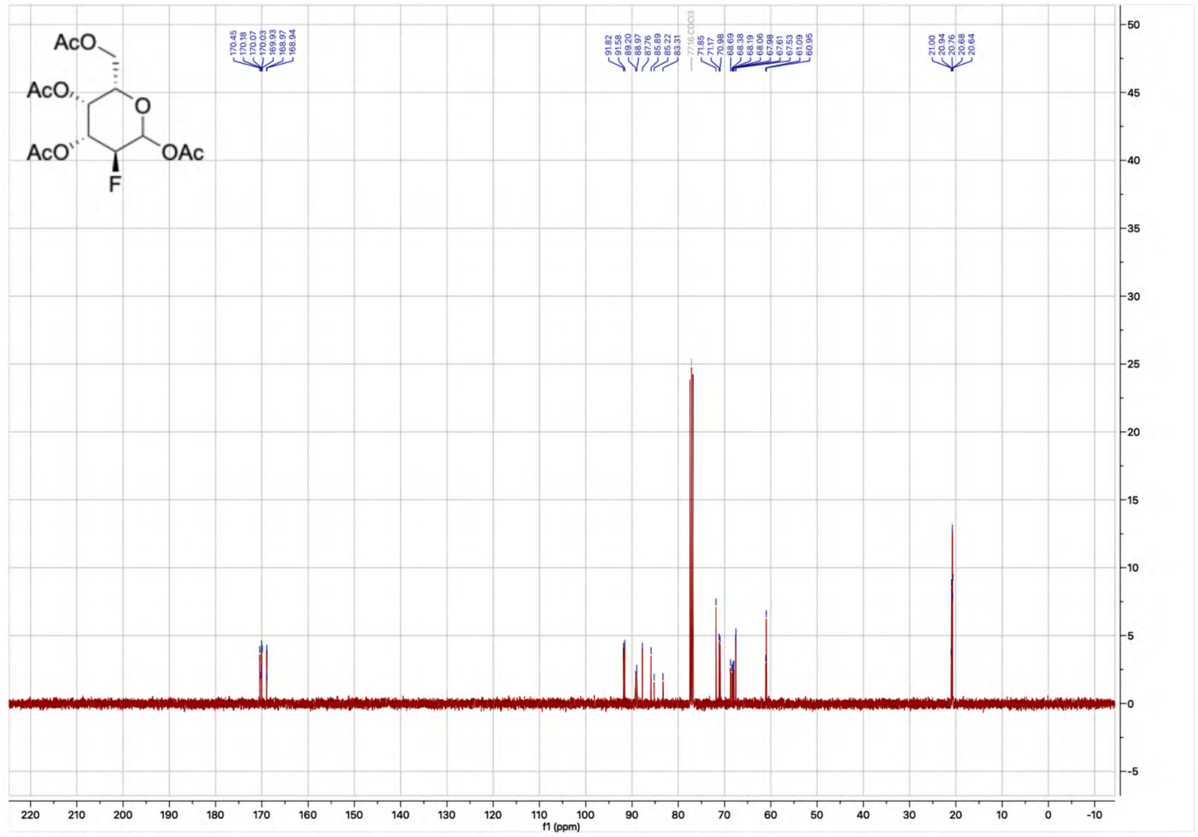

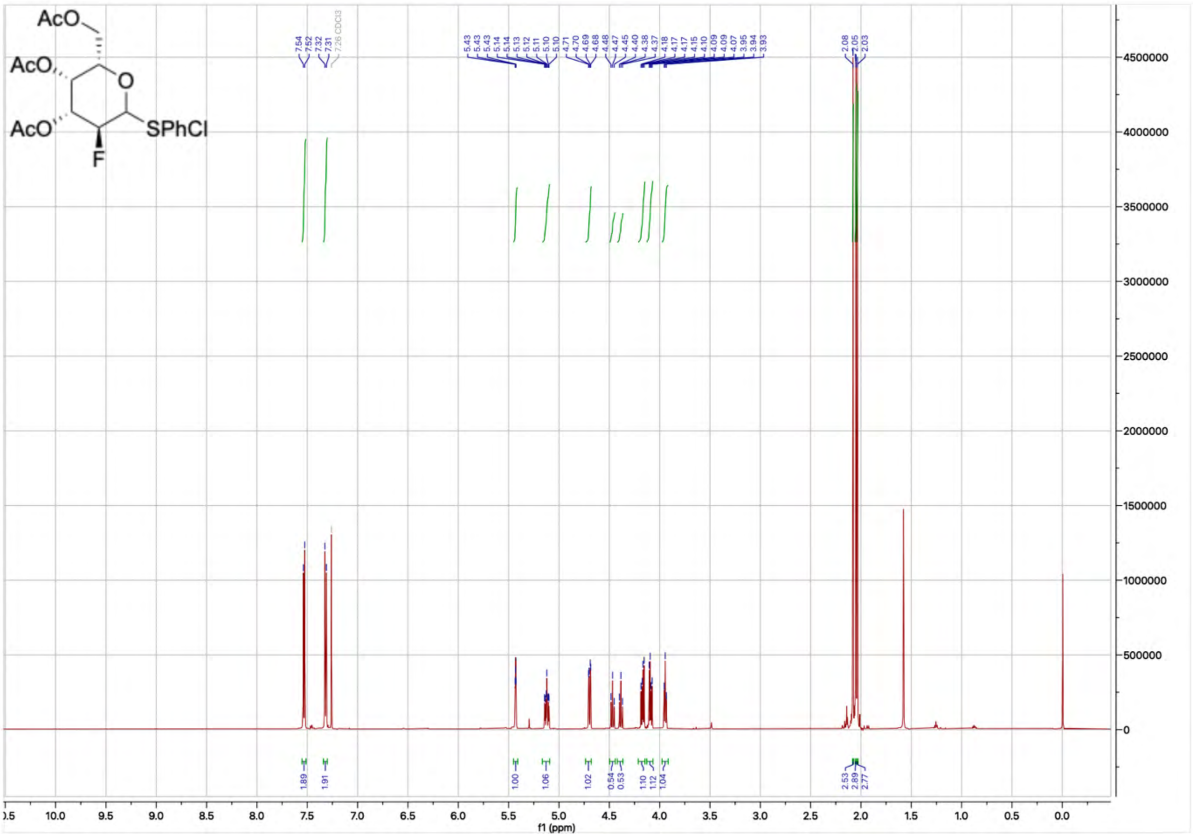

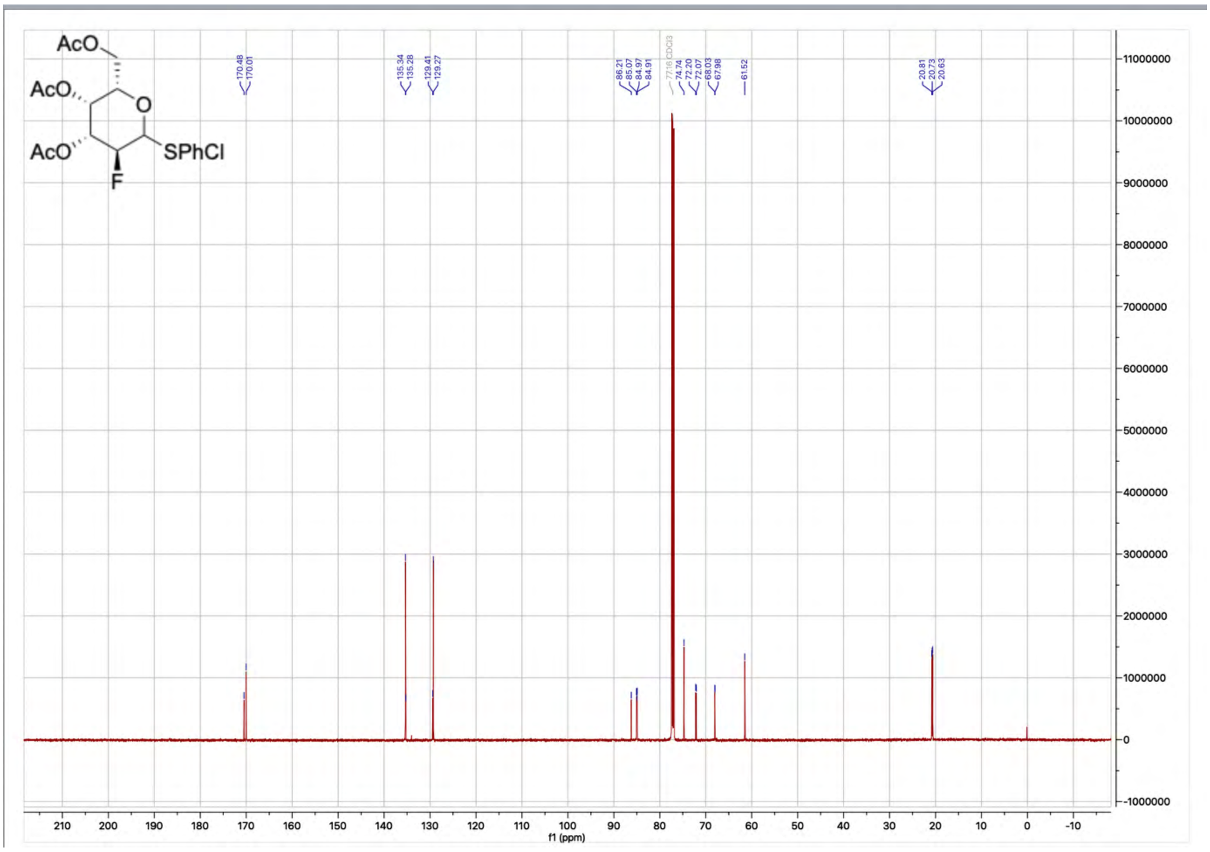

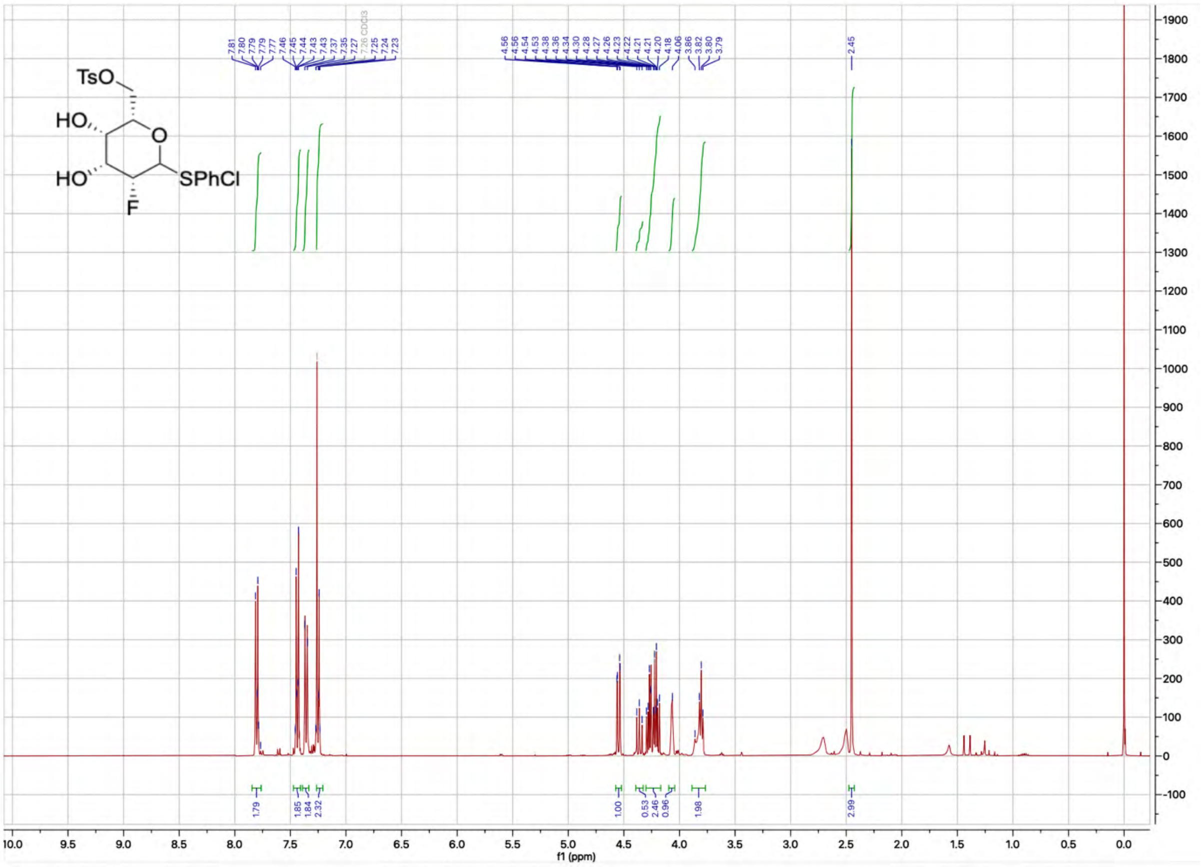
TsO
HO
HO
O
SPhCl
F
7.81
7.80
7.79
7.79
7.77
7.46
7.45
7.44
7.43
7.43
7.37
7.35
7.27
7.26 CDCl3
7.25
7.24
7.23
4.56
4.56
4.54
4.53
4.38
4.36
4.34
4.30
4.28
4.27
4.26
4.23
4.22
4.21
4.21
4.20
4.18
4.06
3.86
3.82
3.80
3.79
2.45
1.79
1.85
1.84
2.32
1.00
0.53
2.46
0.96
1.98
2.99
f1 (ppm)

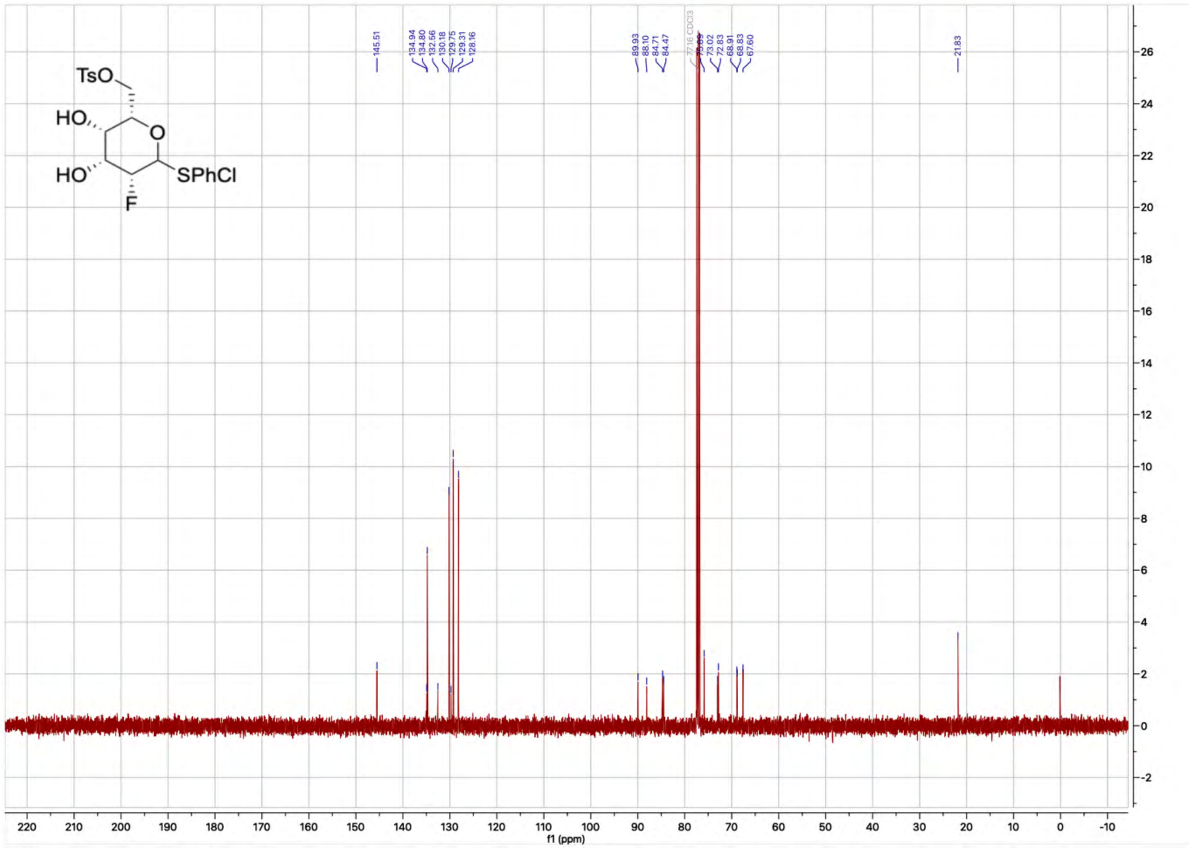
TsO
HO
HO
O
SPhCl
F
145.51
134.94
134.80
132.56
130.18
129.75
129.31
128.16
89.93
88.10
84.71
84.47
77.16 CDCl3
75.69
73.02
72.83
68.91
68.83
67.60
21.83
f1 (ppm)

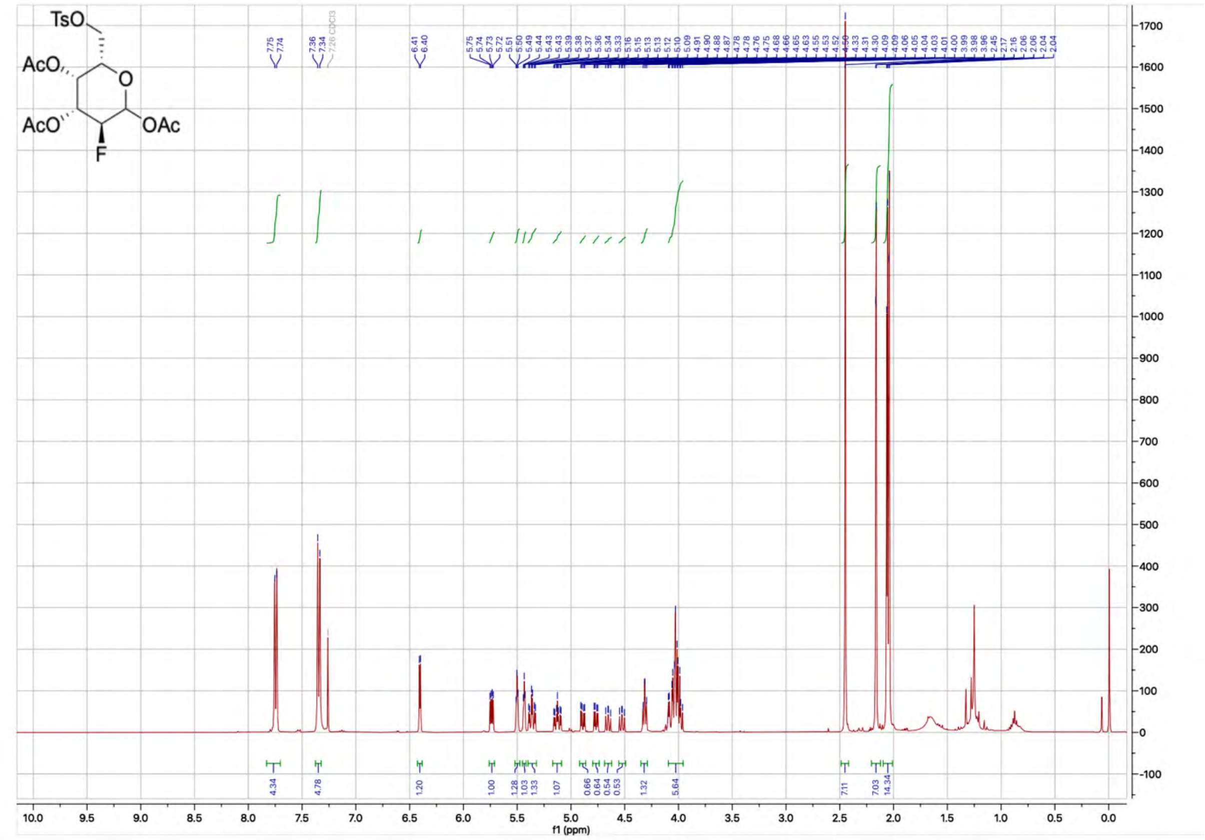
TsO
AcO
AcO
OAc
F
f1 (ppm)

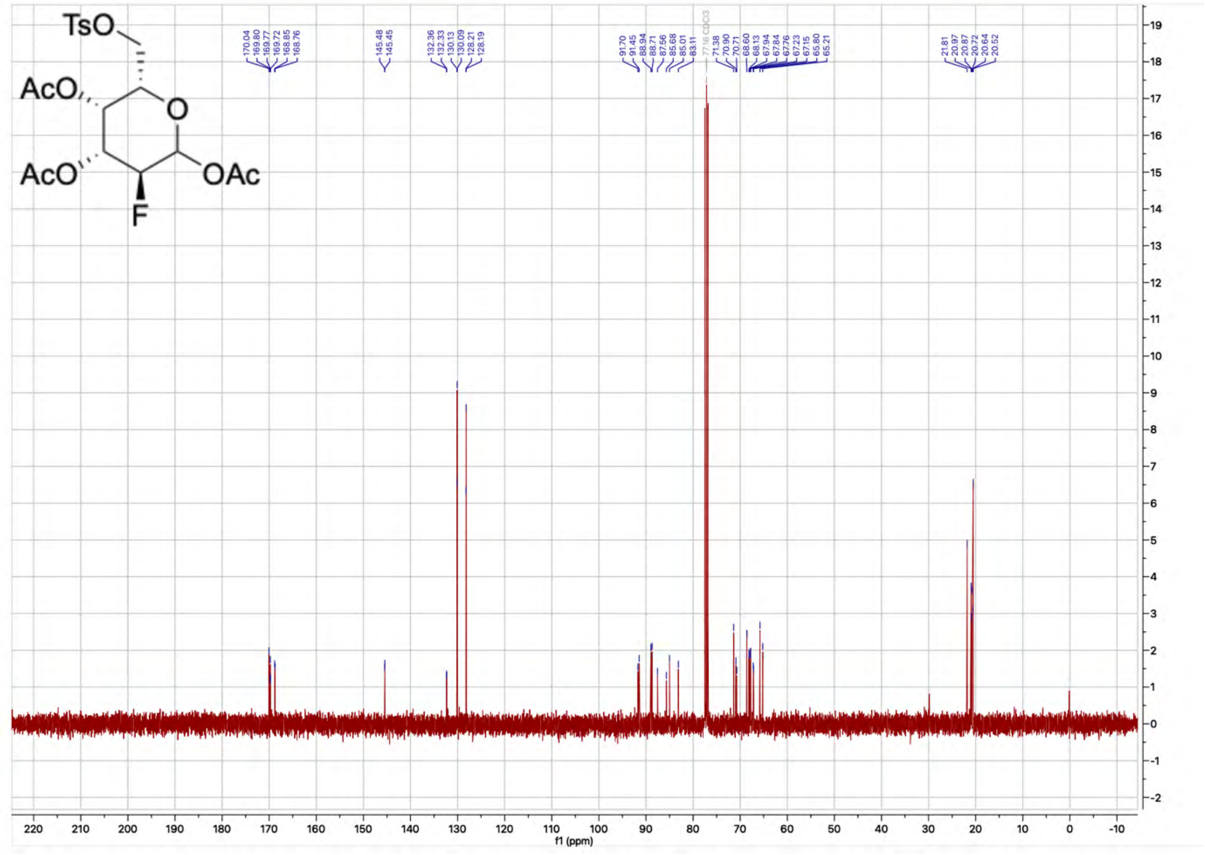
TsO
AcO
AcO
OAc
F
f1 (ppm)

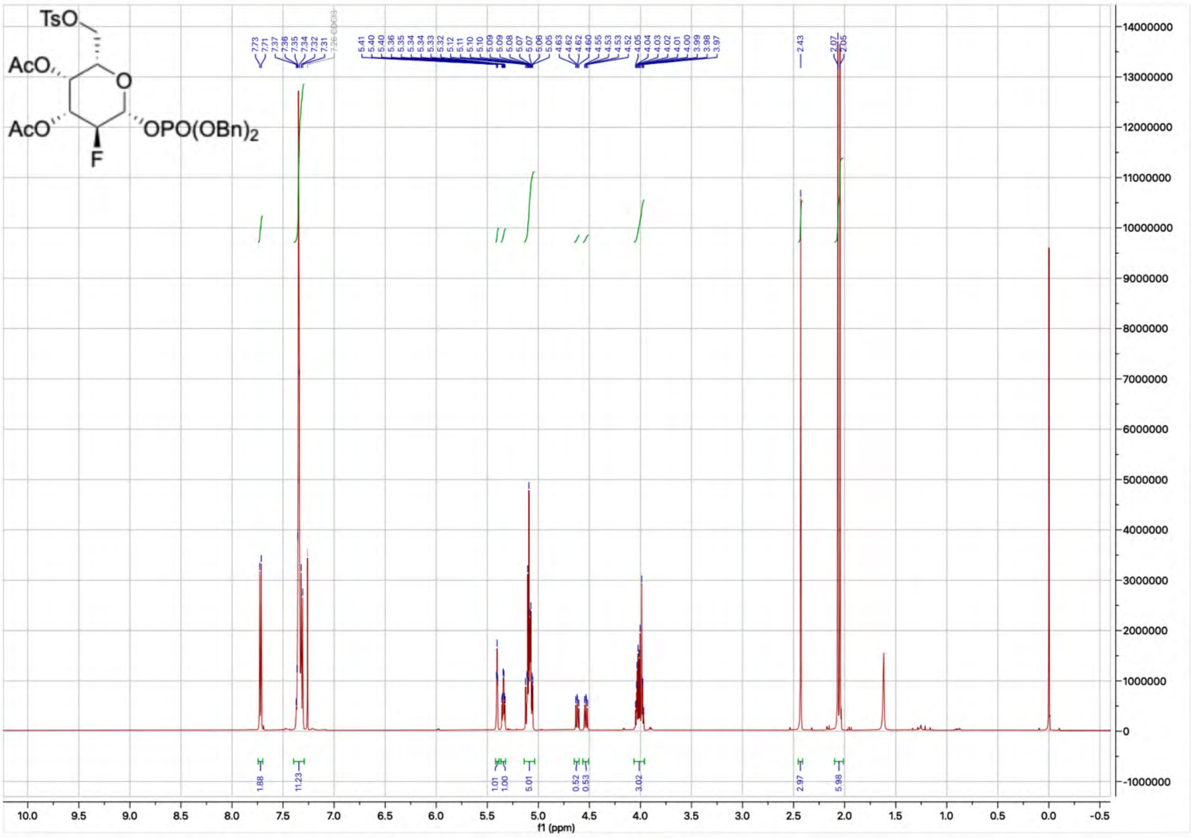
TsO
AcO
AcO
O
OPO(OBn)2
F
f1 (ppm)

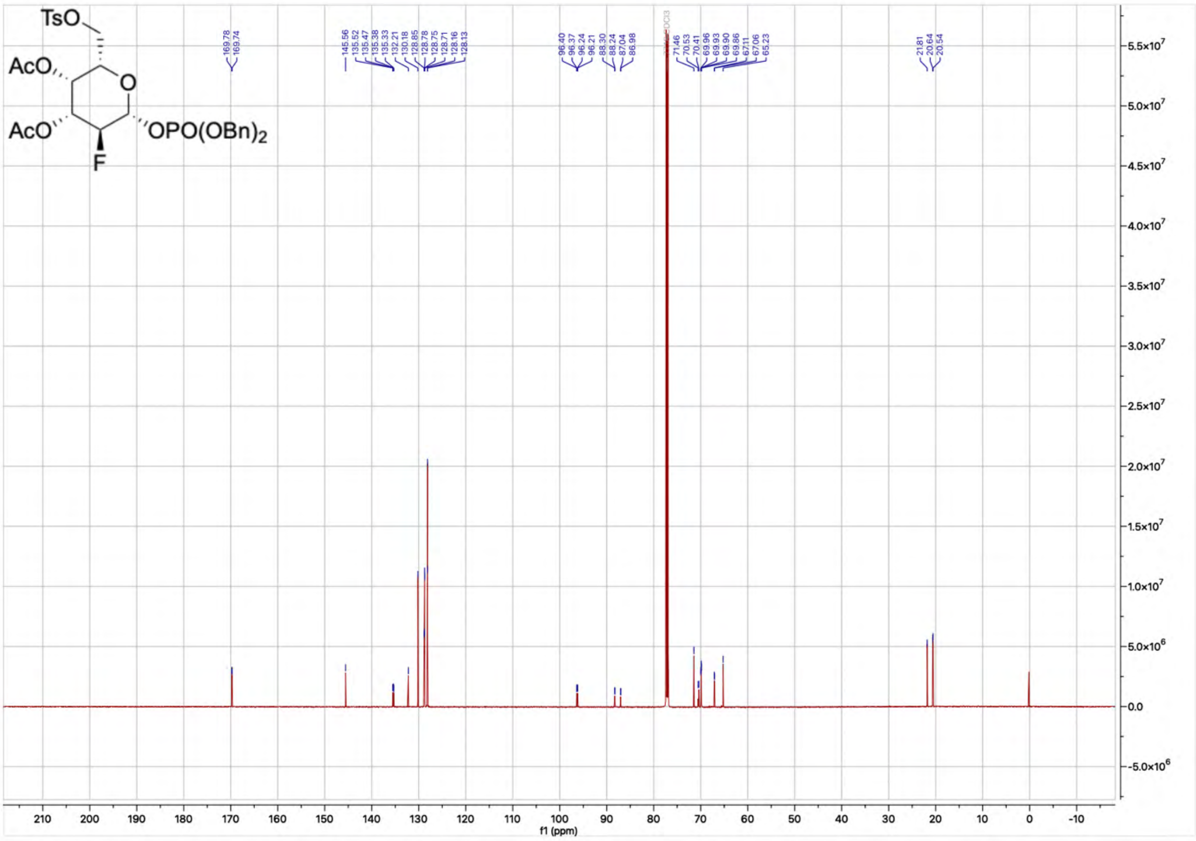
TsO
AcO
AcO
O
OPO(OBn)2
F
f1 (ppm)

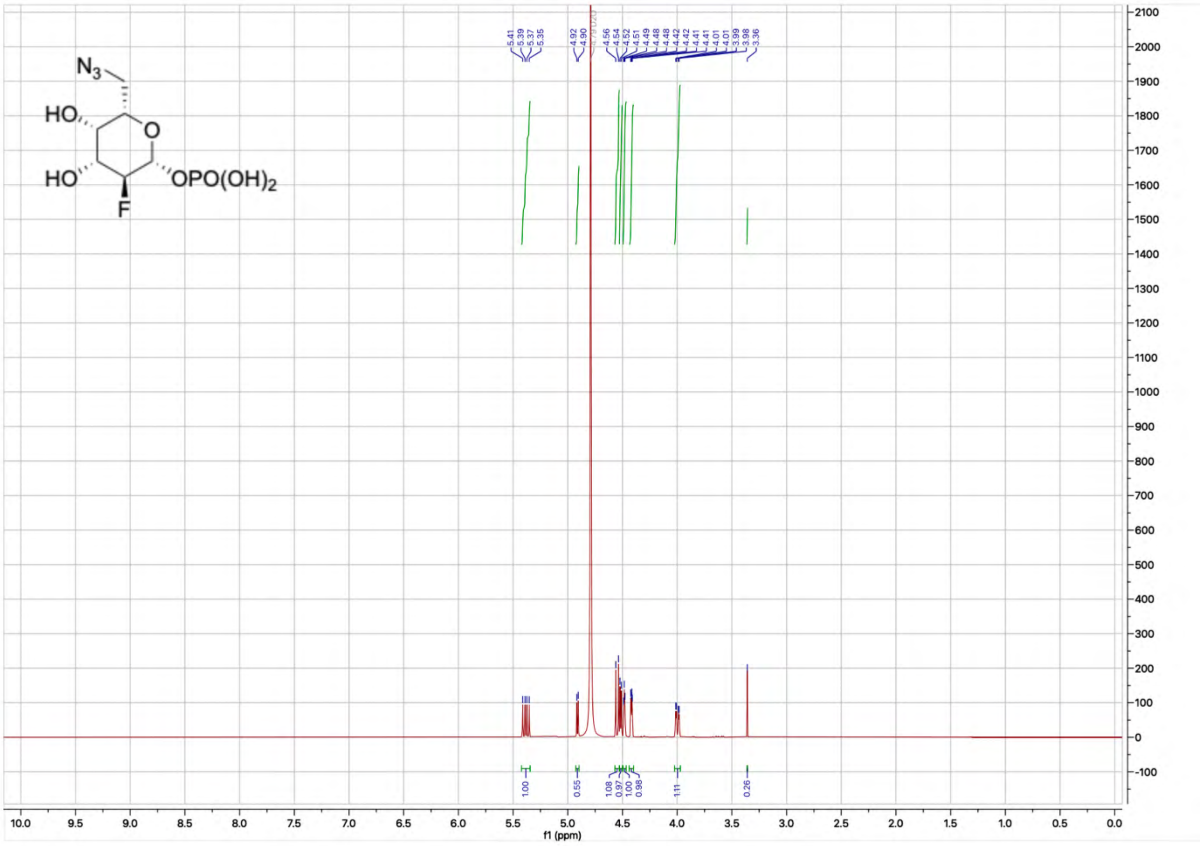

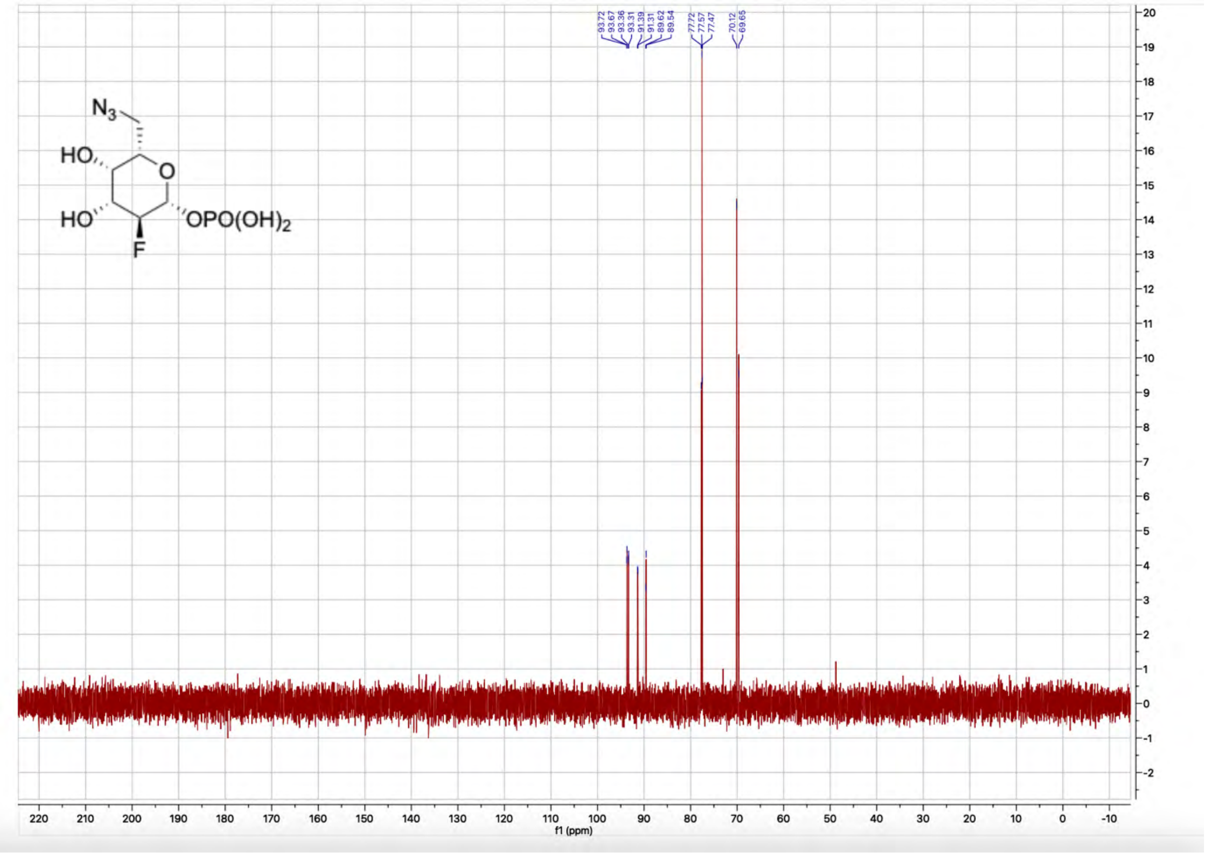

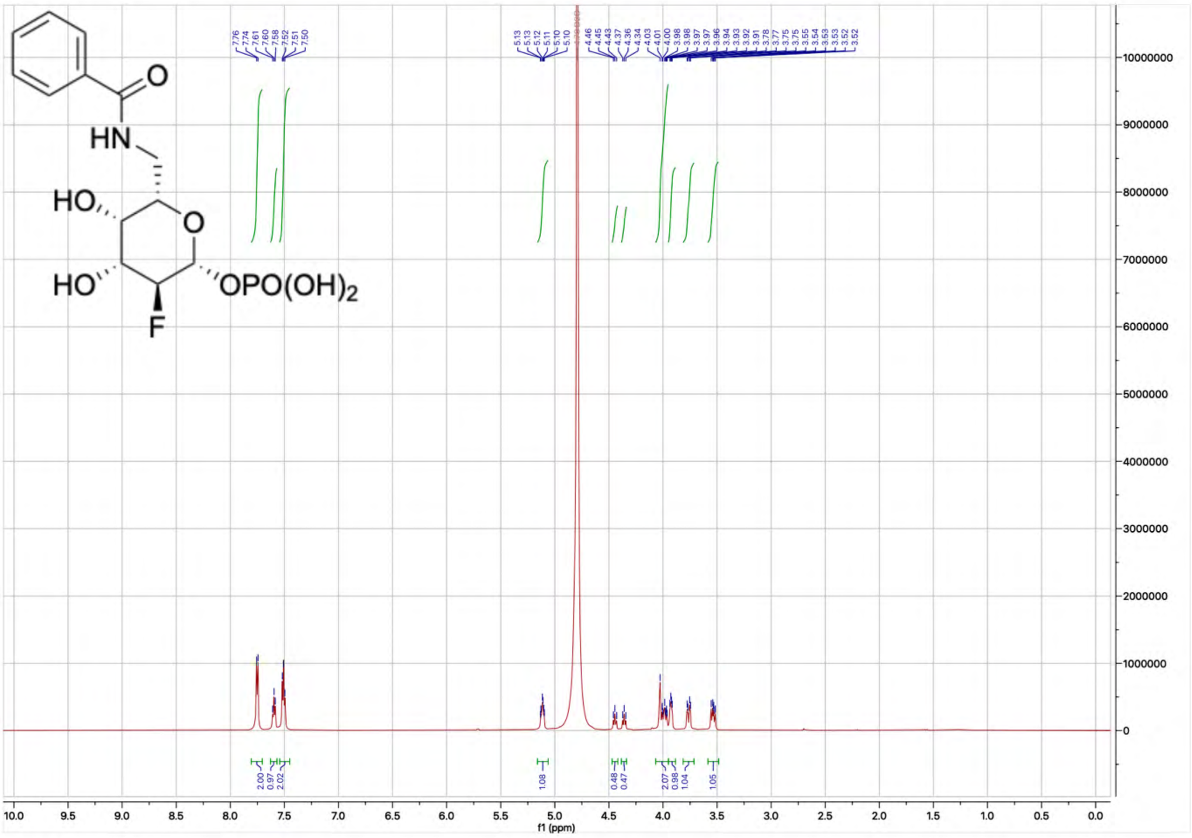

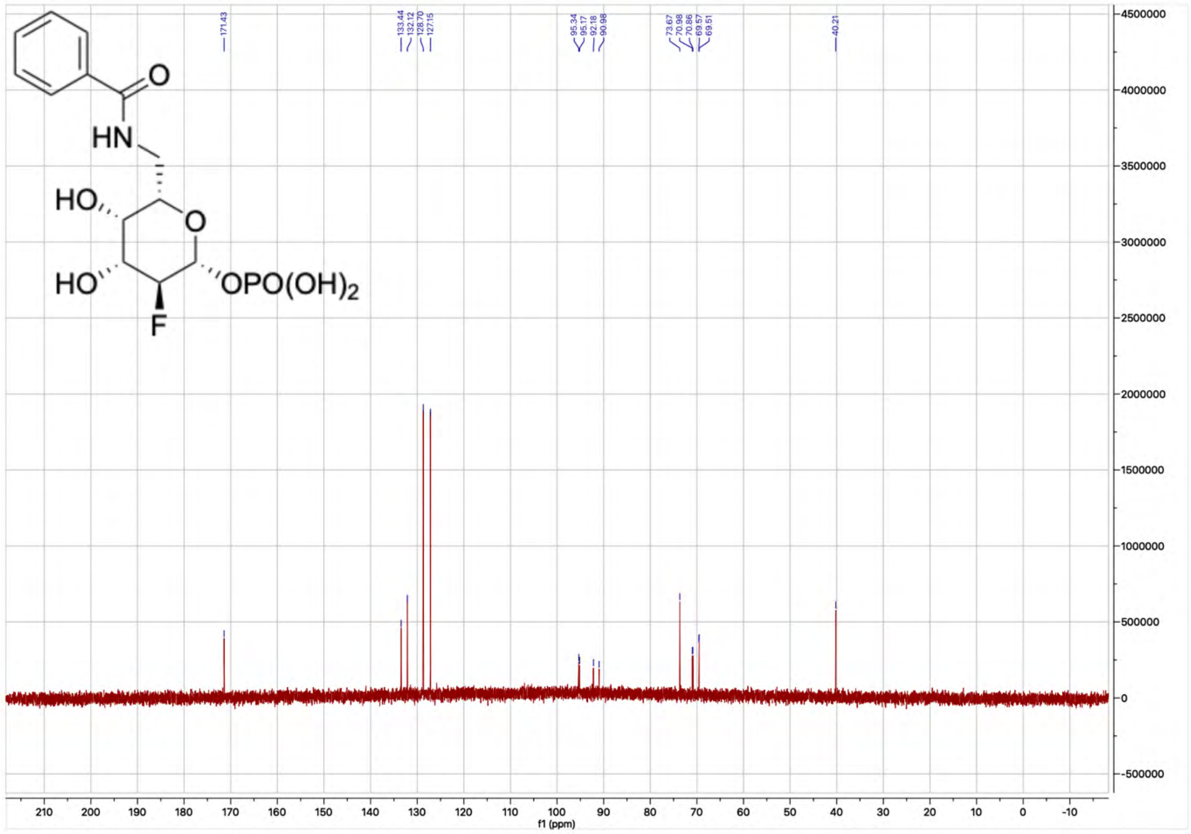

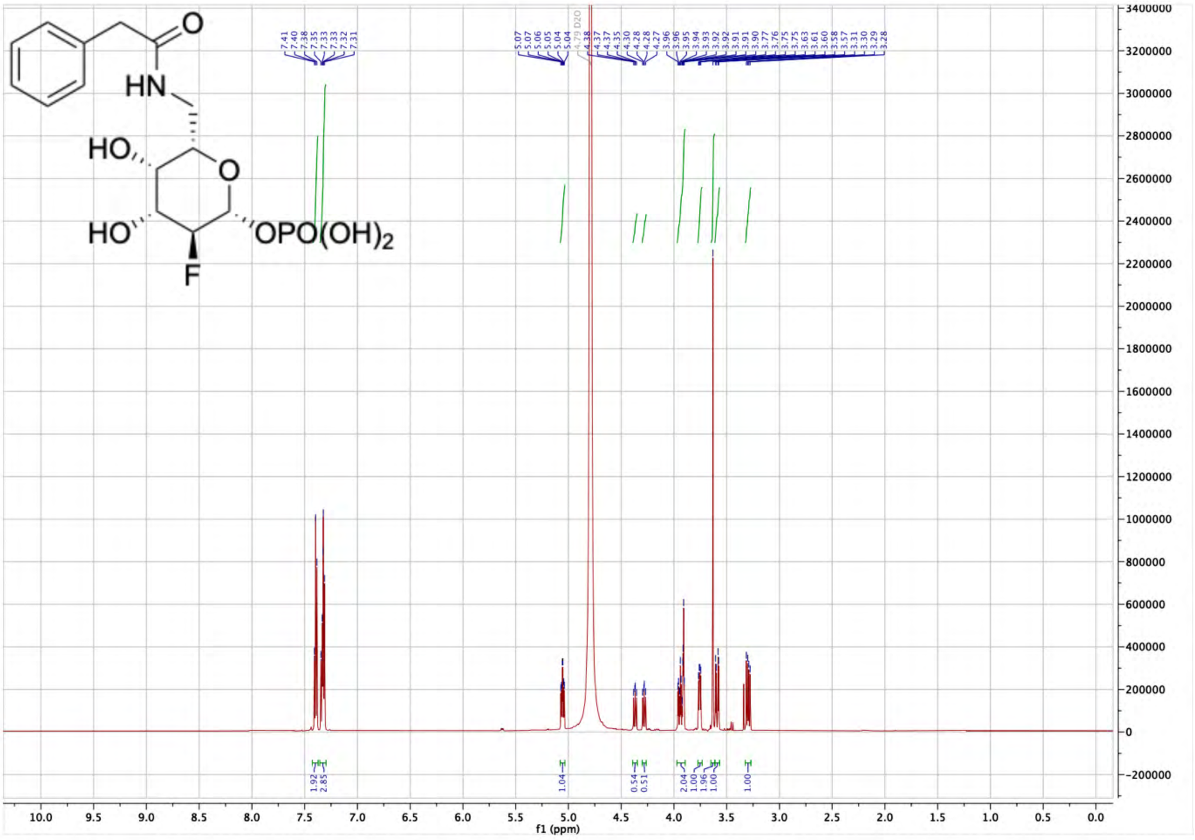

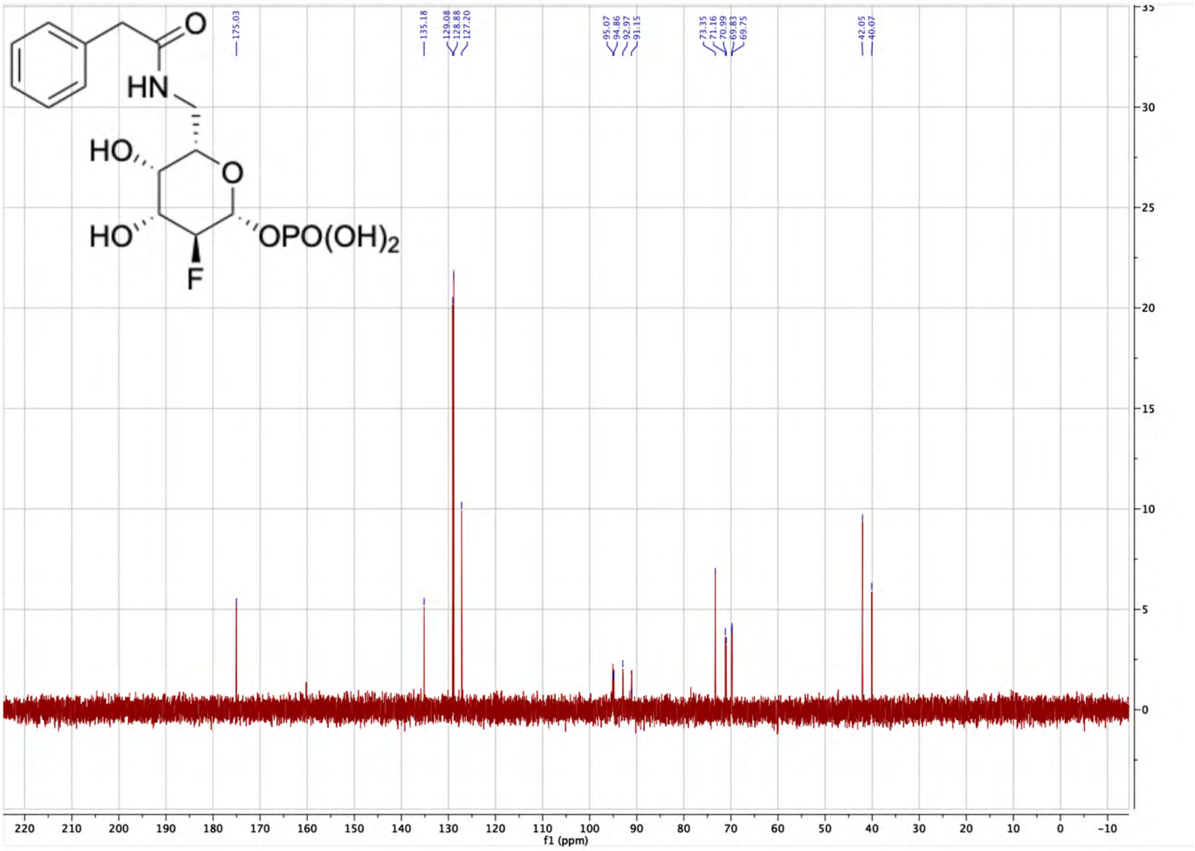

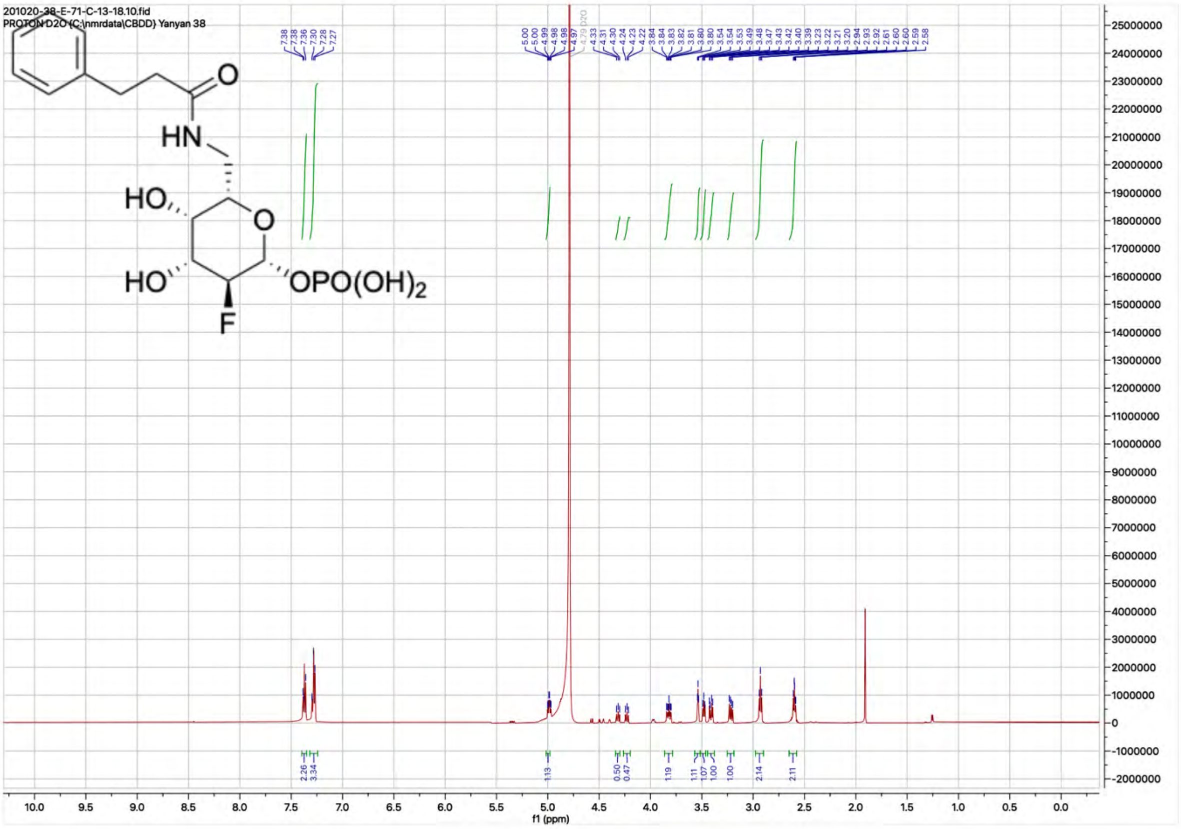
201020-38-E-71-C-13-18.10.fid
PROTON D2O {C:\nmrdata\CBDD} Yanyan 38
HN
HO
O
HO
OPO(OH)2
F
f1 (ppm)

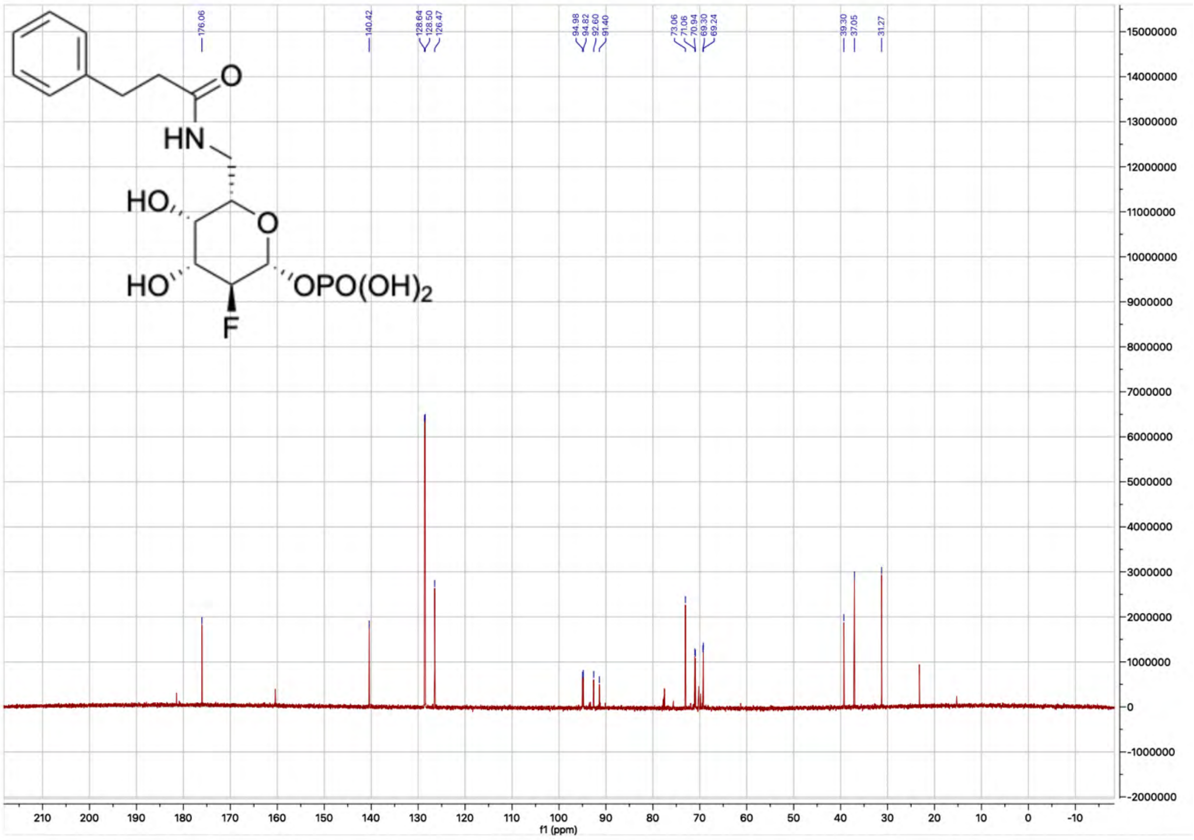
HN
HO
O
HO
OPO(OH)2
F
f1 (ppm)

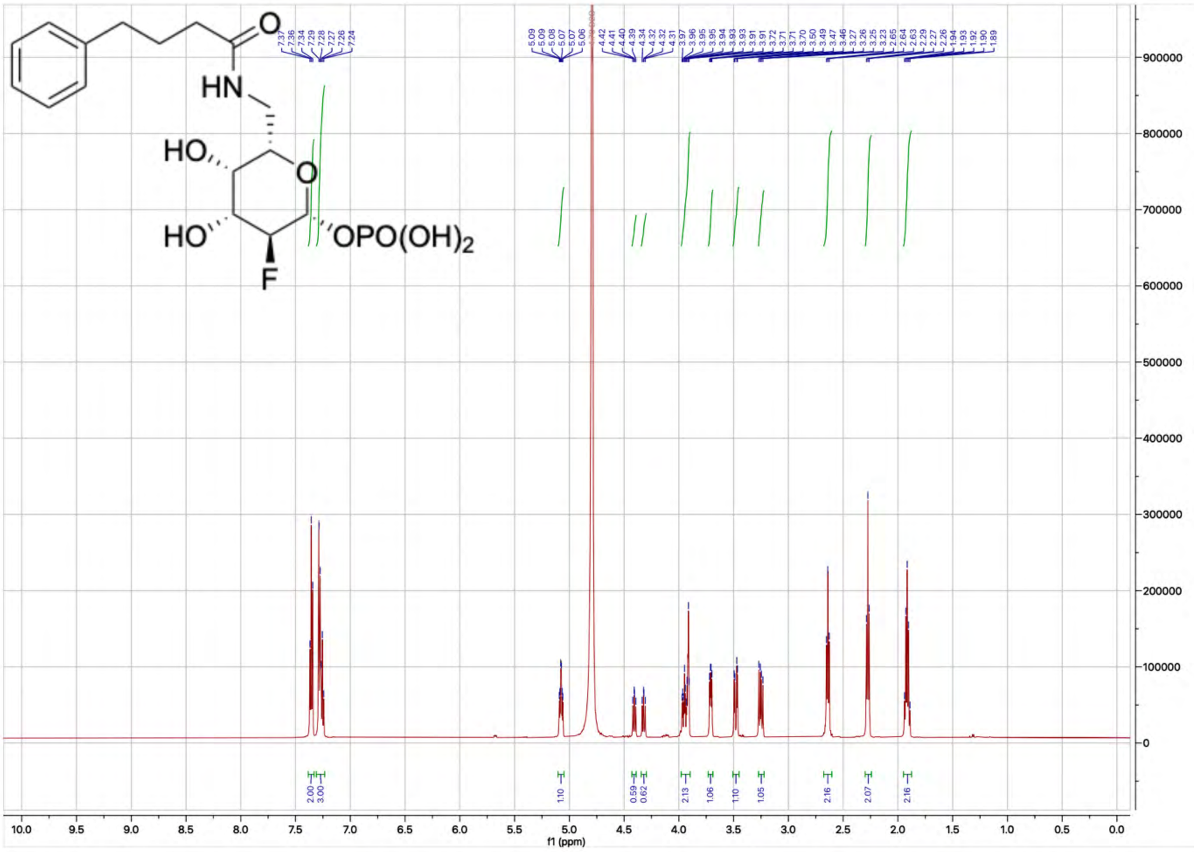

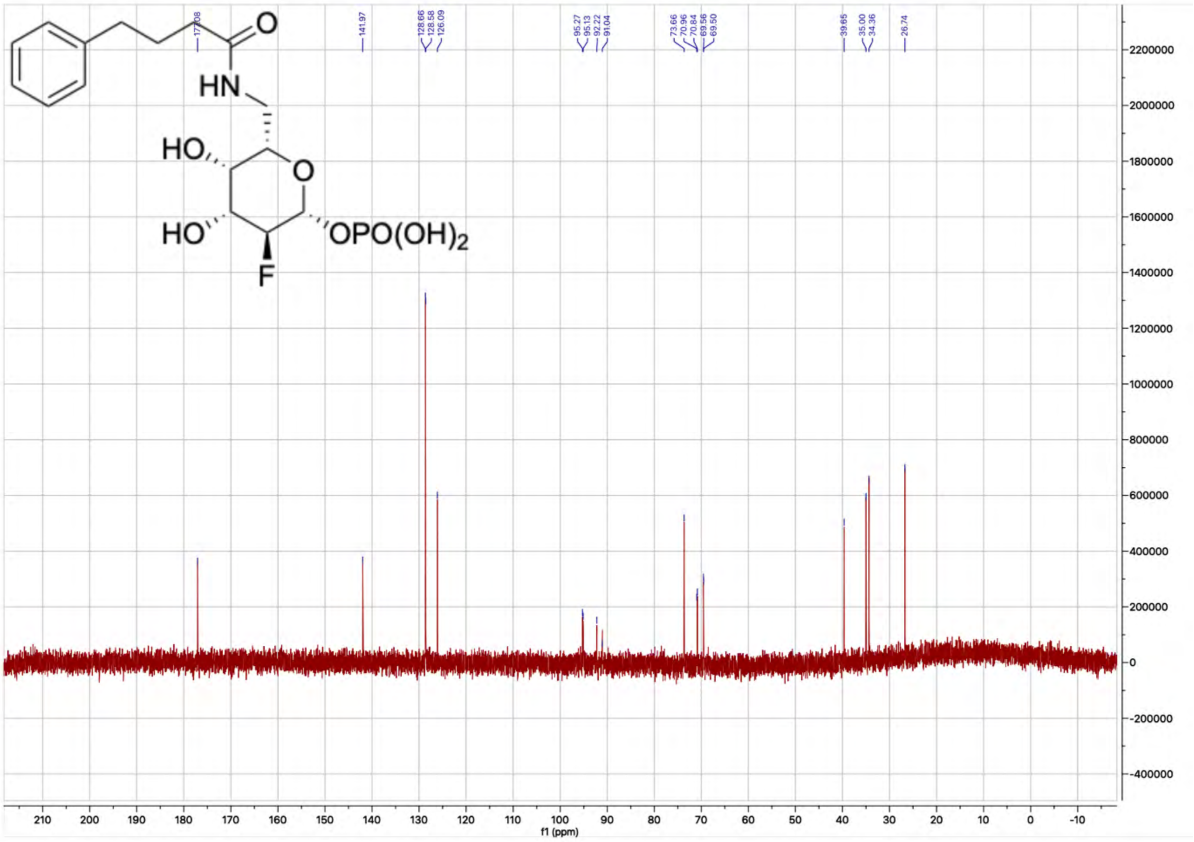

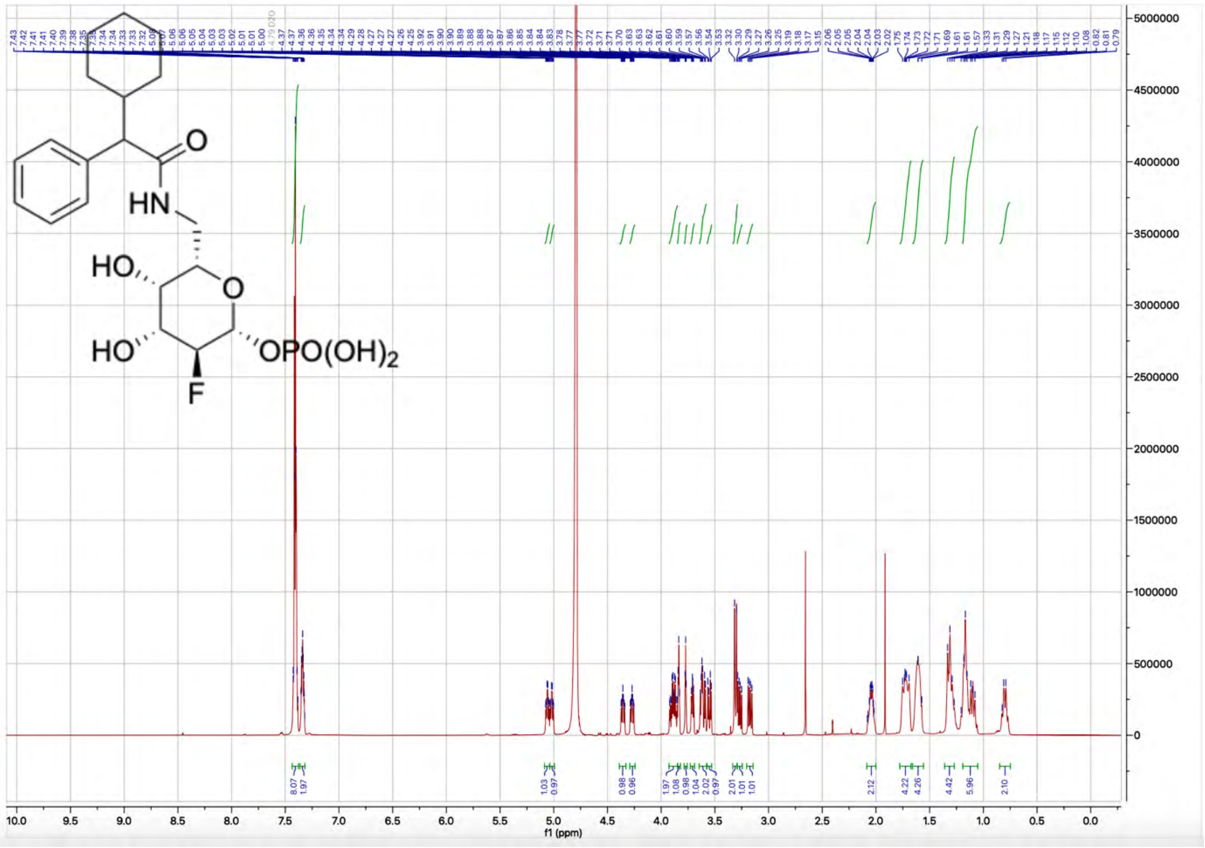

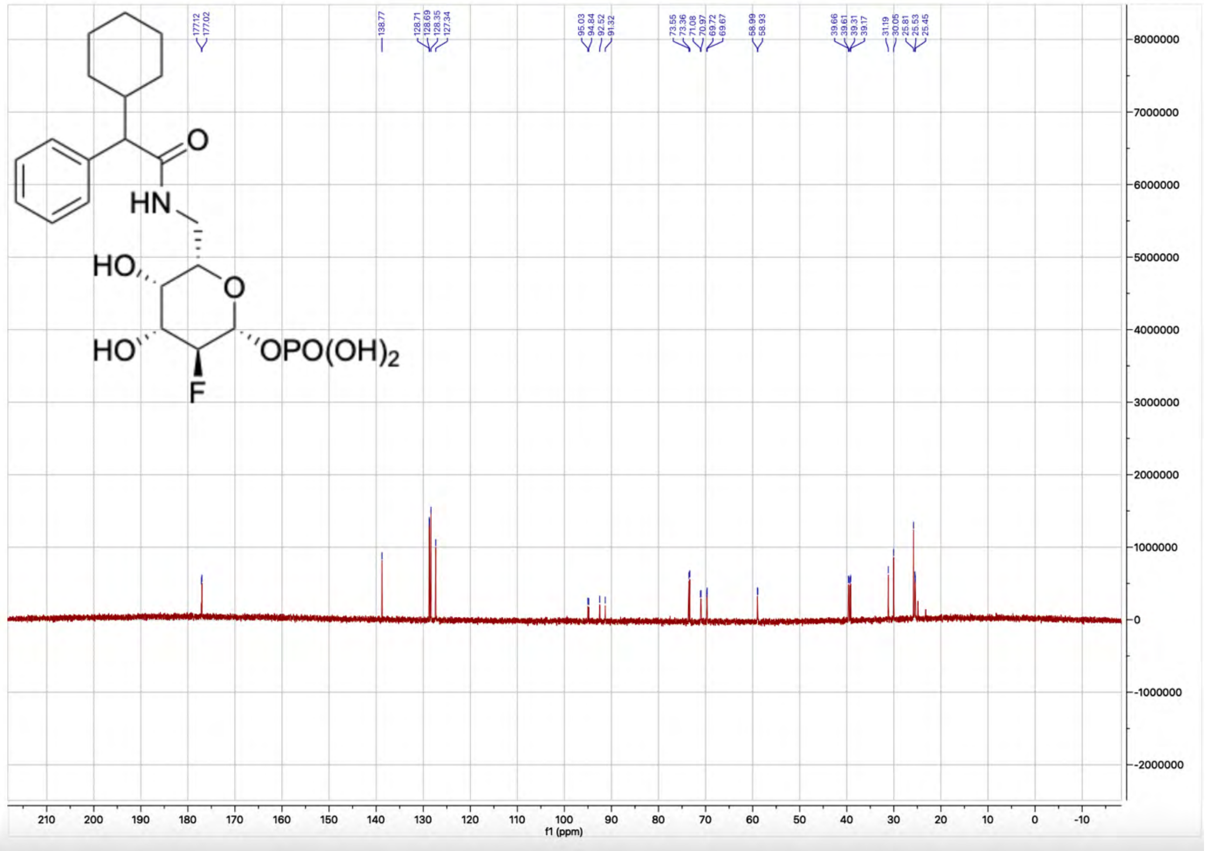

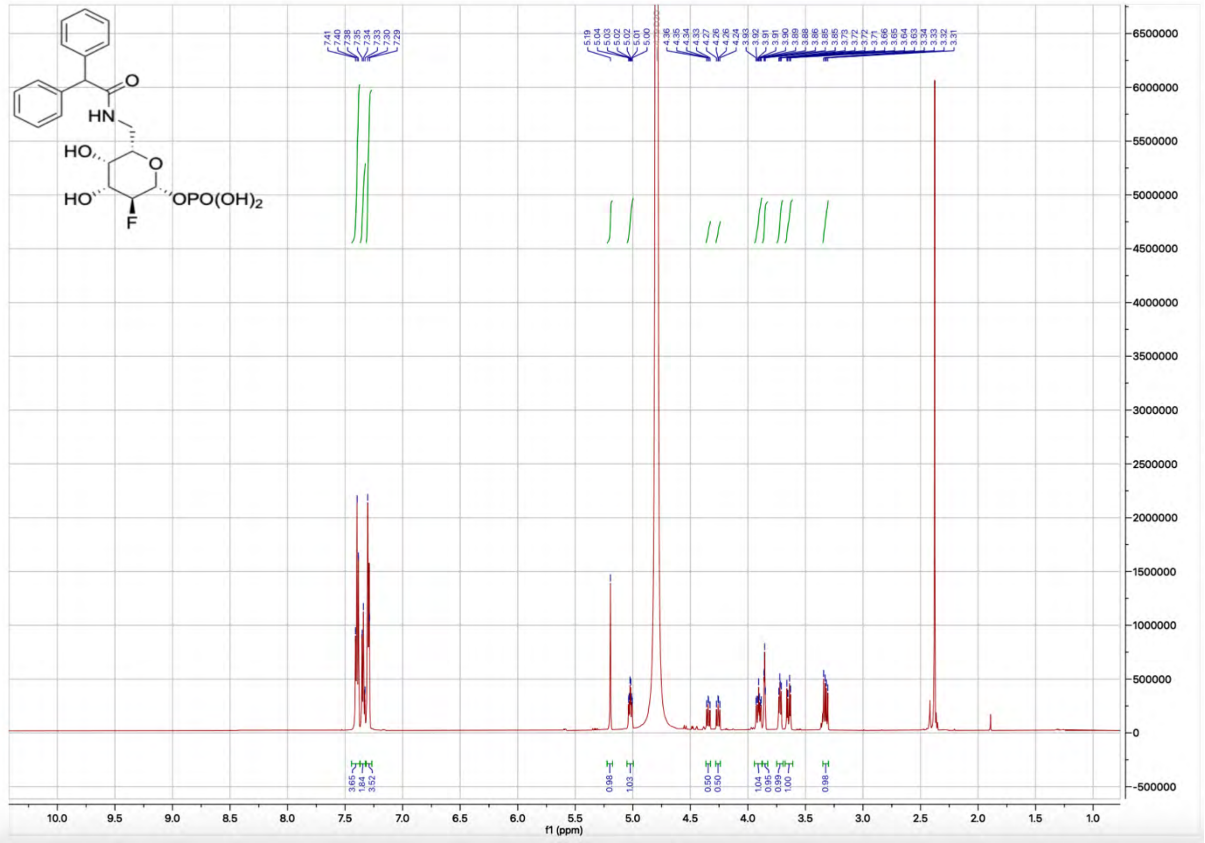
HN
HO
O
HO
OPO(OH)2
F
7.41 7.40 7.38 7.35 7.34 7.33 7.30 7.29
5.19 5.04 5.03 5.02 5.02 5.01 5.00
4.36 4.35 4.34 4.33 4.27 4.26 4.26 4.24
3.93 3.92 3.91 3.91 3.90 3.89 3.88 3.86 3.85 3.85 3.73 3.72 3.72 3.71 3.66 3.65 3.64 3.63 3.34 3.33 3.32 3.31
3.65 1.84 3.52
0.98 1.03
0.50 0.50
1.04 0.95 0.99 1.00
0.98
f1 (ppm)

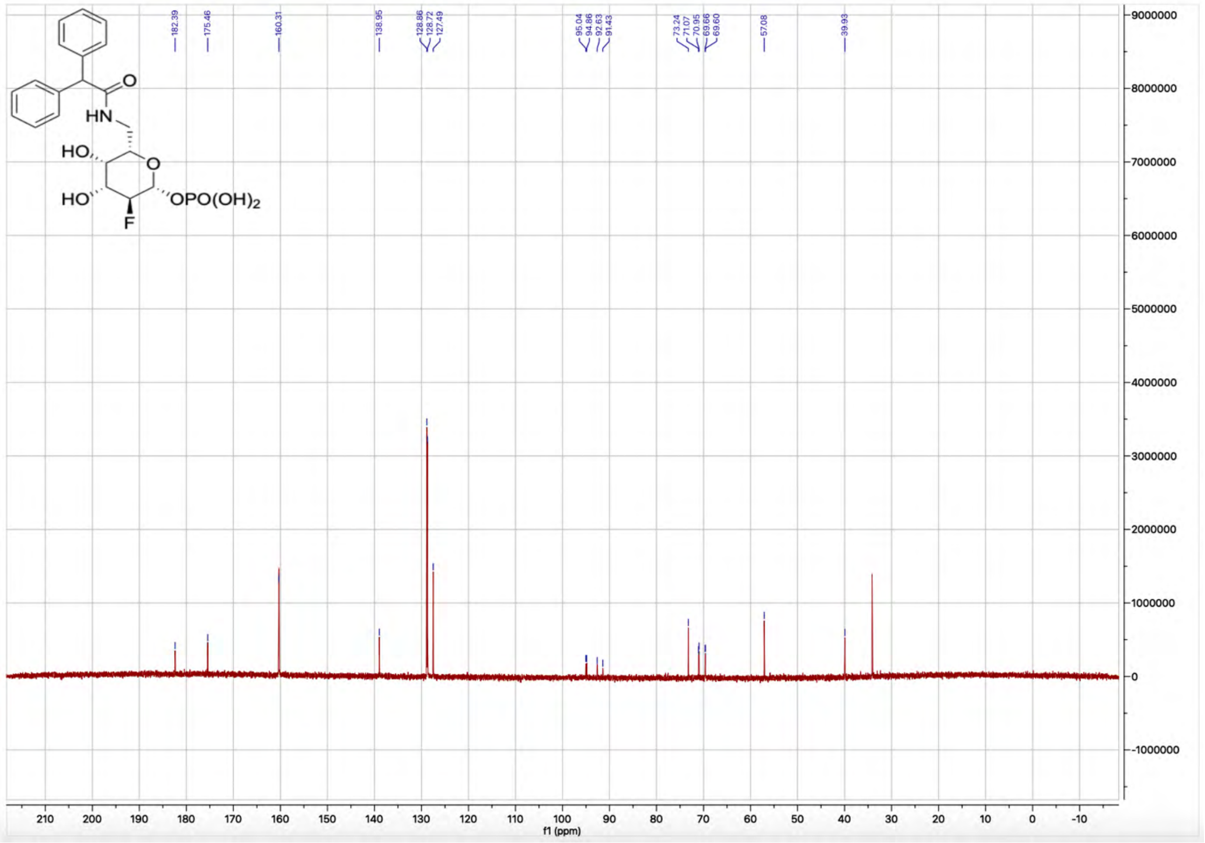
HN
HO
O
HO
OPO(OH)2
F
182.39
175.46
160.31
138.95
128.86 128.72 127.49
95.04 94.86 92.63 91.43
73.24 71.07 70.95 69.66 69.60
57.08
39.93
f1 (ppm)

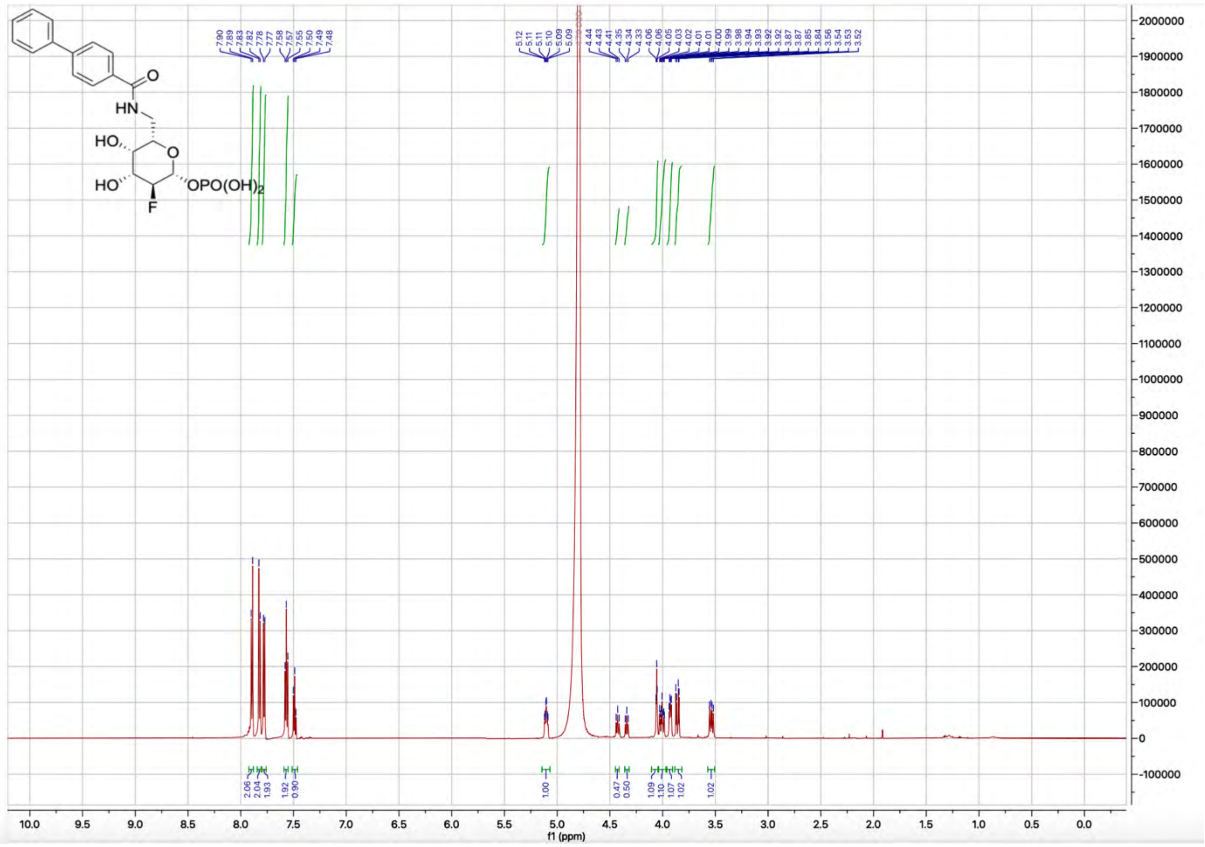

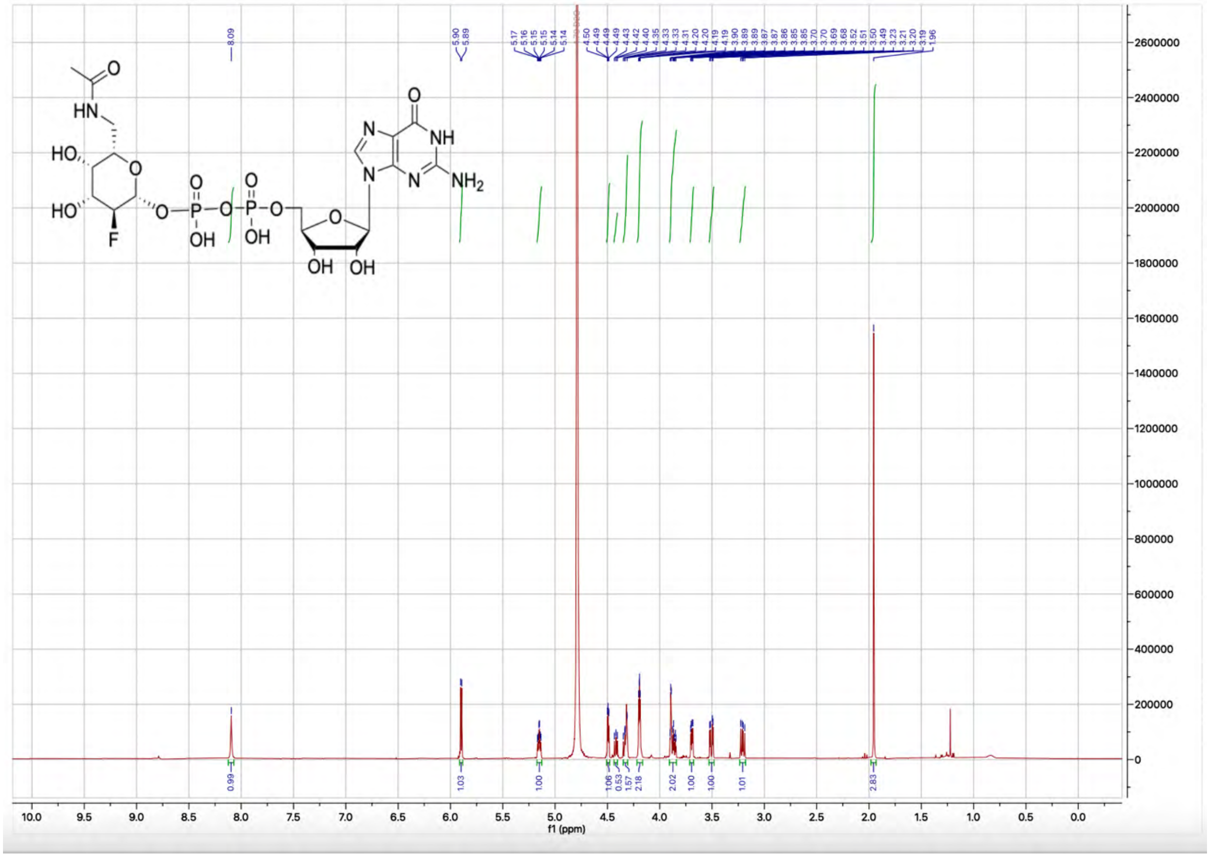

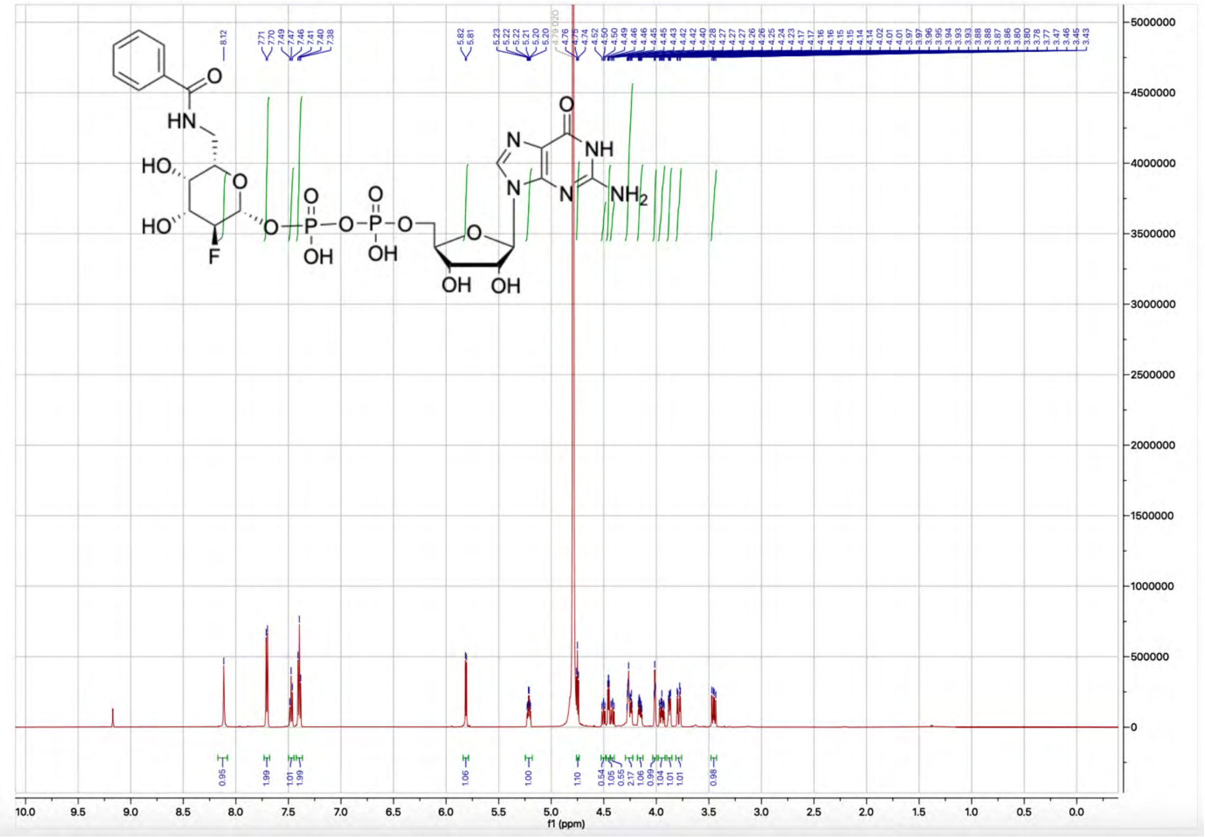

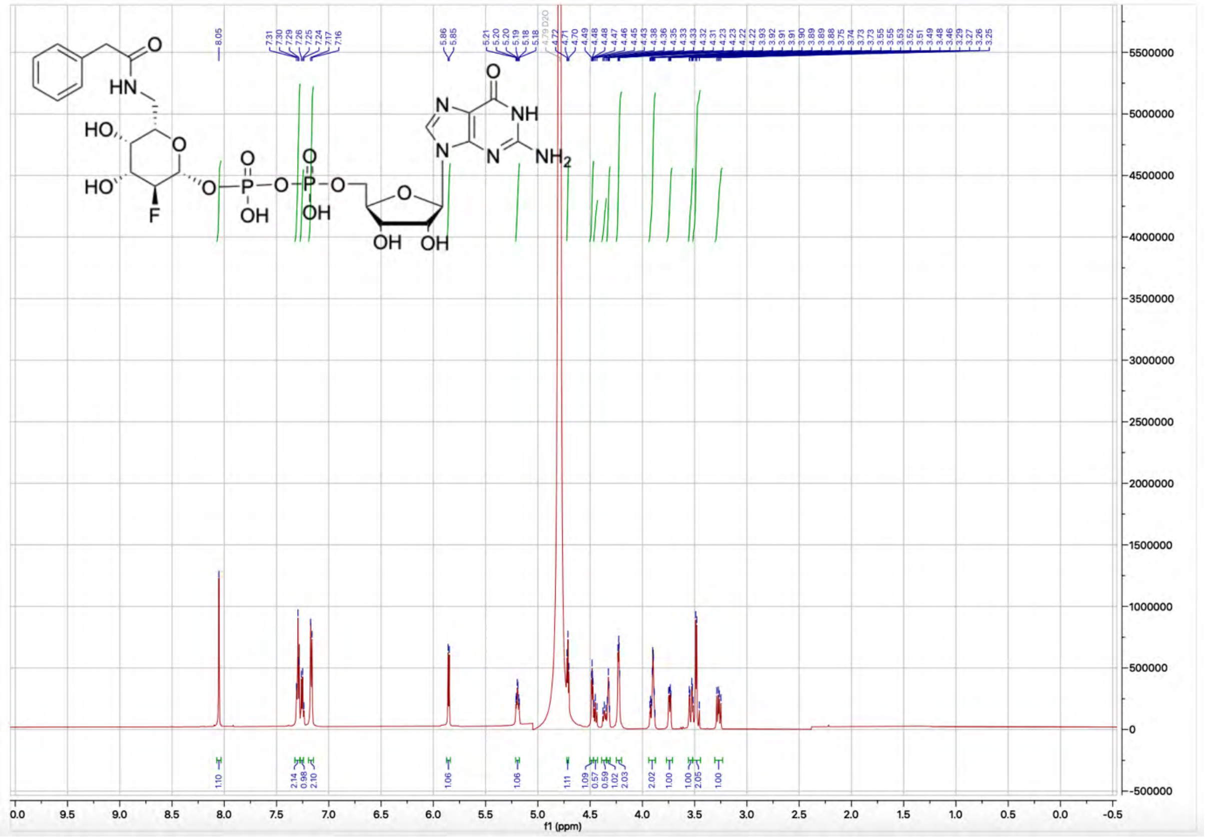

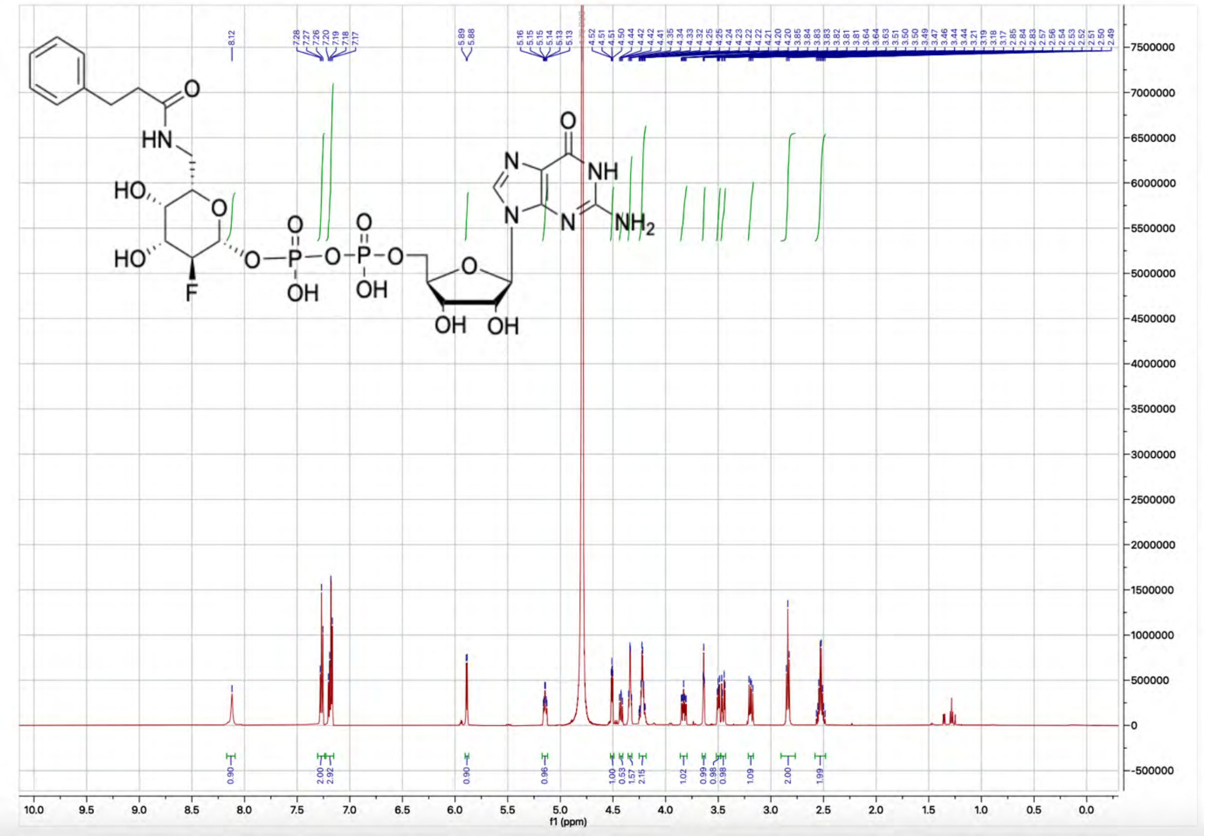

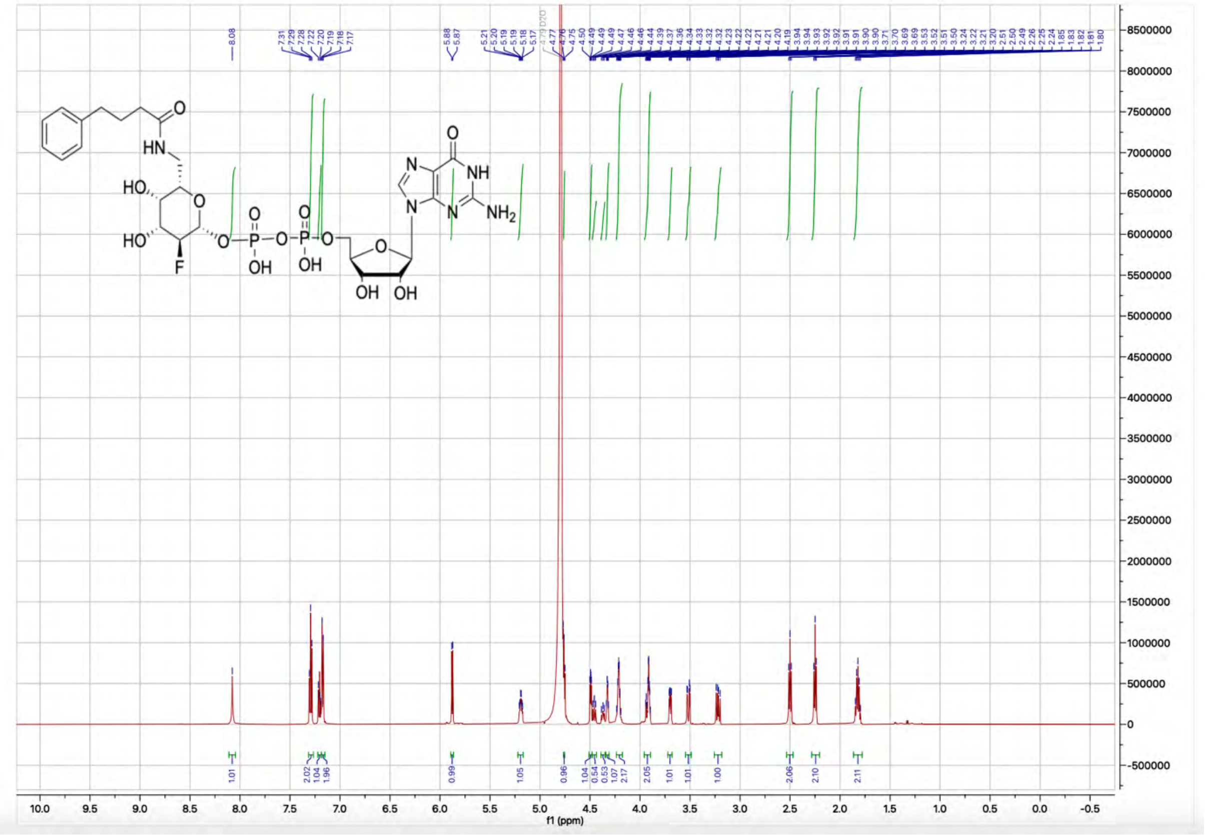

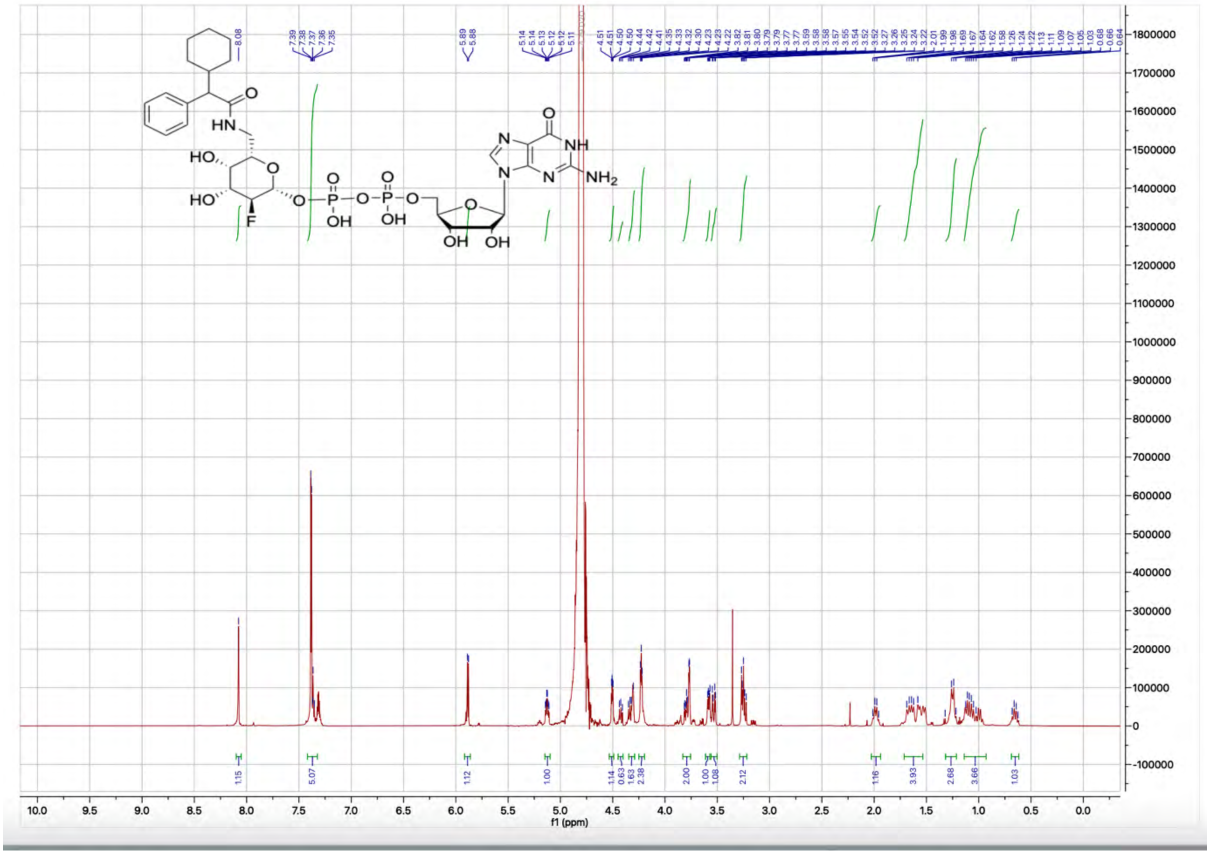

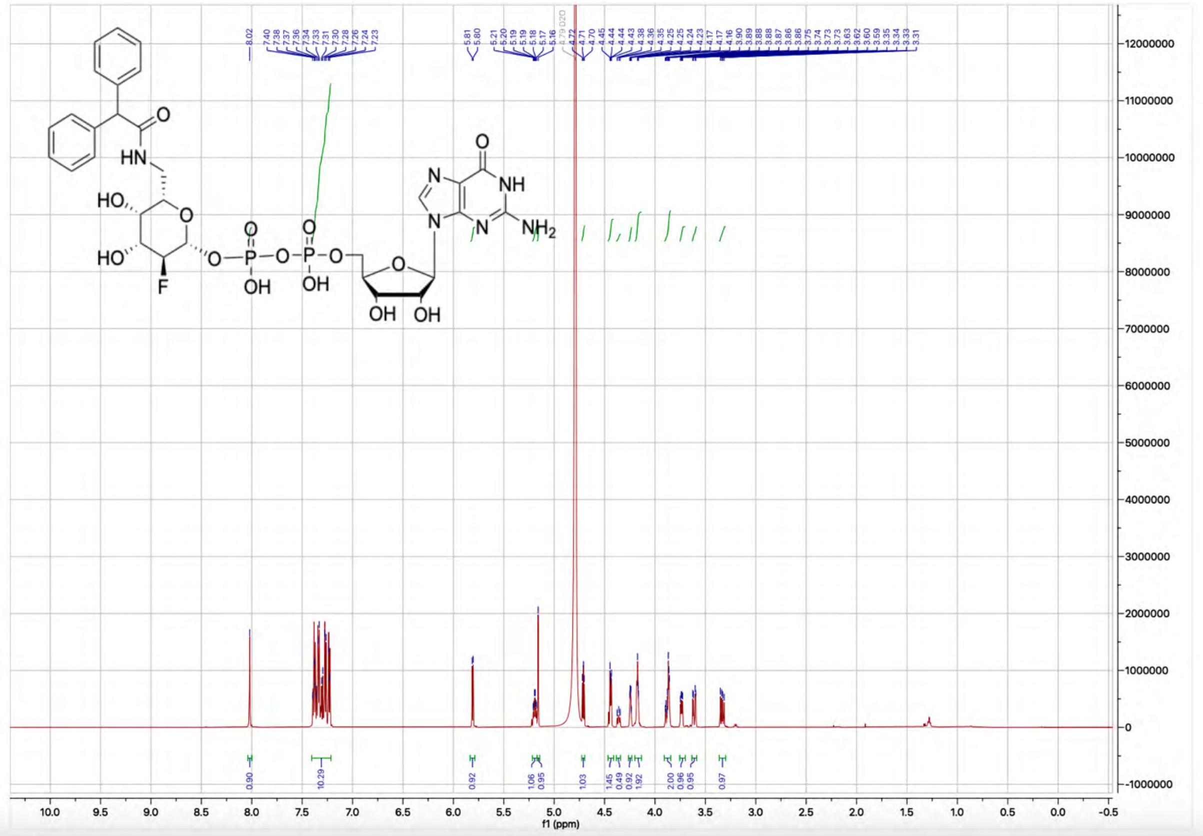

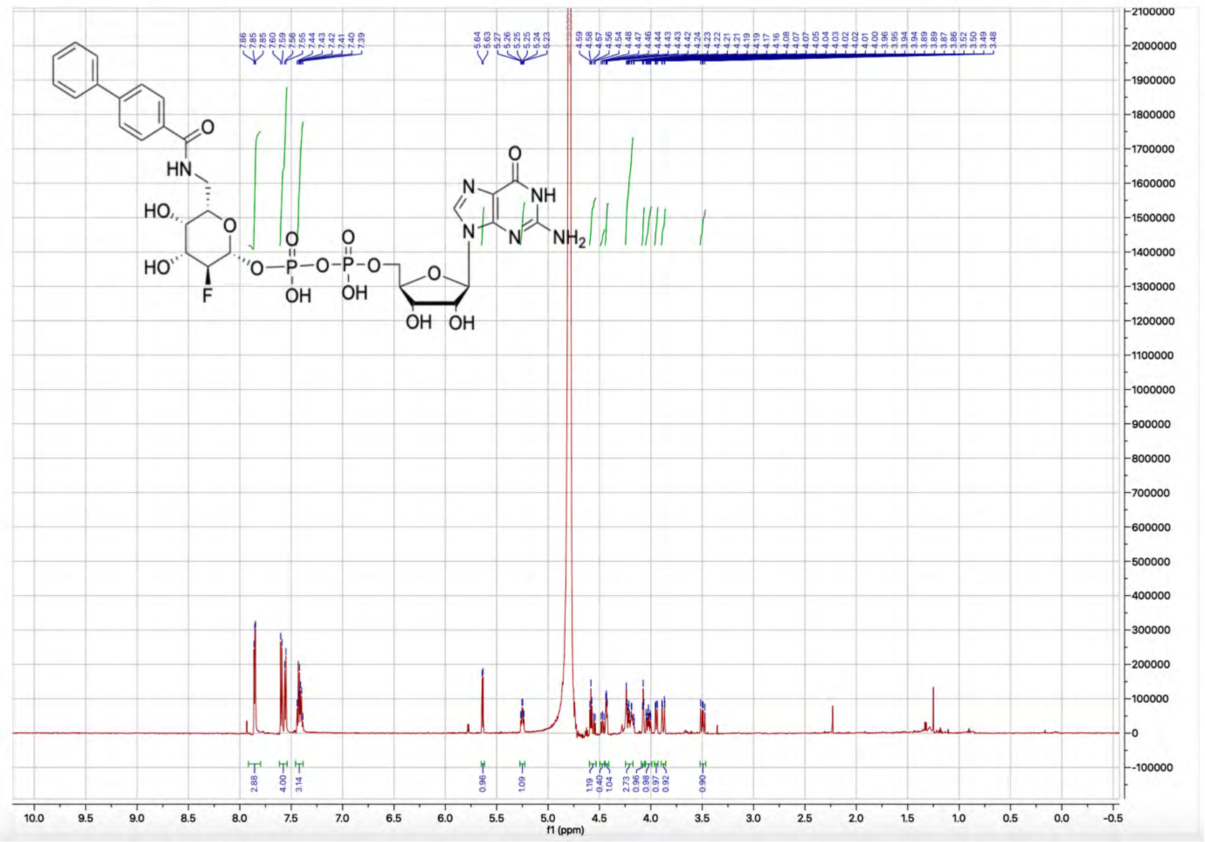

HN
HO
HO
F
O
OH
OH
OH
NH
N
NH2
f1 (ppm)


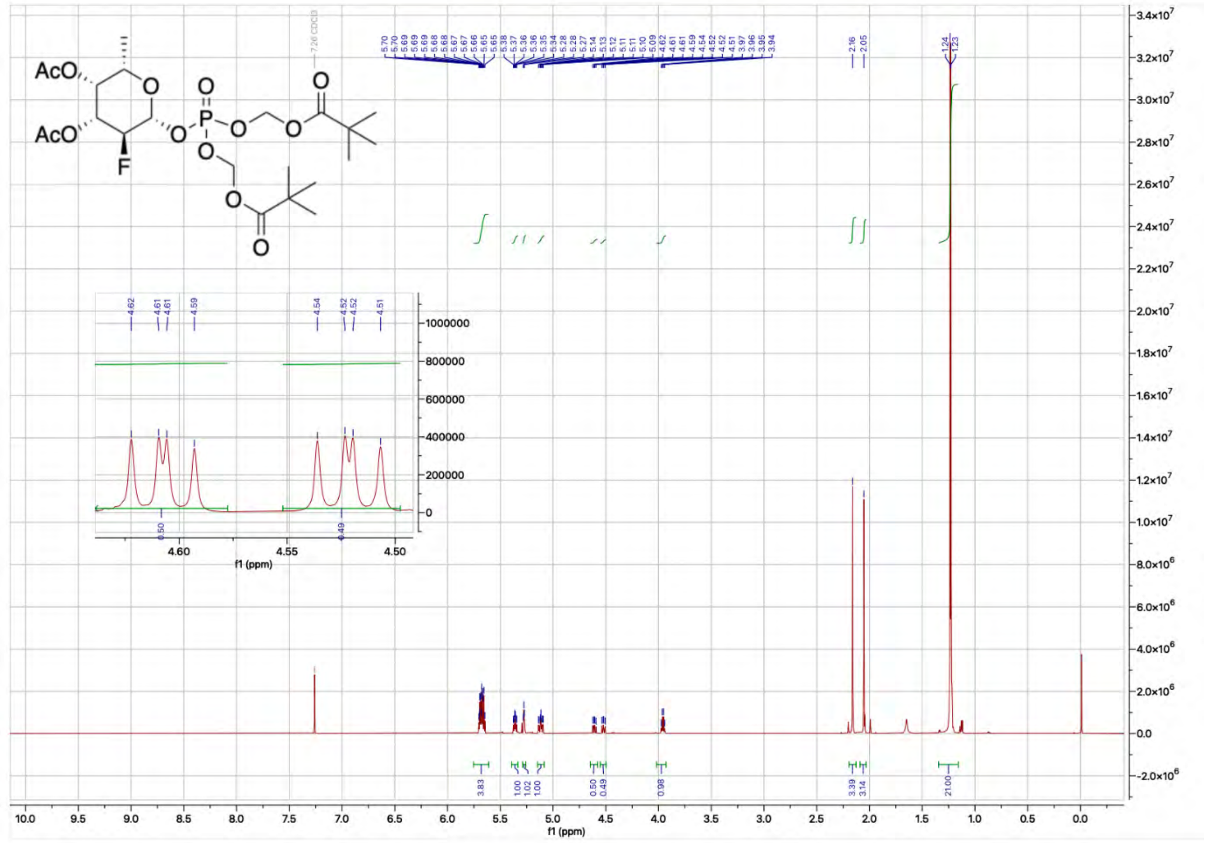

AcO
AcO
F
O
7.26 CDCl3
f1 (ppm)

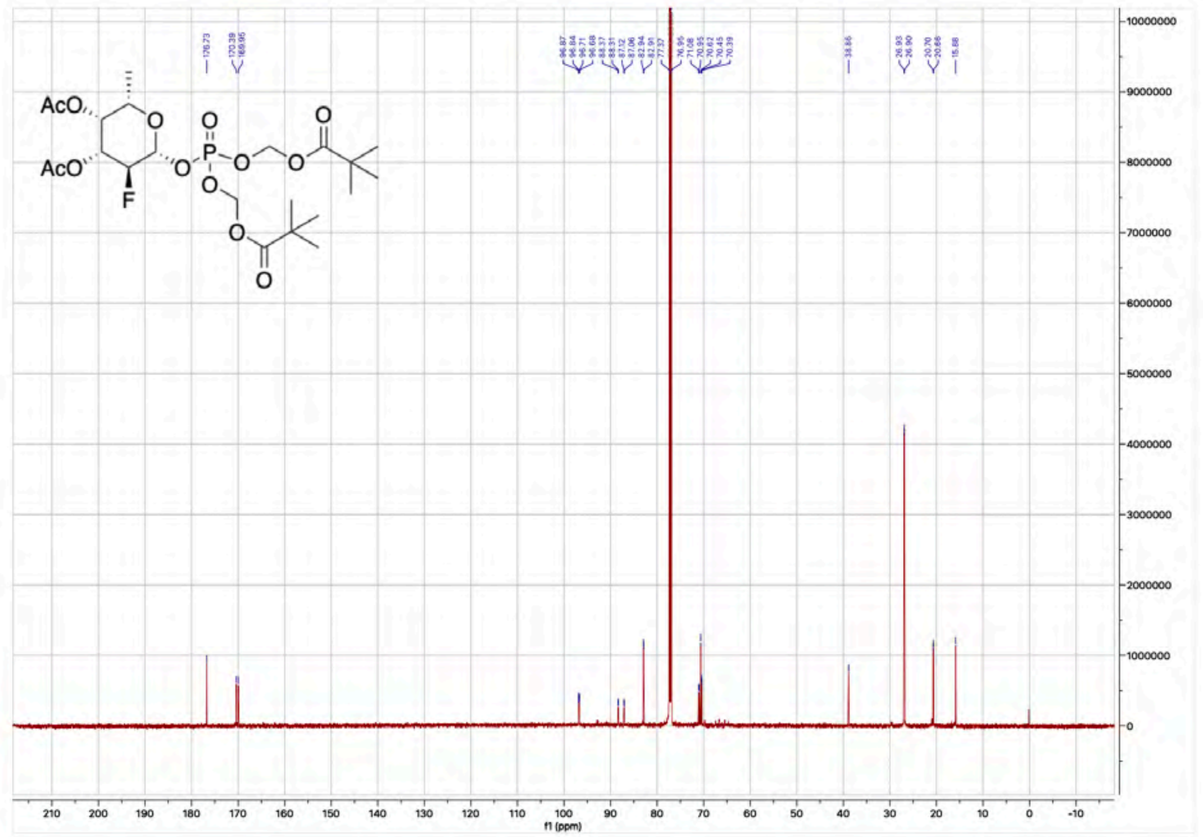
AcO
AcO
F
O
P
f1 (ppm)

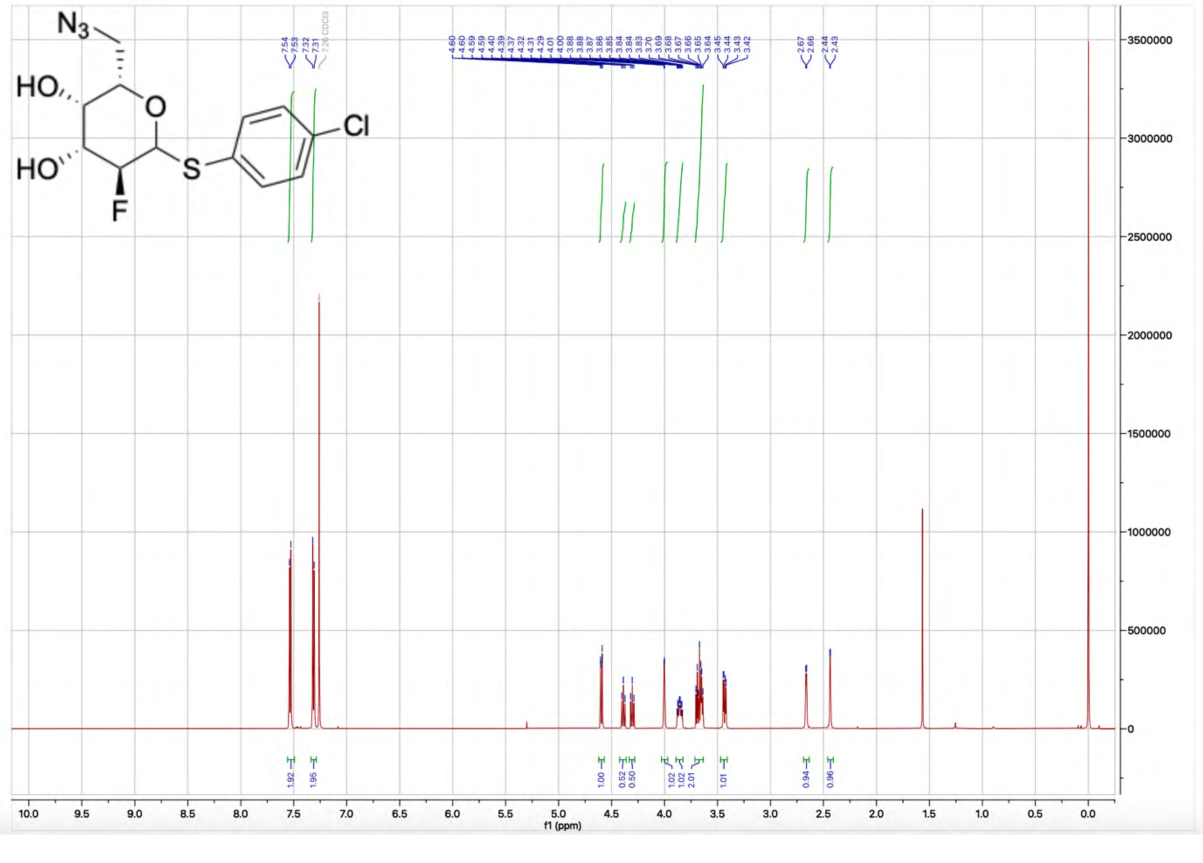
N3
HO
HO
F
S
Cl
f1 (ppm)

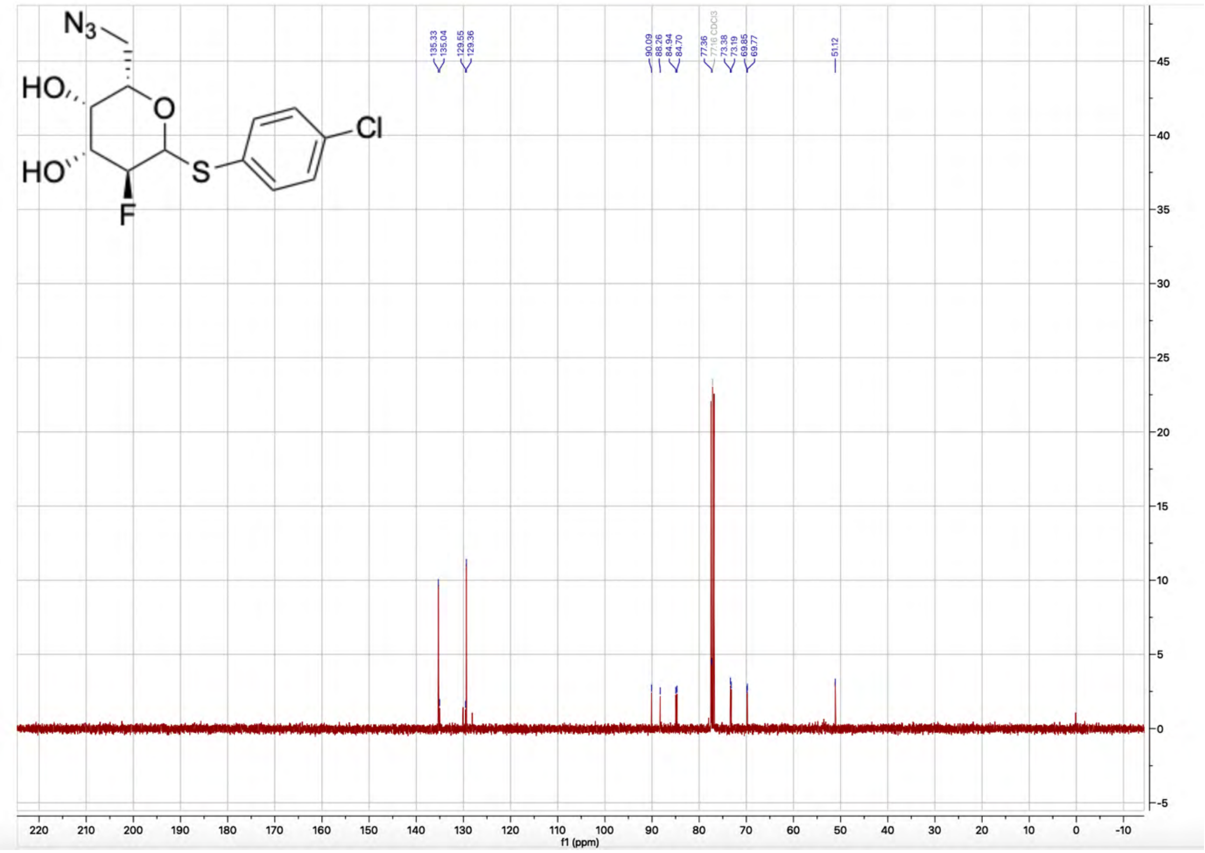

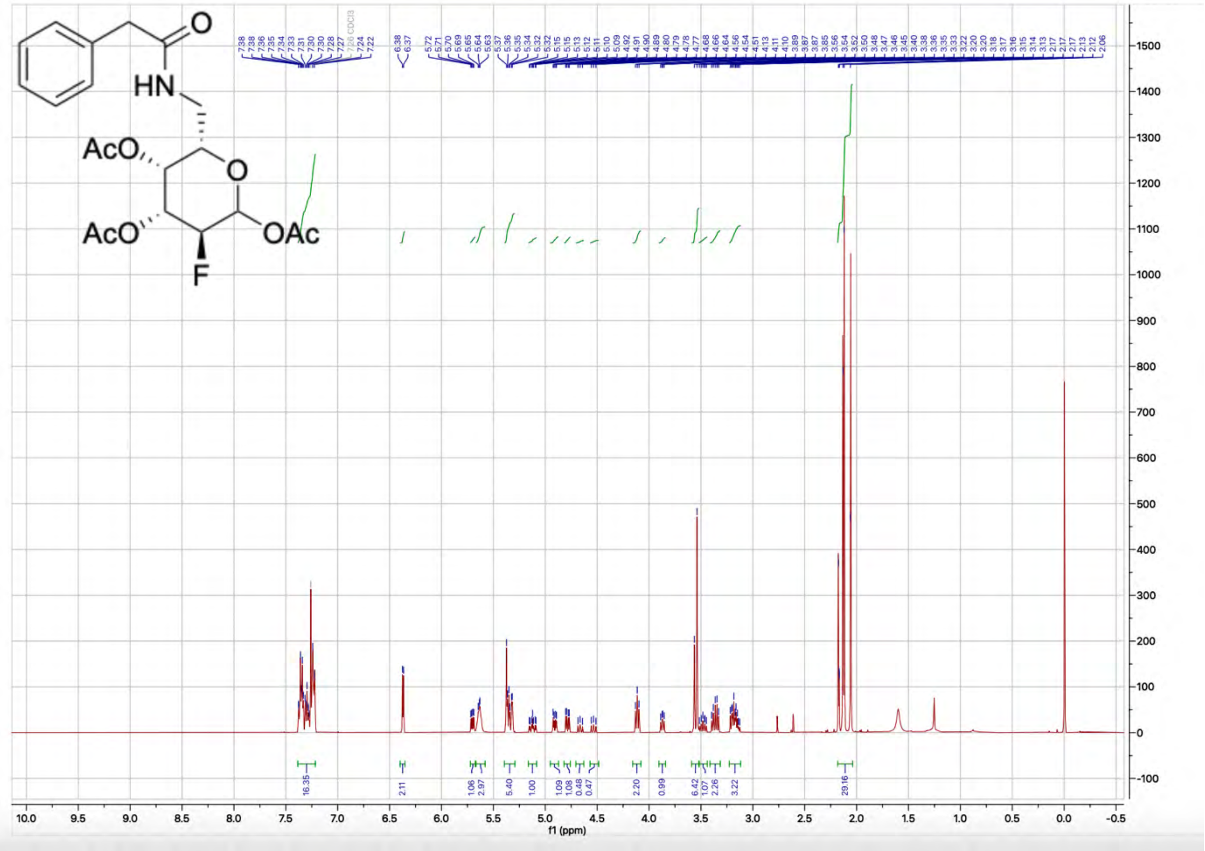

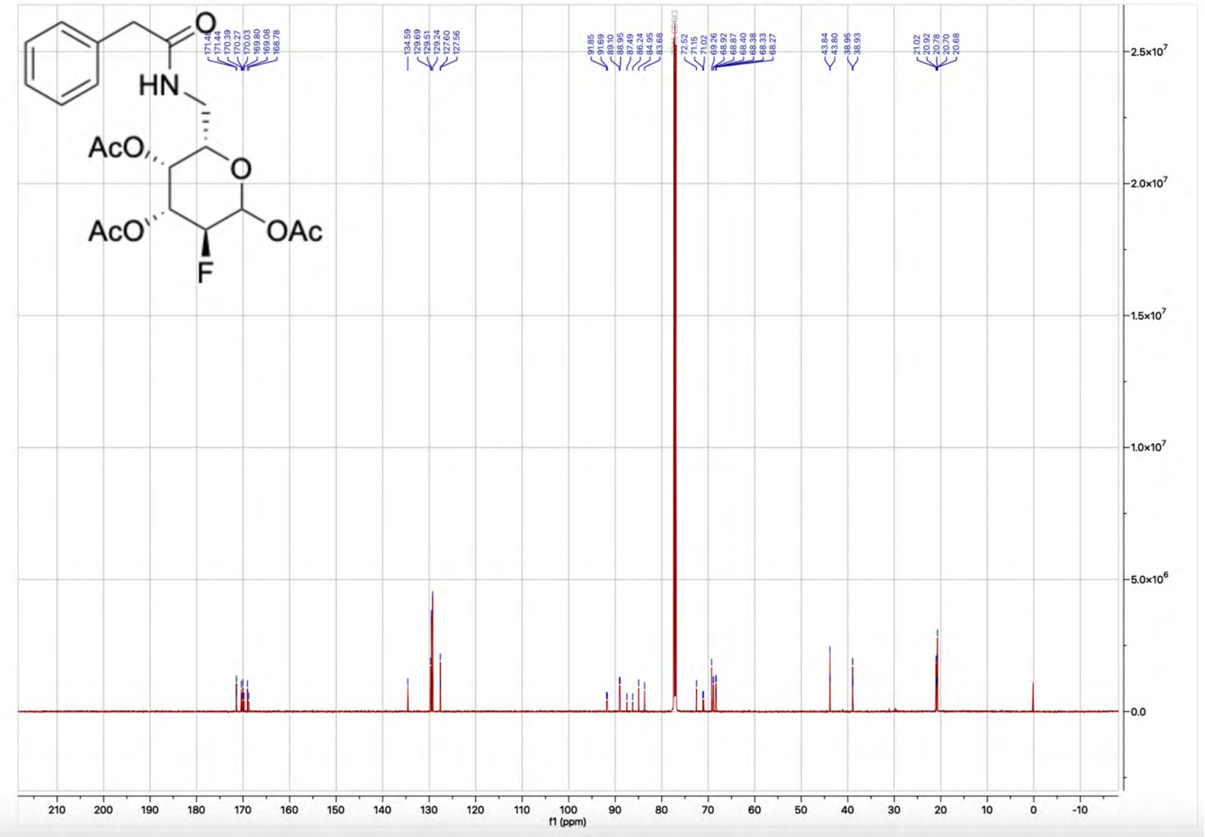

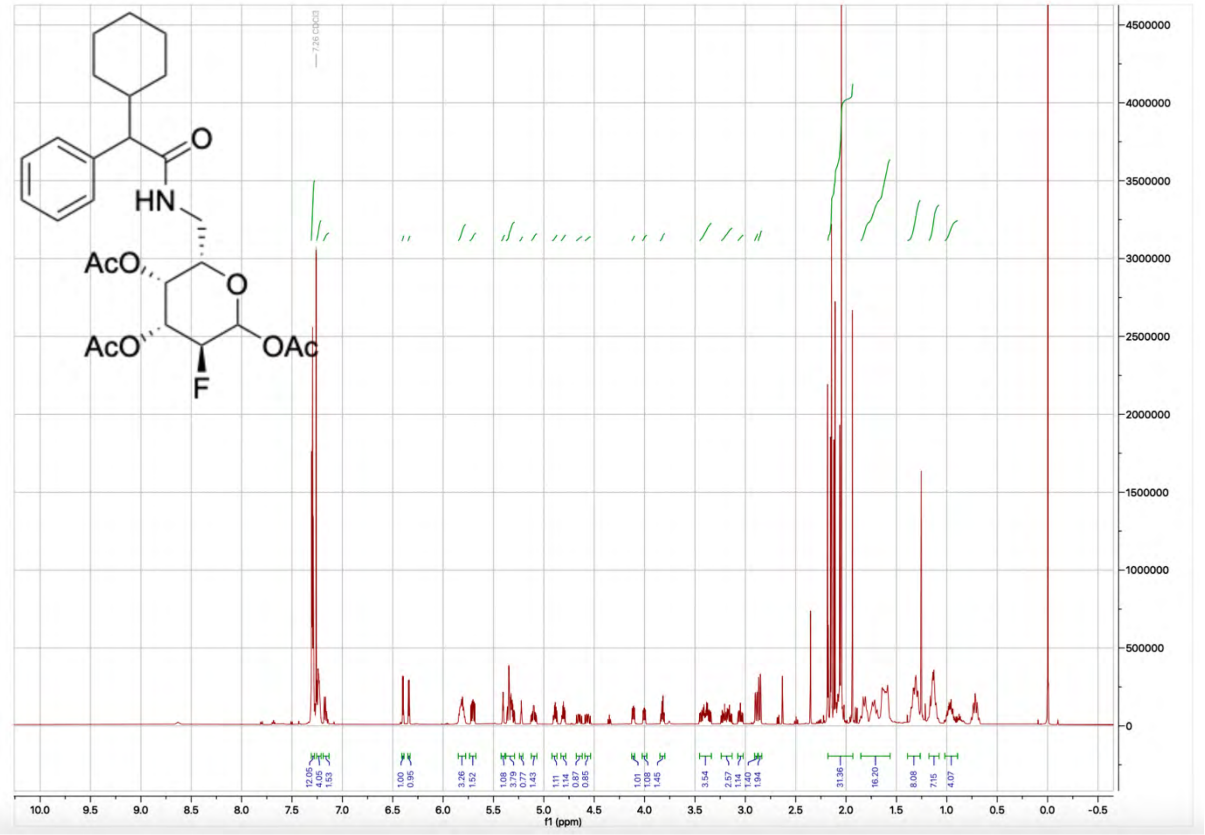

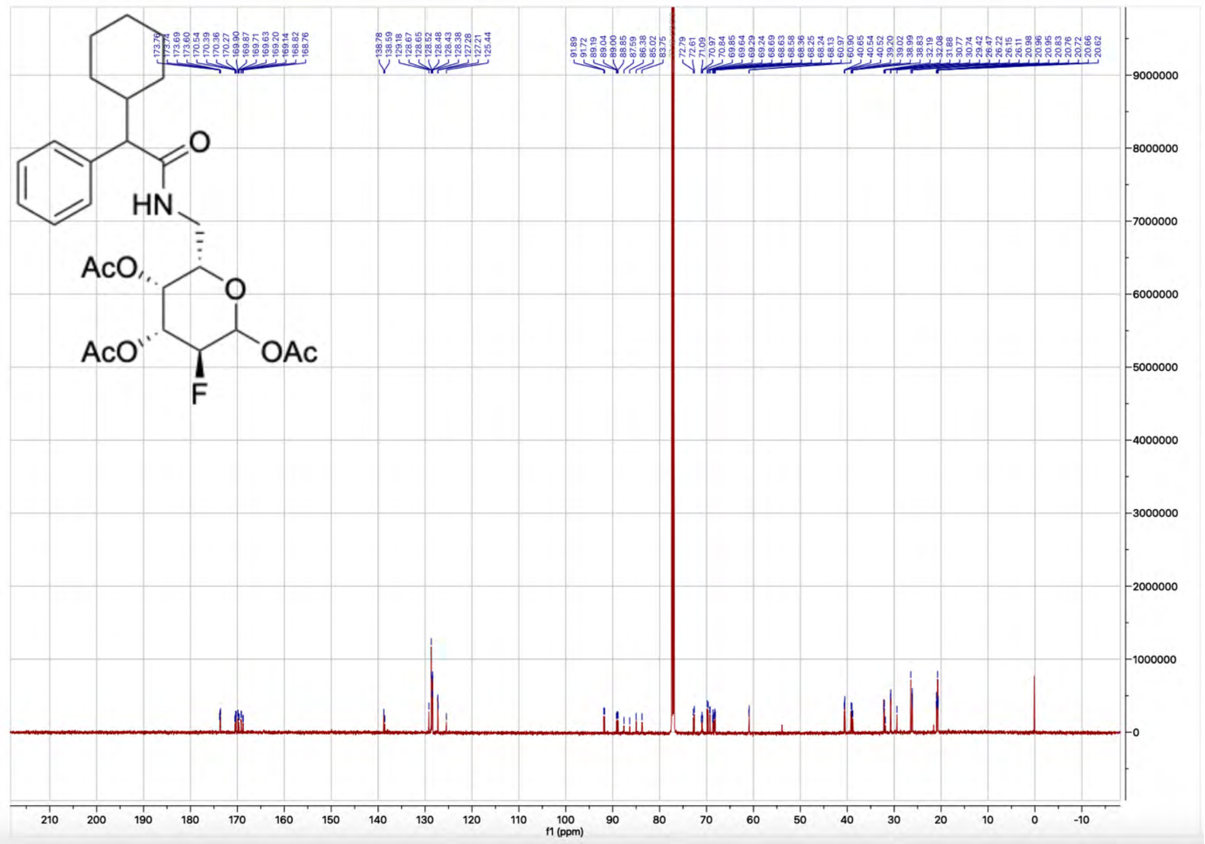
AcO
AcO
OAc
F
HN
O
f1 (ppm)

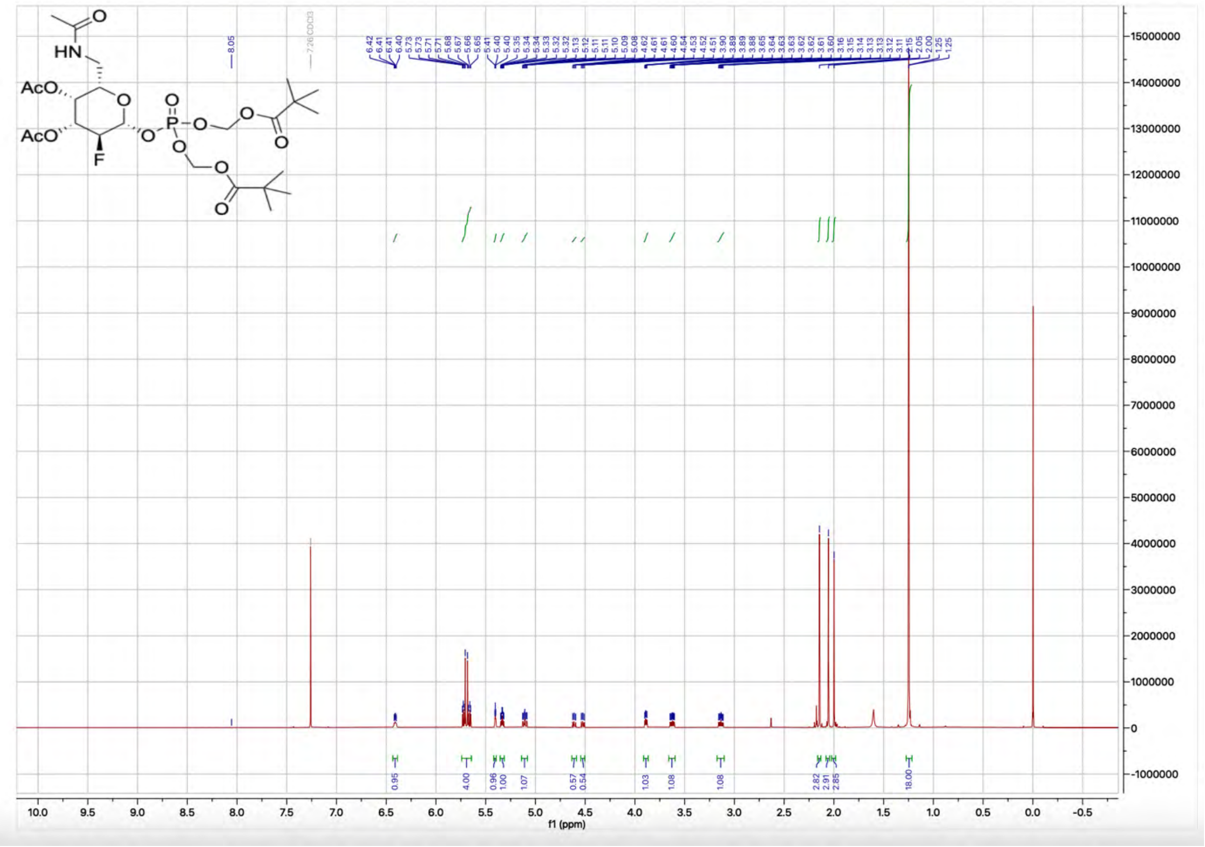
HN
O
AcO
AcO
F
P
f1 (ppm)

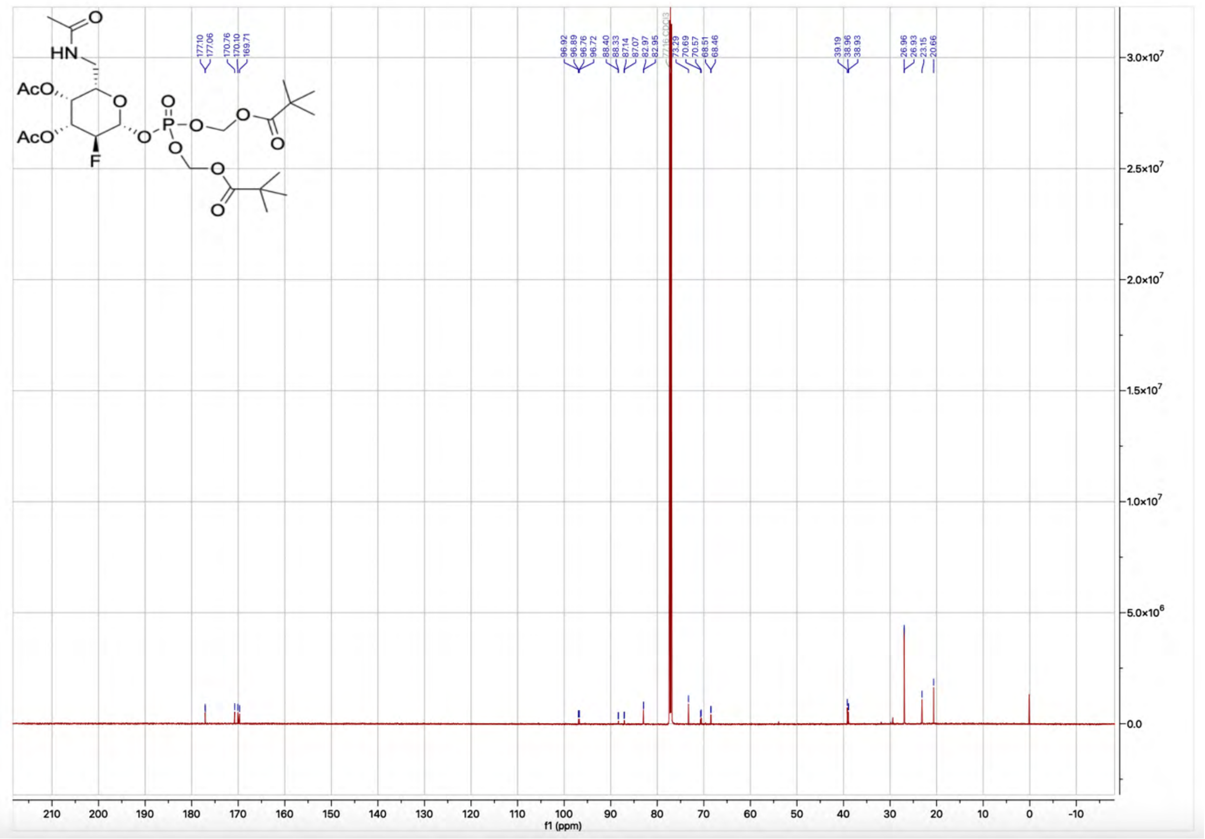
AcO
AcO
HN
F
O
P
f1 (ppm)

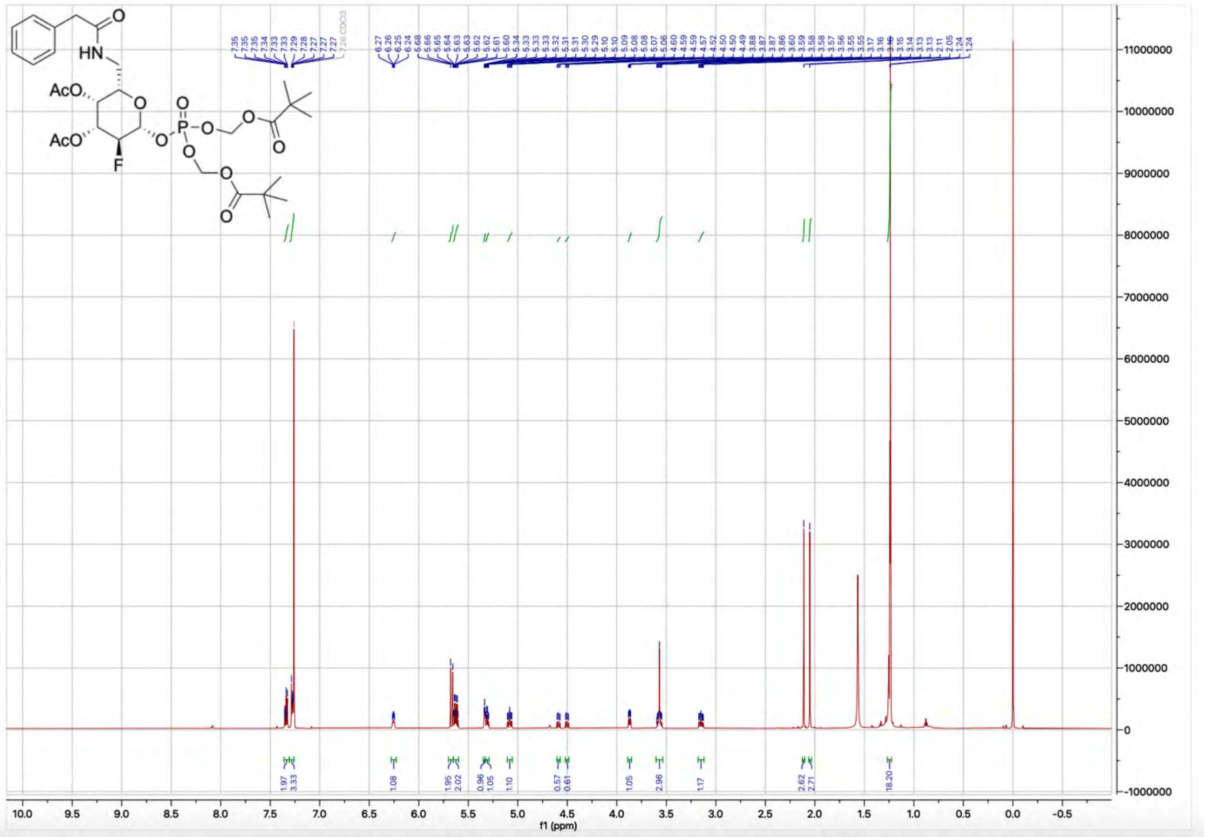
AcO
AcO
HN
F
O
P
f1 (ppm)

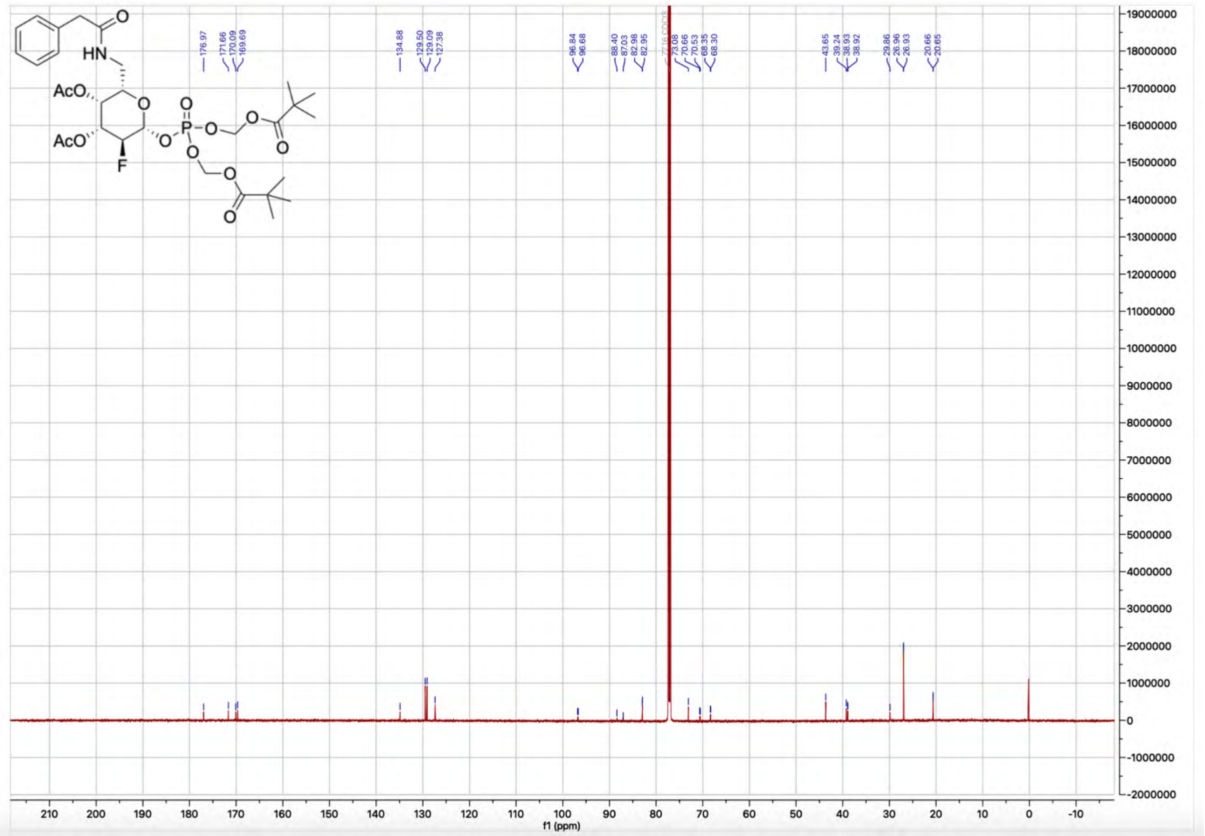

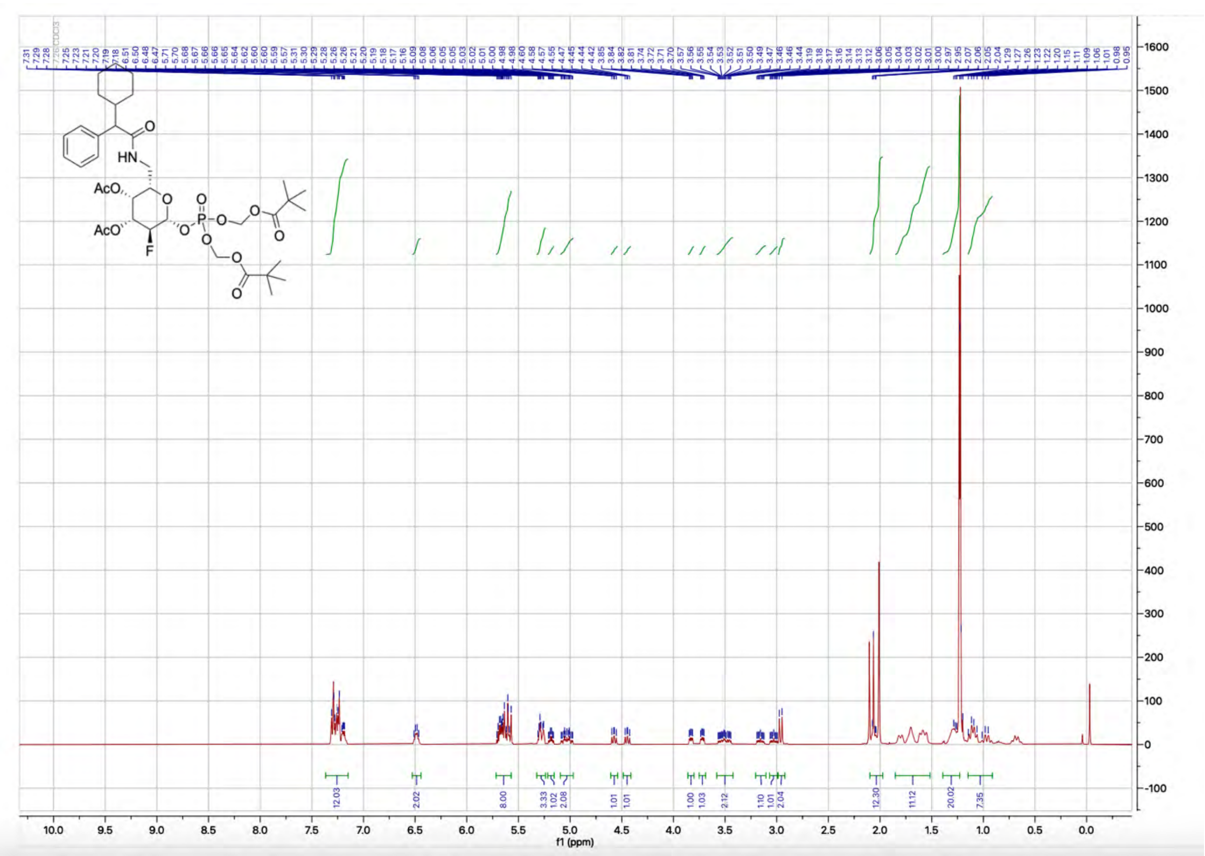

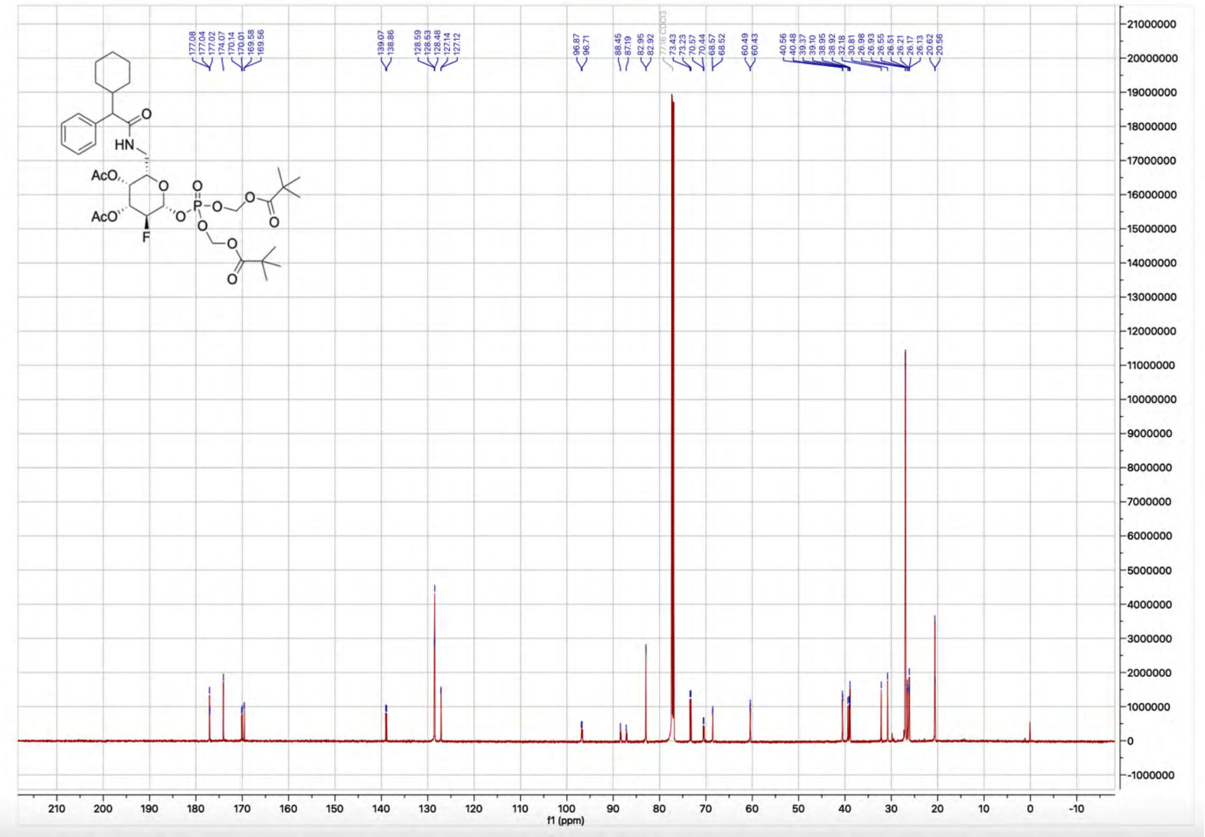

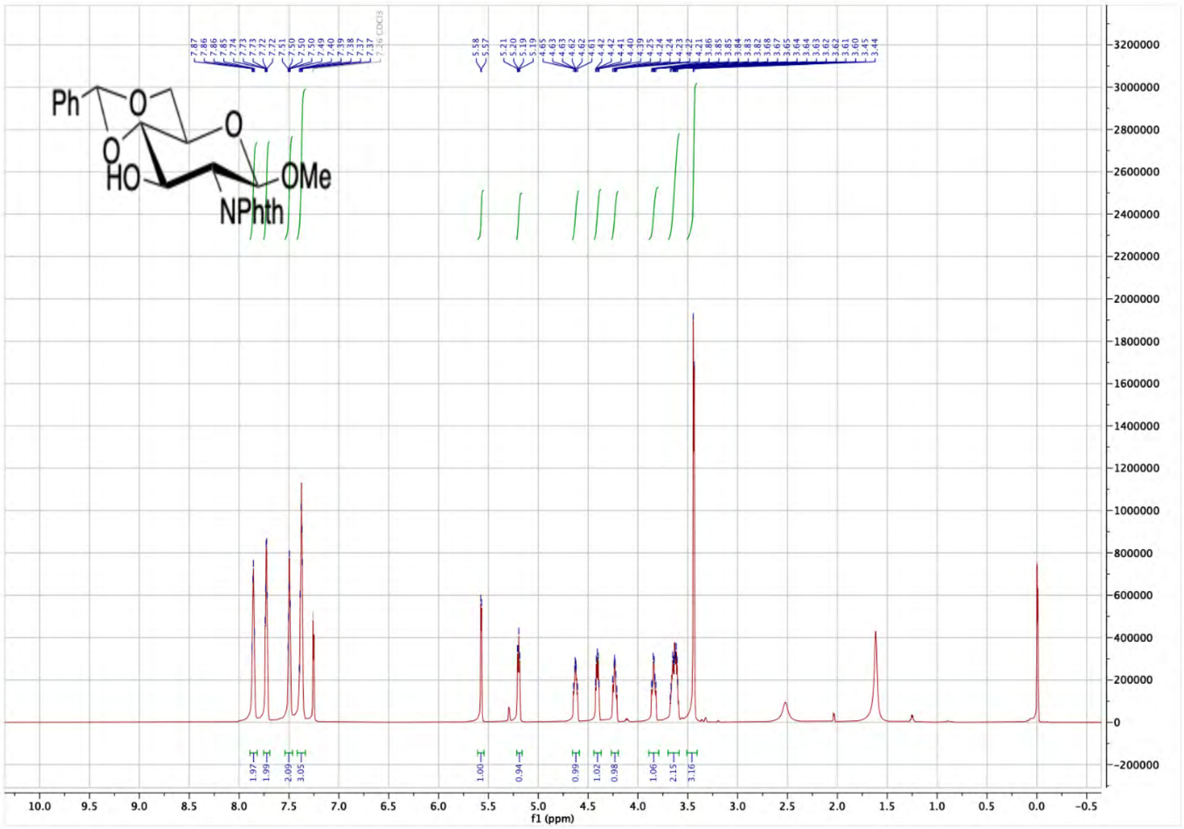

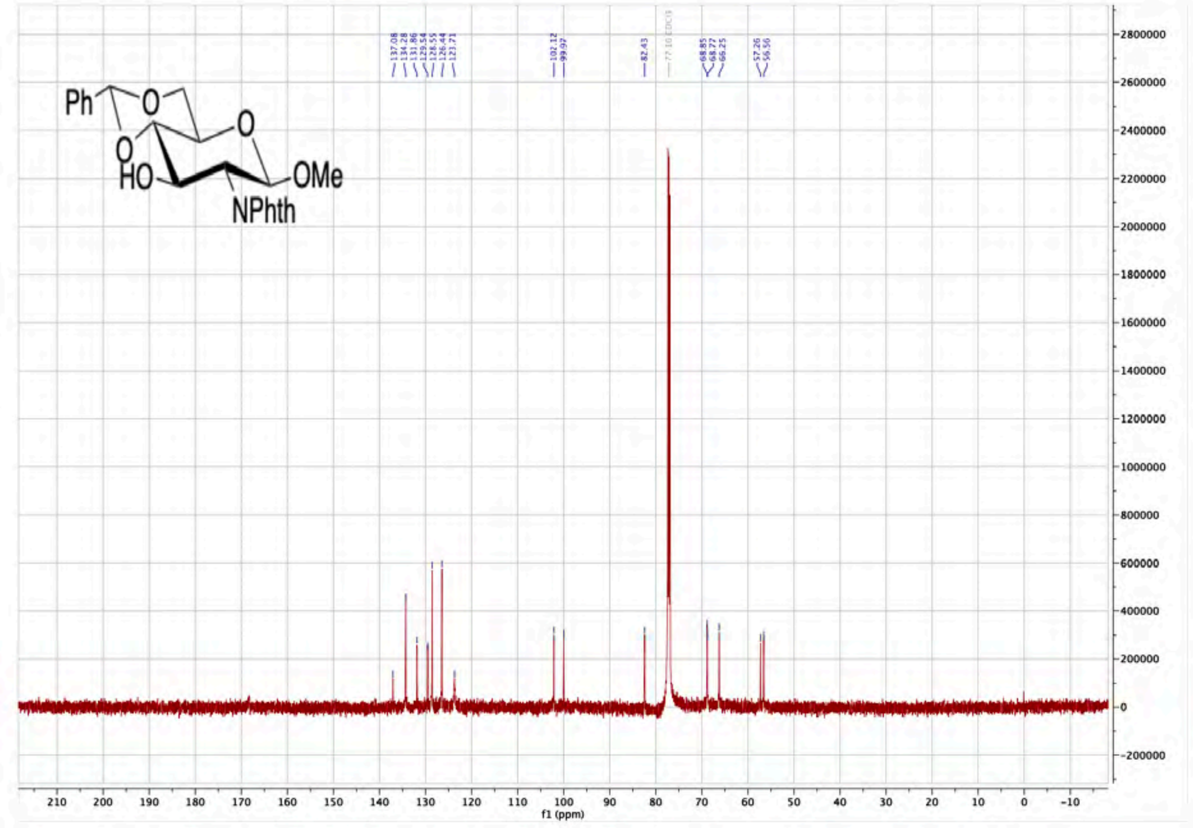
Ph
O
O
O
HO
OMe
NPhth
f1 (ppm)

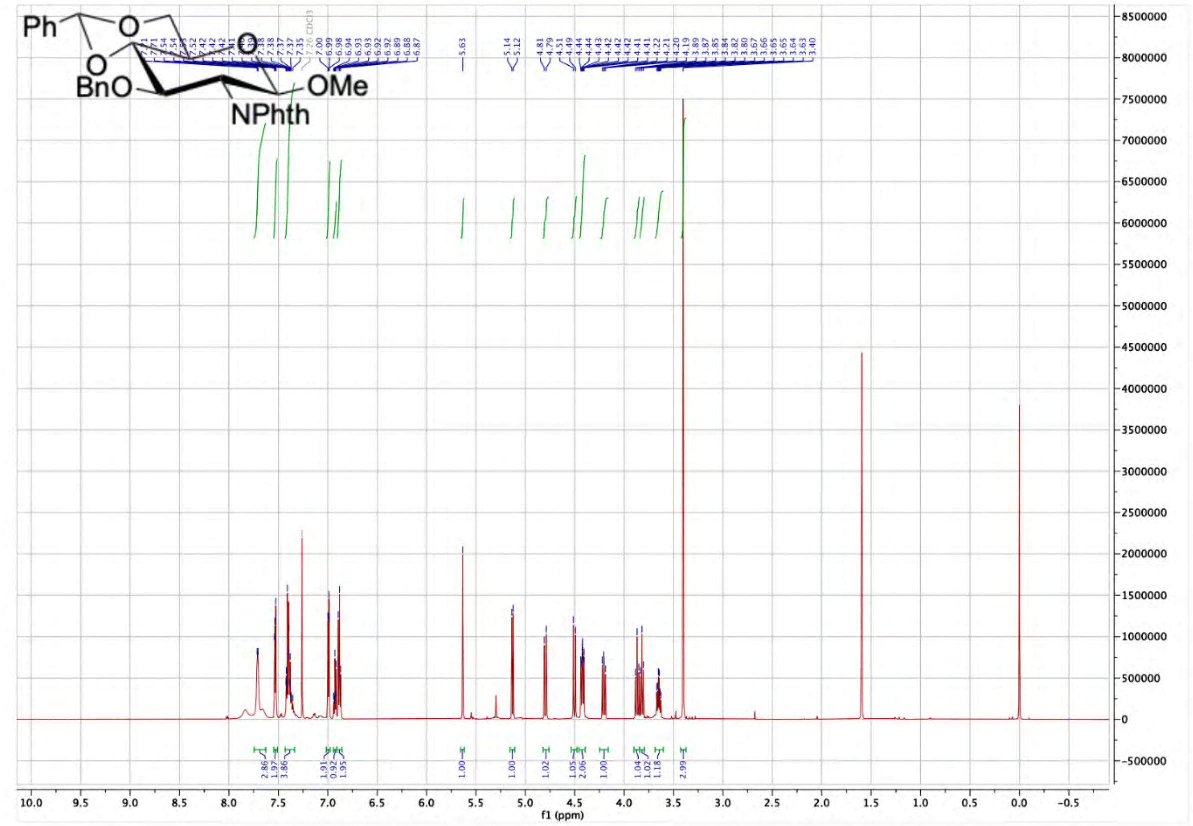
Ph
O
O
O
BnO
OMe
NPhth
f1 (ppm)

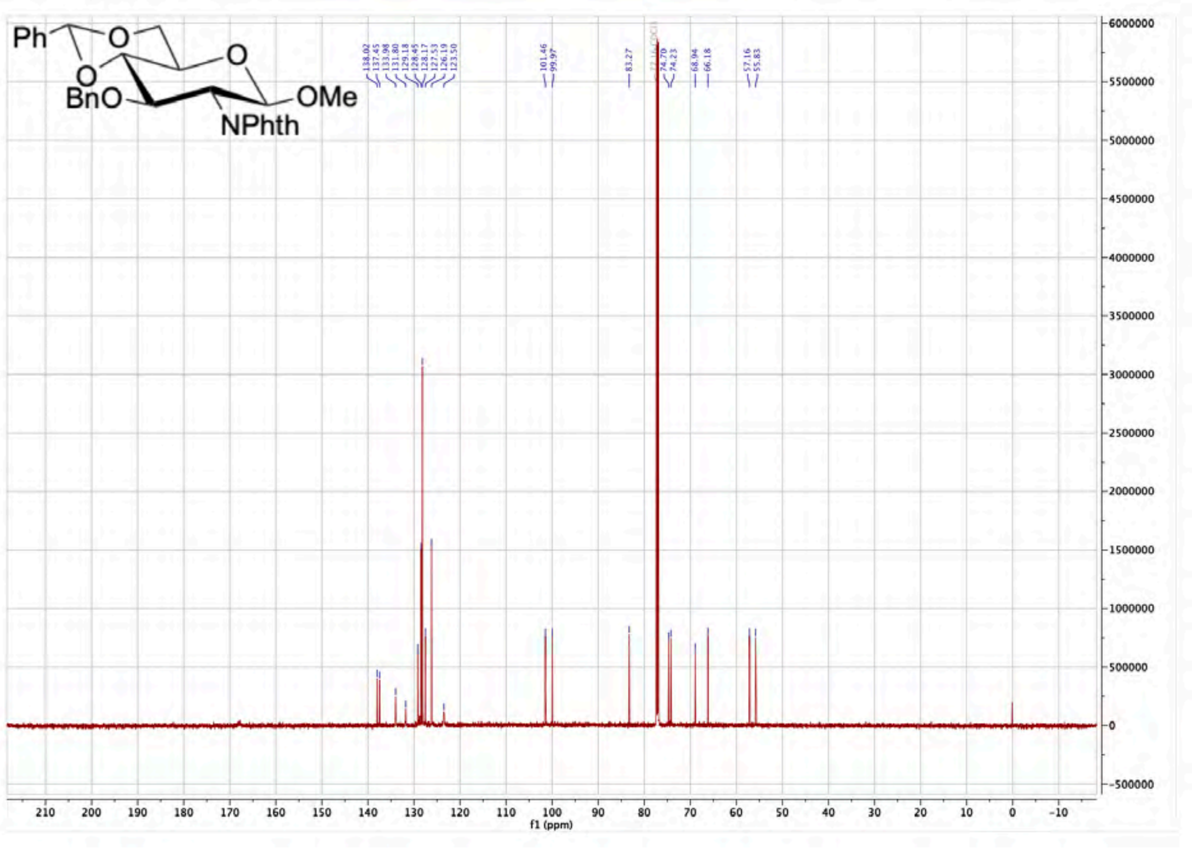

Ph
O
O
O
BnO
OMe
NPhth
138.02
137.45
133.98
131.80
129.18
128.45
128.17
127.53
126.19
123.50
101.46
99.97
83.27
74.70
74.23
68.94
66.18
57.16
55.83
f1 (ppm)


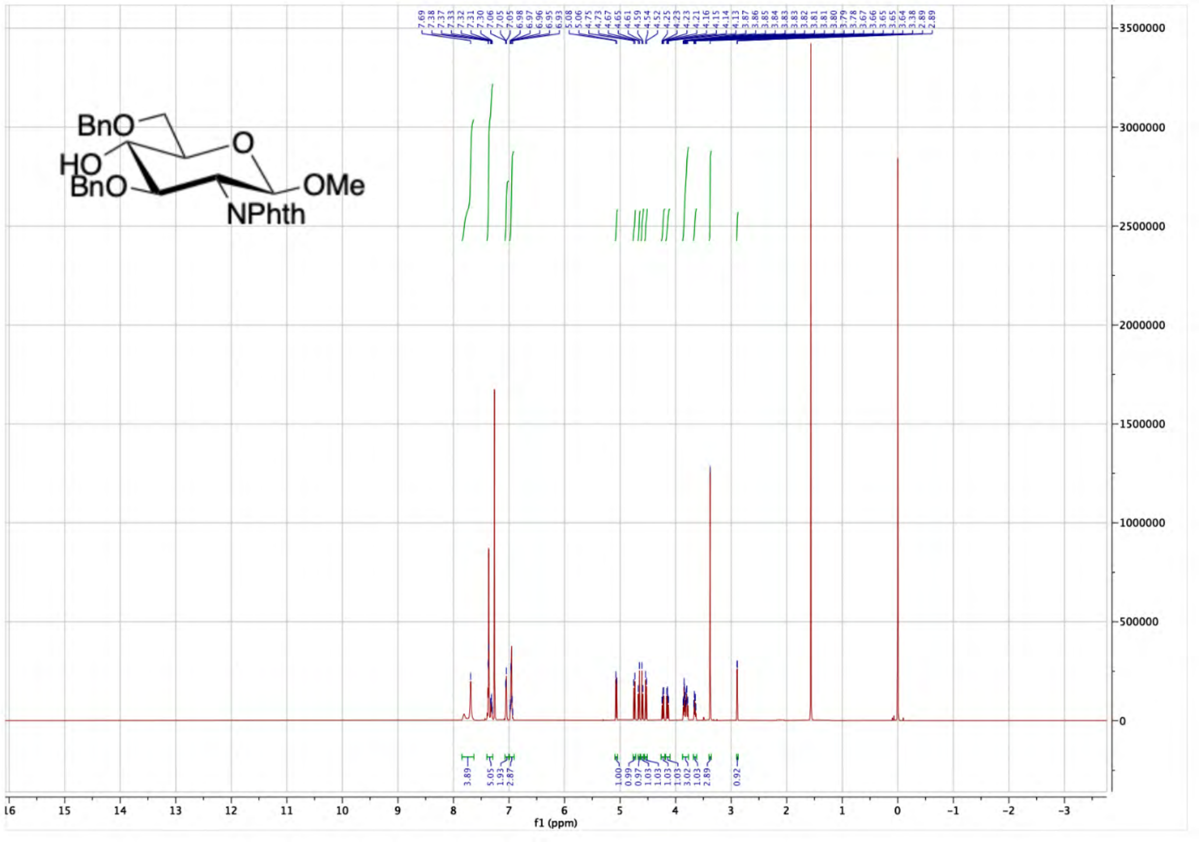

BnO
HO
BnO
O
OMe
NPhth
f1 (ppm)

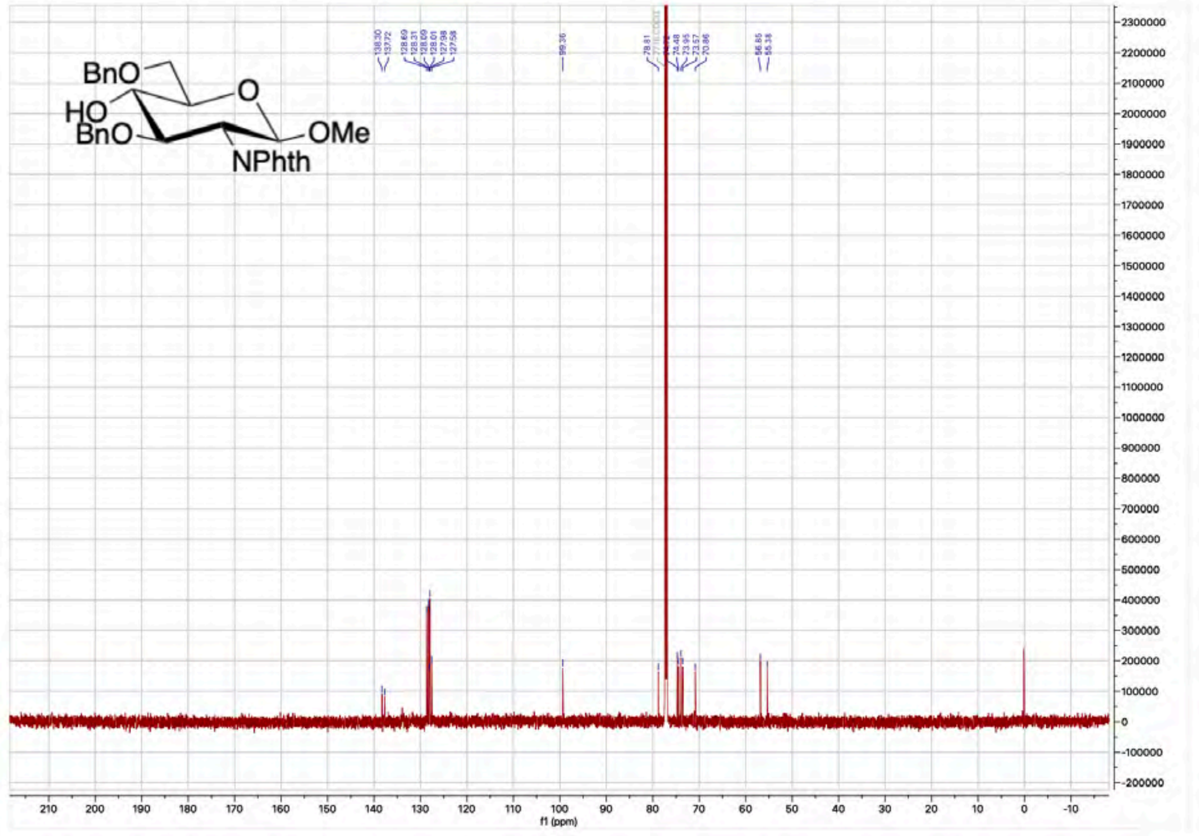

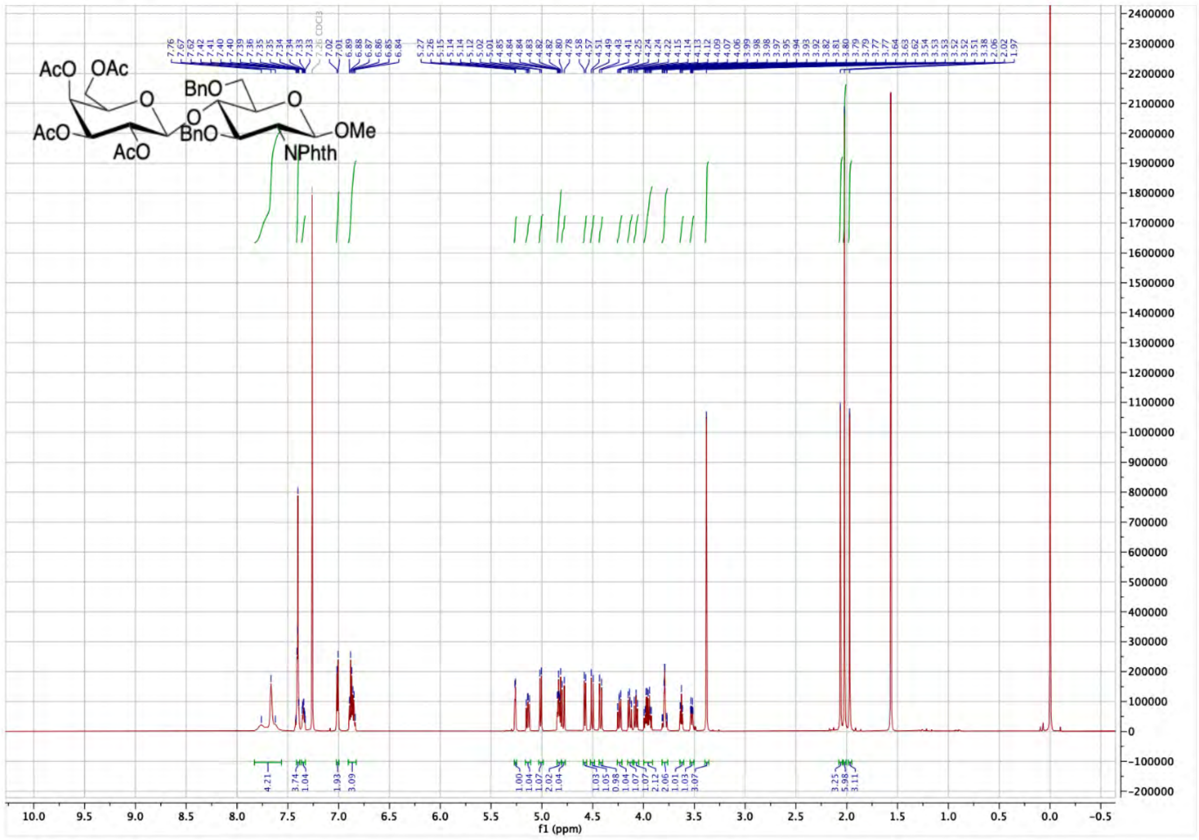

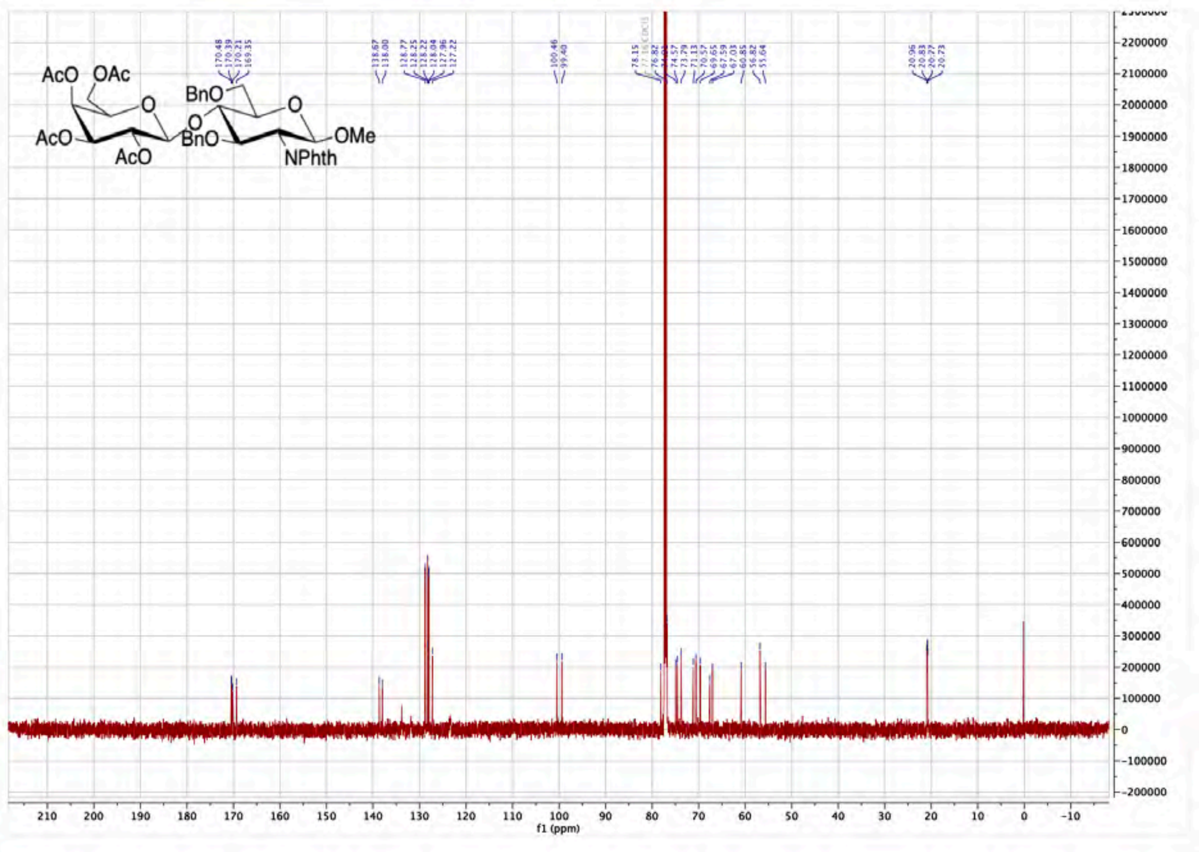
AcO OAc
BnO
O
AcO
AcO
O
BnO
O
OMe
NPhth
f1 (ppm)

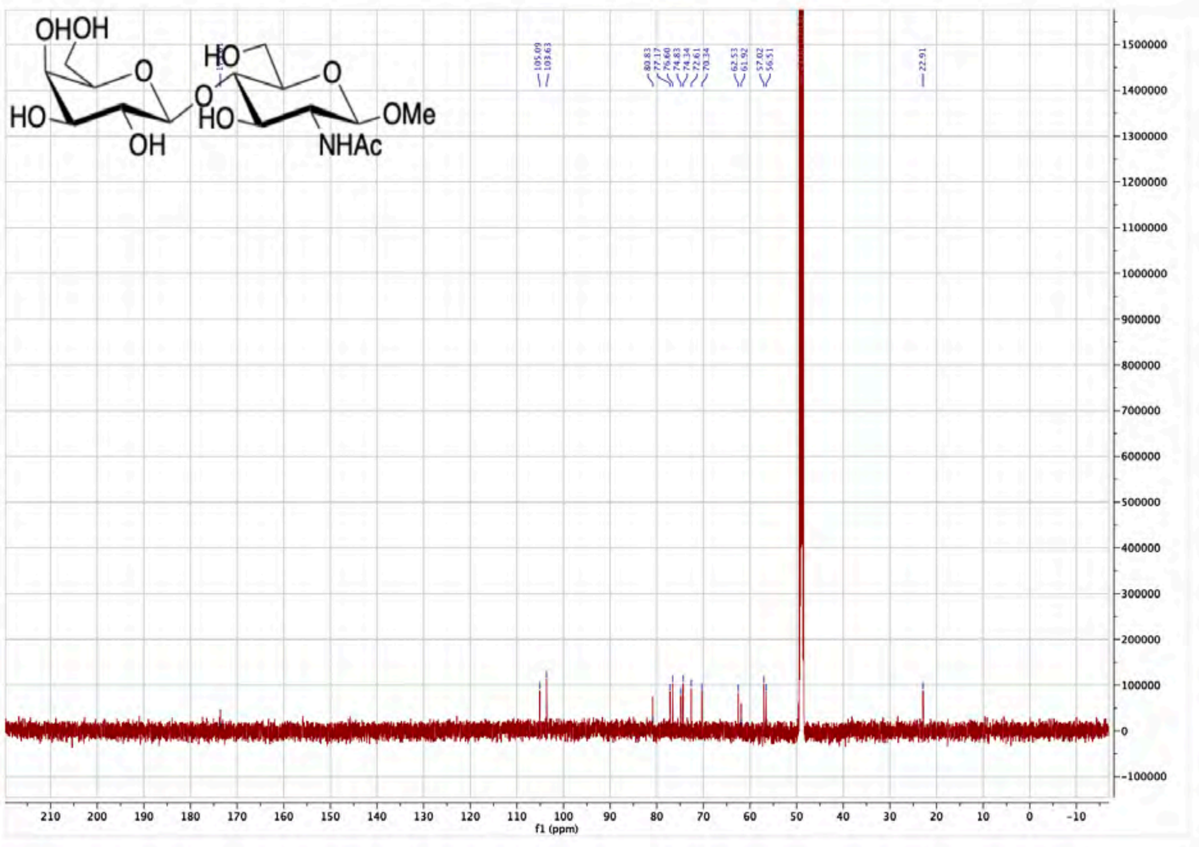
OHOH
HO
O
O
HO
OH
HO
O
OMe
NHAc
f1 (ppm)

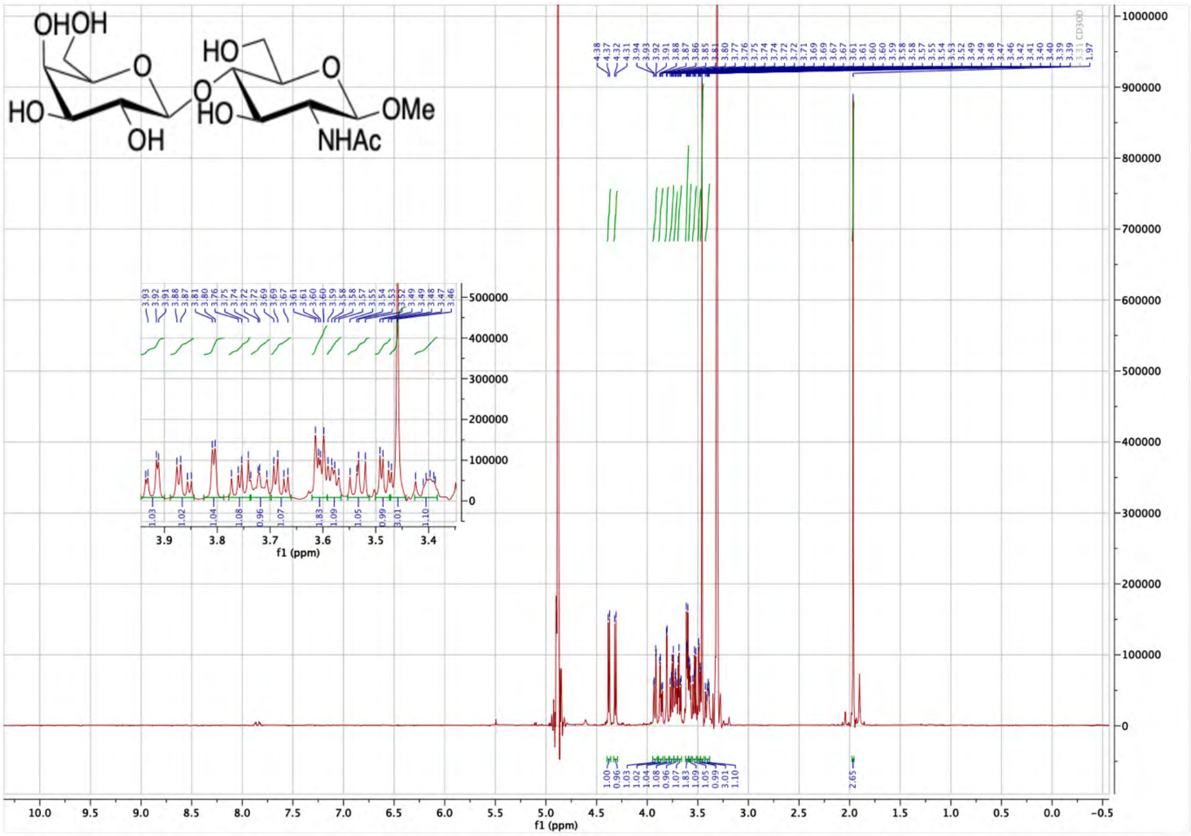

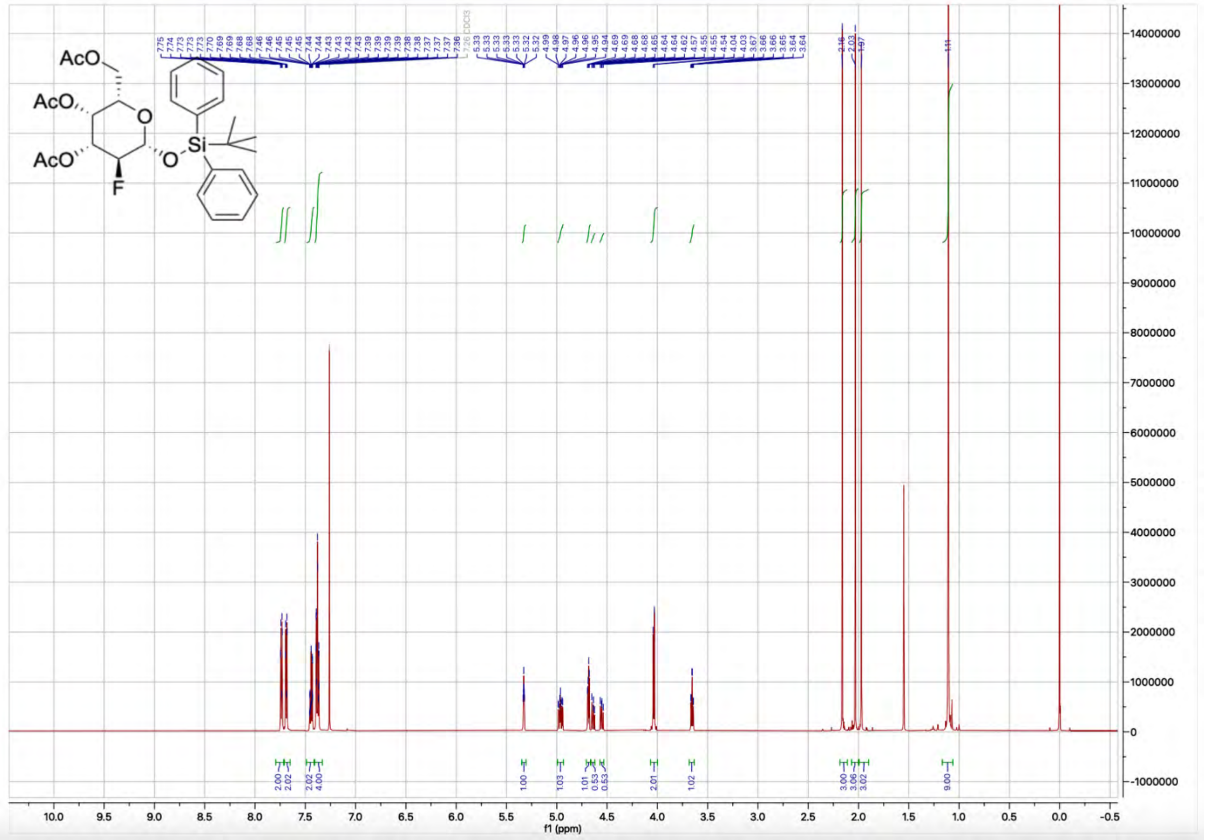

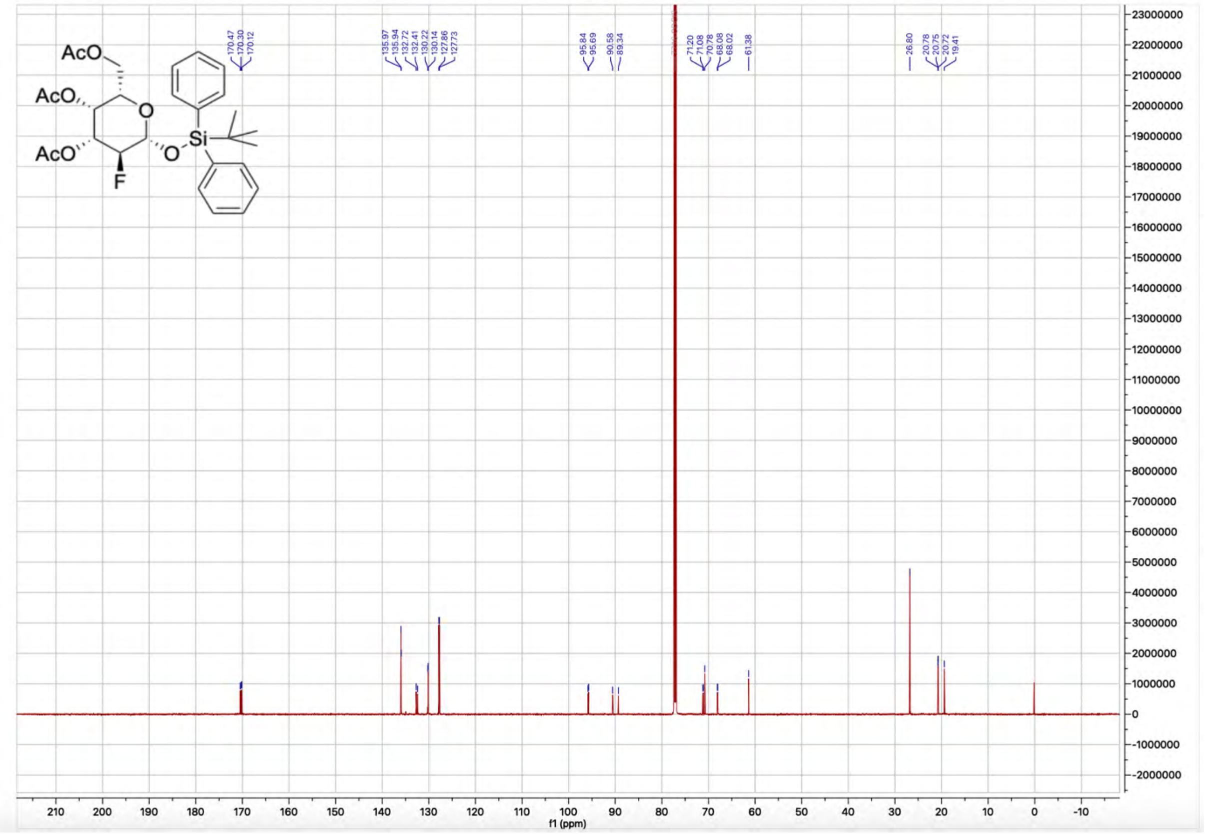
AcO
AcO
AcO
O
O
Si
F
170.47
170.30
170.12
135.97
135.94
132.72
132.41
130.22
130.14
127.86
127.73
95.84
95.69
90.58
89.34
71.20
71.08
70.78
68.08
68.02
61.38
26.80
20.78
20.75
20.72
19.41
f1 (ppm)

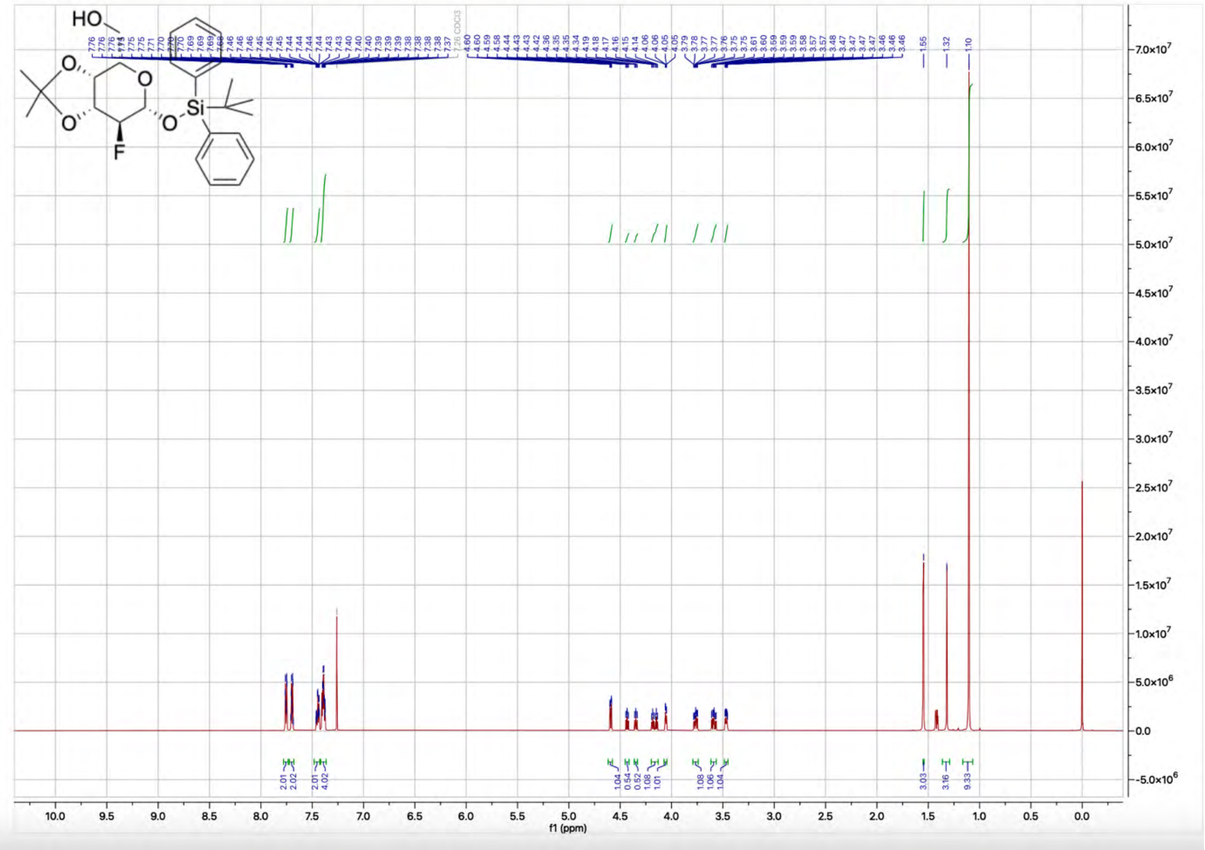
HO
O
O
O
O
Si
F
1.55
1.32
1.10
f1 (ppm)

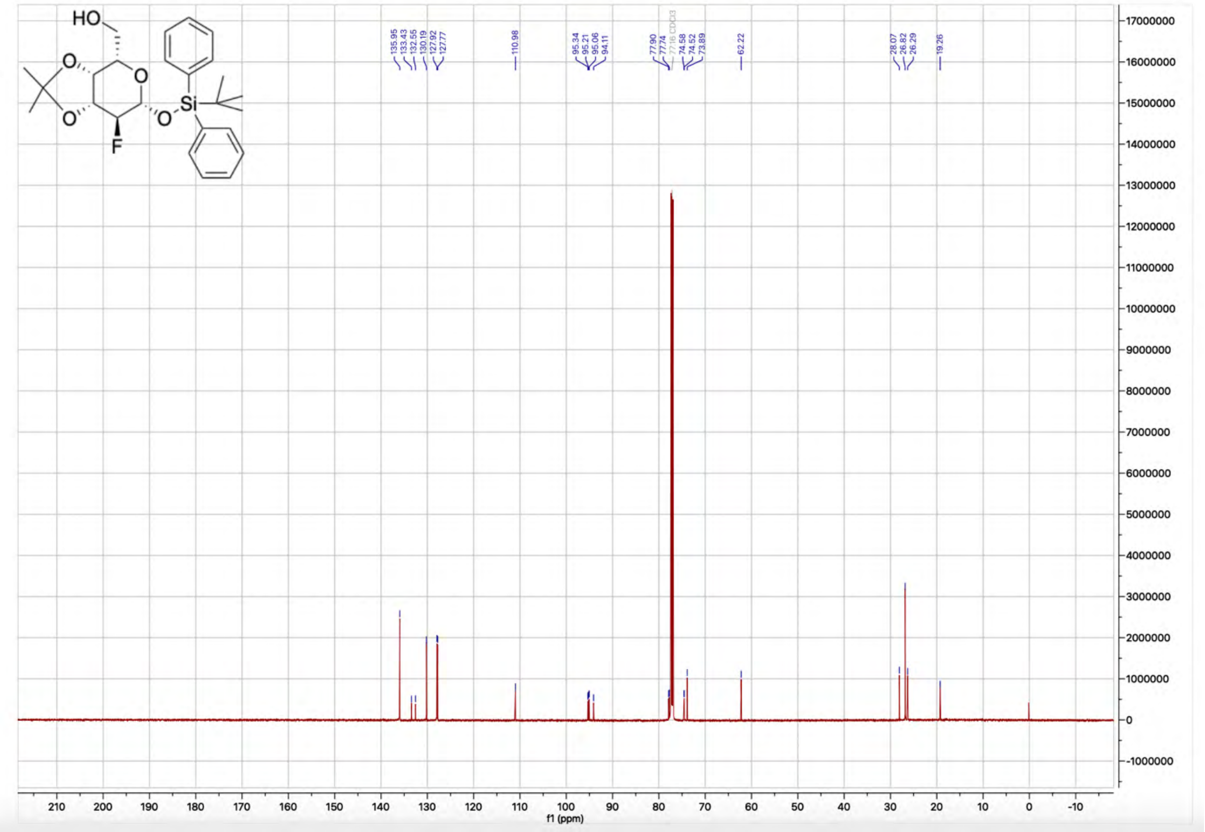
HO
O
O
O
F
O
Si
135.95
133.43
132.55
130.19
127.92
127.77
110.98
95.34
95.21
95.06
94.11
77.90
77.74
77.16 CDCl3
74.58
74.52
73.89
62.22
28.07
26.82
26.29
19.26
f1 (ppm)

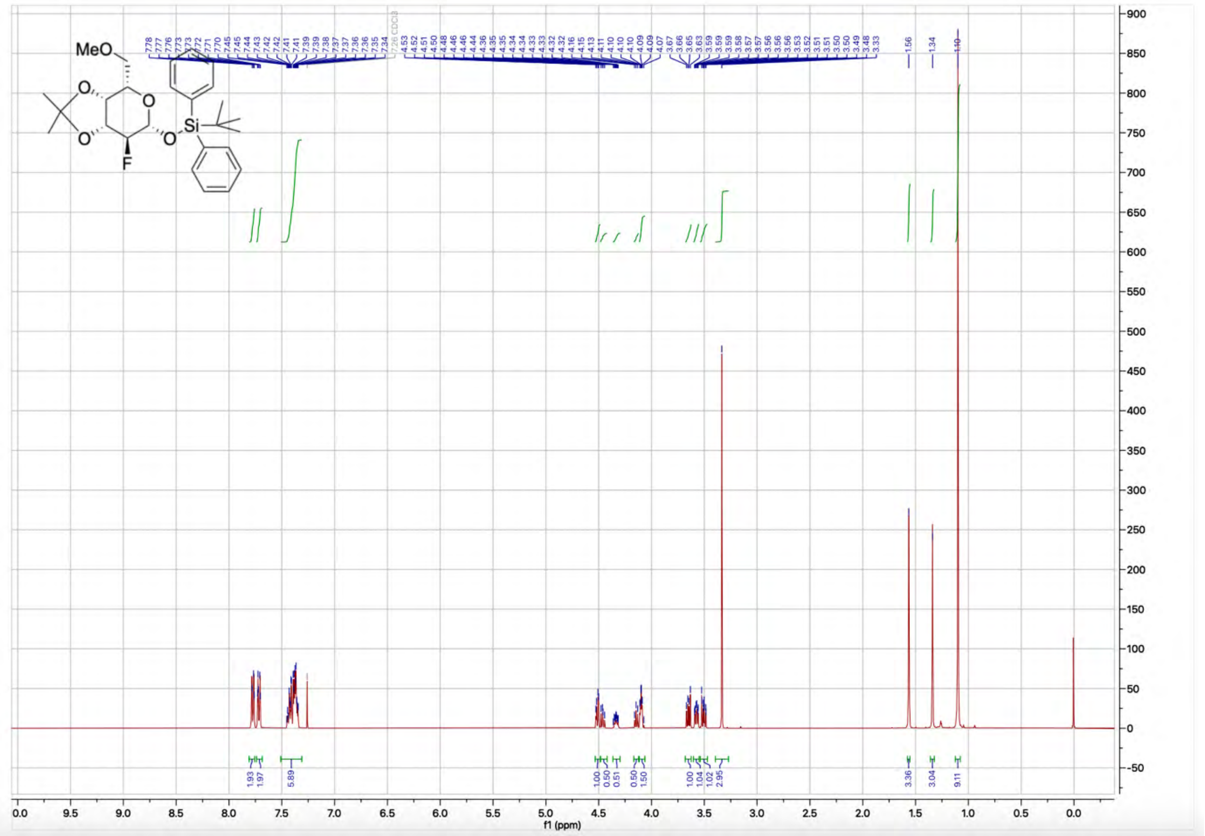
MeO
O
O
O
F
O
Si
1.56
1.34
1.10
1.93
1.97
5.89
1.00
0.50
0.51
0.50
1.50
1.00
1.04
1.02
2.95
3.36
3.04
9.11
f1 (ppm)

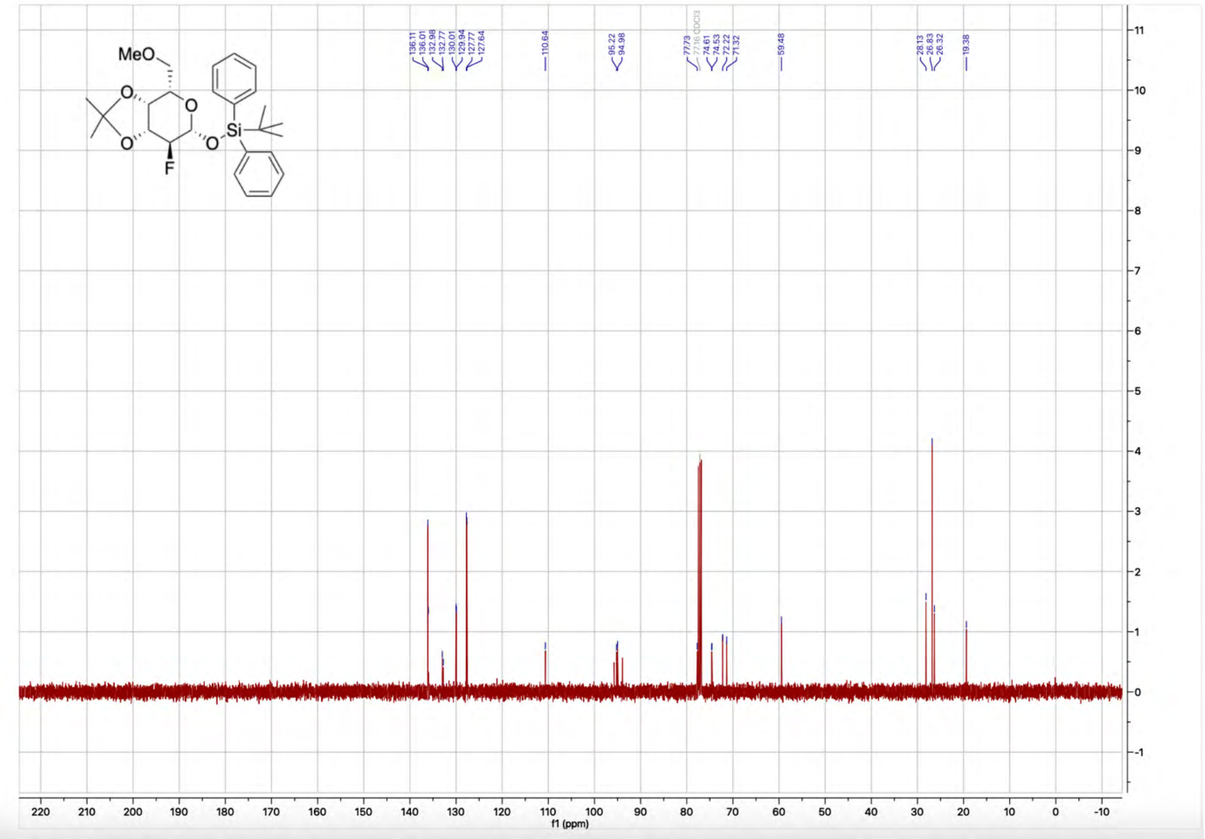
MeO
O
O
O
F
O
Si
136.11
136.01
132.98
132.77
130.01
129.94
127.77
127.64
110.64
95.22
94.98
77.73
74.61
74.53
72.22
71.32
59.48
28.13
26.83
26.32
19.38
f1 (ppm)

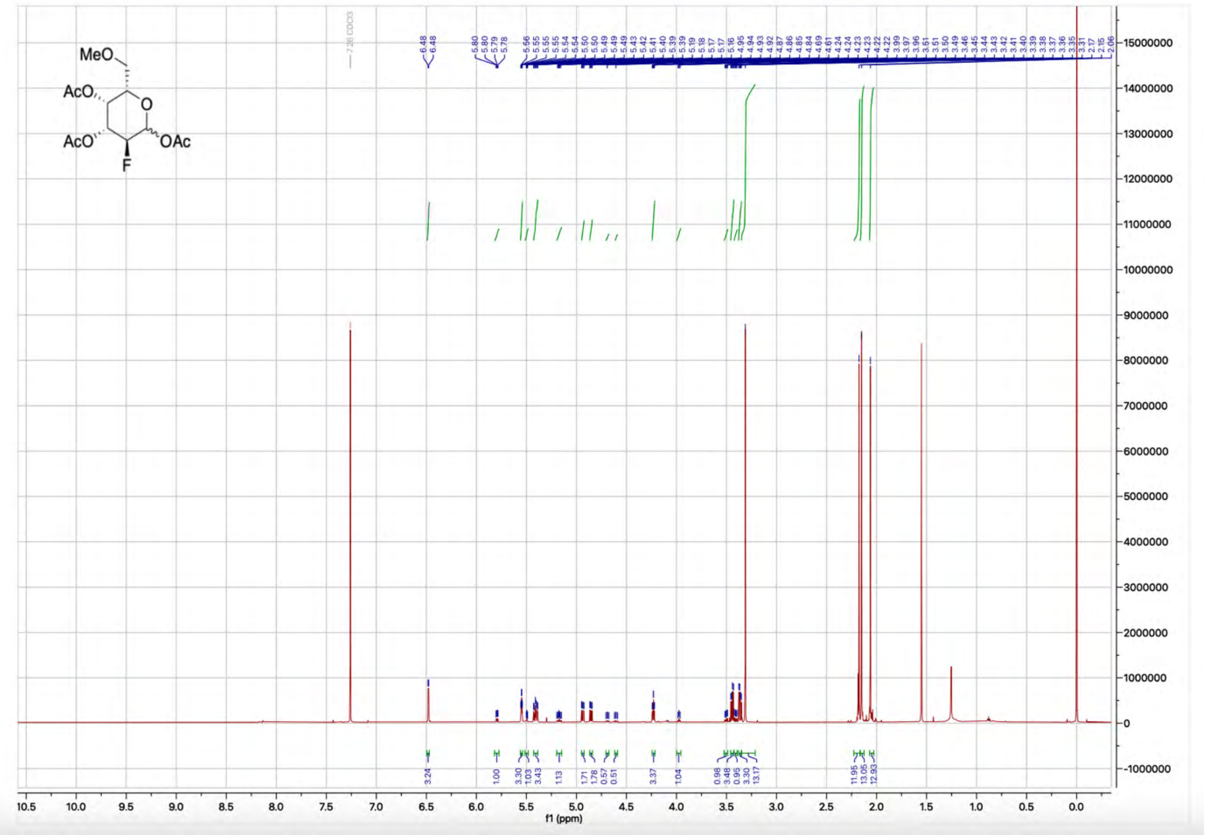
MeO
AcO
AcO
F
O
OAc
7.26 CDCl3
6.48
6.48
5.80
5.80
5.79
5.78
3.24
1.00
3.30
1.03
3.43
1.13
1.71
1.78
0.57
0.51
3.37
1.04
0.98
3.48
0.95
3.30
13.17
11.95
13.05
12.93
f1 (ppm)

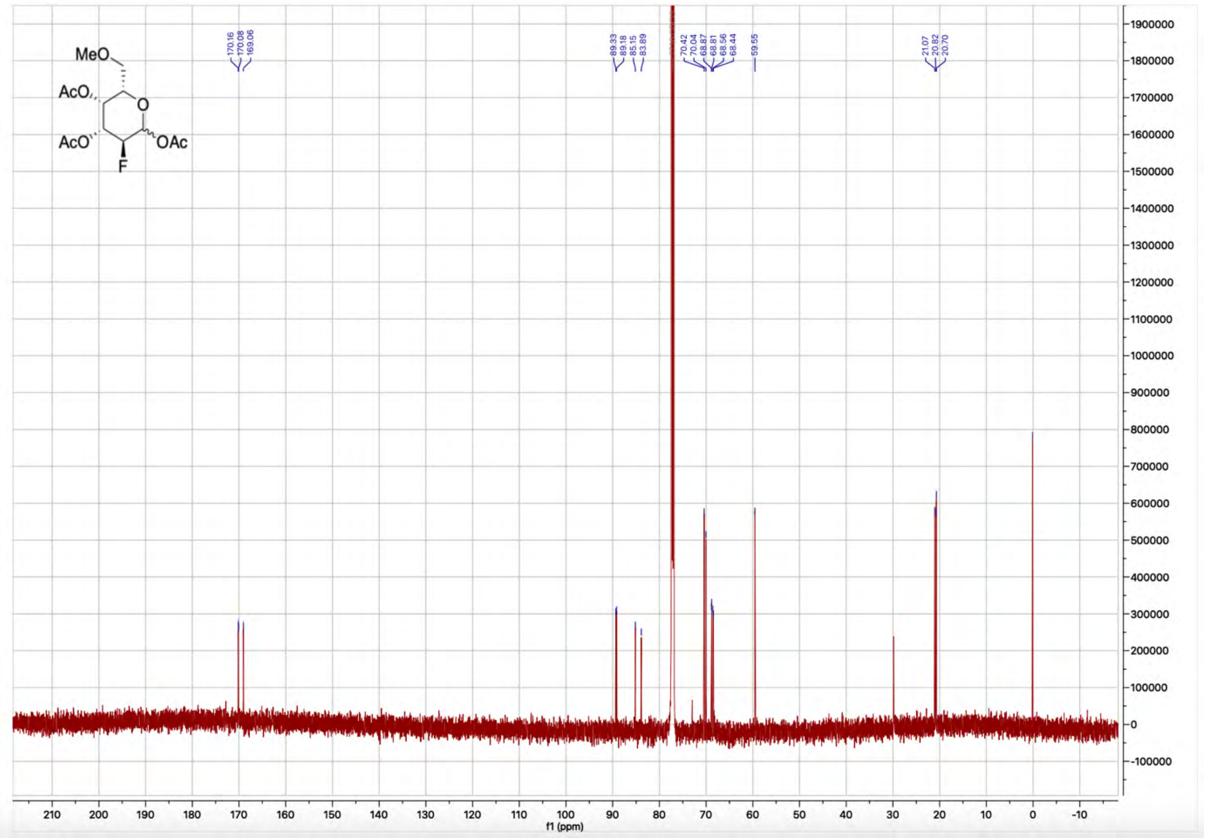

MeO
AcO
O
AcO
F
OAc
170.16
170.08
169.06
89.33
89.18
85.15
83.89
70.42
70.04
68.87
68.81
68.56
68.44
59.55
21.07
20.82
20.70
f1 (ppm)


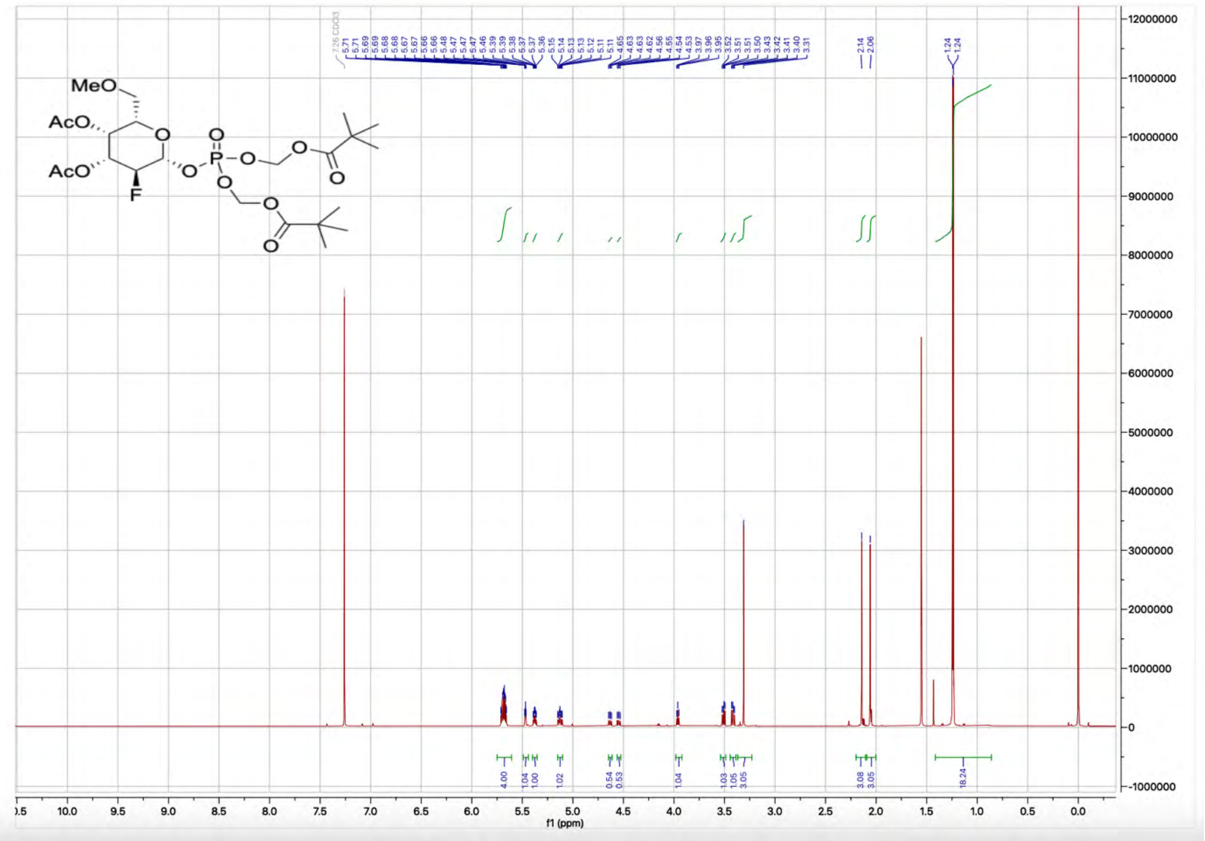

MeO
AcO
O
AcO
F
O
P
O
O
O
O
O
O
O
O
7.26 CDCl3
5.71
5.69
5.68
5.68
5.67
5.67
5.66
5.66
5.48
5.47
5.47
5.47
5.46
5.39
5.39
5.38
5.37
5.37
5.36
5.15
5.14
5.13
5.13
5.12
5.11
5.11
4.65
4.63
4.63
4.62
4.56
4.55
4.54
4.53
3.97
3.96
3.95
3.52
3.51
3.51
3.50
3.43
3.42
3.41
3.40
3.31
2.14
2.06
1.24
1.24
4.00
1.04
1.00
1.02
0.54
0.53
1.04
1.03
1.05
3.05
3.08
3.05
18.24
f1 (ppm)

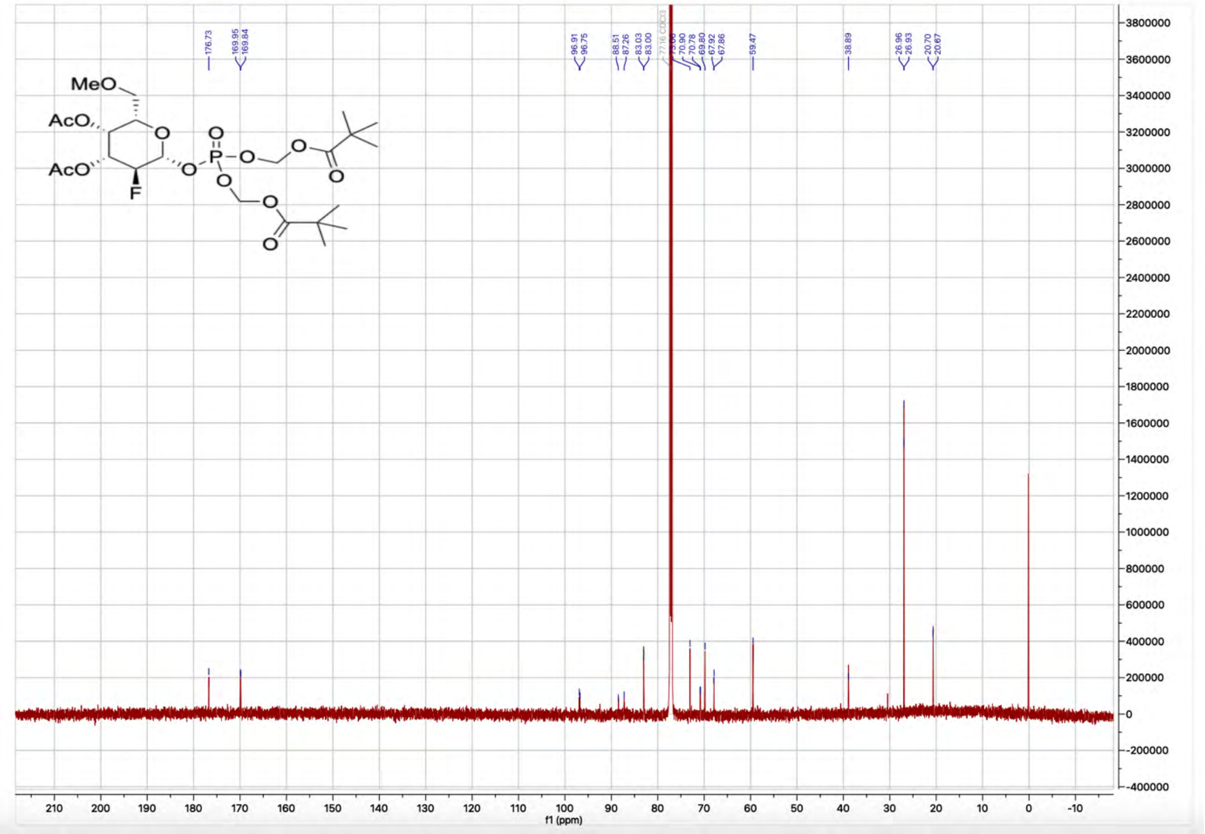


#### 5) Graphs depicting the determination of Ki

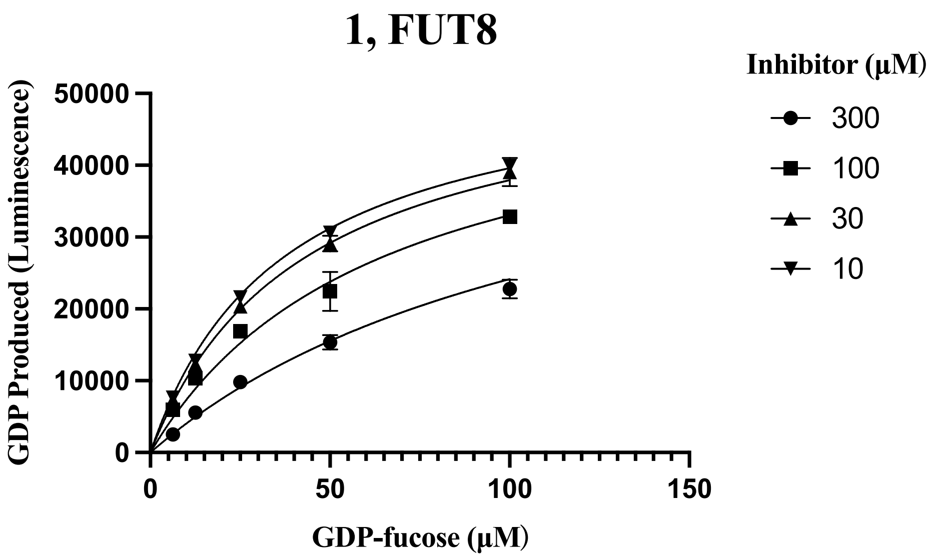

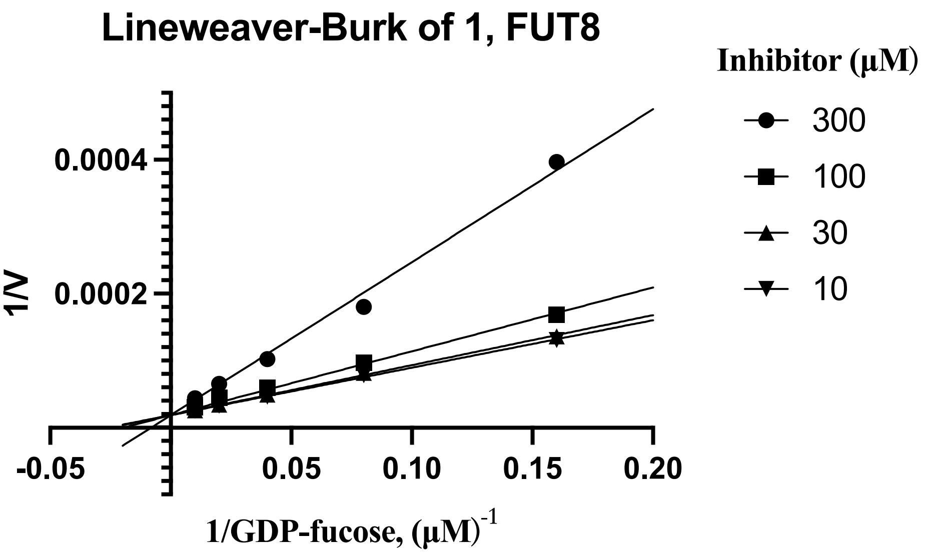
Lineweaver-Burk of 1, FUT8
Inhibitor (μM)
300
100
30
10
1/V
0.0004
0.0002
-0.05
0.05
0.10
0.15
0.20
1/GDP-fucose, (μM)-1

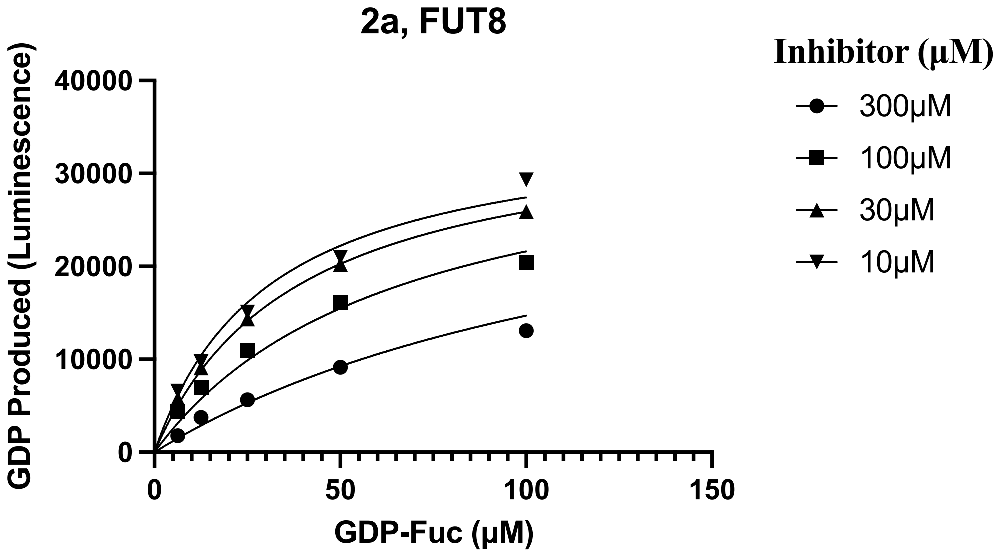
2a, FUT8
Inhibitor (μM)
300μM
100μM
30μM
10μM
GDP Produced (Luminescence)
40000
30000
20000
10000
0
0
50
100
150
GDP-Fuc (μM)

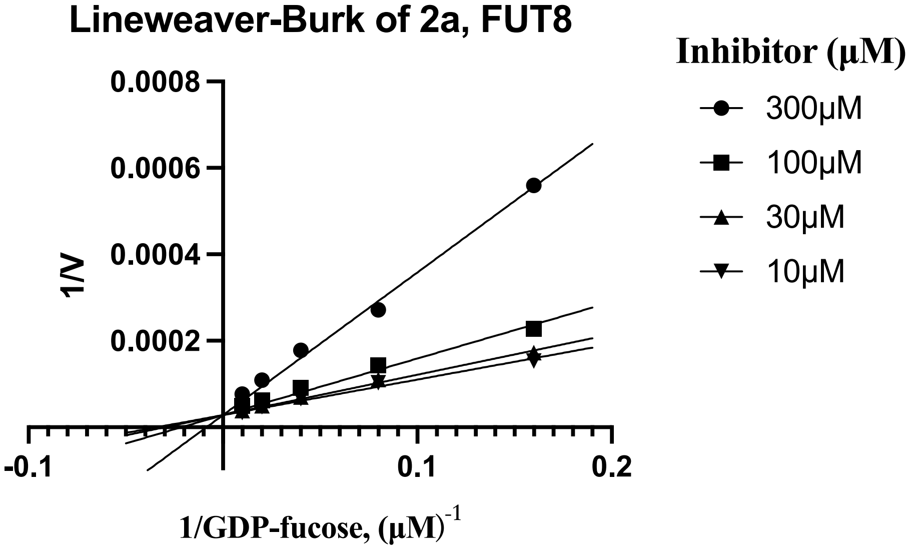
Lineweaver-Burk of 2a, FUT8
Inhibitor (μM)
300μM
100μM
30μM
10μM
1/V
0.0008
0.0006
0.0004
0.0002
-0.1
0.1
0.2
1/GDP-fucose, (μM)-1

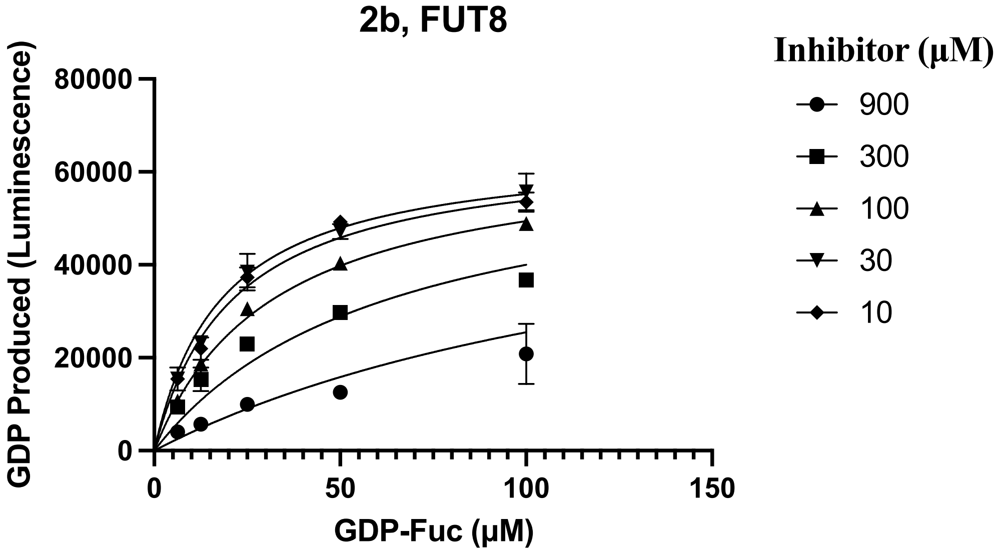
2b, FUT8
GDP Produced (Luminescence)
80000
60000
40000
20000
0
0
50
100
150
GDP-Fuc (μM)
Inhibitor (μM)
900
300
100
30
10

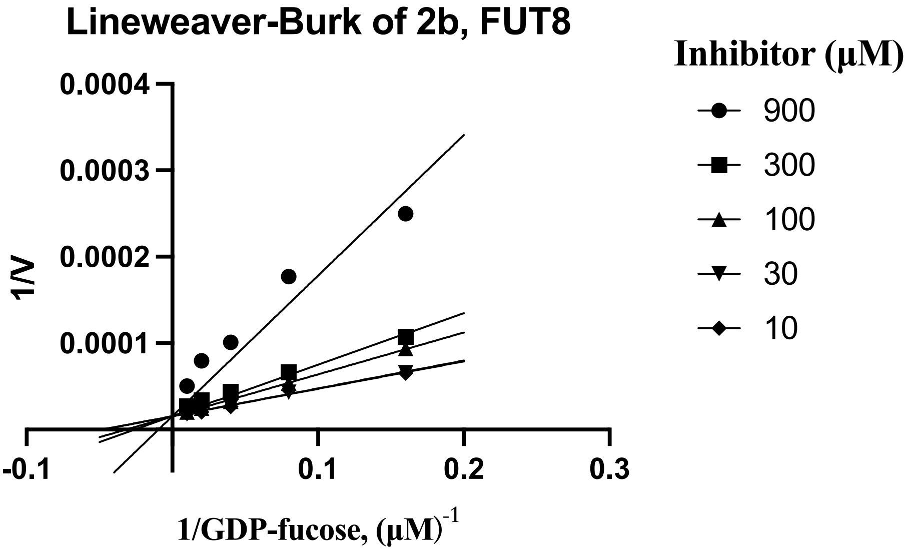
Lineweaver-Burk of 2b, FUT8
1/V
0.0004
0.0003
0.0002
0.0001
-0.1
0.1
0.2
0.3
1/GDP-fucose, (μM)-1
Inhibitor (μM)
900
300
100
30
10

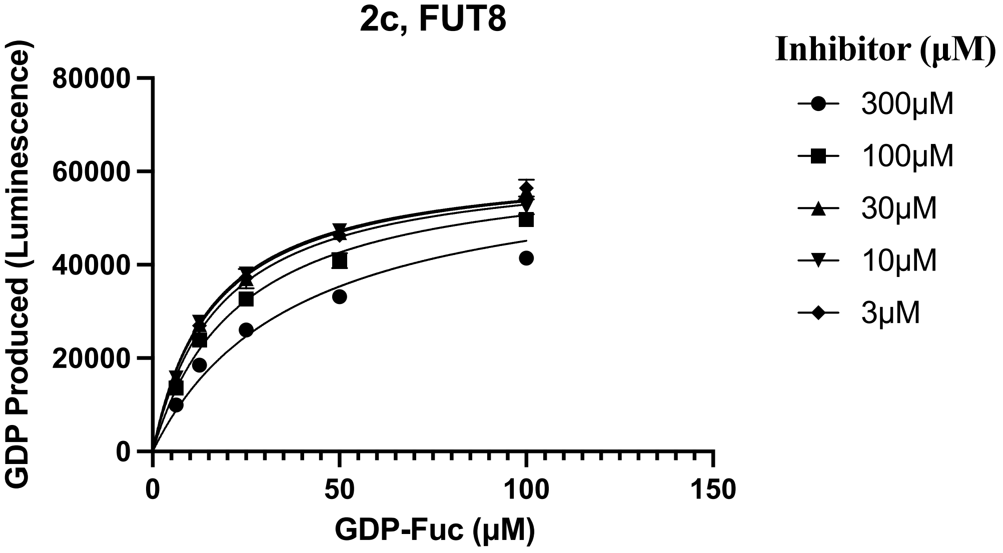
2c, FUT8
GDP Produced (Luminescence)
80000
60000
40000
20000
0
0
50
100
150
GDP-Fuc (μM)
Inhibitor (μM)
300μM
100μM
30μM
10μM
3μM

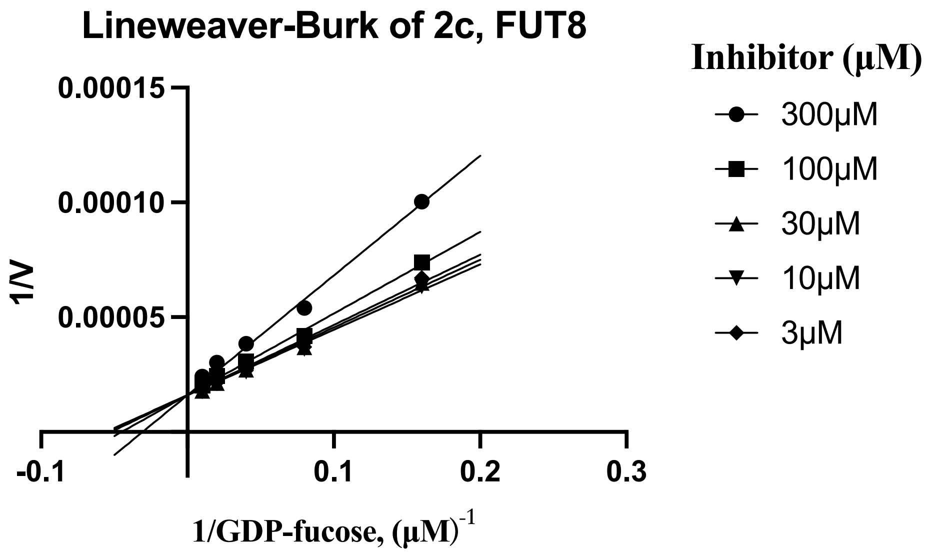
Lineweaver-Burk of 2c, FUT8
0.00015
0.00010
0.00005
1/V
-0.1
0.1
0.2
0.3
1/GDP-fucose, (μM)-1
Inhibitor (μM)
300μM
100μM
30μM
10μM
3μM

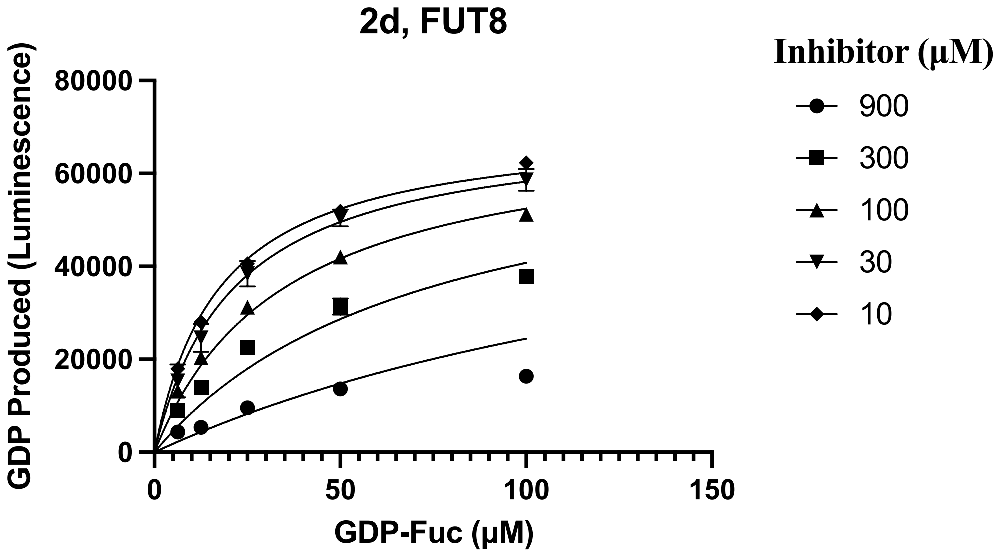
2d, FUT8
80000
60000
40000
20000
0
GDP Produced (Luminescence)
0
50
100
150
GDP-Fuc (μM)
Inhibitor (μM)
900
300
100
30
10

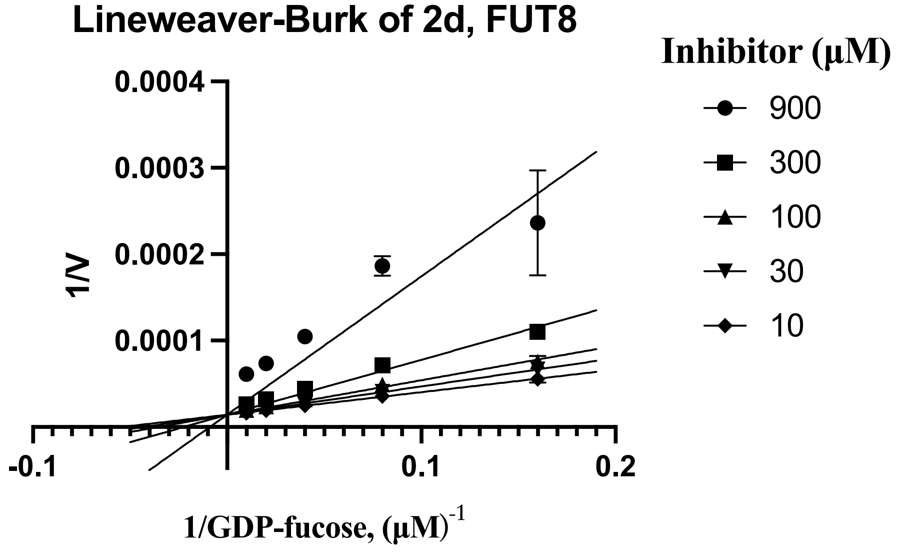
Lineweaver-Burk of 2d, FUT8
0.0004
0.0003
0.0002
0.0001
1/V
-0.1
0.1
0.2
1/GDP-fucose, (μM)-1
Inhibitor (μM)
900
300
100
30
10

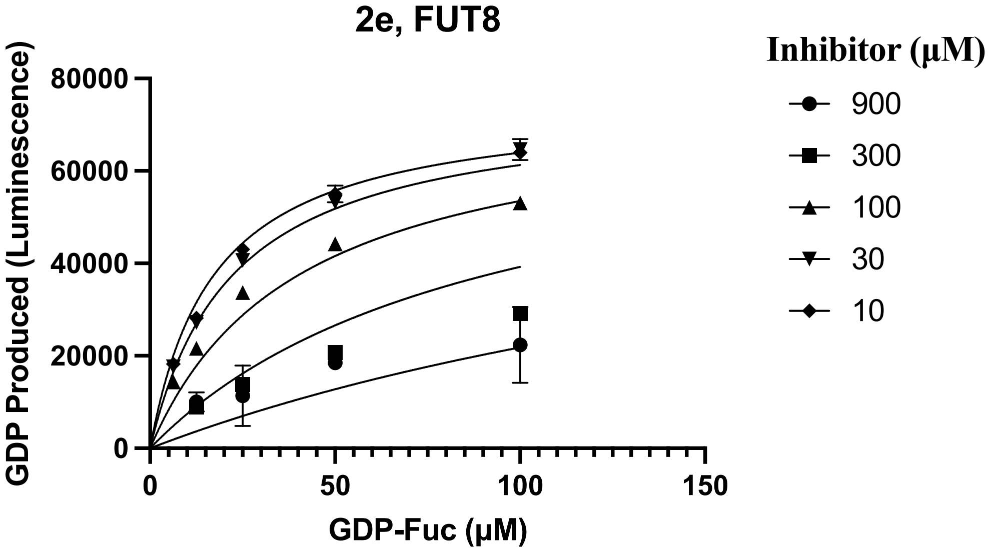
2e, FUT8
GDP Produced (Luminescence)
80000
60000
40000
20000
0
0
50
100
150
GDP-Fuc (μM)
Inhibitor (μM)
900
300
100
30
10

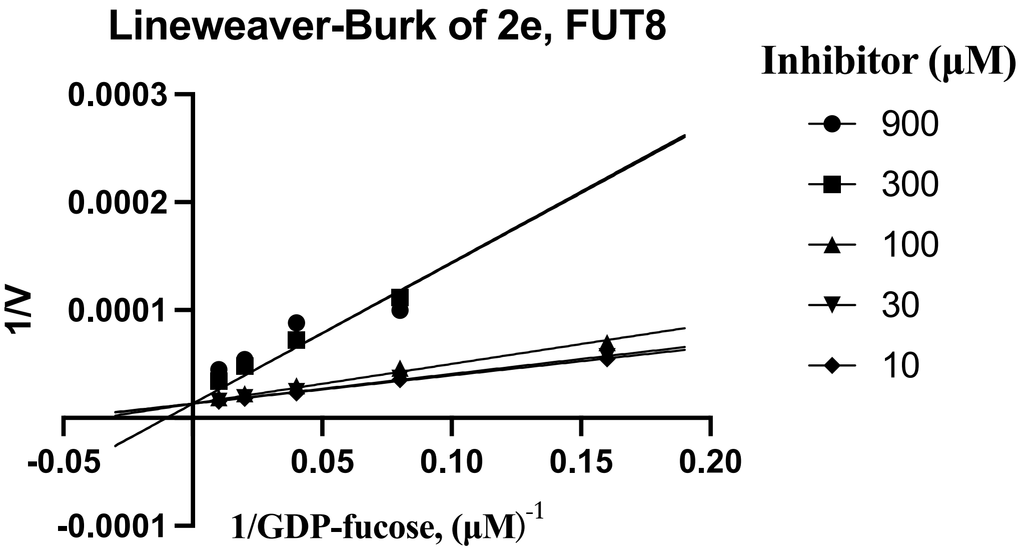
Lineweaver-Burk of 2e, FUT8
1/V
0.0003
0.0002
0.0001
-0.0001
-0.05
0.05
0.10
0.15
0.20
1/GDP-fucose, (μM)-1
Inhibitor (μM)
900
300
100
30
10

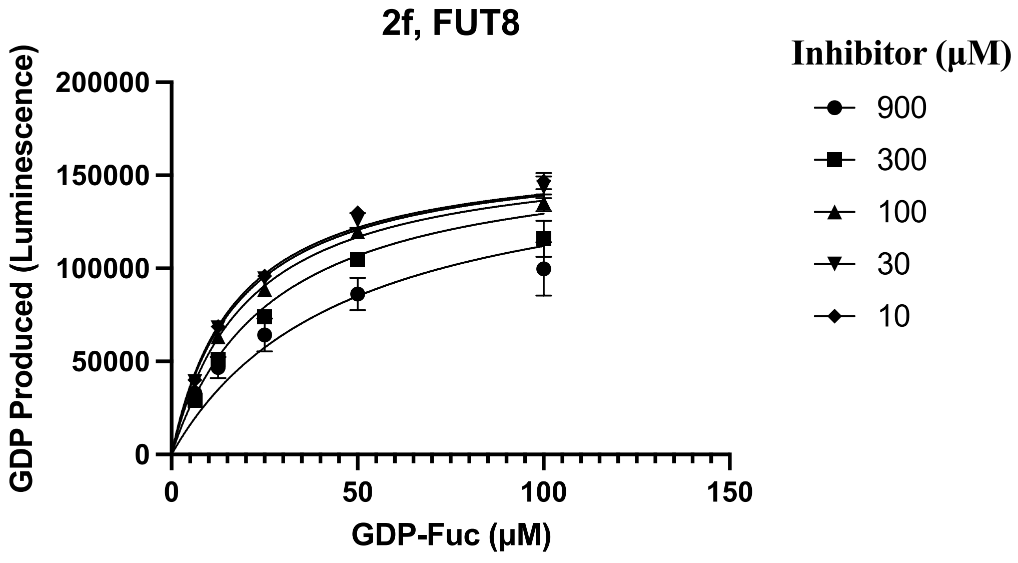
2f, FUT8
GDP Produced (Luminescence)
200000
150000
100000
50000
0
0
50
100
150
GDP-Fuc (μM)
Inhibitor (μM)
900
300
100
30
10

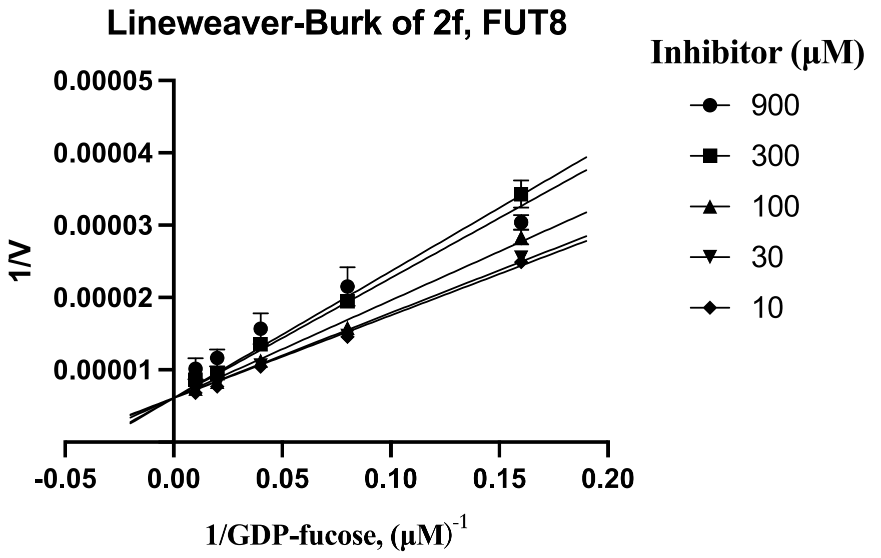

Lineweaver-Burk of 2f, FUT8
Inhibitor (μM)
900
300
100
30
10
1/V
0.00005
0.00004
0.00003
0.00002
0.00001
-0.05
0.00
0.05
0.10
0.15
0.20
1/GDP-fucose, (μM)-1


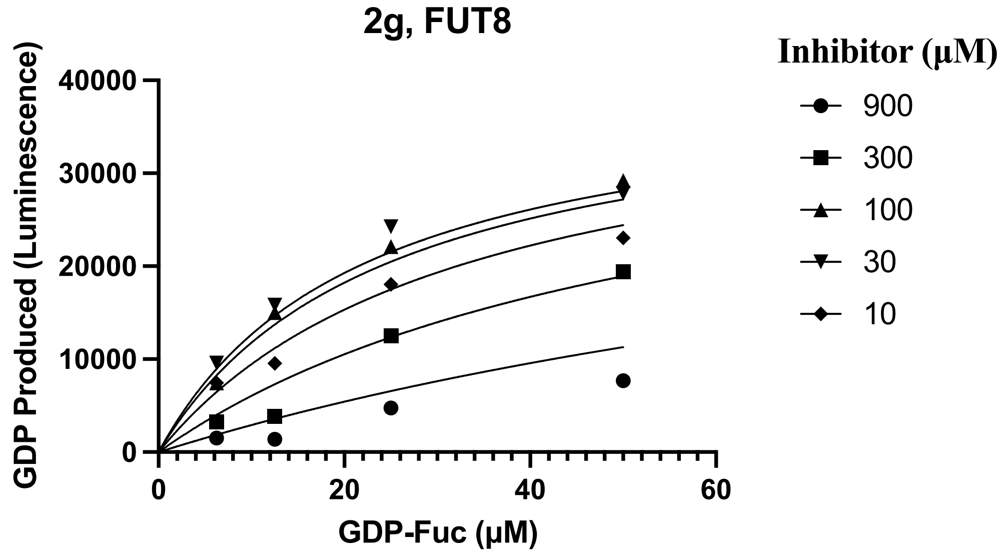

2g, FUT8
Inhibitor (μM)
900
300
100
30
10
GDP Produced (Luminescence)
40000
30000
20000
10000
0
0
20
40
60
GDP-Fuc (μM)


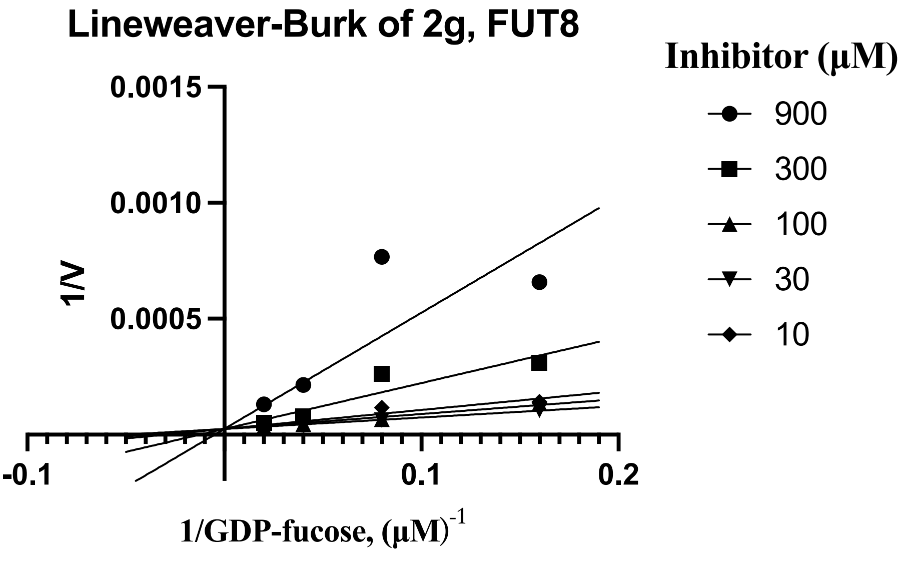

Lineweaver-Burk of 2g, FUT8
Inhibitor (μM)
900
300
100
30
10
1/V
0.0015
0.0010
0.0005
-0.1
0.1
0.2
1/GDP-fucose, (μM)-1

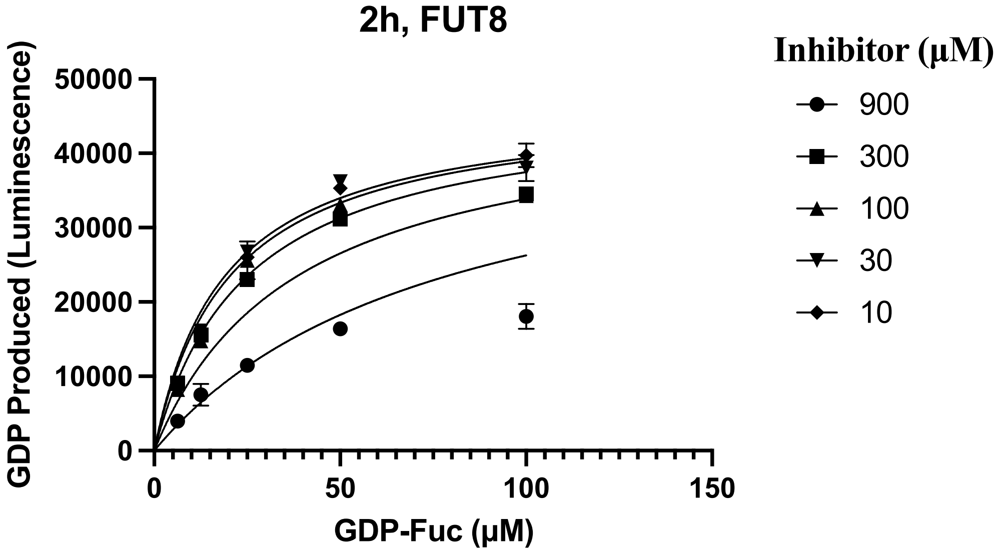
2h, FUT8
Inhibitor (μM)
900
300
100
30
10
GDP Produced (Luminescence)
50000
40000
30000
20000
10000
0
0
50
100
150
GDP-Fuc (μM)

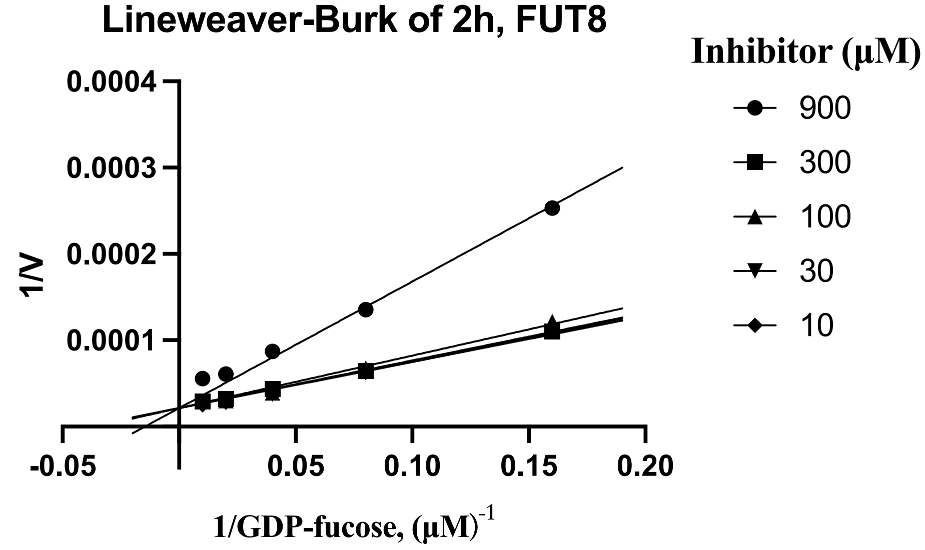
Lineweaver-Burk of 2h, FUT8
Inhibitor (μM)
900
300
100
30
10
1/V
0.0004
0.0003
0.0002
0.0001
-0.05
0.05
0.10
0.15
0.20
1/GDP-fucose, (μM)-1

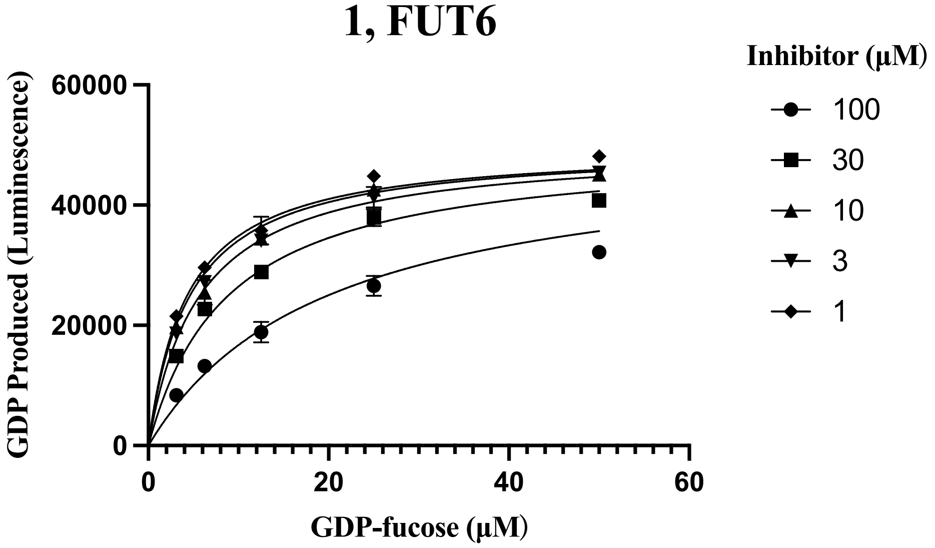
1, FUT6
Inhibitor (μM)
100
30
10
3
1
GDP Produced (Luminescence)
60000
40000
20000
0
0
20
40
60
GDP-fucose (μM)

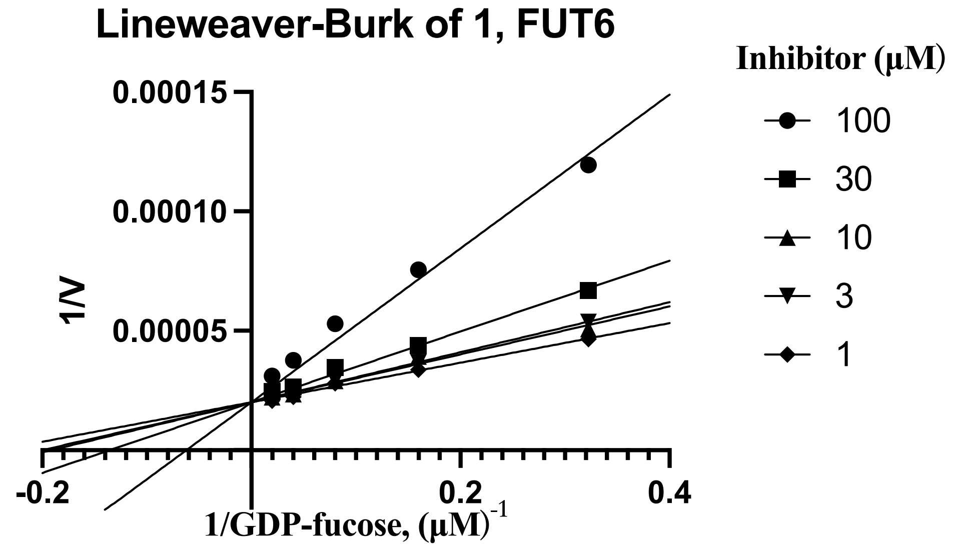
Lineweaver-Burk of 1, FUT6
Inhibitor (μM)
100
30
10
3
1
0.00015
0.00010
0.00005
1/V
-0.2
0.2
0.4
1/GDP-fucose, (μM)-1

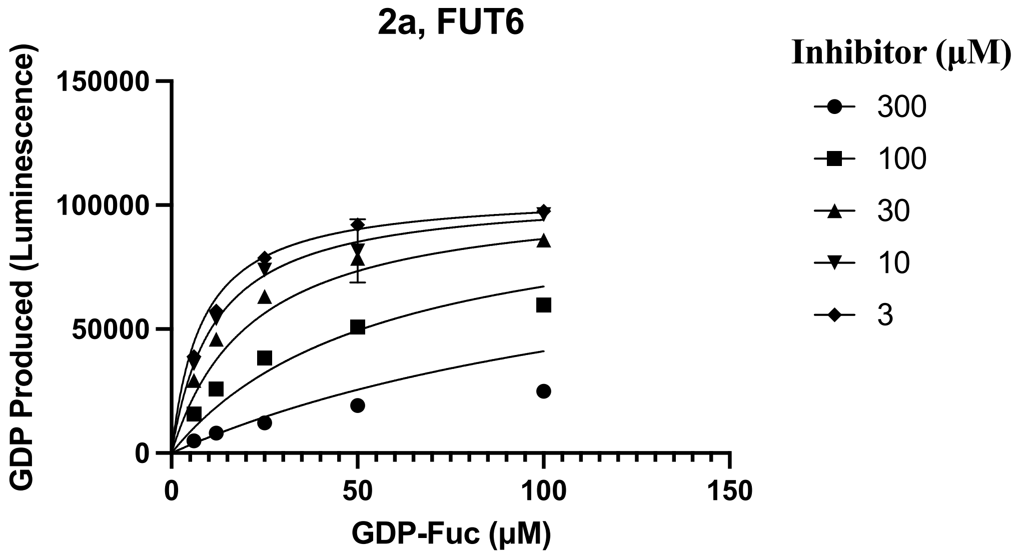
2a, FUT6
Inhibitor (μM)
300
100
30
10
3
150000
100000
50000
0
GDP Produced (Luminescence)
0
50
100
150
GDP-Fuc (μM)

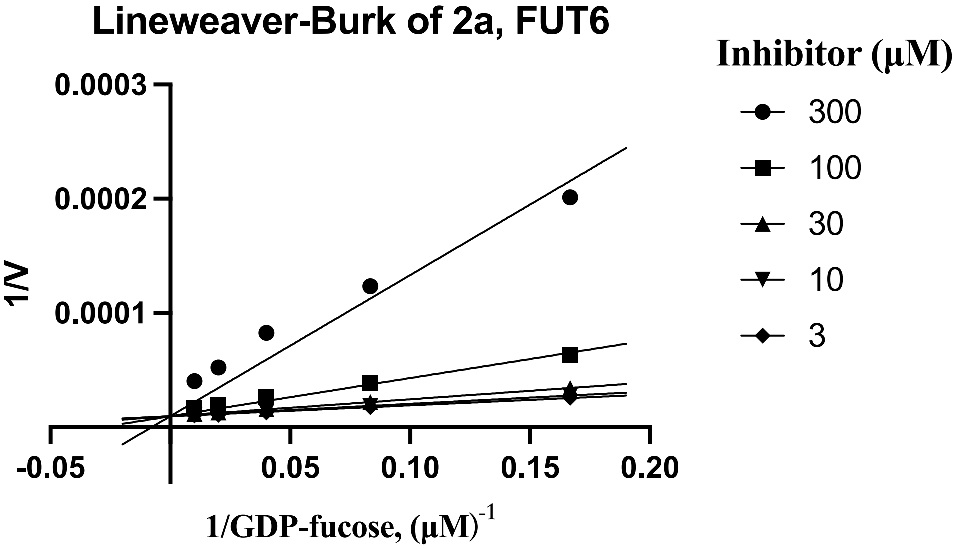
Lineweaver-Burk of 2a, FUT6
Inhibitor (μM)
300
100
30
10
3
0.0003
0.0002
0.0001
1/V
-0.05
0.05
0.10
0.15
0.20
1/GDP-fucose, (μM)-1

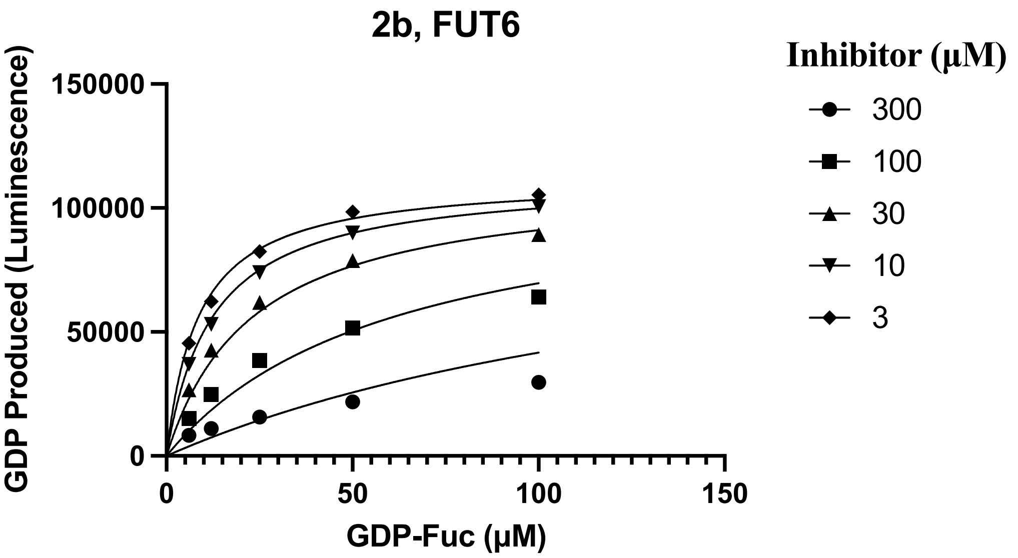
2b, FUT6
GDP Produced (Luminescence)
150000
100000
50000
0
0
50
100
150
GDP-Fuc (μM)
Inhibitor (μM)
300
100
30
10
3

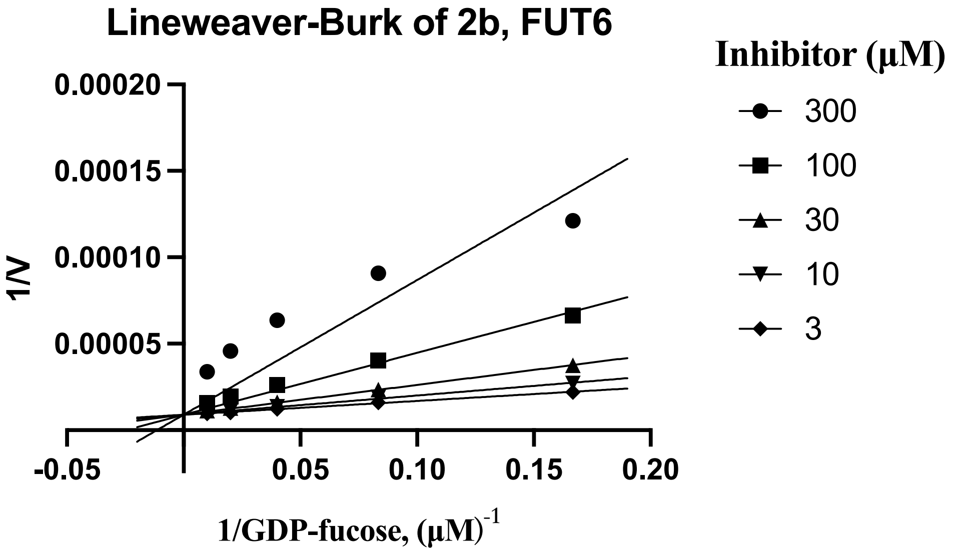
Lineweaver-Burk of 2b, FUT6
1/V
0.00020
0.00015
0.00010
0.00005
-0.05
0.05
0.10
0.15
0.20
1/GDP-fucose, (μM)-1
Inhibitor (μM)
300
100
30
10
3

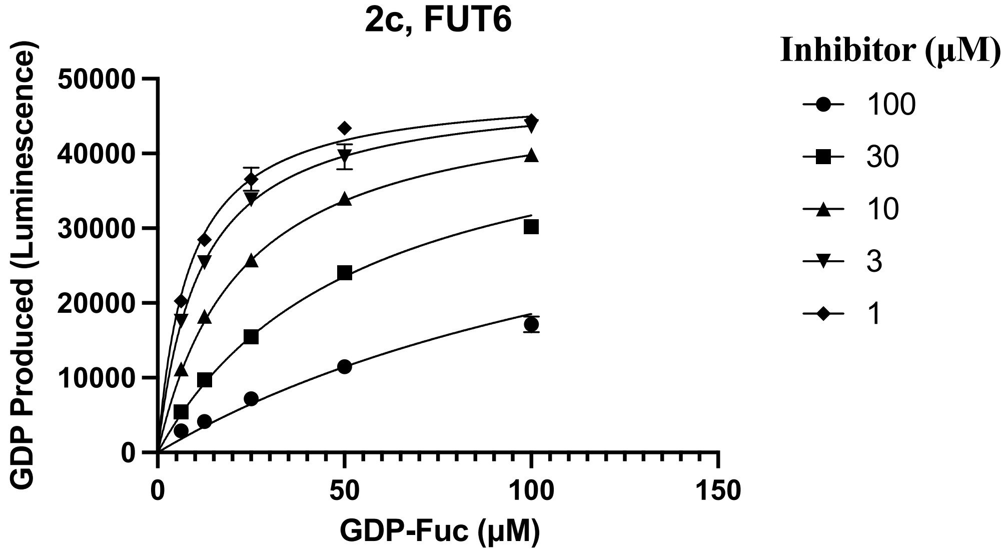
2c, FUT6
GDP Produced (Luminescence)
50000
40000
30000
20000
10000
0
0
50
100
150
GDP-Fuc (μM)
Inhibitor (μM)
100
30
10
3
1

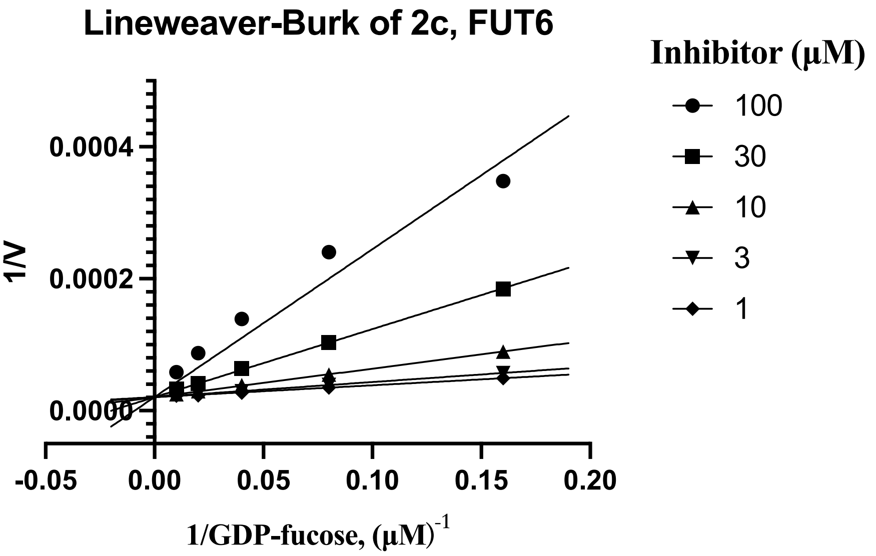
Lineweaver-Burk of 2c, FUT6
1/V
0.0004
0.0002
0.0000
-0.05
0.00
0.05
0.10
0.15
0.20
1/GDP-fucose, (μM)-1
Inhibitor (μM)
100
30
10
3
1

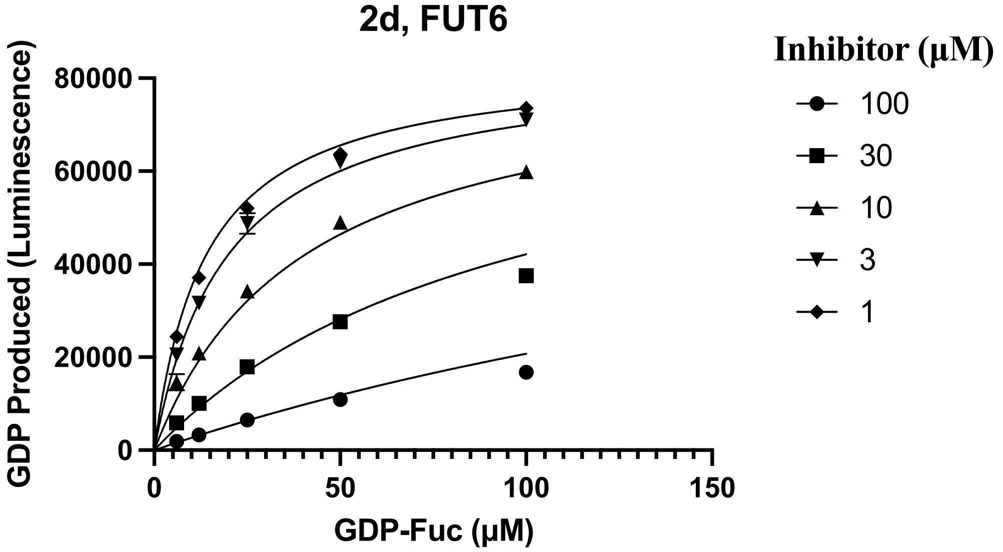
2d, FUT6
GDP Produced (Luminescence)
80000
60000
40000
20000
0
0
50
100
150
GDP-Fuc (μM)
Inhibitor (μM)
100
30
10
3
1

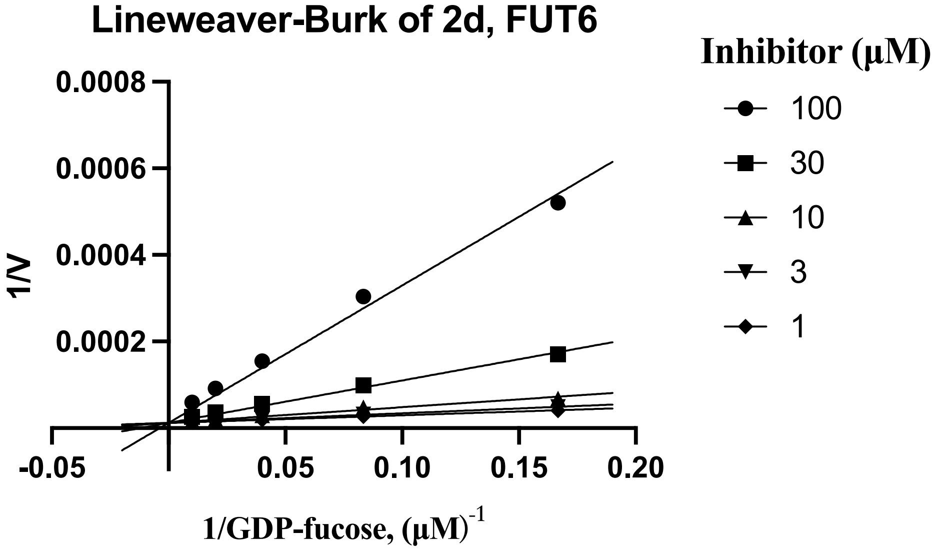
Lineweaver-Burk of 2d, FUT6
1/V
0.0008
0.0006
0.0004
0.0002
-0.05
0.05
0.10
0.15
0.20
1/GDP-fucose, (μM)-1
Inhibitor (μM)
100
30
10
3
1

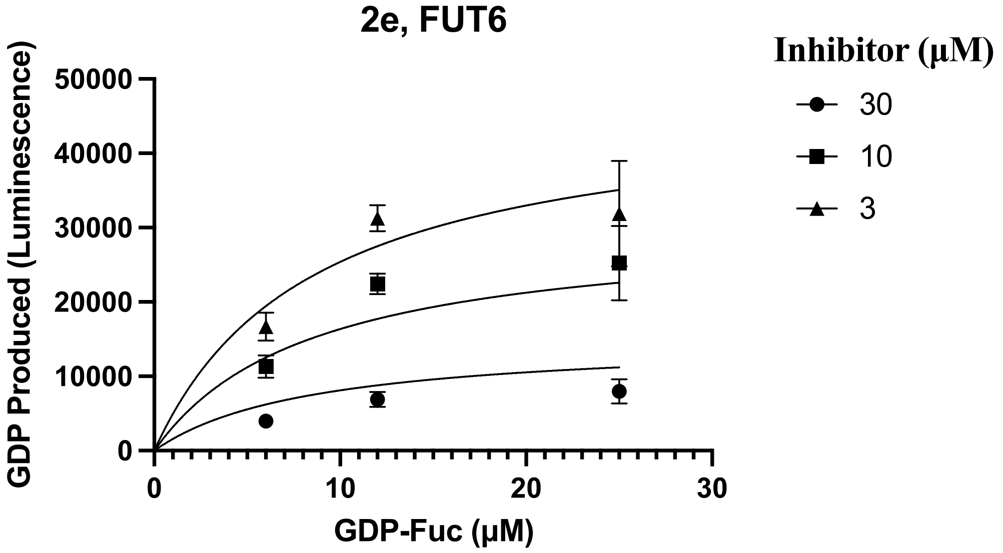
2e, FUT6
Inhibitor (μM)
30
10
3
GDP Produced (Luminescence)
50000
40000
30000
20000
10000
0
0
10
20
30
GDP-Fuc (μM)

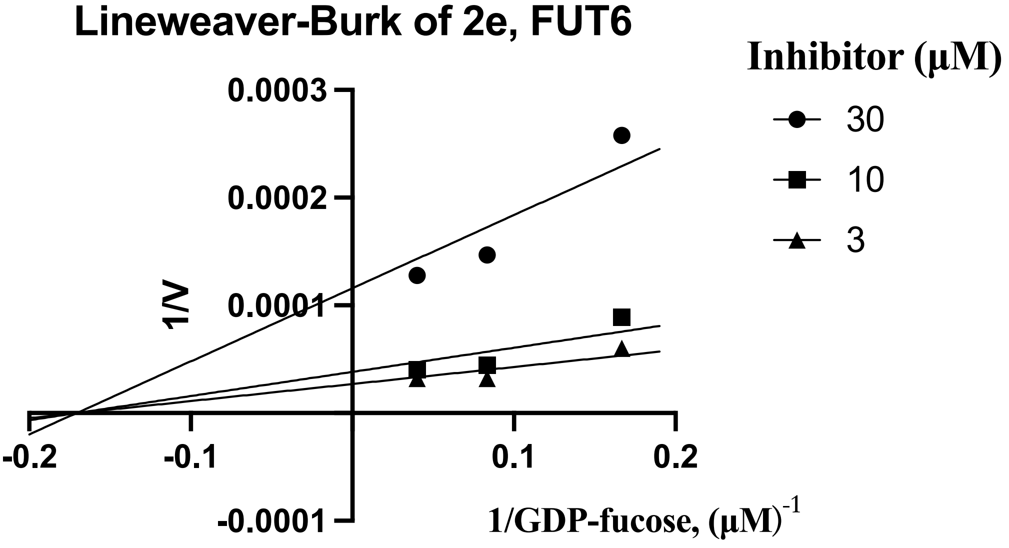
Lineweaver-Burk of 2e, FUT6
Inhibitor (μM)
30
10
3
1/V
0.0003
0.0002
0.0001
-0.0001
-0.2
-0.1
0.1
0.2
1/GDP-fucose, (μM)-1

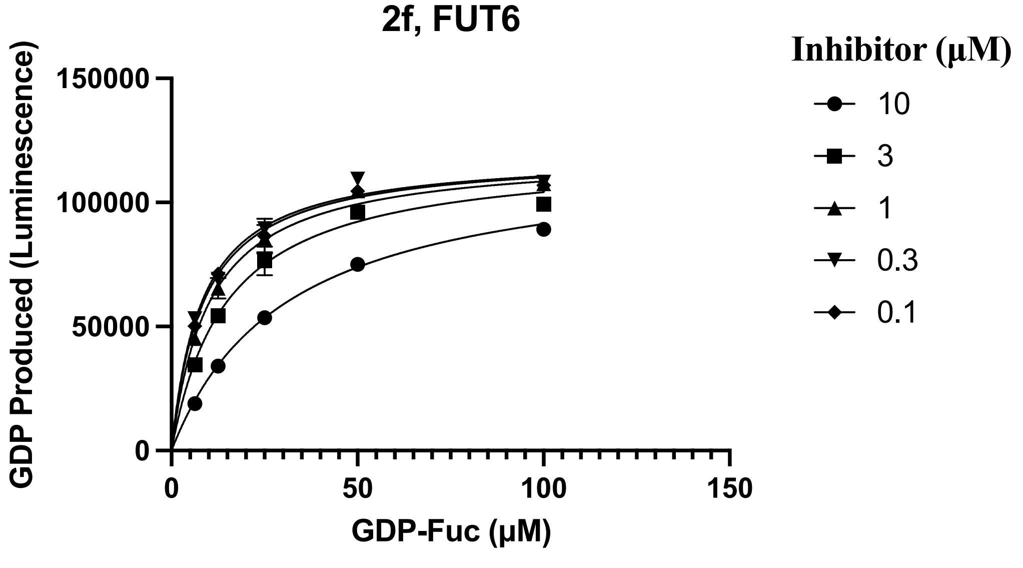
2f, FUT6
Inhibitor (μM)
10
3
1
0.3
0.1
GDP Produced (Luminescence)
150000
100000
50000
0
0
50
100
150
GDP-Fuc (μM)

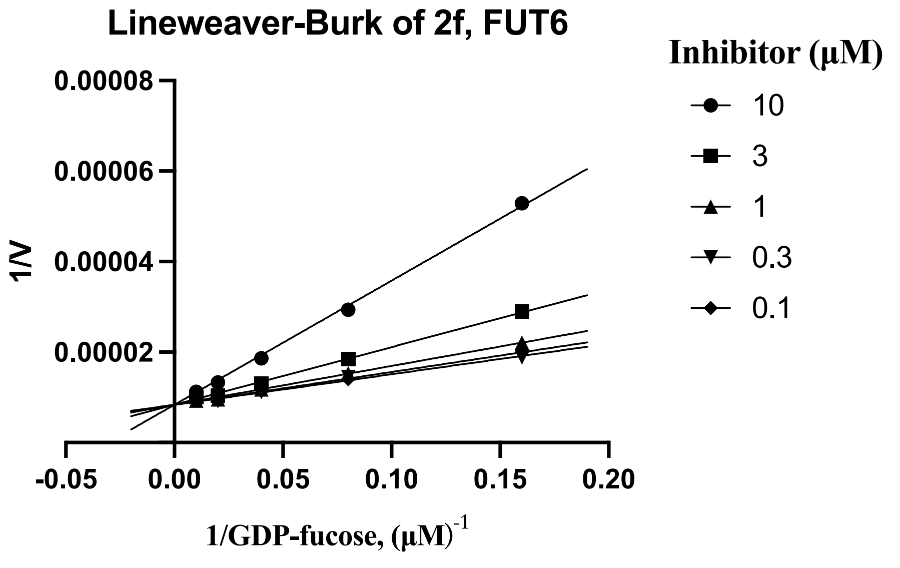

Lineweaver-Burk of 2f, FUT6
0.00008
0.00006
0.00004
0.00002
1/V
-0.05
0.00
0.05
0.10
0.15
0.20
1/GDP-fucose, (μM)-1
Inhibitor (μM)
10
3
1
0.3
0.1


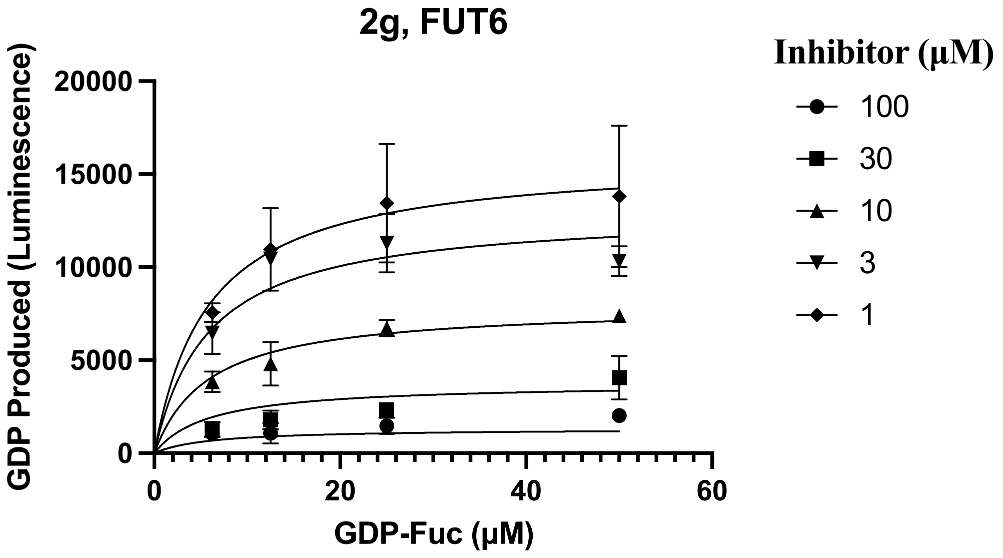

2g, FUT6
20000
15000
10000
5000
0
GDP Produced (Luminescence)
0
20
40
60
GDP-Fuc (μM)
Inhibitor (μM)
100
30
10
3
1


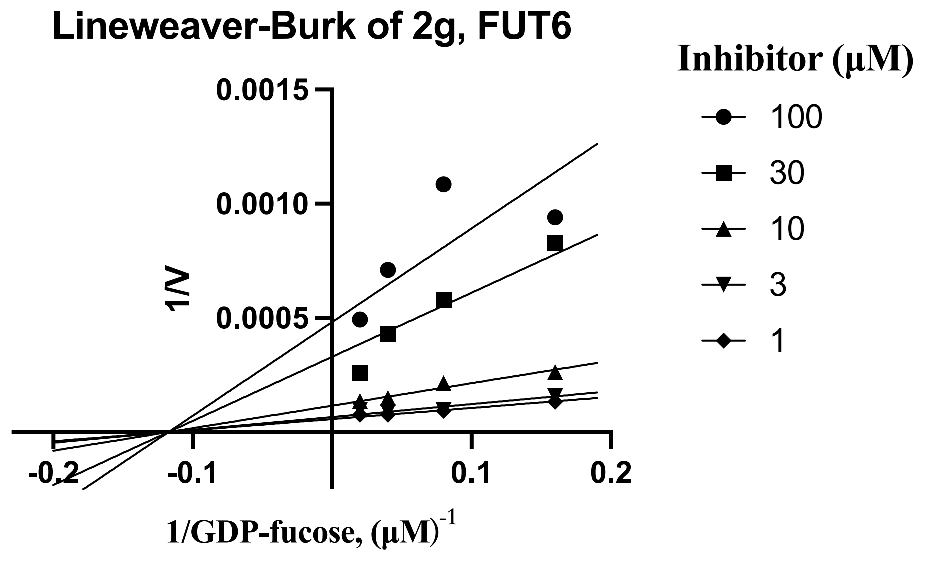

Lineweaver-Burk of 2g, FUT6
0.0015
0.0010
0.0005
1/V
-0.2
-0.1
0.1
0.2
1/GDP-fucose, (μM)-1
Inhibitor (μM)
100
30
10
3
1

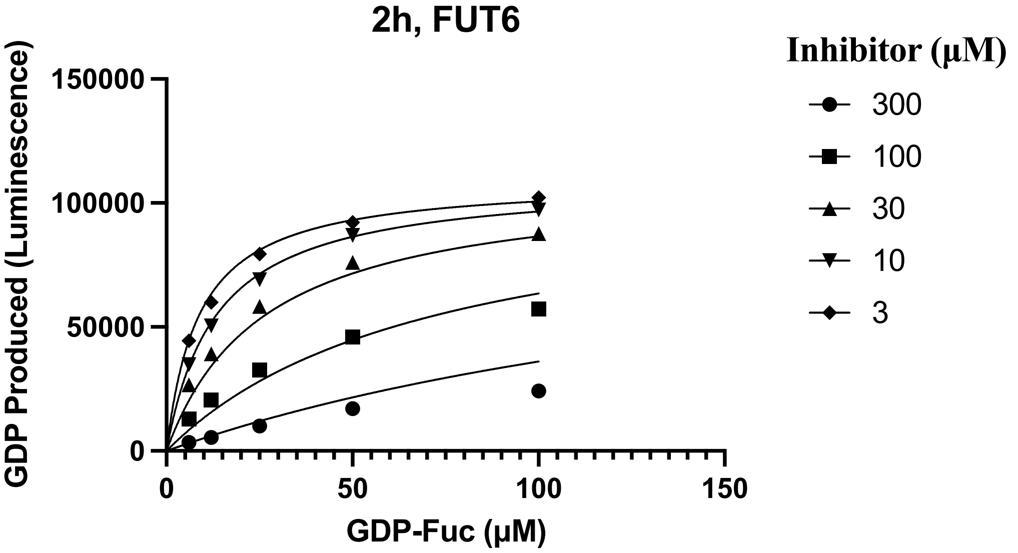
2h, FUT6
Inhibitor (µM)
300
100
30
10
3
GDP Produced (Luminescence)
150000
100000
50000
0
0
50
100
150
GDP-Fuc (µM)

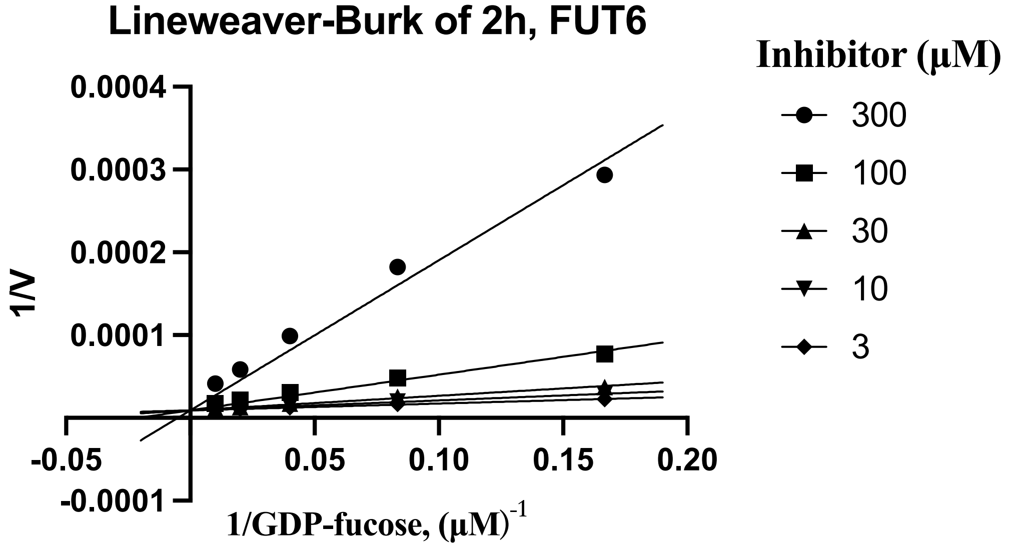
Lineweaver-Burk of 2h, FUT6
Inhibitor (µM)
300
100
30
10
3
1/V
0.0004
0.0003
0.0002
0.0001
-0.0001
-0.05
0.05
0.10
0.15
0.20
1/GDP-fucose, (µM)-1

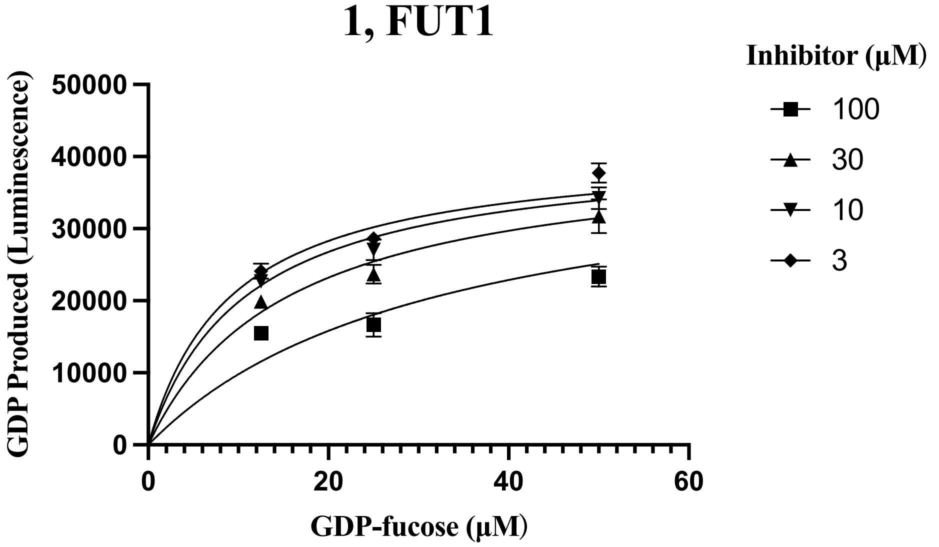
1, FUT1
Inhibitor (µM)
100
30
10
3
GDP Produced (Luminescence)
50000
40000
30000
20000
10000
0
0
20
40
60
GDP-fucose (µM)

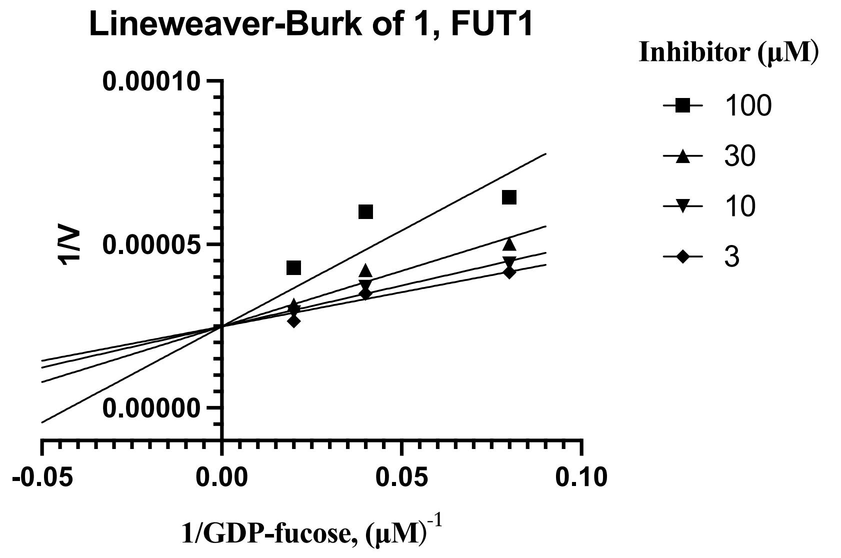

Lineweaver-Burk of 1, FUT1
Inhibitor (μM)
100
30
10
3
1/V
0.00010
0.00005
0.00000
-0.05
0.00
0.05
0.10
1/GDP-fucose, (μM)-1


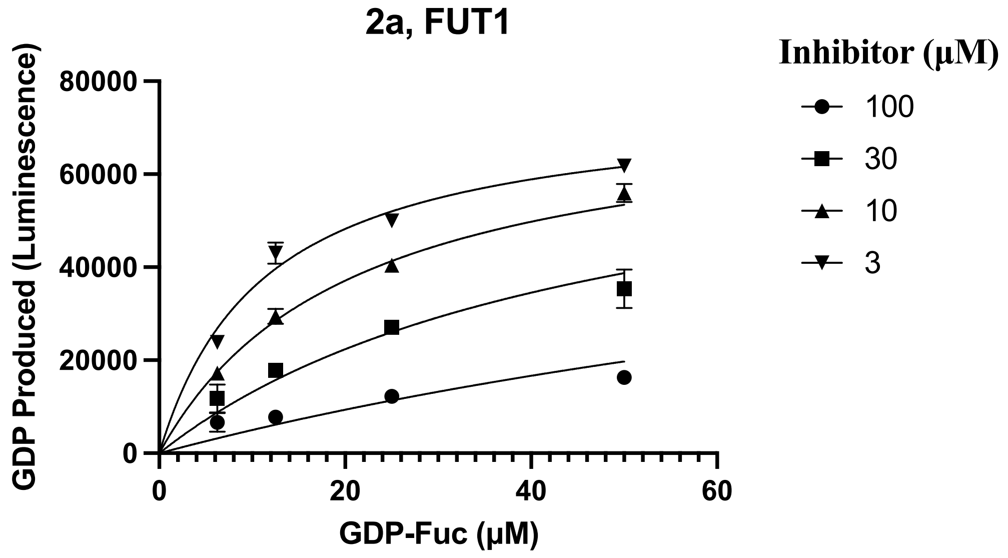

2a, FUT1
Inhibitor (μM)
100
30
10
3
GDP Produced (Luminescence)
80000
60000
40000
20000
0
0
20
40
60
GDP-Fuc (μM)


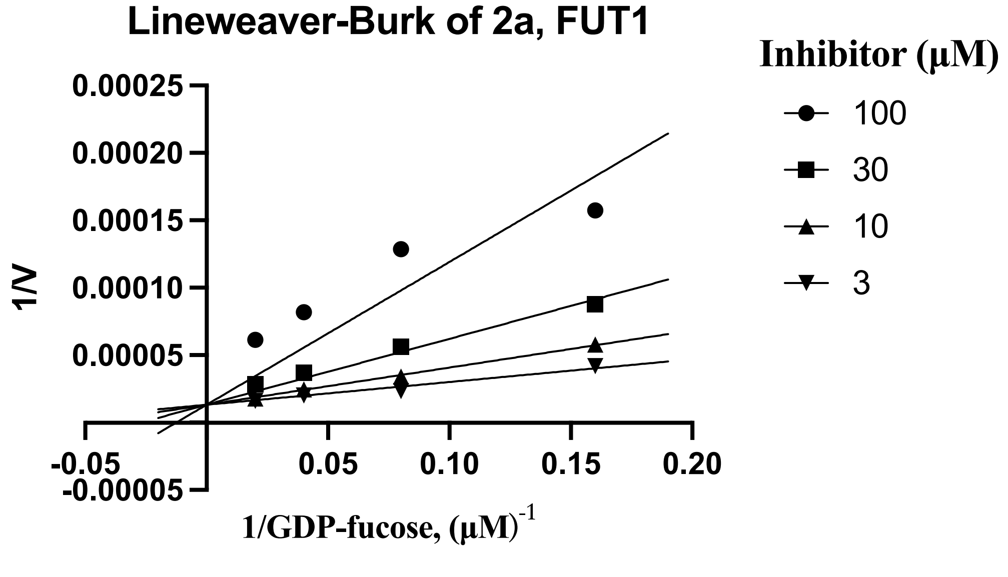

Lineweaver-Burk of 2a, FUT1
Inhibitor (μM)
100
30
10
3
1/V
0.00025
0.00020
0.00015
0.00010
0.00005
-0.00005
-0.05
0.05
0.10
0.15
0.20
1/GDP-fucose, (μM)-1

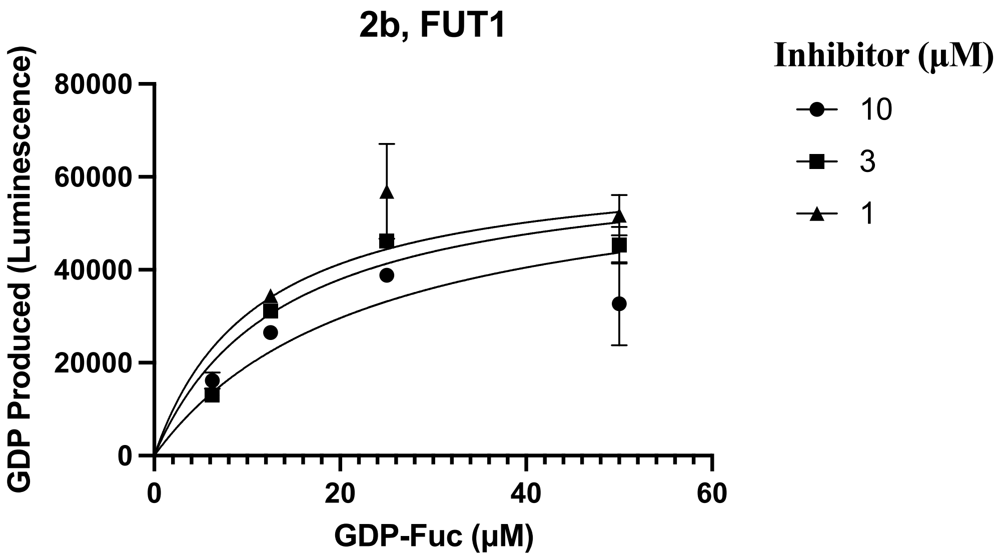
2b, FUT1
GDP Produced (Luminescence)
80000
60000
40000
20000
0
0
20
40
60
GDP-Fuc (μM)
Inhibitor (μM)
10
3
1

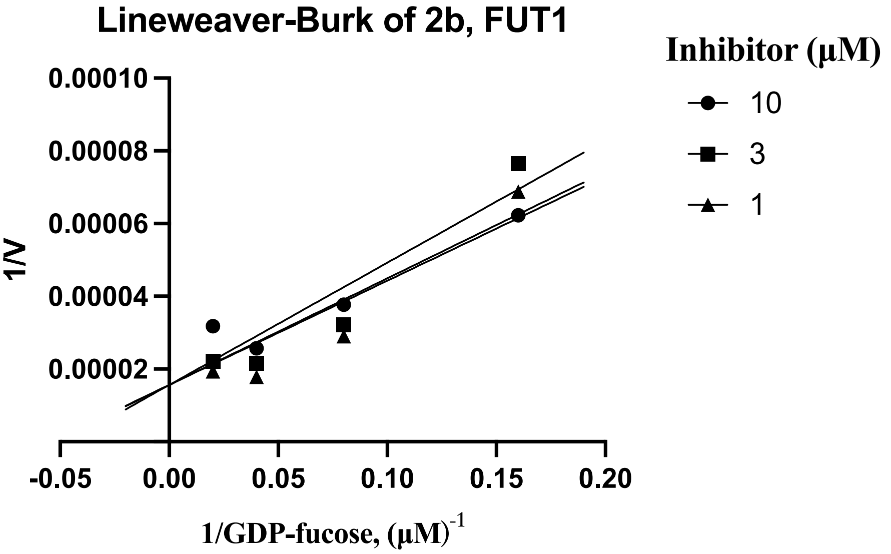
Lineweaver-Burk of 2b, FUT1
1/V
0.00010
0.00008
0.00006
0.00004
0.00002
-0.05
0.00
0.05
0.10
0.15
0.20
1/GDP-fucose, (μM)-1
Inhibitor (μM)
10
3
1

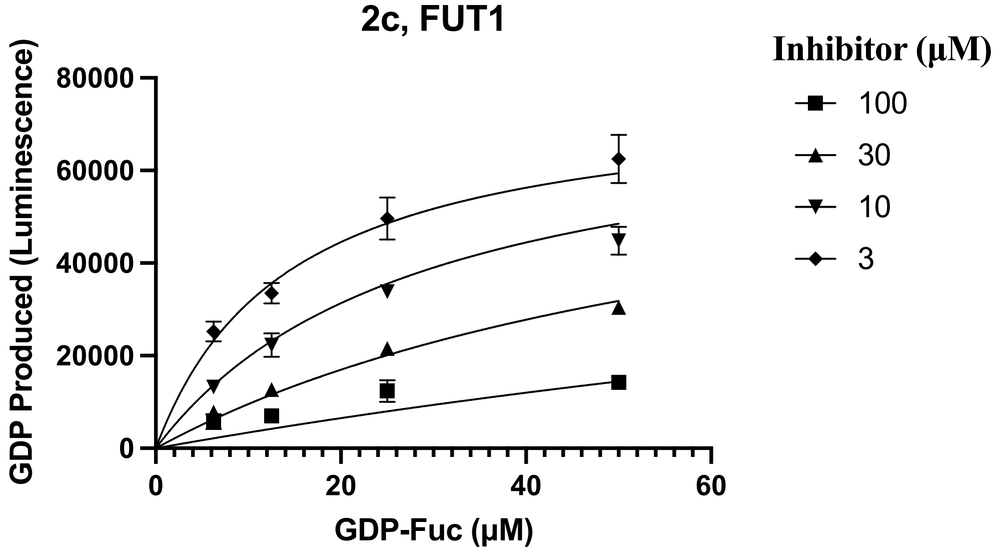
2c, FUT1
GDP Produced (Luminescence)
80000
60000
40000
20000
0
0
20
40
60
GDP-Fuc (μM)
Inhibitor (μM)
100
30
10
3

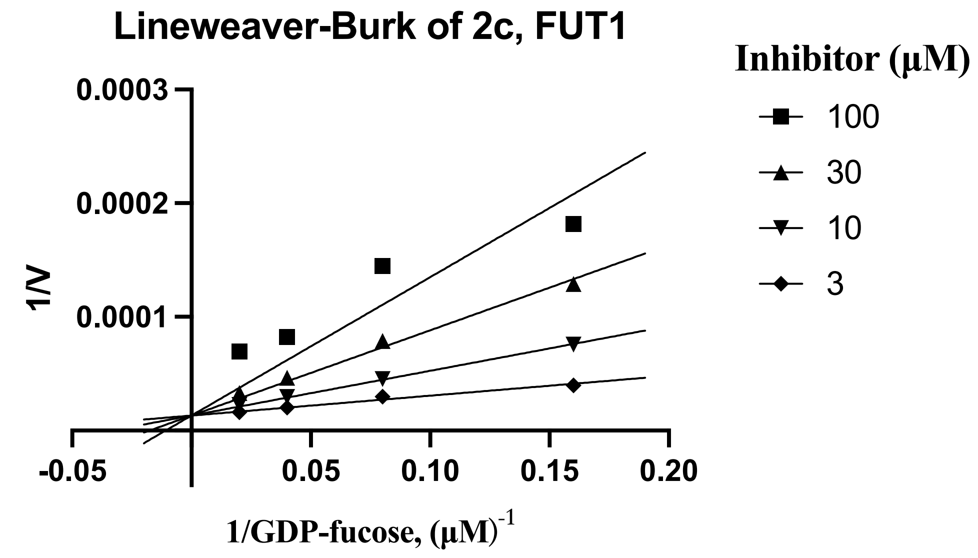
Lineweaver-Burk of 2c, FUT1
0.0003
0.0002
0.0001
1/V
-0.05
0.05
0.10
0.15
0.20
1/GDP-fucose, (µM)-1
Inhibitor (µM)
100
30
10
3

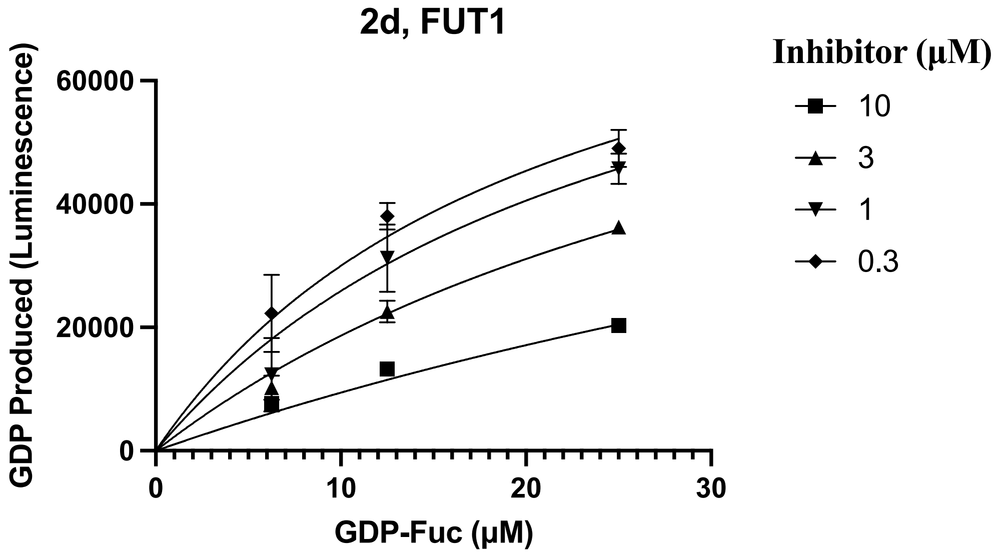
2d, FUT1
60000
40000
20000
0
GDP Produced (Luminescence)
0
10
20
30
GDP-Fuc (µM)
Inhibitor (µM)
10
3
1
0.3

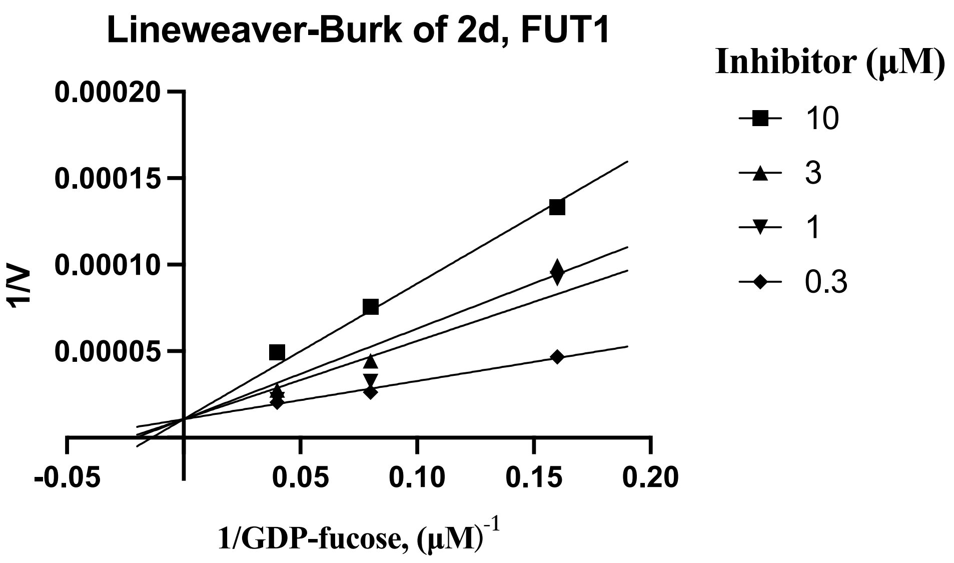
Lineweaver-Burk of 2d, FUT1
0.00020
0.00015
0.00010
0.00005
1/V
-0.05
0.05
0.10
0.15
0.20
1/GDP-fucose, (µM)-1
Inhibitor (µM)
10
3
1
0.3

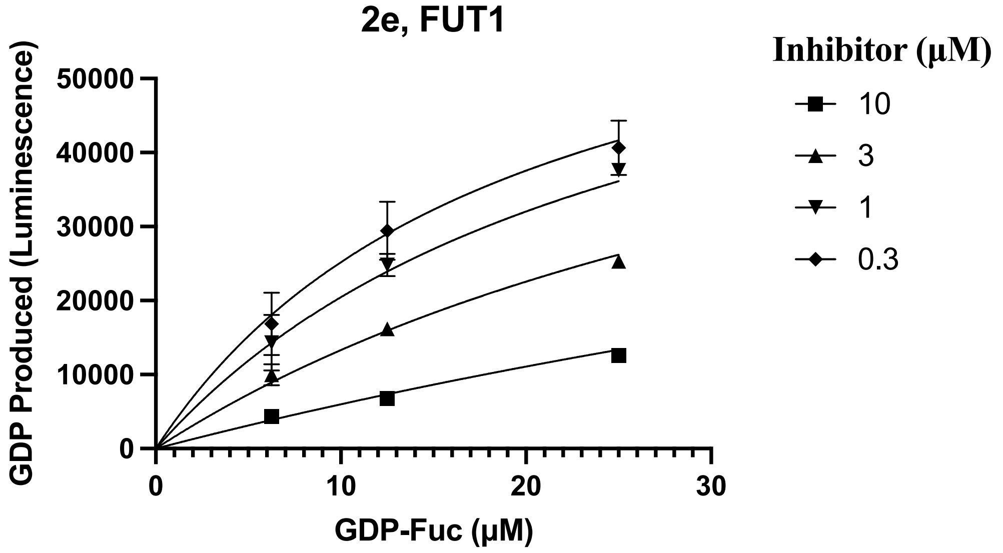
2e, FUT1
Inhibitor (µM)
10
3
1
0.3
GDP Produced (Luminescence)
50000
40000
30000
20000
10000
0
0
10
20
30
GDP-Fuc (µM)

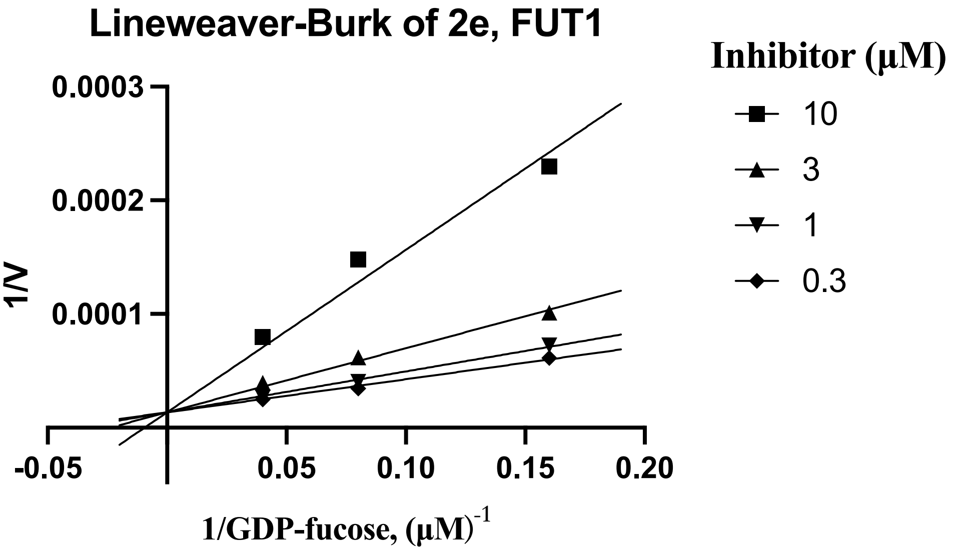
Lineweaver-Burk of 2e, FUT1
Inhibitor (µM)
10
3
1
0.3
1/V
0.0003
0.0002
0.0001
-0.05
0.05
0.10
0.15
0.20
1/GDP-fucose, (µM)-1

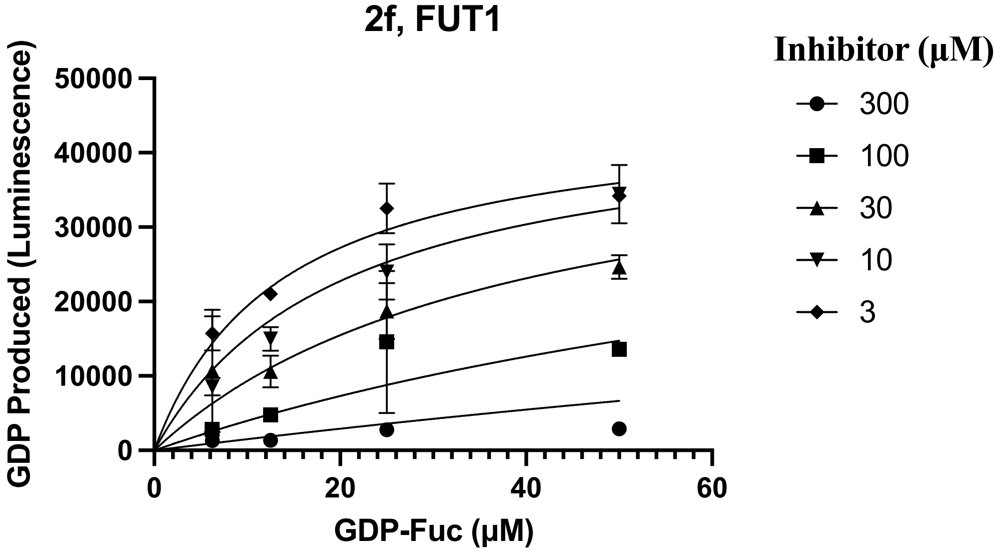
2f, FUT1
Inhibitor (µM)
300
100
30
10
3
GDP Produced (Luminescence)
50000
40000
30000
20000
10000
0
0
20
40
60
GDP-Fuc (µM)

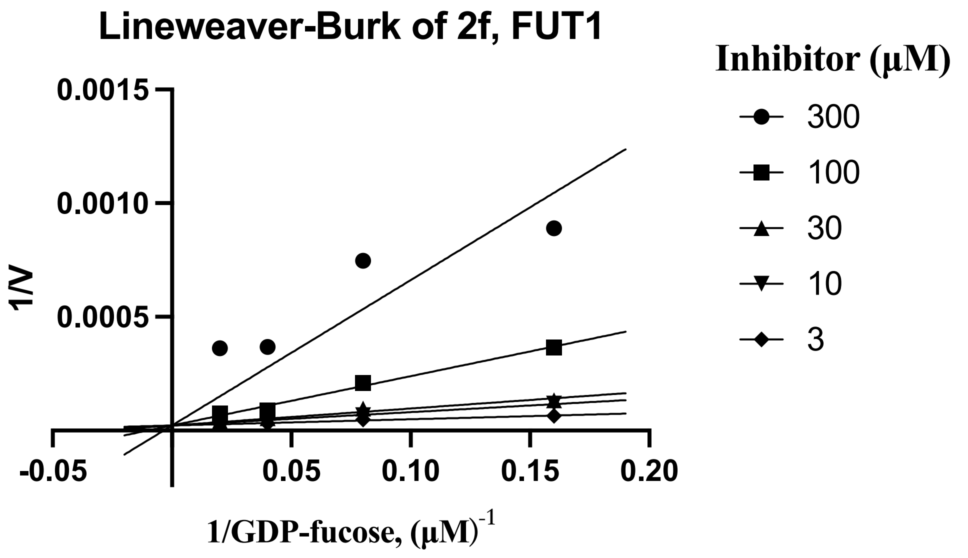
Lineweaver-Burk of 2f, FUT1
Inhibitor (µM)
300
100
30
10
3
1/V
0.0015
0.0010
0.0005
-0.05
0.05
0.10
0.15
0.20
1/GDP-fucose, (µM)-1

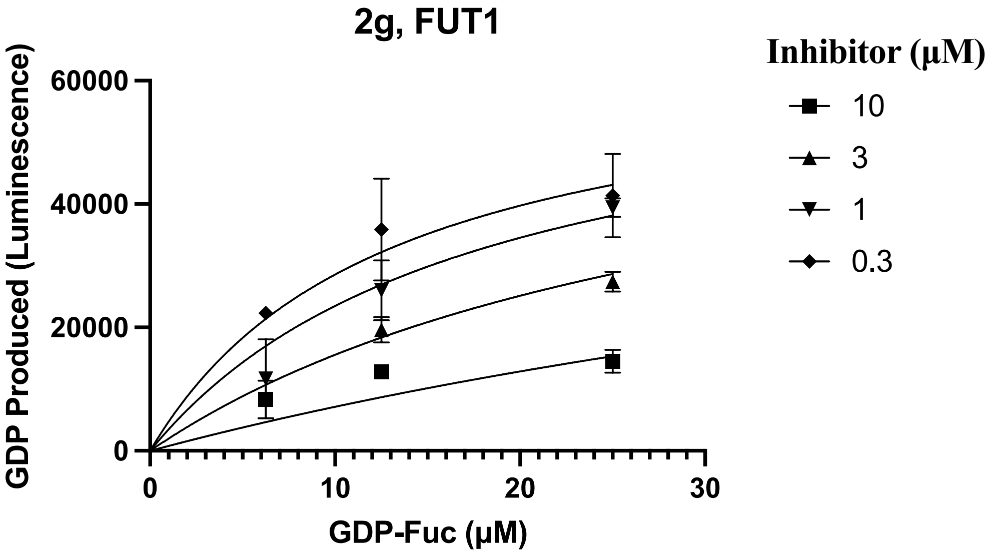
2g, FUT1
Inhibitor (µM)
10
3
1
0.3
GDP Produced (Luminescence)
60000
40000
20000
0
0
10
20
30
GDP-Fuc (µM)

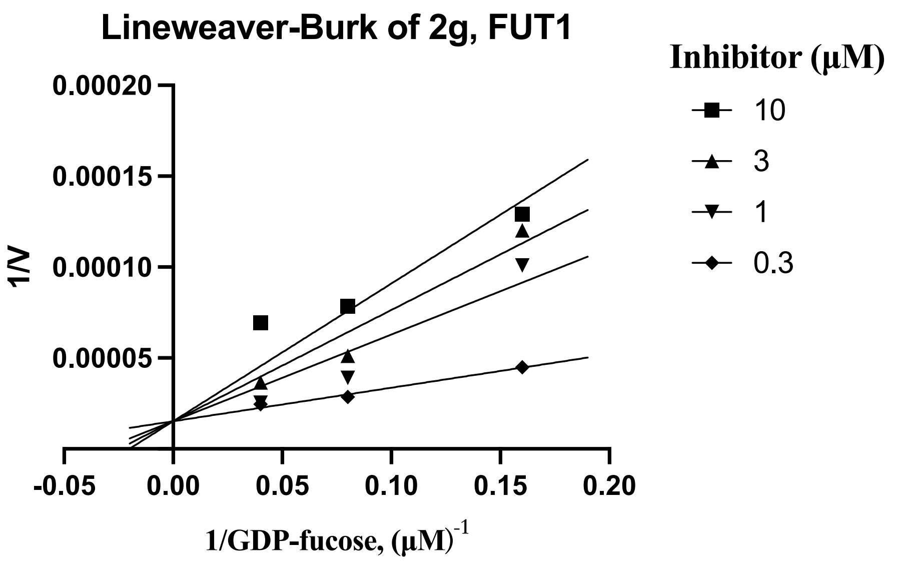
Lineweaver-Burk of 2g, FUT1
Inhibitor (µM)
10
3
1
0.3
1/V
0.00020
0.00015
0.00010
0.00005
-0.05
0.00
0.05
0.10
0.15
0.20
1/GDP-fucose, (µM)-1

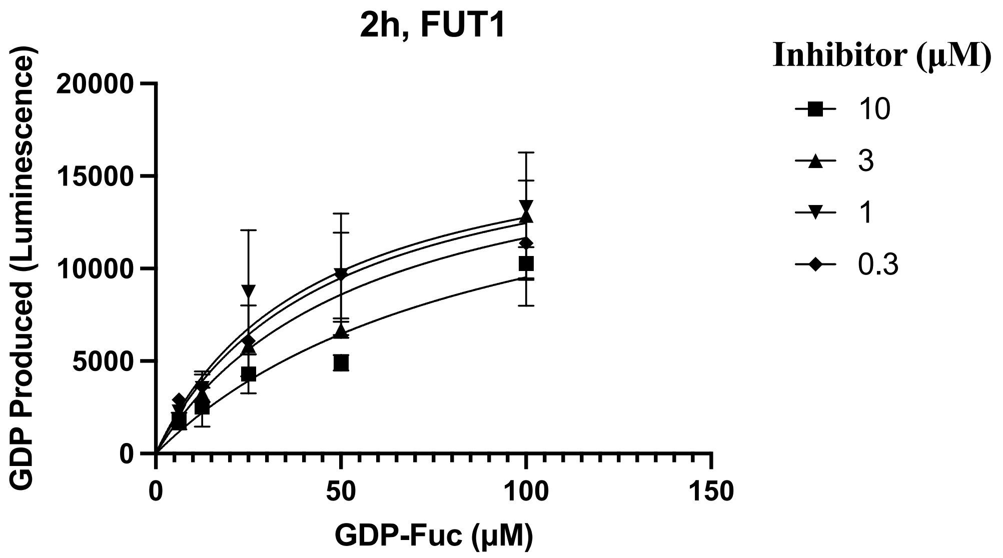
2h, FUT1
GDP Produced (Luminescence)
20000
15000
10000
5000
0
0
50
100
150
GDP-Fuc (µM)
Inhibitor (µM)
10
3
1
0.3

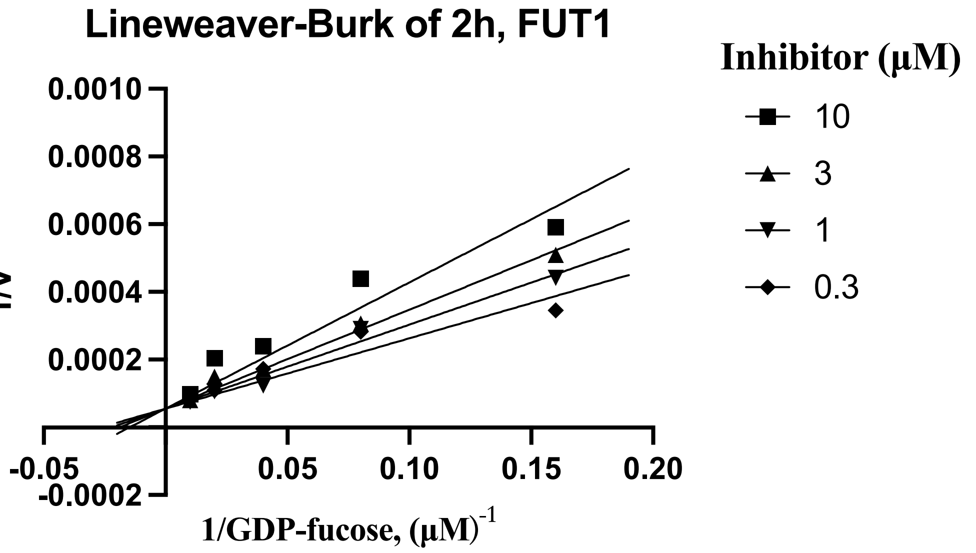
Lineweaver-Burk of 2h, FUT1
1/V
0.0010
0.0008
0.0006
0.0004
0.0002
-0.0002
-0.05
0.05
0.10
0.15
0.20
1/GDP-fucose, (µM)-1
Inhibitor (µM)
10
3
1
0.3

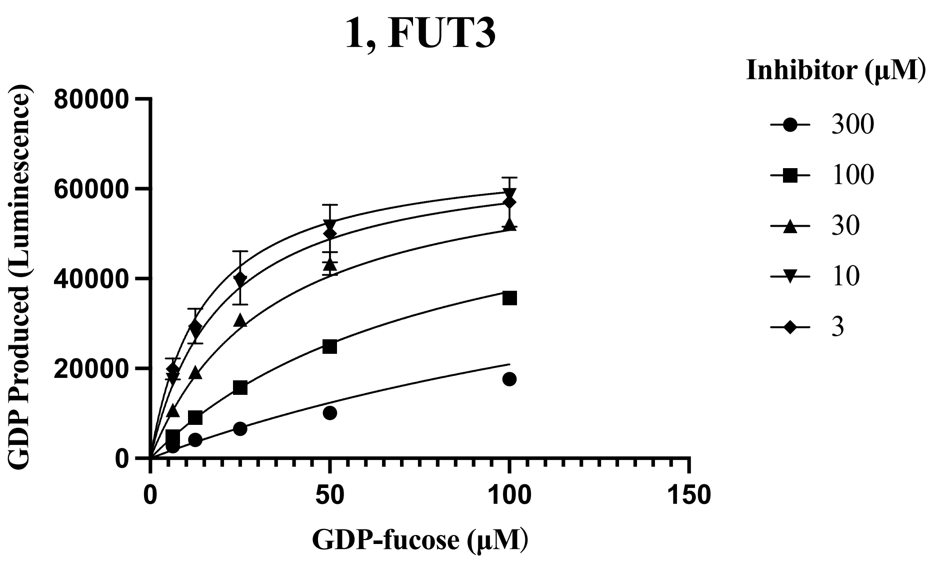
1, FUT3
GDP Produced (Luminescence)
80000
60000
40000
20000
0
0
50
100
150
GDP-fucose (µM)
Inhibitor (µM)
300
100
30
10
3

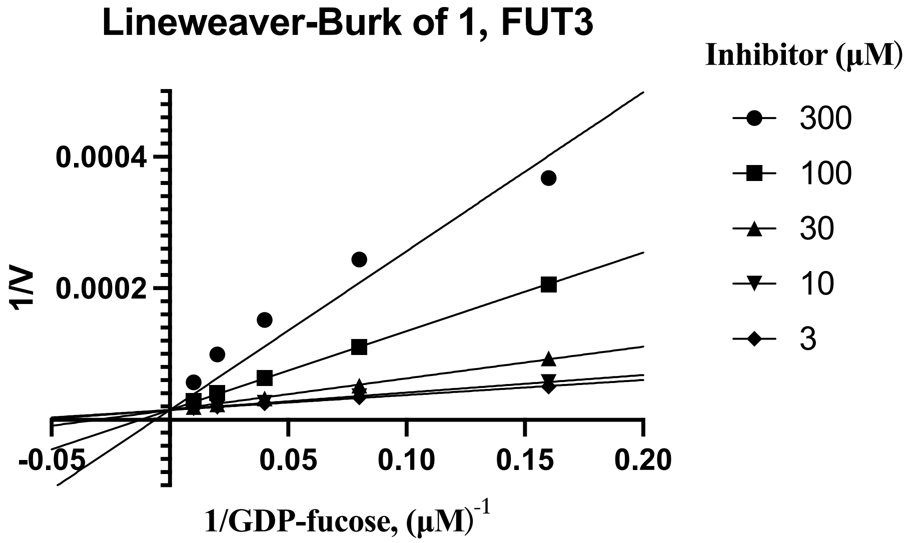
Lineweaver-Burk of 1, FUT3
Inhibitor (µM)
300
100
30
10
3
1/V
0.0004
0.0002
-0.05
0.05
0.10
0.15
0.20
1/GDP-fucose, (µM)-1

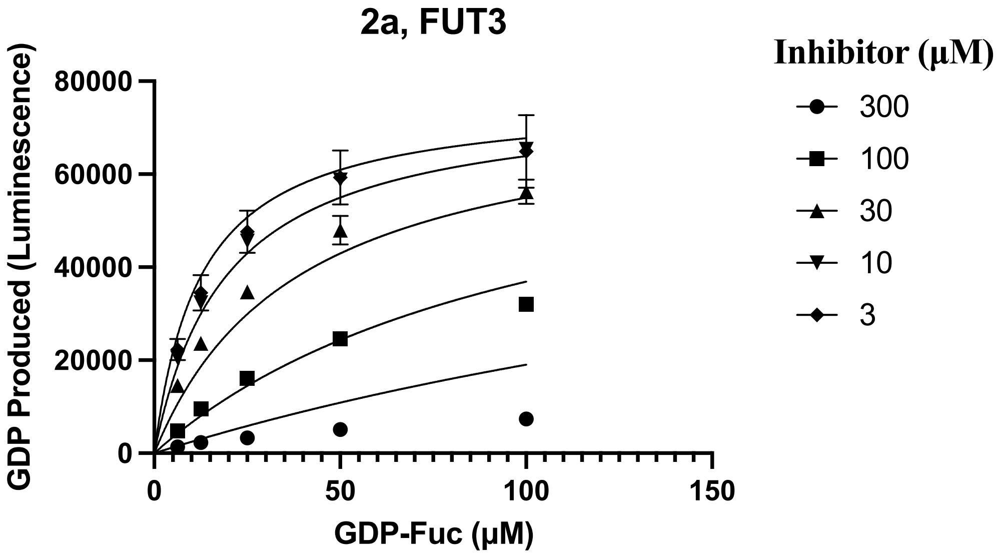
2a, FUT3
Inhibitor (µM)
300
100
30
10
3
GDP Produced (Luminescence)
80000
60000
40000
20000
0
0
50
100
150
GDP-Fuc (µM)

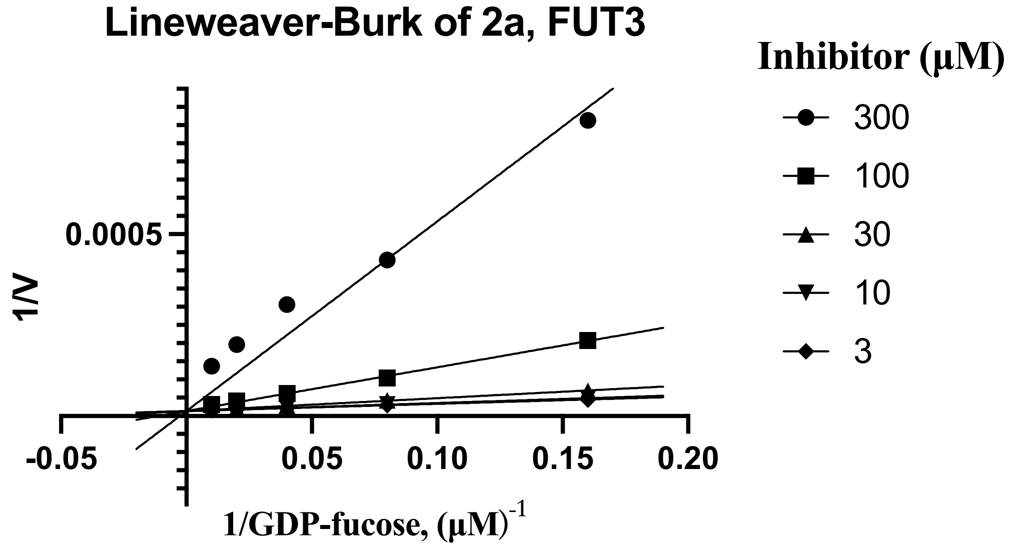
Lineweaver-Burk of 2a, FUT3
Inhibitor (µM)
300
100
30
10
3
1/V
0.0005
-0.05
0.05
0.10
0.15
0.20
1/GDP-fucose, (µM)-1

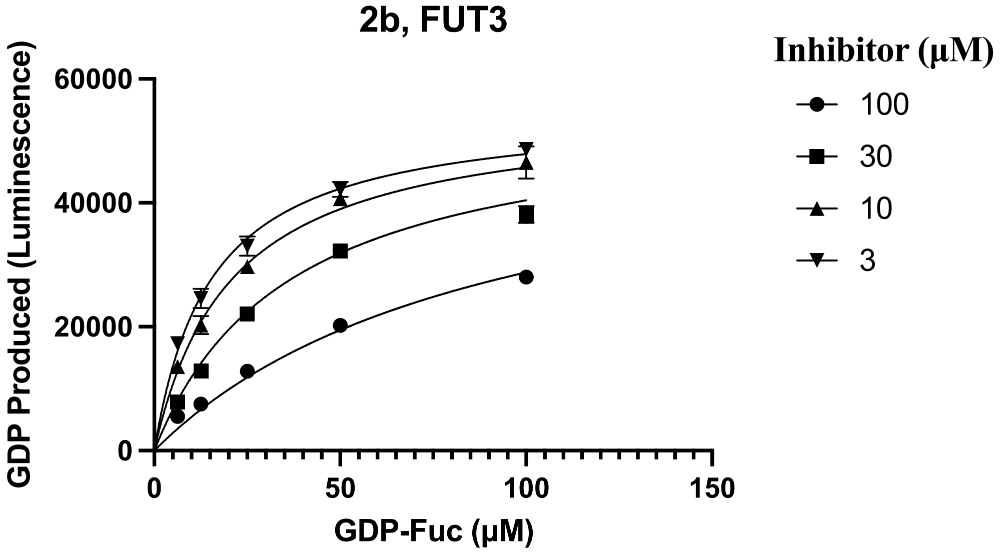
2b, FUT3
GDP Produced (Luminescence)
60000
40000
20000
0
0
50
100
150
GDP-Fuc (μM)
Inhibitor (μM)
100
30
10
3

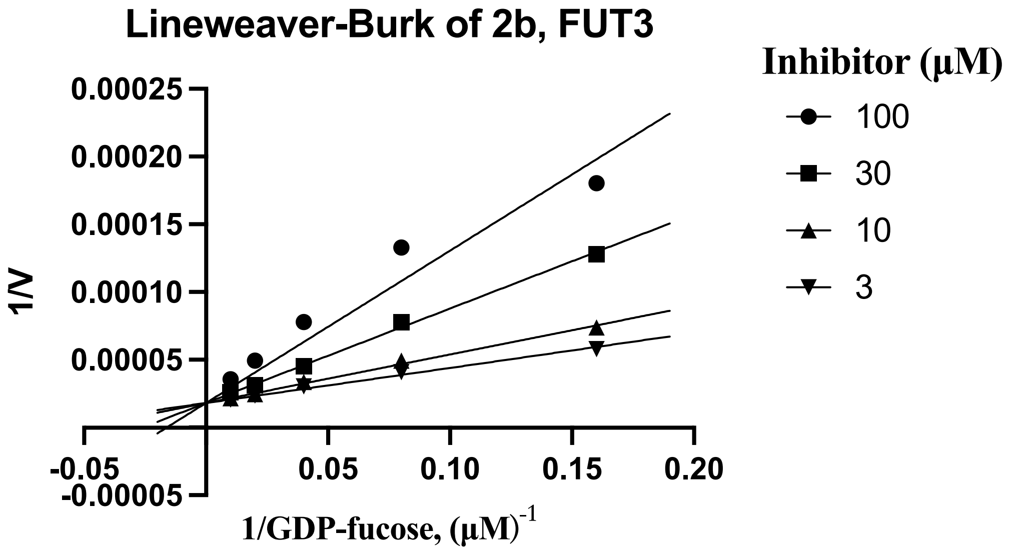
Lineweaver-Burk of 2b, FUT3
1/V
0.00025
0.00020
0.00015
0.00010
0.00005
-0.00005
-0.05
0.05
0.10
0.15
0.20
1/GDP-fucose, (μM)-1
Inhibitor (μM)
100
30
10
3

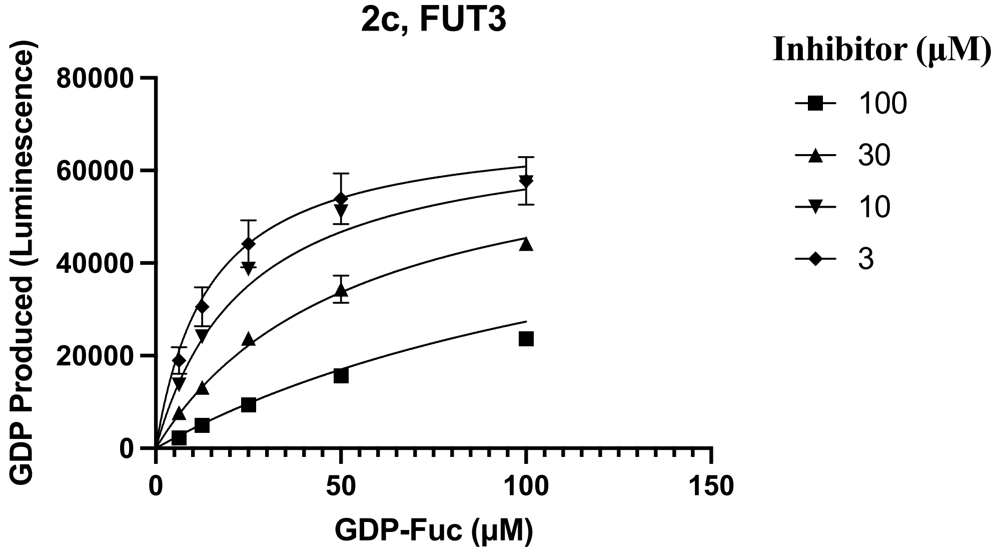
2c, FUT3
GDP Produced (Luminescence)
80000
60000
40000
20000
0
0
50
100
150
GDP-Fuc (μM)
Inhibitor (μM)
100
30
10
3

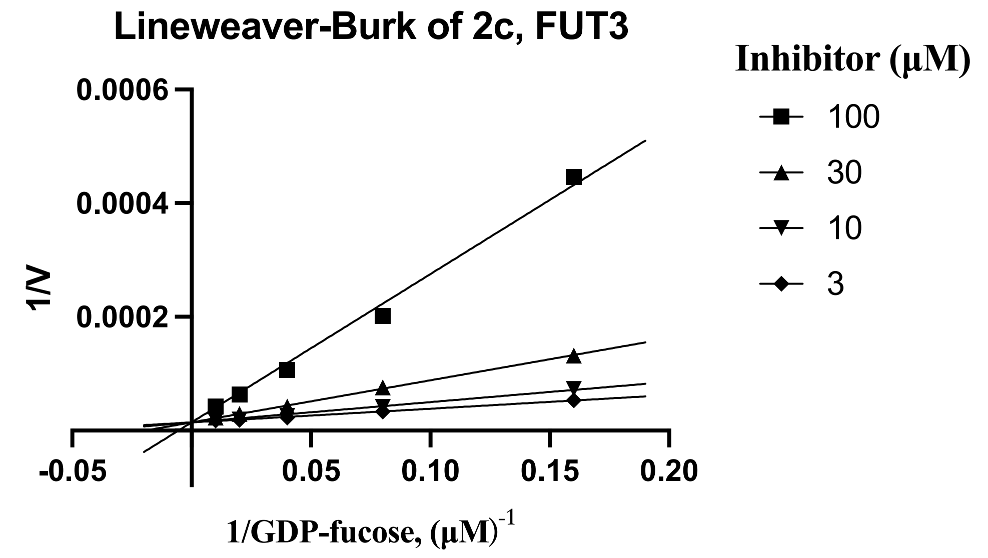

Lineweaver-Burk of 2c, FUT3
Inhibitor (µM)
100
30
10
3
0.0006
0.0004
0.0002
1/V
-0.05
0.05
0.10
0.15
0.20
1/GDP-fucose, (µM)-1


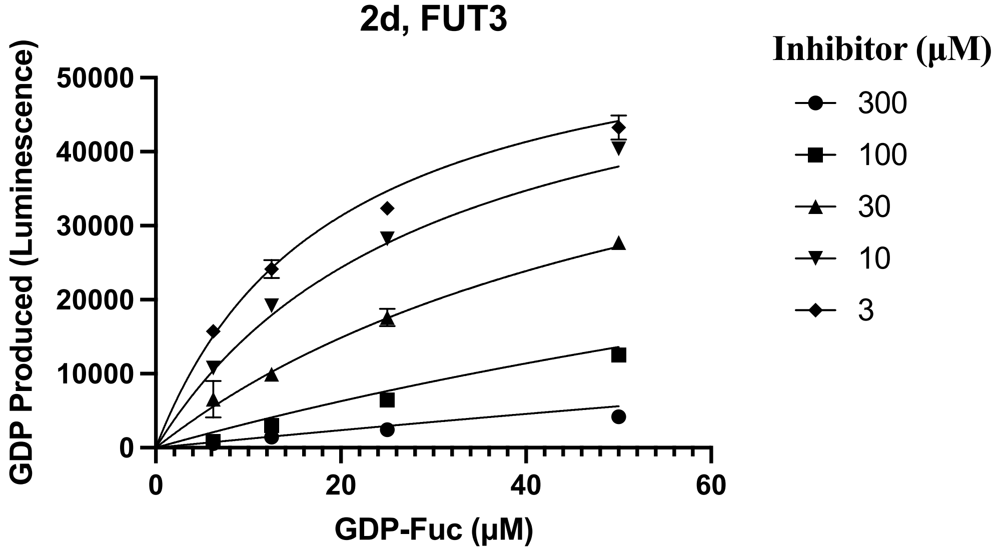

2d, FUT3
Inhibitor (µM)
300
100
30
10
3
50000
40000
30000
20000
10000
0
GDP Produced (Luminescence)
0
20
40
60
GDP-Fuc (µM)


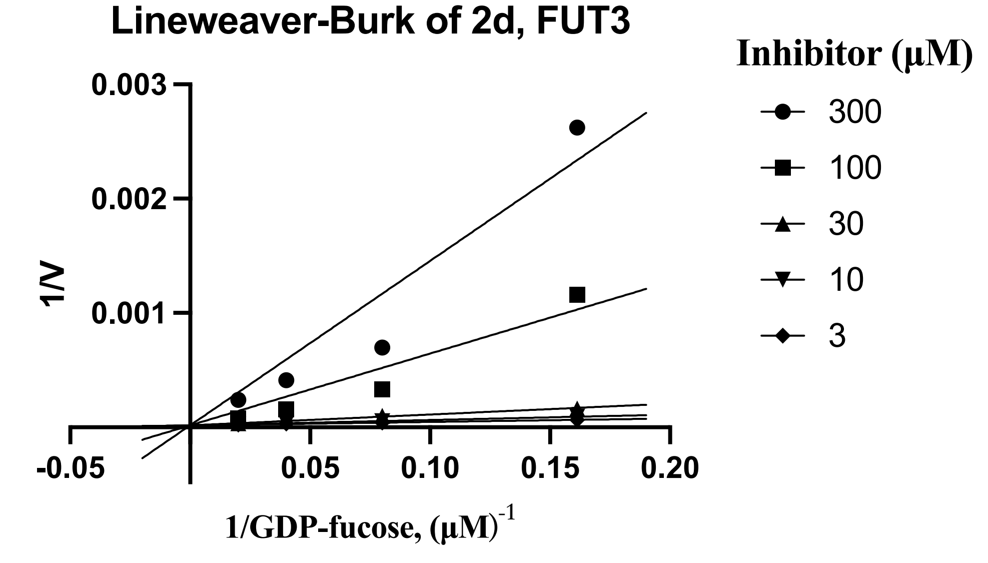

Lineweaver-Burk of 2d, FUT3
Inhibitor (µM)
300
100
30
10
3
0.003
0.002
0.001
1/V
-0.05
0.05
0.10
0.15
0.20
1/GDP-fucose, (µM)-1

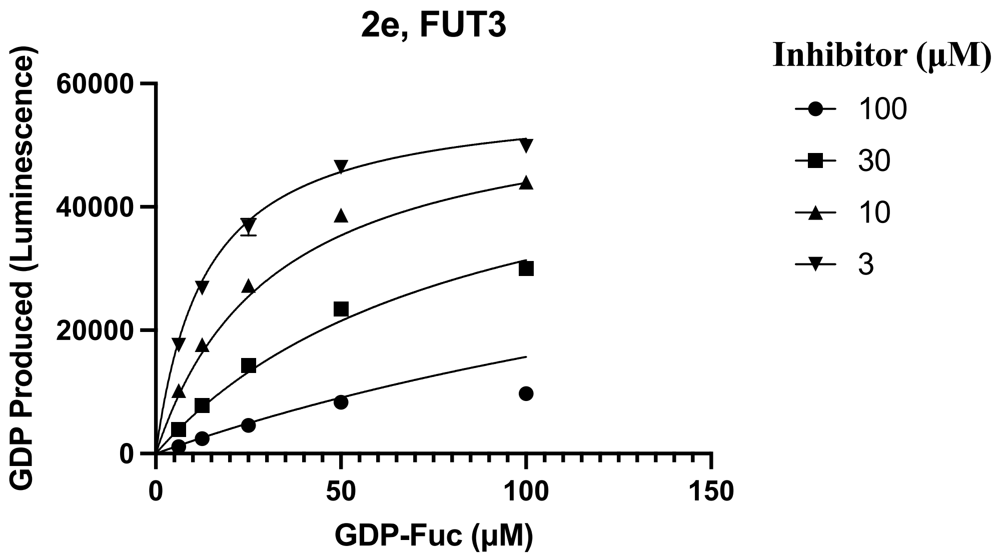
2e, FUT3
GDP Produced (Luminescence)
60000
40000
20000
0
0
50
100
150
GDP-Fuc (µM)
Inhibitor (µM)
100
30
10
3

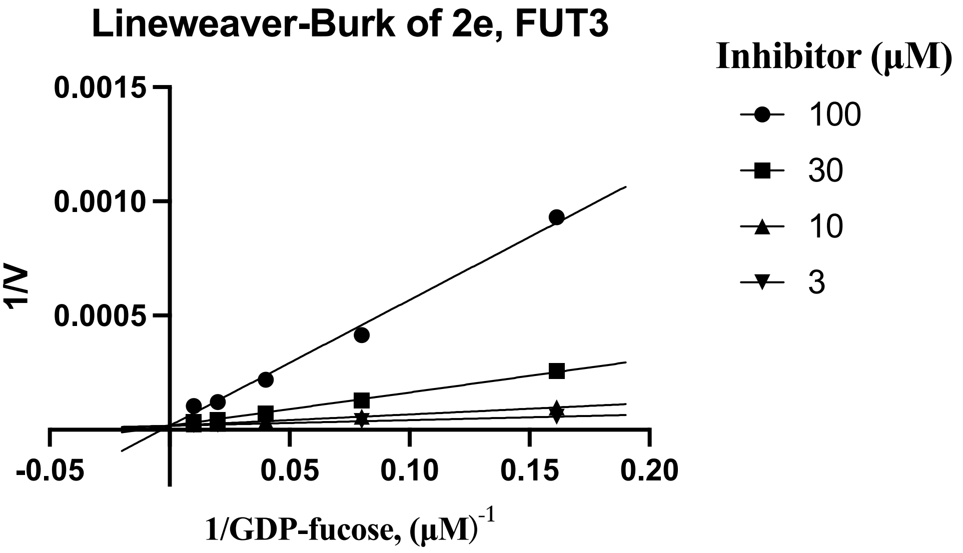
Lineweaver-Burk of 2e, FUT3
1/V
0.0015
0.0010
0.0005
-0.05
0.05
0.10
0.15
0.20
1/GDP-fucose, (µM)-1
Inhibitor (µM)
100
30
10
3

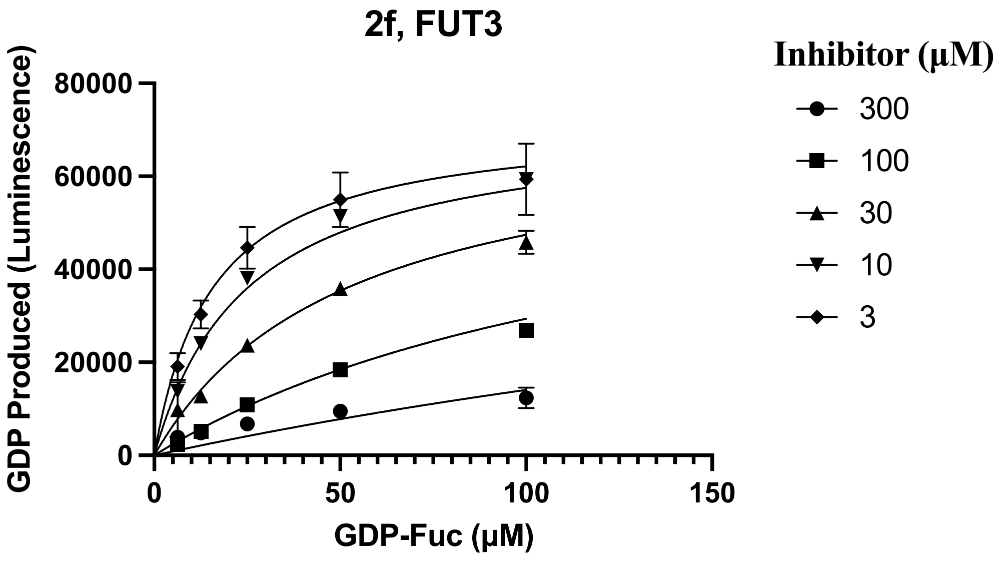
2f, FUT3
GDP Produced (Luminescence)
80000
60000
40000
20000
0
0
50
100
150
GDP-Fuc (µM)
Inhibitor (µM)
300
100
30
10
3

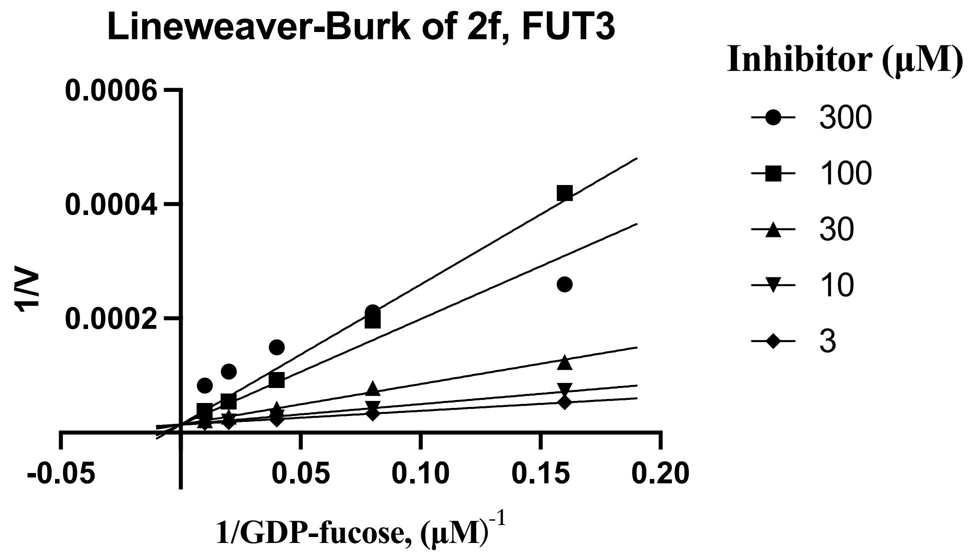
Lineweaver-Burk of 2f, FUT3
Inhibitor (μM)
300
100
30
10
3
0.0006
0.0004
0.0002
1/V
-0.05
0.05
0.10
0.15
0.20
1/GDP-fucose, (μM)-1

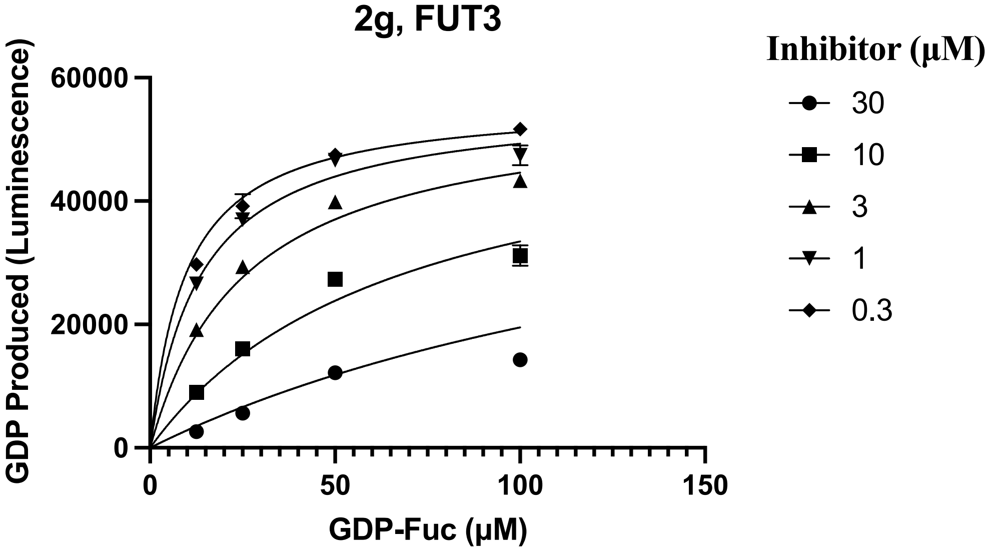
2g, FUT3
Inhibitor (μM)
30
10
3
1
0.3
60000
40000
20000
0
GDP Produced (Luminescence)
0
50
100
150
GDP-Fuc (μM)

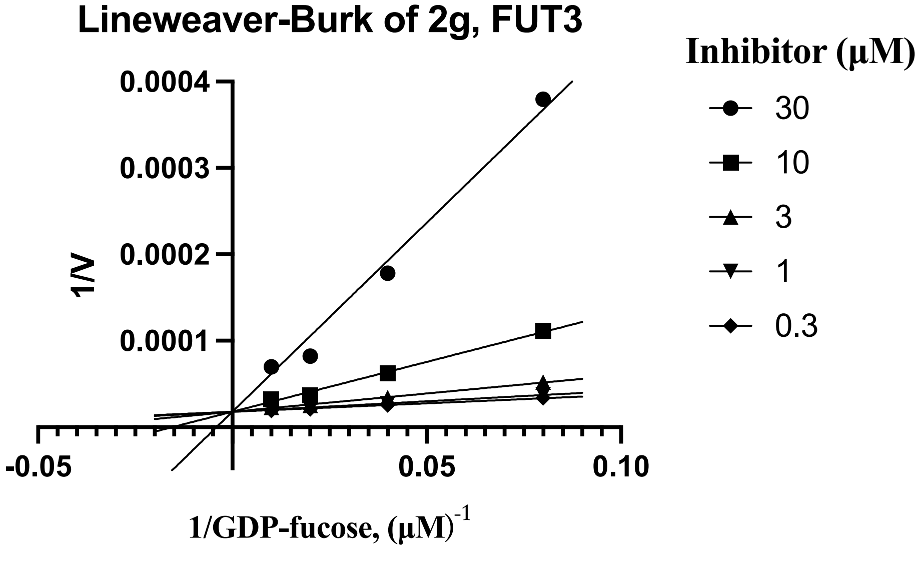
Lineweaver-Burk of 2g, FUT3
Inhibitor (μM)
30
10
3
1
0.3
0.0004
0.0003
0.0002
0.0001
1/V
-0.05
0.05
0.10
1/GDP-fucose, (μM)-1

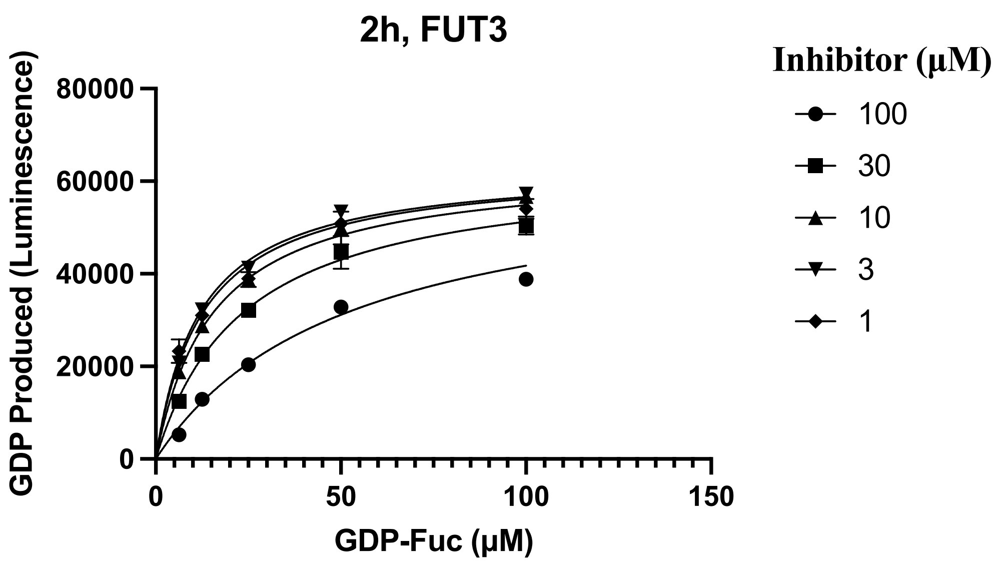

2h, FUT3
GDP Produced (Luminescence)
80000
60000
40000
20000
0
0
50
100
150
GDP-Fuc (µM)
Inhibitor (µM)
100
30
10
3
1


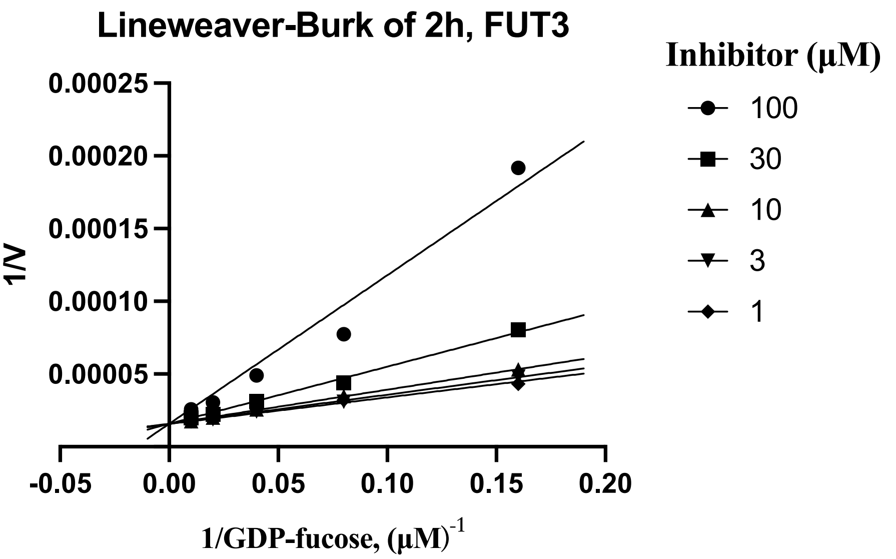

Lineweaver-Burk of 2h, FUT3
1/V
0.00025
0.00020
0.00015
0.00010
0.00005
-0.05
0.00
0.05
0.10
0.15
0.20
1/GDP-fucose, (µM)-1
Inhibitor (µM)
100
30
10
3
1


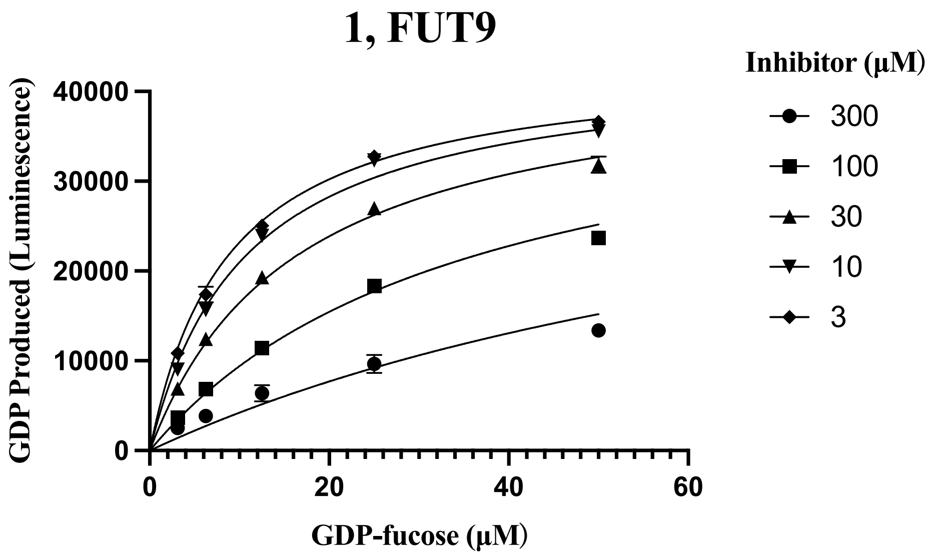

1, FUT9
GDP Produced (Luminescence)
40000
30000
20000
10000
0
0
20
40
60
GDP-fucose (µM)
Inhibitor (µM)
300
100
30
10
3

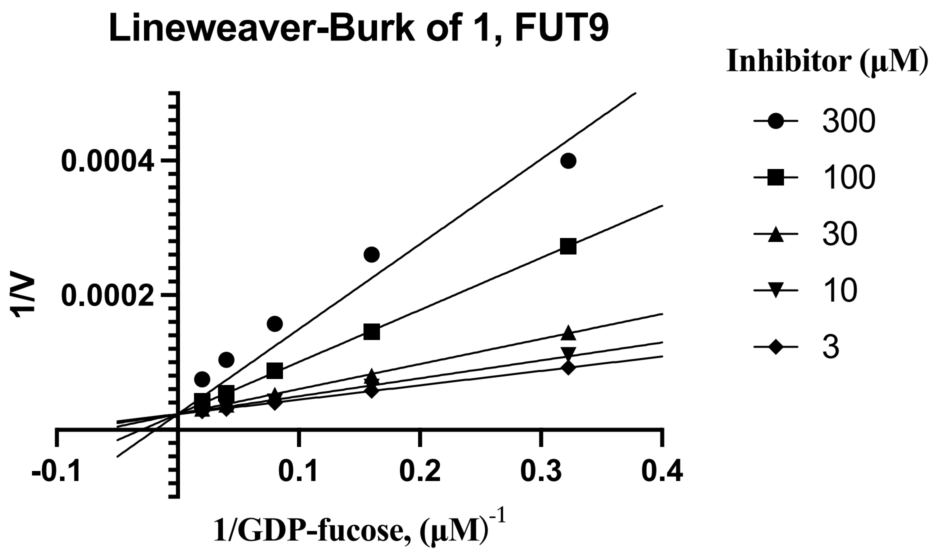
Lineweaver-Burk of 1, FUT9
Inhibitor (μM)
300
100
30
10
3
1/V
0.0004
0.0002
-0.1
0.1
0.2
0.3
0.4
1/GDP-fucose, (μM)-1

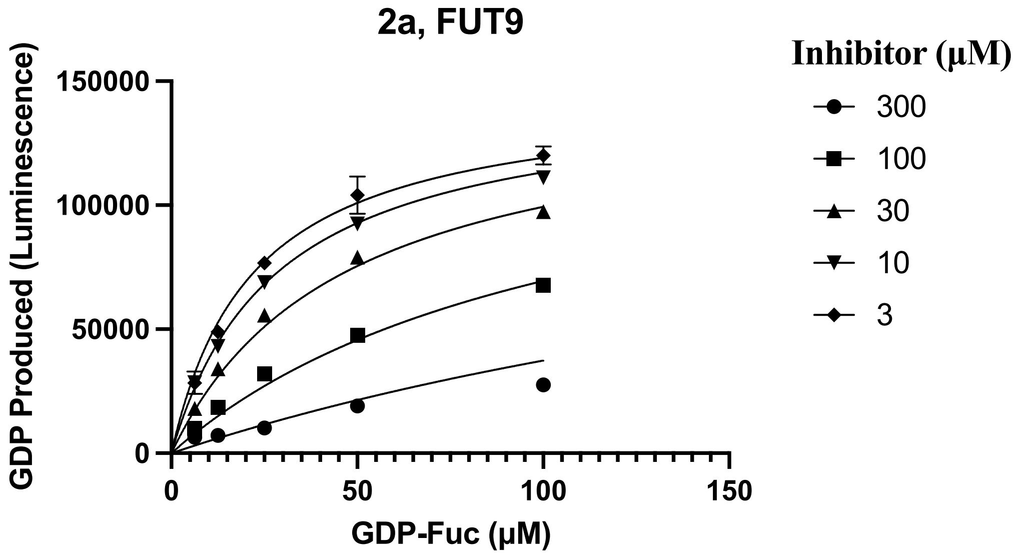
2a, FUT9
Inhibitor (μM)
300
100
30
10
3
GDP Produced (Luminescence)
150000
100000
50000
0
0
50
100
150
GDP-Fuc (μM)

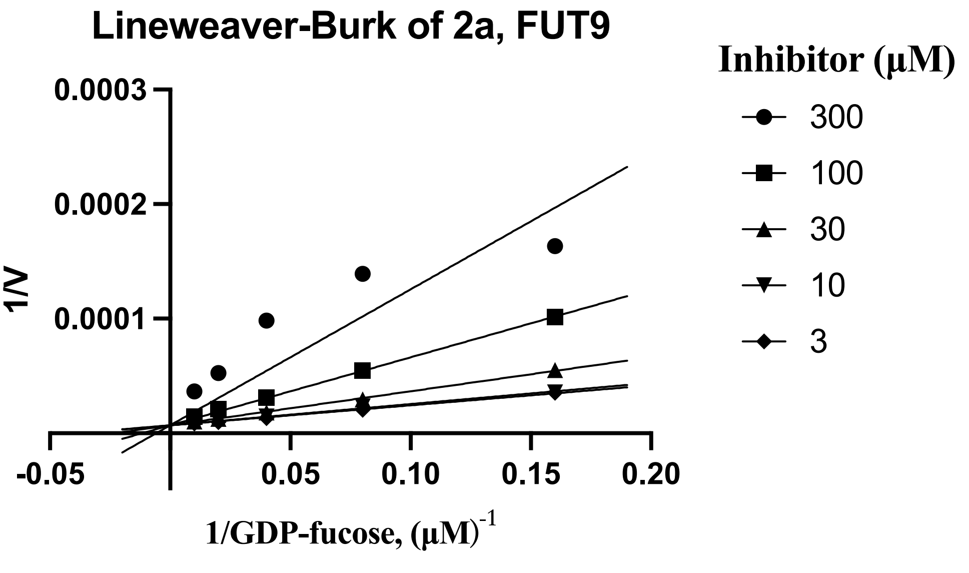
Lineweaver-Burk of 2a, FUT9
Inhibitor (μM)
300
100
30
10
3
1/V
0.0003
0.0002
0.0001
-0.05
0.05
0.10
0.15
0.20
1/GDP-fucose, (μM)-1

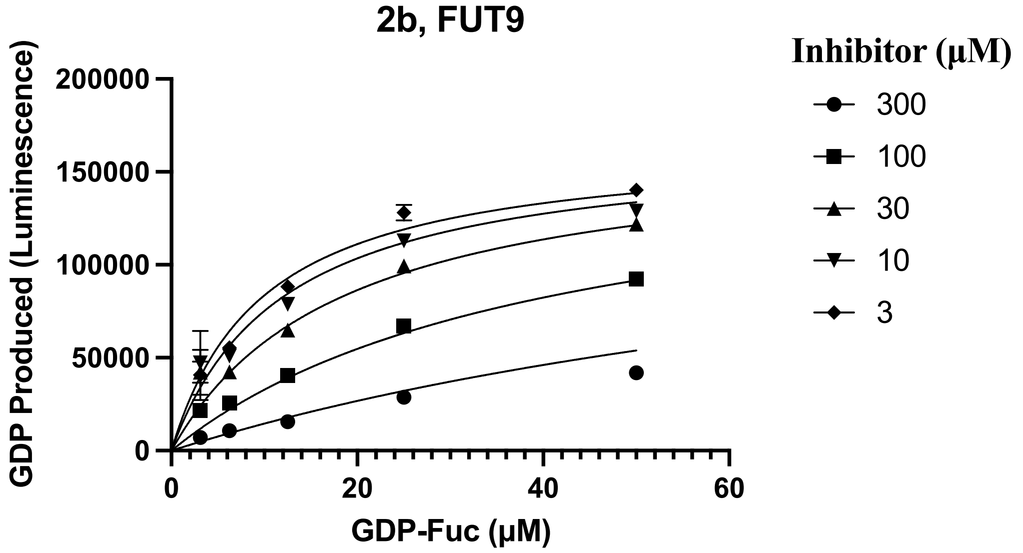
2b, FUT9
GDP Produced (Luminescence)
200000
150000
100000
50000
0
0
20
40
60
GDP-Fuc (µM)
Inhibitor (µM)
300
100
30
10
3

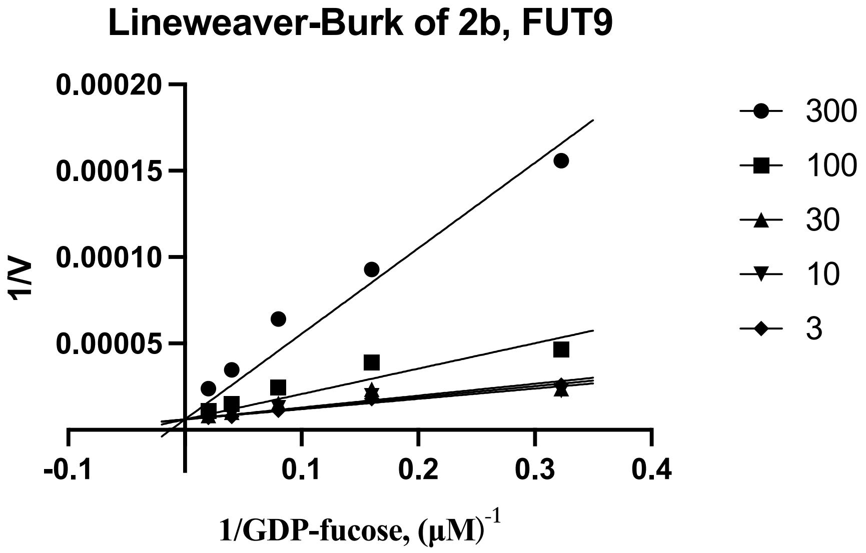
Lineweaver-Burk of 2b, FUT9
1/V
0.00020
0.00015
0.00010
0.00005
-0.1
0.1
0.2
0.3
0.4
1/GDP-fucose, (µM)-1
300
100
30
10
3

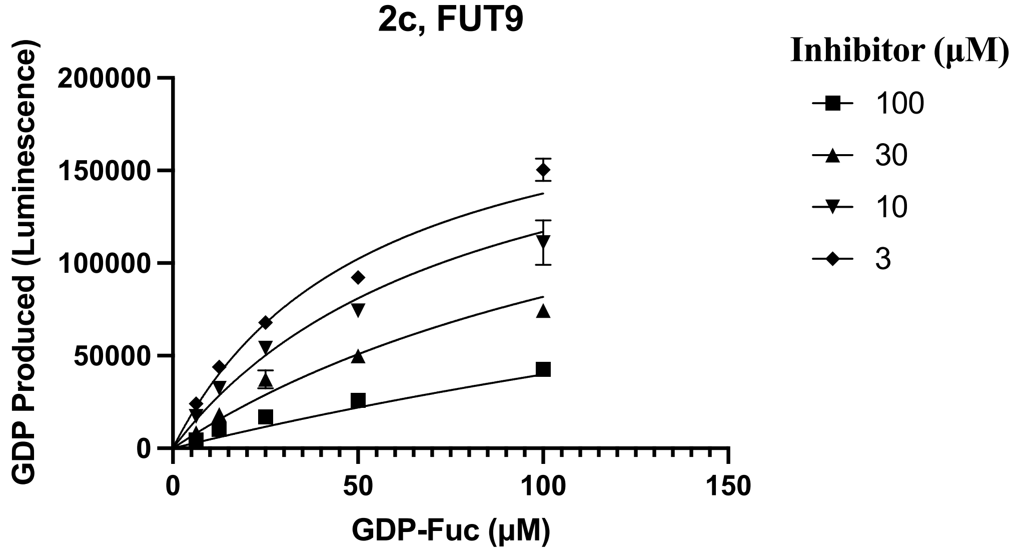
2c, FUT9
GDP Produced (Luminescence)
200000
150000
100000
50000
0
0
50
100
150
GDP-Fuc (µM)
Inhibitor (µM)
100
30
10
3

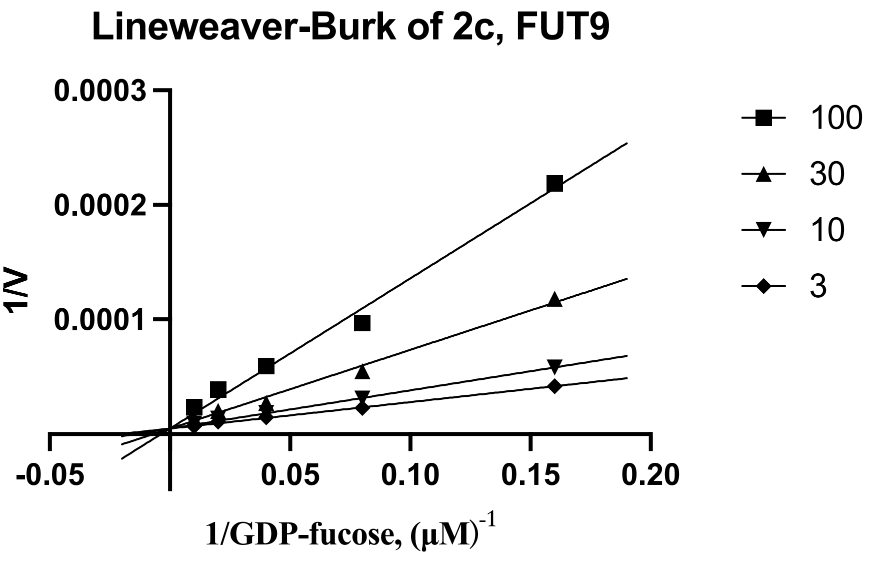
Lineweaver-Burk of 2c, FUT9
0.0003
0.0002
0.0001
1/V
-0.05
0.05
0.10
0.15
0.20
1/GDP-fucose, (μM)-1
100
30
10
3

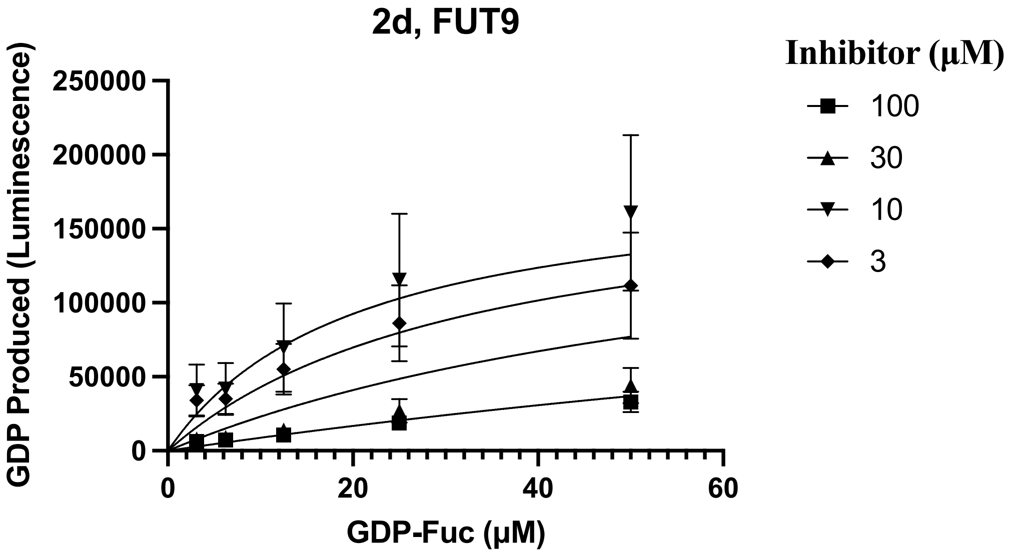
2d, FUT9
GDP Produced (Luminescence)
250000
200000
150000
100000
50000
0
0
20
40
60
GDP-Fuc (μM)
Inhibitor (μM)
100
30
10
3

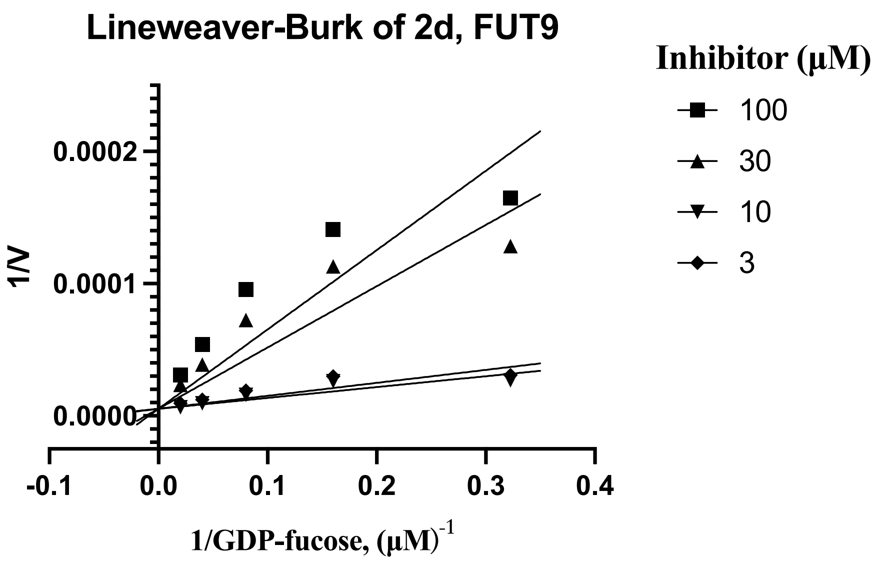
Lineweaver-Burk of 2d, FUT9
0.0002
0.0001
0.0000
1/V
-0.1
0.0
0.1
0.2
0.3
0.4
1/GDP-fucose, (μM)-1
Inhibitor (μM)
100
30
10
3

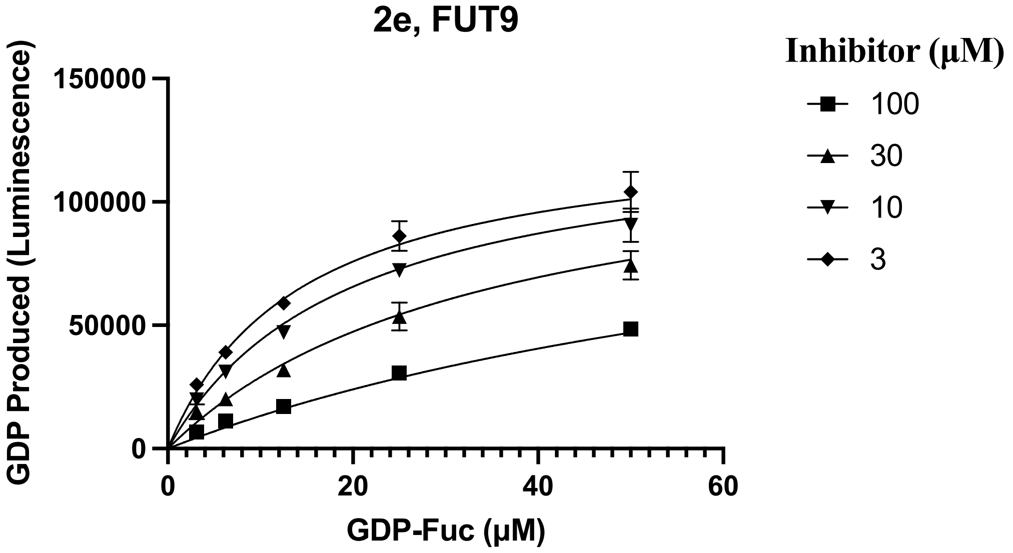
2e, FUT9
GDP Produced (Luminescence)
150000
100000
50000
0
0
20
40
60
GDP-Fuc (μM)
Inhibitor (μM)
100
30
10
3

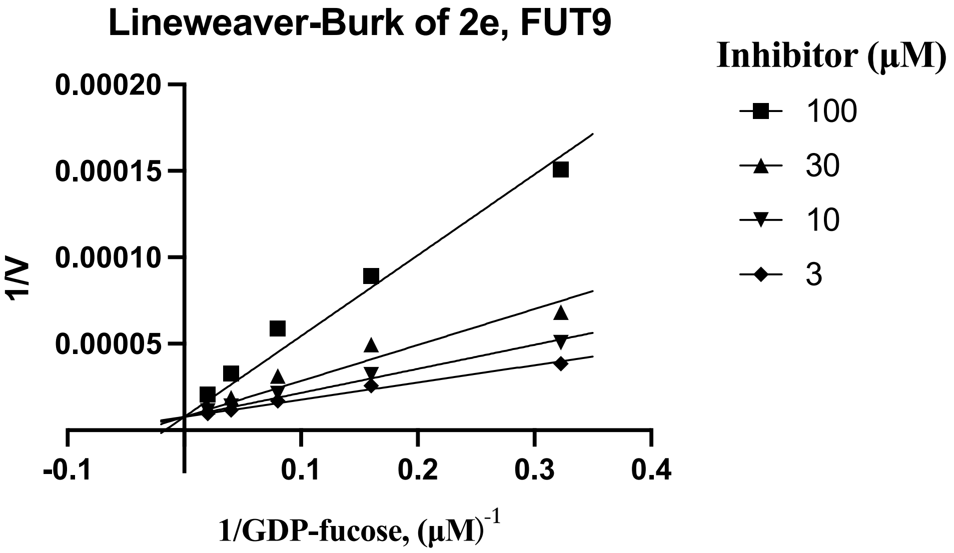
Lineweaver-Burk of 2e, FUT9
1/V
0.00020
0.00015
0.00010
0.00005
-0.1
0.1
0.2
0.3
0.4
1/GDP-fucose, (μM)-1
Inhibitor (μM)
100
30
10
3

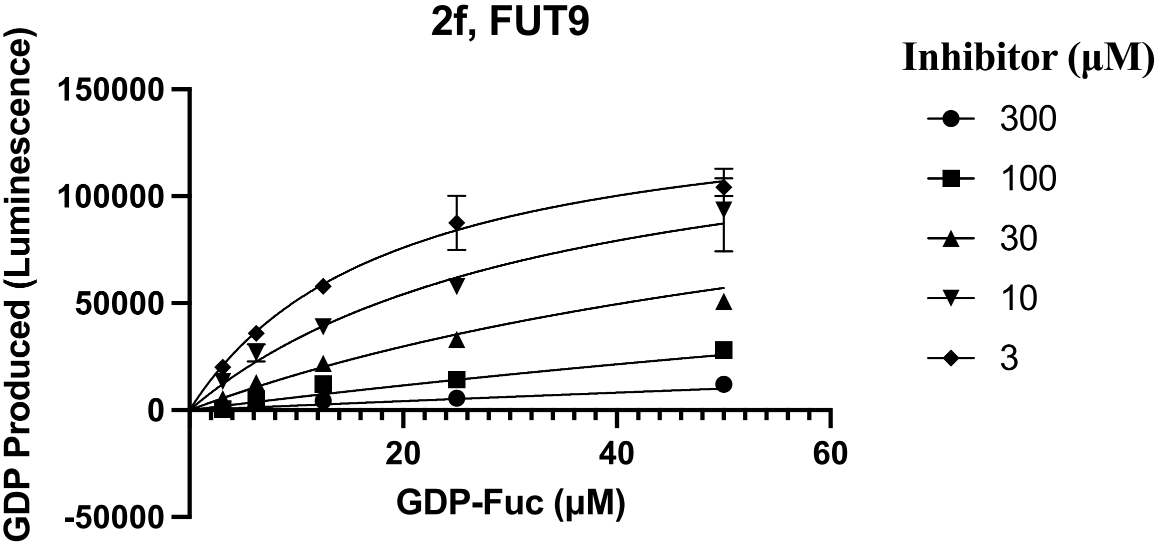
2f, FUT9
GDP Produced (Luminescence)
150000
100000
50000
0
-50000
20
40
60
GDP-Fuc (μM)
Inhibitor (μM)
300
100
30
10
3

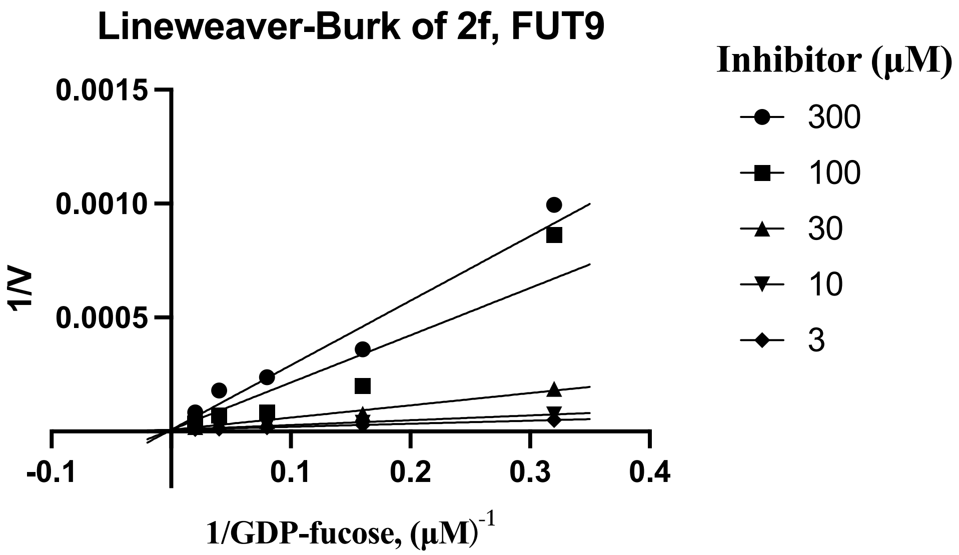

Lineweaver-Burk of 2f, FUT9
Inhibitor (µM)
300
100
30
10
3
0.0015
0.0010
0.0005
1/V
-0.1
0.1
0.2
0.3
0.4
1/GDP-fucose, (µM)-1


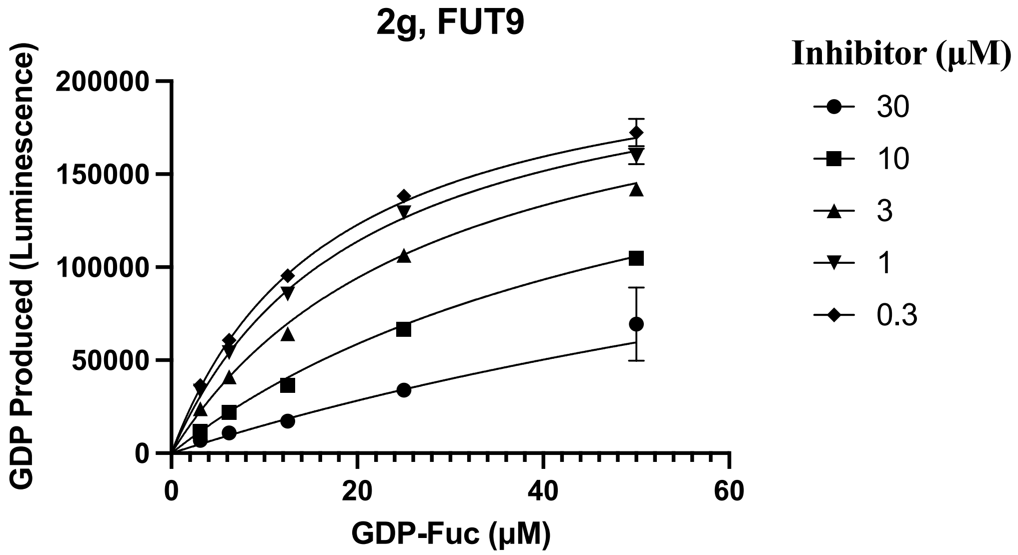

2g, FUT9
Inhibitor (µM)
30
10
3
1
0.3
200000
150000
100000
50000
0
GDP Produced (Luminescence)
0
20
40
60
GDP-Fuc (µM)


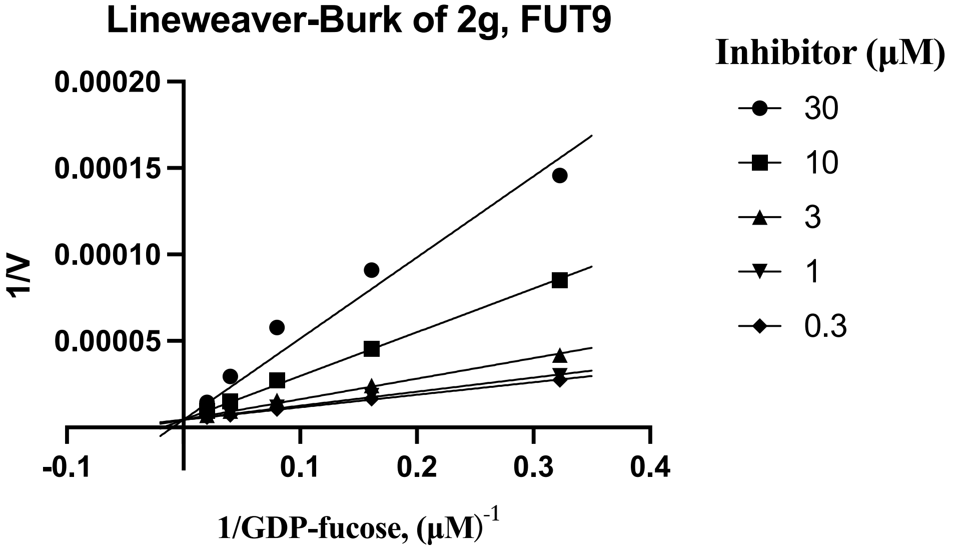

Lineweaver-Burk of 2g, FUT9
Inhibitor (µM)
30
10
3
1
0.3
0.00020
0.00015
0.00010
0.00005
1/V
-0.1
0.1
0.2
0.3
0.4
1/GDP-fucose, (µM)-1

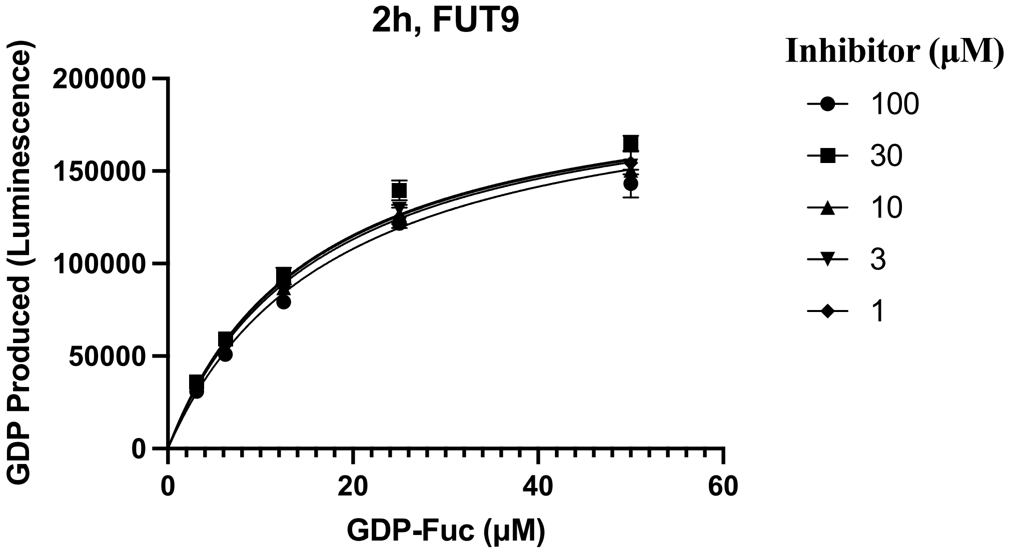

2h, FUT9
GDP Produced (Luminescence)
200000
150000
100000
50000
0
0
20
40
60
GDP-Fuc (μM)
Inhibitor (μM)
100
30
10
3
1


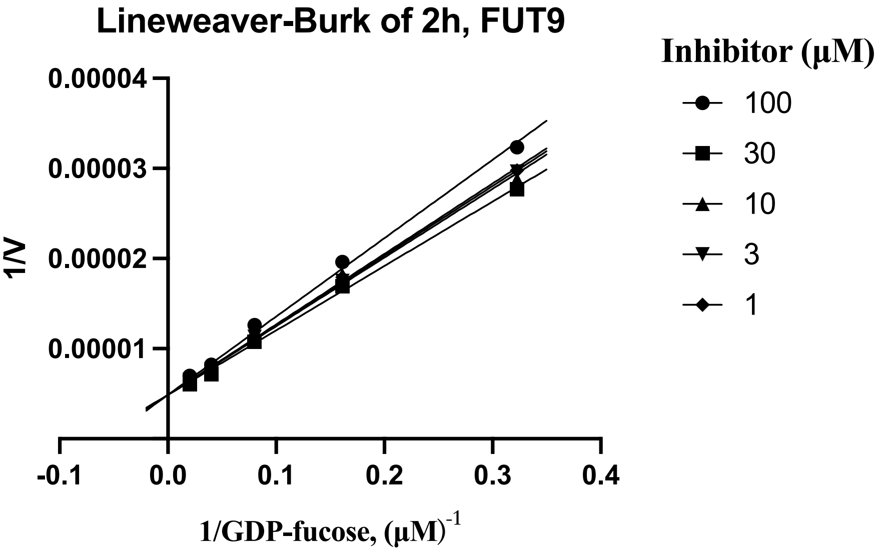

Lineweaver-Burk of 2h, FUT9
1/V
0.00004
0.00003
0.00002
0.00001
-0.1
0.0
0.1
0.2
0.3
0.4
1/GDP-fucose, (μM)-1
Inhibitor (μM)
100
30
10
3
1
